# Pervasive findings of directional selection realize the promise of ancient DNA to elucidate human adaptation

**DOI:** 10.1101/2024.09.14.613021

**Authors:** Ali Akbari, Alison R. Barton, Steven Gazal, Zheng Li, Mohammadreza Kariminejad, Annabel Perry, Yating Zeng, Alissa Mittnik, Nick Patterson, Matthew Mah, Xiang Zhou, Alkes L. Price, Eric S. Lander, Ron Pinhasi, Nadin Rohland, Swapan Mallick, David Reich

## Abstract

We present a method for detecting evidence of natural selection in ancient DNA time-series data that leverages an opportunity not utilized in previous scans: testing for a consistent trend in allele frequency change over time. By applying this to 8433 West Eurasians who lived over the past 14000 years and 6510 contemporary people, we find an order of magnitude more genome-wide significant signals than previous studies: 347 independent loci with >99% probability of selection. Previous work showed that classic hard sweeps driving advantageous mutations to fixation have been rare over the broad span of human evolution, but in the last ten millennia, many hundreds of alleles have been affected by strong directional selection. Discoveries include an increase from ∼0% to ∼20% in 4000 years for the major risk factor for celiac disease at *HLA-DQB1*; a rise from ∼0% to ∼8% in 6000 years of blood type B; and fluctuating selection at the *TYK2* tuberculosis risk allele rising from ∼2% to ∼9% from ∼5500 to ∼3000 years ago before dropping to ∼3%. We identify instances of coordinated selection on alleles affecting the same trait, with the polygenic score today predictive of body fat percentage decreasing by around a standard deviation over ten millennia, consistent with the “Thrifty Gene” hypothesis that a genetic predisposition to store energy during food scarcity became disadvantageous after farming. We also identify selection for combinations of alleles that are today associated with lighter skin color, lower risk for schizophrenia and bipolar disease, slower health decline, and increased measures related to cognitive performance (scores on intelligence tests, household income, and years of schooling). These traits are measured in modern industrialized societies, so what phenotypes were adaptive in the past is unclear. We estimate selection coefficients at 9.9 million variants, enabling study of how Darwinian forces couple to allelic effects and shape the genetic architecture of complex traits.

---

Ancient DNA data hold extraordinary promise for revealing adaptation, making it possible to track effects across time and to obtain direct measurements of selection coefficients^1–3^. Rather than being trapped in the present and studying the scars left by selection on the genomes of descendants—for example, searching for alleles too differentiated in frequency across populations^4,5^, or too common given their estimated age^6–8^, or gene genealogies distorted from the expectation for random drift^9^—ancient DNA makes it possible to test if frequencies of variants shifted more than could be expected by chance. Such data also make it easier to measure selection on variants not of recent mutational origin, which is challenging to detect using retrospective methods^10^. Most previous ancient DNA selection studies focused on two time-points—comparing allele frequencies in earlier to later groups—to search for alleles with extreme shifts compared to expectation from the genomic background^11,12^. We search for a consistently non-zero derivative over time, fully embracing the time-series nature of ancient DNA and using information differently affected by confounding factors.

Ancient DNA studies in West Eurasia^11–14^ (Europe and its neighbors in the Near East) have identified dozens of alleles influenced by selection^15,16^. But despite growth in the number of ancient individuals with data from zero before 2010 to more than 10,000 today, the number of genome-wide significant loci reported in a single study grew only mildly: from 12 in the first genome scan in 2015^11^, to 21 in a scan in 2024^14^. The small numbers raise the concern that the power of ancient DNA to detect selection might be reaching a plateau, and that this approach might not in fact be able to deliver broad insights into the nature of adaptation.

## Three innovations increase statistical power and minimize false signals of selection

Our improved yield of discoveries comes from more power (due to a qualitatively new method and larger sample size), and fewer artifactual signals (due to intensive data cleaning). First, we increased power by testing for a consistent trend in allele-frequency change over time. Most past studies of selection in ancient DNA dealt with the challenge posed by admixture by treating more recent populations as linear combinations of more ancient ones, then searching for alleles whose frequencies were outliers compared to what would be expected from this history. However, changes in frequency due to selection are often less than what can be expected from random genetic drift, and in this context, increasing sample size helps little. We employed a qualitatively different approach, using the genetic similarity of each individual to every other, and testing if the date when they lived provides additional predictive power for the allele frequencies of their population beyond what is expected from the empirical population structure. Our test is simple: at each variant we ask if hypothesizing a non-zero selection coefficient *s*—causing allele frequency to trend in the same direction over all times and places—predicts frequency differences across populations significantly better than empirically measured population structure alone (Methods).

Second, we increased power through a five-fold increase in sample size. We analyzed 8433 unrelated ancient individuals from the last 14000 years^17^ (Online Table 1). Data for 6686 come from enriching ancient DNA libraries for more than a million single nucleotide polymorphisms (SNPs) where median coverage is 3.6-fold (at least 0.44-fold); the remaining 1747 individuals are shotgun sequenced with median 1.6-fold coverage (at least 0.11-fold). For 3644 ancient individuals, sequences are previously reported. For 318, we increased data quality on previously reported individuals, largely from 300 newly reported shotgun genomes with median 4.9-fold coverage (40 at >17-fold coverage) (Online Table 2). For 4471 ancient individuals obtained by sequencing 5227 newly reported libraries (Online Table 3), we make data available for studies of selection with the support of sample custodians; archaeological contextual information will be provided in future publications which should be the references for analyses of their population history. We co-analyzed with 6510 modern people: 575 largely from the 1000 Genomes Project^18^, and 5935 from the UK Biobank^19^ (subsampled so their countries of origin were evenly spread over West Eurasia) (Extended Data Figure 1a,b).

**Table 1.**
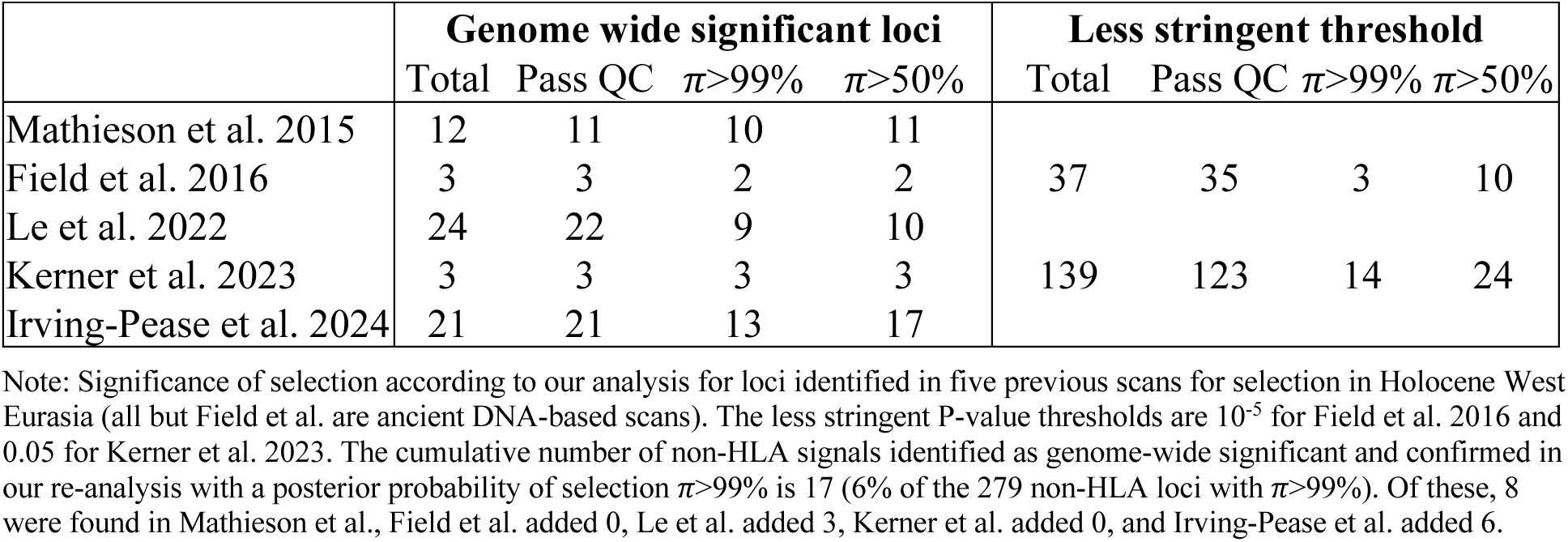
Re-evaluation of signals from five scans for selection in Holocene West Eurasia.

Third, we increased power and reduced false positives by data cleaning and imputation. We applied multiple data quality filters, including restricting to sites with similar frequencies in ancestry-matched modern and ancient people and giving consistent signals of selection with and without modern people (Supplementary Information section 1). We filled in missing data by leveraging known patterns of allelic correlation, using GLIMPSE^20^ to impute diploid genotypes and thereby increase allele counts at every locus (for imputation we used sequences aligning everywhere in the genome as we found that this greatly enhances information even for samples analyzed using in-solution enrichment). We analyzed 8,212,921 SNPs and 1,713,563 insertions/deletions (indels) imputed at high quality across chromosomes 1-22 in all individuals (we did not analyze the sex chromosomes or mitochondrial DNA).

## A test for directional selection with a negligible false-positive rate

For each SNP in the genome, we estimate a selection coefficient, which we found has a standard error typically around 0.1% for common variants (Figure 2, Extended Data Figure 1c). In theory, a valid test for selection should be *Z,* the number of standard errors this quantity is from zero, and we can use a normal distribution to identify scores that pass the standard threshold of genome-wide significance (P<5×10^−8^) for Genome-Wide Association Studies (GWAS). In practice, the median χ^2^ statistic (squared *Z-*score for number of standard errors *s* is from zero) is inflated by 5.26-fold relative to a χ^2^ distribution with one degree of freedom. In human genetics studies, such inflation can arise due to a variety of factors such as uncorrected population structure, and is often addressed by rescaling χ^2^ statistics by the median inflation across the genome^21,22^. However, such rescaling is only appropriate if the great majority of the genome is unaffected by the biological signal being studied. If, instead, a substantial fraction of the genome has real signal^21,23^, random locations in the genome will not provide an appropriate neutral baseline. In fact, we find evidence of exactly this problem: a large proportion of the genome in linkage disequilibrium (LD) with sites with evidence of directional selection (Supplementary Information section 2, Extended Data Figure 2b).

Instead, we calibrated our test by taking advantage of a striking finding about the connection between selection coefficients and associations to phenotypes in living people. We find that the proportion of SNPs showing significant association to a phenotype in a GWAS increases dramatically with our selection statistic Z, plateauing at around 3.9-times the rate of overlap for random SNPs (Figure 1a) (for this analysis we used 1,363,674 SNPs with a genome-wide significant association to at least one phenotype for 452 traits in the Pan-UK Biobank^24^). The increase is observed even after conditioning on minor allele frequency (MAF) to remove artifacts due to both selection and phenotypic associations being easier to detect for higher MAFs. The plateau occurs at the same place when we control for negative selection at linked loci (Extended Data Figure 3a). This is the pattern expected for a true threshold for genome-wide significance: if SNPs beyond this threshold reflect a combination of true signal and false discoveries, we would expect enrichment to continue beyond it. Because this threshold occurs at a value of Z=9.10—1.67-times larger than the standard threshold (5.45) for genome-wide significance for a normal distribution—we rescale the naïve score by this quantity to obtain an X-statistic (Z/1.67) whose significance threshold matches the standard threshold.

**Figure 1:**
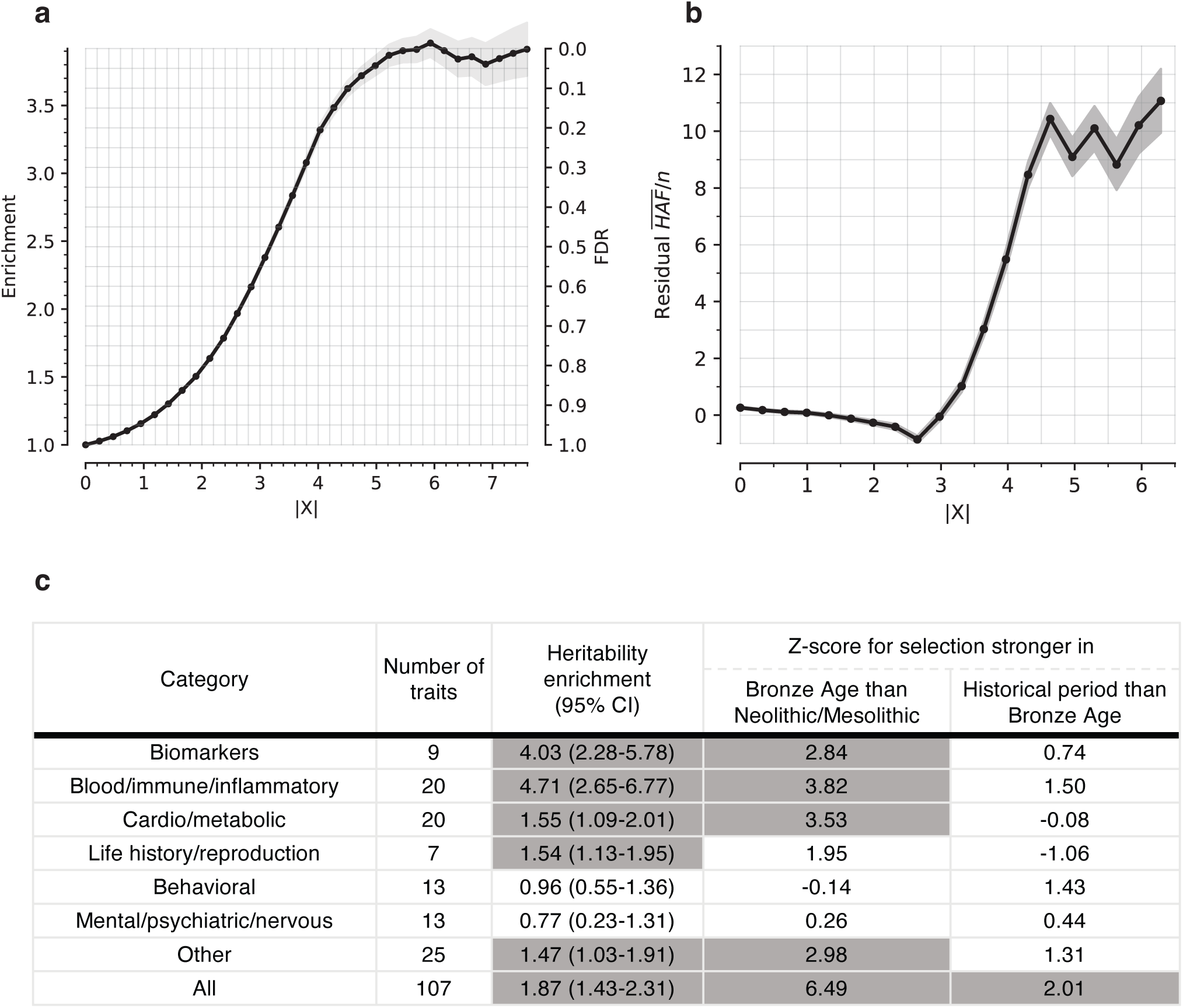
Multiple lines of evidence show we are detecting genuine signals of directional selection. **(a)** Proportion of SNPs significant in at least one of 452 Pan-UKBB GWAS, for SNPs with |X| above the value on the x-axis, and controlling for allele frequency. **(b)** Residual mean HAF-score [(HAF)/n], computed as observed minus expected value, with n the haploid sample size, from a linear regression correcting for background selection, and a window size of 200 kb. **(c)** The heritability enrichment column is a meta-analysis on heritability enrichment for annotations based on a binary selection annotation, with FDR either below 1% (=1) or above 1% (=0). Z-score for change in selection intensity over time is based on a meta-analysis of heritability enrichment comparing key cultural transitions: Mesolithic-Neolithic (MN) to Bronze Age (BA); and Bronze Age (BA) to Historical Era (HE). We annotate each SNP according to whether it is among the top 5% with the highest probability of a stronger magnitude of selection coefficient in one time transect vs. another.

To test whether we set an appropriate threshold for genome-wide significance with this procedure, we used orthogonal information: the sum of squared derived allele frequency in 200 kilobase haplotypes linked to each tested allele: the “Haplotype Allele Frequency” (HAF) score. Previous work^25,26^ showed that directional positive selection on derived alleles can increase HAF scores, while negative selection always decreases it, and we verified this by simulation (Extended Data Figure 3b, Supplementary Information section 3). After computing the residual HAF-score for each variant controlling for negative selection at linked loci^27,28^, we find it increases with the X-statistic and plateaus around 5.45, the standard threshold for genome-wide significance in GWAS (Figure 2b, Extended Data Figure 3c).

**Figure 2:**
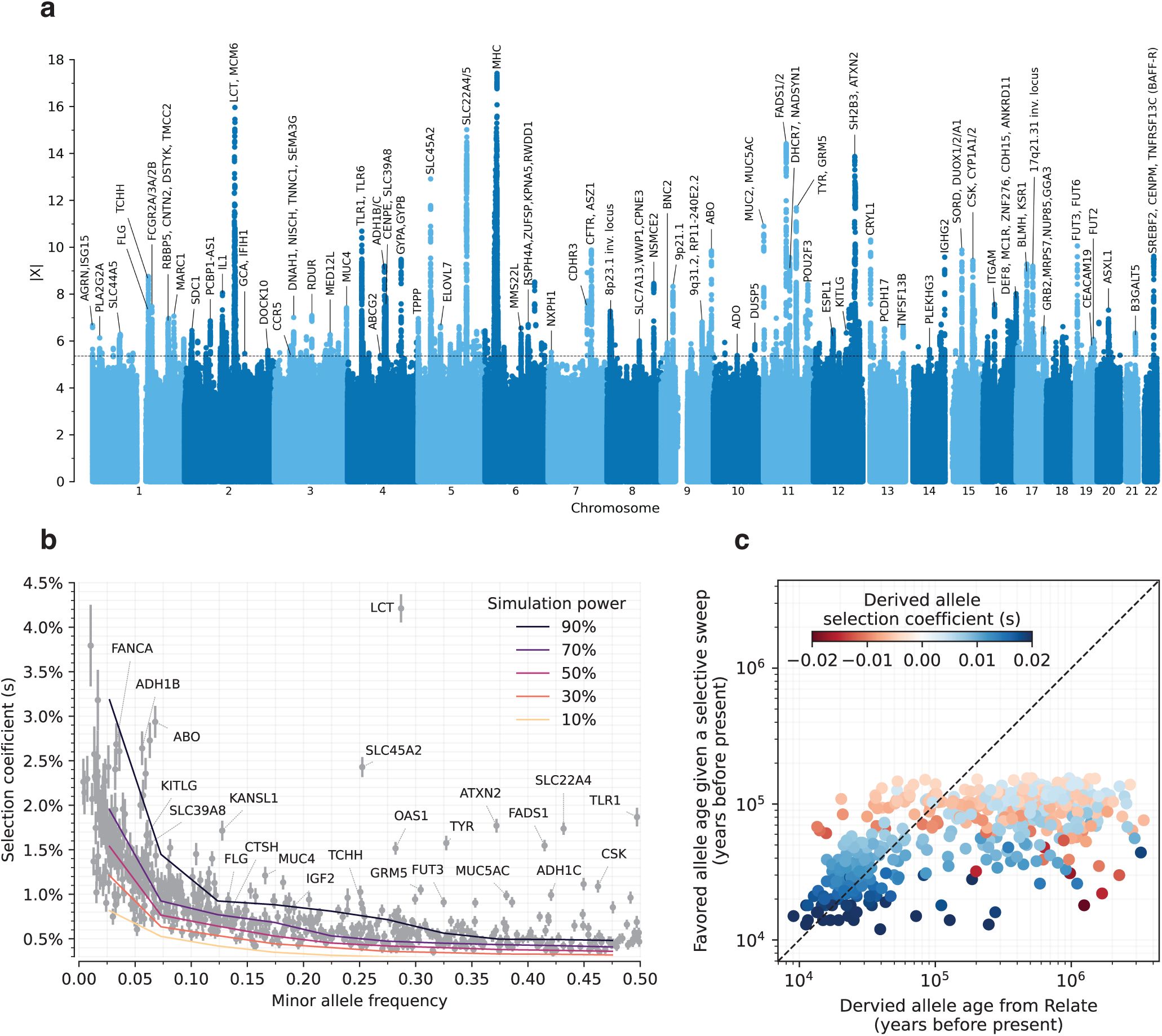
Genome scan for directional selection. **(a)** The x-axis is chromosomal position, and the y-axis the selection signal for each variant. The dotted line indicates our genome-wide significance threshold of |X|=5.45. For clarity, only select loci are annotated. **(b)** Selection coefficient (*s*) estimated from our scan plotted against minor allele frequency of tagging SNPs at independent loci with FDR<5%. Overlaid grids are simulation-based power estimates (90%, 70%, 50%, 30%, and 10% probability of detection). **(c)** The estimated age of the favored allele in a selective sweep versus the date of origin of the mutation is inferred from RELATE^9^, for tagging SNPs with FDR<5% at independent loci. The age of the sweep is defined as the time in the past when the frequency of the favored allele is expected to have been 0.0001 given the present-day frequency in 1000 Genomes Project European populations and assuming the selection coefficient has been constant over time.

To translate X to a posterior probability of selection *π*, we used a False Discovery Rate (FDR) approach (Supplementary Information section 2). We fit a smooth function to the enrichment curve for GWAS signals and estimate that at X-statistic magnitudes greater than our threshold for genome-wide significance of 5.45, *π*>99% (Extended Data Table 1).

We confirmed that our X-statistics are detecting biologically meaningful patterns by showing that signals of selection are unusually associated with specific classes of traits^29^. In particular, we find enrichment for SNPs contributing to blood-immune-inflammatory traits (95% confidence interval (CI) 2.6-6.8)^12,13^, compared to random SNPs with matched characteristics defining the baseline of 1-fold. In contrast, for mental-psychiatric-nervous and behavioral traits, we do not see enrichment (95% CI of 0.2-1.3 and 0.5-1.4) (Figure 1c, Extended Data Figure 4a). These patterns cannot be explained by differences in allele frequencies or purifying selection since we control for these factors. The intensity of selection on blood-immune-inflammatory and cardio-metabolic traits increased in the Bronze Age relative to the pre-farming period (Figure 1c, Extended Data Figure 4b), which may reflect adaptation to new diets, higher population densities, or living closer to domesticated animals.

## Hundreds of loci affected by directional natural selection

We identified 347 independent loci (279 excluding the HLA region) with |X|>5.45, corresponding to a π>99% probability of selection (Figure 2a). To produce this list, we identified the strongest signal in the genome and considered all SNPs in LD with it in modern Europeans from the 1000 Genomes Project (r^2^>0.05) to potentially reflect the same signal. We then found the second-strongest signal excluding these positions, and so on until no more SNPs pass this threshold (Extended Data Figure 2b). We provide visualizations of the trajectories for these 347 loci (Supplementary Information section 5) and summary statistics for 9.9 million imputed variants (Online Table 4), which can be cross-referenced with GWAS and viewed along with their frequency trajectories at the AGES internet browser https://reich-ages.rc.hms.harvard.edu.

The actual number of loci under selection is likely to be much larger. Using a threshold of |X|=3.16, which corresponding to FDR=50%, we identify 10361 non-HLA loci, implying >5000 independent episodes of selection. Moreover, our approach to identifying distinct loci is conservative, because genuinely selected alleles in LD with nearby stronger ones will be missed. Down-sampling analyses show that further increases in sample size are expected to increase the number of loci further, with people living >8000 years ago providing the most added power (Extended Data Figure 1d,e).

To obtain insight into the phenotypic targets of the loci under natural selection, we take advantage of the fact that a high proportion (82%) of the variants with genome-wide evidence of selection are independently associated to a phenotype in at least one Pan-UK Biobank GWAS in living people. However, biological interpretation is complicated since the allele that was the target of selection may differ from the tag SNP we are using to represent the locus (and may even be in a neighboring gene), because some alleles affect multiple phenotypes, or because the relevant modern trait may not be measured in one of the GWAS we are analyzing, or because the phenotype in modern societies may not have existed in the ancient ones where selection acted. The median selection magnitude |*s*| at the tag SNPs is 0.8% (range 0.4-4.2%), and the median minor allele frequency (MAF) is 19%. Standard errors in our estimates of *|s*| for common alleles are ∼0.1%, and we have limited power to detect selection coefficients of magnitude <.5% (Figure 2b) (Extended Data Figure 1c).

We compared our results to those of five previous selection scans in Holocene West Eurasia (four based on ancient DNA) (Table 1). Of 39 unique non-HLA loci that met the formal threshold for genome-wide significance in at least one of the previous studies, 17 pass our π>0.99 threshold. The other 22 do not replicate, in most cases due to what appears to be incompletely controlled population structure driven by mixtures of populations with different allele frequencies before they came together (Supplementary Information section 5). (Two of the previous studies also reported additional candidate loci that did not pass the author’s own genome-wide significance threshold, and we found that only ∼10% of these replicated, suggesting most are false-positives^8,13^.)

We present a gallery of 36 single-allele trajectories of particular interest (Figure 3) as well as estimates of how their selection coefficients changed over time (Extended Data Figure 5). These loci are not necessarily those with the largest X-scores, but are highlighted as they address long-standing debates. They include 24 passing the π>99% threshold, 7 with probable evidence of selection (64%<π<98%), and 5 with surprising negative findings.

**Figure 3:**
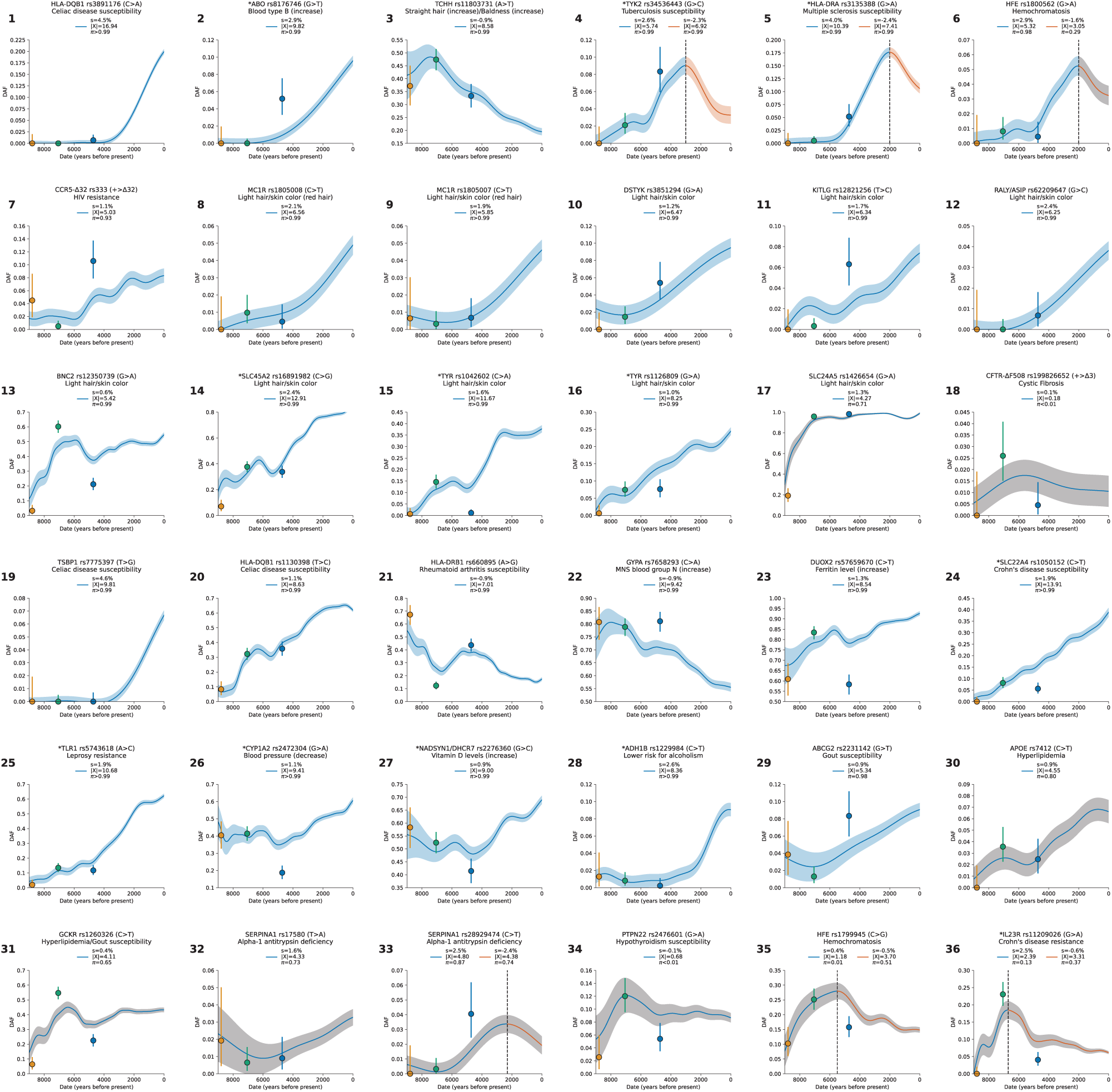
Gallery of notable single-locus selection trajectories. Each panel displays the derived allele frequency trajectory over time for a variant (uncorrected for structure), along with selection coefficient (*s*), selection statistic (X), and posterior probability of selection (*π*). Circles represent frequencies in Western Hunter-Gatherers (orange), Early European Farmers (green), and Steppe Pastoralists (blue). The highlighted loci are not necessarily those with the strongest signals, and even include negative results. We highlight them here because of their biological interest and because they speak to long-standing debates. For Panels 4, 5, 6, 33, 35, and 36 separate analyses are shown for transects before and after a manually selected peak (marked by a black line), with 200-year overlap. In cases where *π*>90%, the confidence interval is shaded blue (or blue for before and red for after the split); otherwise, the shading is gray. Variants reported in other ancient DNA studies are marked with an asterisk.

### *HLB-DQB1:* Selection in favor of the major risk factor for celiac disease (panel 1)

At the HLA region of chromosome 6, densely packed genes play key roles in microbe recognition. rs3891176 (C>A, meaning that the ancestral allele is C and the newly arising mutation is A) is an excellent tag for *HLA-DQB1*02/DQ2*, with individuals carrying two A alleles having a 19-fold higher susceptibility for celiac disease or gluten sensitivity (Extended Data Figure 6a,b). The A allele has a selection coefficient of *s*=4.5% (π>99%), rising from ∼0% to ∼20% in the last 4000 years. These findings speak to the debate about the relationship between agriculture and celiac disease^30–32^, as the results suggest that the pathogenic exposures that drove its rise were not a phenomenon only or largely of the Neolithic.

### *ABO*: Positive selection for B at the expense of the A allele (panel 2)

*ABO* modifies oligosaccharides in glycoproteins on the surface of red blood cells and codes for the A, B, and null (O) alleles that interact in different ways with pathogens^33,34^. We show that the B allele rose from ∼0% to ∼10% over the last ∼6000 years (*s*=2.9%, π>99%), and was matched by a concomitant decrease in A frequency. The A and B alleles are associated with opposite effects on many phenotypes, suggesting that with changing lifestyles and pathogenic exposures, the optimal balance of these alleles changed (Extended Data Figure 6c,d).

### *TCHH*: Selection for an allele that reduced male pattern baldness (panel 3)

An allele at missense SNP rs11803731 (A>T) in *TCHH* is a strong predictor of straight hair and male pattern baldness in Europeans. The derived allele T is rare in African and East Asian populations, and has been hypothesized to have been positively selected, analogous to the straight-hair *EDAR* allele in East Asians^35^. We observe an opposite trend: the derived allele was negatively selected (*s* = −0.9%, π>99%), decreasing from ∼50% to ∼20% in the past 7000 years. This implies a 1.8% decrease in predisposition to baldness over this period.

### *TYK2.* Reversal of selection at a major factor for tuberculosis (panel 4)

Individuals carrying two copies of the rs34536443 G>C allele have >80% prevalence of clinically significant tuberculosis^36^. Previous work^37^ found evidence of negative selection on the C allele and hypothesized it was associated with the time tuberculosis began to be endemic in Europe. We confirm a drop in frequency from ∼9% to ∼3% in the last ∼3000 years (*s* = −2.3%, π>99%), but also identify positive selection from ∼5500 to ∼3000 years ago, from around ∼2% to ∼9% (*s*=2.6%, π>99%). This may reflect changing endemicity of different pathogens over time.

### *HLA-DRB1.* Elevated MS risk in north Europe is not due to selection on the steppe (panel 5)

A previous study^38^ discovered positive selection at the rs3135388 G>A tag SNP for the HLA-DRB1*15:01 risk factor for multiple sclerosis (MS)^39^. Because selection was already occurring in Yamnaya steppe pastoralists, and Yamnaya ancestry is most common in north Europeans today, the authors argued that the genetically higher risk for MS in north than in south Europeans was driven by selection on the steppe. We confirm positive selection at this allele, rising from ∼0% to ∼18% between ∼6000 and ∼2000 years ago (*s*=4.0%, π>99%). However, we also document three features of the selection history missed by previous work (Supplementary Information section 6), and which together show that the primary driver of the north/south differential in this allele’s frequency was not selection on the steppe. First, selection did not begin on the steppe^38^; it was occurring earlier south of the Caucasus mountains in people without steppe ancestry. Second, after Yamnaya ancestry spread west, selection was stronger in north Europe at *s* = 14.5 ± 3.4% than in southwest Europe at *s* = 5.1 ± 2.5% (measured >3500 BP). Third we document negative selection in the last ∼2000 years missed by previous work (*s* = −2.4%, π>99%), likely reflecting new pathogen exposures.

### *HFE*: Reversal of selection at the major risk factor for hemochromatosis (panel 6)

The rs1800562 (G>A) allele predicts pathogenic iron buildup in cells in individuals with two copies, and we find evidence of positive selection from ∼5000 to ∼2000 years ago, rising from ∼1% to ∼5% (*s* =2.9%, π=98%), then dropping to ∼3% today. This reversal is not genome-wide significant (*s* = −1.6%, π=29%), but is compelling as a single hypothesis test at a locus with long-standing speculation regarding selection. It was hypothesized that the causal allele protected against *Yersinia pestis* (the agent of Black Death)^40^, but this is unlikely as its frequency was decreasing by the time of the Justinianic and Medieval pandemics^41,42^.

### *CCR5-*Δ*32*: Positive selection at an allele conferring immunity to HIV-1 infection (panel 7)

The *CCR5-*Δ*32* allele confers complete resistance to HIV-1 infection in people who carry two copies^43–45^. An initial study dated the rise of this allele to medieval times and hypothesized it may have been selected for resistance to Black Death^46^, but improved genetic maps revised its date to >5000 years ago and the signal became non-significant^47,48^. We find that the allele was probably positively selected ∼6000 to ∼2000 years ago, increasing from ∼2% to ∼8% (*s* =1.1%, π=93%). This is too early to be explained by the medieval pandemic, but ancient pathogen studies show *Yersinia* was endemic in West Eurasia for the last ∼5000 years^49–51^, resurrecting the possibility that it was the cause, although other pathogens are possible.

### Selection for light skin at 10 loci (panels 8-17)

We find nine loci with genome-wide signals of selection for light skin, one probable signal, and no loci showing selection for dark skin.

### *CFTR*: No evidence of selection for the major cystic fibrosis risk allele ΔF508 (panel 18)

The major risk allele for this recessive disease in Europeans^52,53^ has been hypothesized to be an example of heterozygote advantage due to advantages in carriers such as resistance to cholera^54^. However, we find no evidence of selection (π<1%), with the earliest direct observation at ∼2200 years ago in Great Britain and the earliest imputed one ∼10100 years ago in Anatolia. It seems unlikely that cholera was endemic in West Eurasia this long; another explanation is needed for the persistence of this allele which in two copies also causes male infertility.

Fourteen other selection discoveries are highlighted in panels 19-32 of Figure 3. Most pass our threshold for genome-wide significance at π>99%: *TSBP1* (Celiac disease, *s*=4.6%); *HLA-DQB1* (Celiac disease, *s* = 1.1%); *HLA-DRB1* (Rheumatoid arthritis, *s* = −0.9%); *GYPA* (increases MNS blood group N, *s* = −0.9%); *DUOX2* (increases Ferritin level, *s*=1.3%); *SLC22A4* (Crohn’s disease, *s*=1.9%); *TLR1* (Leprosy resistance, *s*=1.9%); *CYP1A2* (decreases blood pressure, *s*=1.1%); *NADSYN1/ DHCR7* (increases vitamin D, *s*=0.9%); and *ADH1B* (lower risk for alcoholism, *s*=2.6%). Four more signals are probable: *ABCG2* (gout, *s*=0.9%, π=98%); *APOE* (hyperlipidemia, *s*=0.9%, π=80%); *GCKR* (hyperlipidemia/gout, *s*=0.4%, π=65%), and *SERPINA1* (alpha-1 antitrypsin deficiency, *s*=1.6%, π=73%).

Panels 33-36 highlight four negative signals at loci previously hypothesized to have been selected: a second locus at *SERPINA1* (alpha-1 antitrypsin deficiency); *PTPN22* (hypothyroidism); a second locus at *HFE* (hemochromatosis); and *IL23R* (Crohn’s disease).

## Directional selection shaped dozens of complex traits

Having examined selection on individual loci, we searched for evidence that groups of alleles with similar influence on traits today trended in the same direction in the past, as would be expected if a phenotype with a similar genetic underpinning was the target of selection. To study this, we leveraged GWAS data for 452 mostly quantitative traits in the Pan-UK Biobank, and 107 dichotomous traits from studies especially of common disease^55^ (Online Table 5). How phenotypes manifest in modern societies may be very different from how they manifested in past populations living in different environments with different lifestyles, so any signals discovered by this approach should not be interpreted as evidence for selection on the exact phenotype being tested.

We used three statistics to test for coordinated selection on alleles affecting the same trait. First, we computed a polygenic score (PGS) for each GWAS: a linear combination of allelic values, weighted by estimated effect size. We evaluated whether the change in PGS over time γ (which we scaled so one-unit corresponds to a one standard deviation change over ten millennia) is more than could be expected by genetic drift alone. To test if the observed deviation is significant, we repeated the test 100 times with randomly flipped signs of GWAS effect sizes, to correct for LD among neighboring sites. As a second test, we repeated the procedure without using the magnitudes of the GWAS effects, and instead only the sign, generating a statistic *γ*_sign_ that may be less affected by concerns about transferability of PGS across groups^12,56–58^. Third, we performed a SNP-by-SNP comparison for each trait, using cross-trait LD Score Regression (LDSC) to estimate genetic correlation (*r_s_*) between selection summary statistics and GWAS summary statistics^59^, accounting for non-independence of SNPs. We computed a standard error from a Block Jackknife to test if this correlation is significantly different from zero. We find high Pearson’s correlation for all three tests (75-91%; Extended Data Figure 7).

For 31 of the 559 traits examined, we were able to carry out a further test of robustness by leveraging data from East Asian GWAS. Early studies claimed selection for greater height in north than in south Europeans, but this was later shown to be a false-positive due to uncorrected population structure in GWAS (ancestry differentially carried by north and south Europeans) that is correlated to structure in the groups tested for selection^60,61^. However, population structure in East Asia should be almost completely uncorrelated to that in the ancient West Eurasians, so it is difficult to see how validation by this test could be anything but a real signal of selection^12,56^.

We identified 12 traits with significant signals from all three tests after correction for number of traits tested (p<10^−4^, correcting for ∼500 hypotheses) (Figure 4, Extended Data Figure 8).

**Figure 4:**
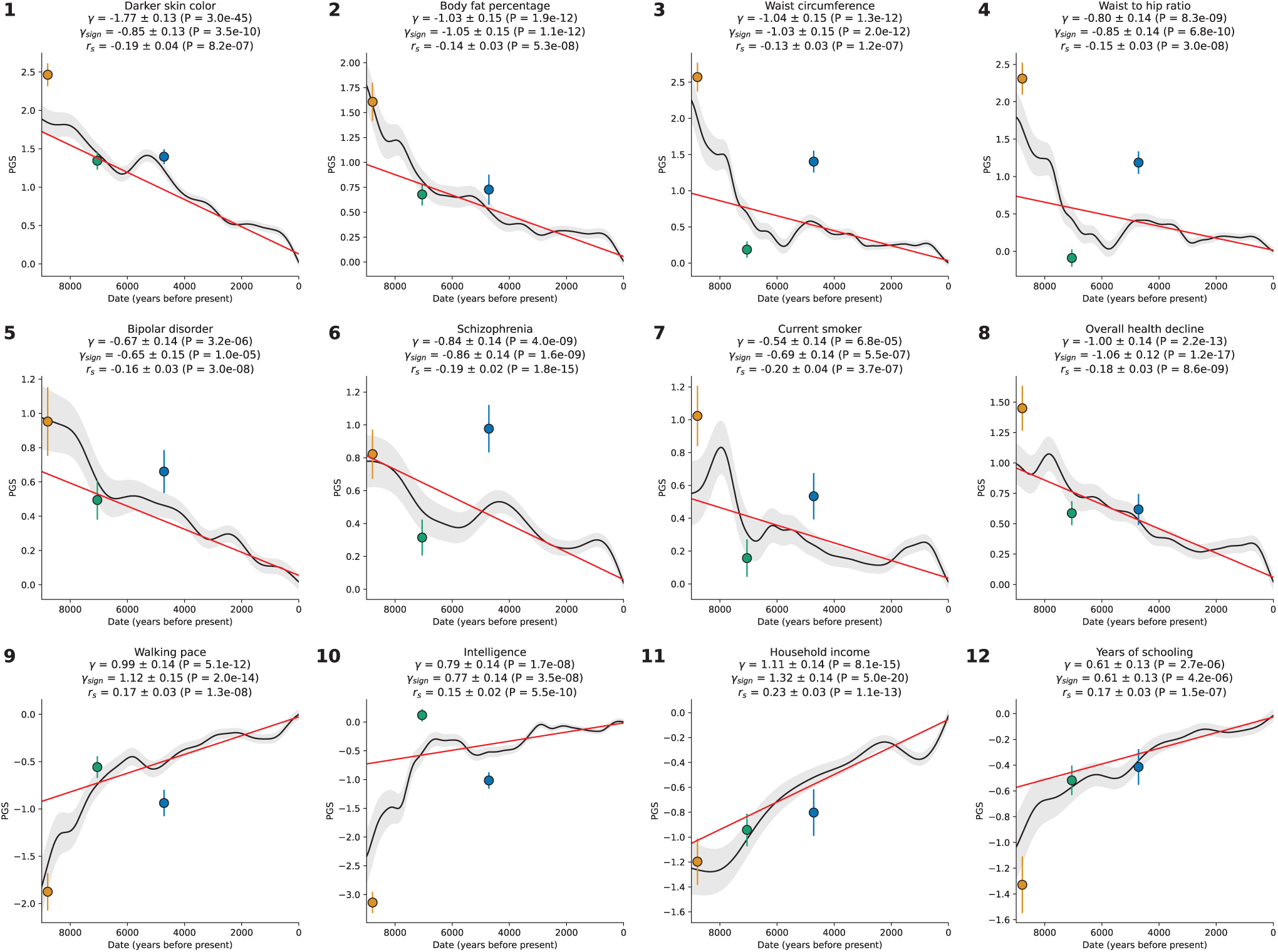
Coordinated selection on alleles affecting same traits (polygenic adaptation). The polygenic score of Western Eurasians over 14000 years in black, with 95% confidence interval in gray. Red represents the linear mixed model regression, adjusted for population structure, with slope *γ*. Three tests of polygenic selection—*γ*, *γ*_sign_, and *r_s_*—are all significant for each of these twelve traits, with the relevant statistics at the top of each panel.

One of the strongest signals is an increase over time in the PGS for light skin pigmentation (*γ* =1.77 **±** 0.13 standard deviations increase in mean PGS in ten millennia, P=3.0×10^−45^; Figure 4, Extended Data Figure 8). This plausibly reflects selection for increased synthesis of vitamin D in regions of low sunlight in farmers with little of it in their diets. Previous ancient DNA analysis^57^ found most of the phenotypic shift is driven by a few loci. Our results agree: 50% of the shift is due to *SLC45A2* alone, and 69% by the top 7 loci (Extended Data Figure 9). However, the selection was extraordinarily polygenic as we need to drop the top 104 loci before the signal disappears (Extended Data Figure 10). A model in which selection for pigmentation impacted all variants in proportion to their effect size fits the data (P=0.10).

Type 2 diabetes risk factors give compelling signals of negative selection. Thus, we observe negative selection on combinations of alleles that today increase body fat percentage (*γ* = −1.03 **±** 0.15), waist circumference (*γ* = −1.04 **±** 0.15), and waist-to-hip ratio (*γ* = −0.80 **±** 0.14), supporting the “Thrifty Gene” hypothesis that a genetic adaptation to store fat in times of plenty, became deleterious after the transition to food-production (Figure 4). For type 2 diabetes itself, the signal (*γ* = −0.40 **±** 0.11) just misses the multiple hypothesis-testing corrected threshold, but the other two exceed it (*γ*_sign_ = −0.51 **±** 0.12; *r_s_* = −0.16 ± 0.04).

We find signals of negative polygenic selection against alleles associated today with psychoses such as bipolar disorder (*γ* = −0.67 **±** 0.14) and schzophrenia (*γ* = −0.84 **±** 0.14) (Figure 4). Superficially this is in tension with the finding that variants with genome-wide significant of selection are not enriched for variants known to modulate psychiatric traits (Figure 2b). However, for variants with weaker signals, we do observe heritability enrichment (Extended Data Figure 4a). Brain traits have qualitatively different genetic architectures than blood-immune-inflammatory ones, with a higher total proportion of sites modulating them and smaller effect sizes on average per allele^62^. If brain traits tend to be associated with many alleles with small selection coefficients, this may reduce heritability enrichment at precisely the loci in the genome giving the strongest selection signals. These traits too are extraordinarily polygenic: we have to drop 740 loci for bipolar disorder and 726 loci for schizophrenia for the signals to become non-significant (Extended Data Figure 10).

We observe signals of selection for combinations of alleles that at today associated with healthy lifestyles into old age. This includes selection for alleles that promote faster walking pace (*γ* = 0.99 **±** 0.14), against alleles that today are associated with smoking (*γ* = −0.54 **±** 0.14), and against alleles contributing to overall health decline (*γ* =-1.00 **±** 0.14).

We finally observe signals of selection for combinations of alleles that today predict three correlated behavioral traits: scores on intelligence tests (increasing 0.79 ± 0.14), household income (increasing 1.11 ± 0.14), and years of schooling (increasing 0.61 ± 0.13). These signals are all highly polygenic, and we have to drop 463 to 1109 loci for the signals to become nonsignificant (Extended Data Figure 10). We also tested for a correlation of East Asian GWAS effect size measurements to West Eurasian selection. We observe a significant correlation for γ_sign_ (P=3.8×10^−6^) and *r_s_* (P=1.9×10^−10^) (Extended Data Figure 11), which is very difficult to explain as an artifact of population structure.

There are caveats when interpreting signals of polygenic adaptation, especially for the three genetically correlated traits of scores on intelligence tests, household income, and years of schooling. These traits—for which there is evidence of significant negative selection in the last century, for example in Iceland, in the opposite direction to the long-term increase we detect^63–65^—are only relevant to modern societies, and would have been unmeasurable in the preliterate societies over the vast majority of the period during which selection acted. The difficulty of interpretation is enhanced by the fact that the alleles driving down the frequency of type 2 diabetes-related traits, are highly correlated to those contributing to the increased scores for years of school, household income, and intelligence tests (Extended Data Figure 12). We could not gain meaningful additional insight into the selection mechanism by repeating analyses in family-based GWAS^66^ due to the limited sample sizes in these studies (Extended Data Figure 13).

## Discussion

Previous work has shown that classic selective sweeps driving alleles to fixation have been rare over the broad span of human evolution^67,68^. Thus, we were surprised that over the last 14,000 years in West Eurasia there have been many hundreds of instances of directional selection with coefficients on the order of 0.5% or more (Figure 2b). This is large enough that if a similarly dense landscape of directionally selected variants had existed tens of thousands of years ago, and if the selection coefficients had been constant since then, we would expect many fixed differences across populations, despite the fact that previous studies have shown there are only a handful—hardly more than would be expected based on random drift^68^.

The simplest way to resolve this paradox is to recognize that selection coefficients are unlikely to have been constant over time, even though we make this simplifying assumption to make it possible to detect selection. By sliding a 2000-year window through our time transect and re-estimating selection coefficient within each window, we can already see that there have in fact been changes in selection pressures at a number of the loci we analyze (Extended Data Figure 6), including at *HLA-DRB1, TYK2* and *HFE* (Figure 3). By comparing the estimated age of the mutation that contributed each selected allele^9^, to the extrapolated time to reach fixation given its estimated *s-*value, we find that around half of the mutations have true ages an order of magnitude larger than the expected sweep age, which means that selection coefficients on the alleles must have shifted over time (Figure 2c).

An alternative explanation for this paradox is to hypothesize that West Eurasians have been experiencing qualitatively more and different natural selection in the Holocene than in earlier periods because of rapidly changing lifestyles and economies. Without a comparable time transect before the advent of food production and societies with high population densities, it is impossible to test this directly. However, this hypothesis is consistent with our evidence of particular intense selection for blood-immune-inflammatory traits, and our evidence that selection for these traits becoming even stronger in the Bronze Age than it was in earlier periods (Figure 1c, Extended Data Figure 4b).

We project that there are at least 5000 independent signals of directional selection (half of the 10361 non-HLA loci found at the FDR=50% threshold) that are in linkage disequilibrium with the overwhelming majority of variants in the genome (Extended Data Figure 2b). This seem to be at odds with findings that there has been relatively little contribution from directional selection to allele frequency changes in genome compared to much larger forces of gene flow, genetic drift, and purifying or stabilizing selection^69^. In fact, there is no conflict. Our method allows us to partition the effects of selection at each SNP into the effects of directional selection (*s*), and the combined effects of fluctuating selection and drift (σ^2^). We estimate that only 2.35 ± 0.13% (jackknife standard deviation) of allele frequency changes are due to directional selection. These results suggest that selection is so rampant that even if a tiny fraction of allele-frequency change is due to directional selection, this corresponds to many hundreds of loci. A corollary is that recent studies finding that stabilizing selection is relatively more important than directional selection in shaping the human allele frequency spectrum^70^ are fully reconcilable with our analyses.

It is important to apply similar approaches to ancient DNA time series over longer times and to other world regions. Comparison of ancient DNA time transects would allow more generalizable insights by identifying which patterns of selection are shared and which are distinctive to the human population history of Holocene West Eurasia.

## Methods

### Testing for selection while correcting for population structure

We used a generalized linear mixed model (GLMM) approach to correct for population structure, a major confounder in scans for significant changes in frequency over time especially as major migration and population mixture have been common in almost all parts of the world. Previous studies have corrected for structure in ancient DNA time transects by modeling the population history and estimating mixture proportions, which works optimally only if there are data from the true source population, which is rarely the case. It is tempting to use an unsupervised approach like Principal Component (PC) to address population structure. However, after experimentation we found this is not effective as PCs are correlated with sample dates which creates collinearity with the quantity we are most interested in (the time-varying component), inflating the empirically estimated variance and reducing power.

The mixed model approach, which is often deployed in the context of genetic association studies to address similar challenges^71^, offers a way to address these issues by combining the structured data in an unsupervised manner and estimating fewer parameters over a wider span of time which results in greater power compared to employing separate regression analyses for each population or comparing the estimated means from different groups. Our simulations show that under simplifying assumptions, a GLMM is more powerful in controlling for population structure and detecting change in allele frequency compared to a generalized linear model using the top principal components (PC) as covariates (Extended Data Figure 14). Thus, despite the fact that the model fitted by the GLMM is far from that expected under true selection, and will miss real signals at sites with fluctuating selection like *TYK2* rs34536443, the method has advantages, and we found in practice that it detected many loci.

We used our GLMM to fit a linear time-varying component to the logit (log-odds) transformation of allele frequency at each position in the genome, and then to test if there is evidence for a consistent trend in allele frequency change over time for all populations. We search for evidence of such a trend beyond the prediction based on population structure and associated genetic drift relating sampled individuals in space and time as measured by the covariance of genotypes over all the individuals, known as the Genetic Relationship Matrix (GRM). In our GLMM, the response variable for each tested allele *j* is the allele count. The allele counts for an individual *i* are drawn from a binomial distribution *B*(2*, p_ij_*), where 2 is the number of chromosomes each person carries at each position, and *p_ij_* is the unknown frequency of allele *j* in the population in which the tested individual *i* lives. A logit link function allows the frequency *p_ij_* to be modeled as a linear combination of covariates. This is a generalization of the Logistic Mixed Model where the response variable is binary:

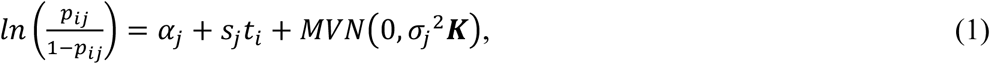

The logit function, 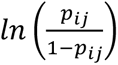, transforms allele frequency so its expected change per generation is proportional to the selection coefficient *s_j_* (regardless of *p_ij_*)^72,73^. *⍺_j_* is a constant related to the average logit transformation of allele frequency in sampled individuals at time *t*=0 today. *s_j_* is the per-generation selection strength at the allele, assumed constant over time and space during the period of our time transect; our test for selection is simply a test for whether the equation fits significantly better if *s_j_* is non-zero than if it is zero. *t_i_* is the negative sampling date in the past, in units of twice the generation interval^72,73^ (assuming 29 years per generation). *g_ij_* is a random effect, an error term capturing individual-specific variability not explained by fixed effects (*⍺_j_* + *s_j_t_i_*). It differs from the error term in a Generalized Linear Model, which is independently and identically distributed following a normal distribution. In our GLMM, the error term is drawn from the vector ***g_j_**∼MVN*(0, *σ_j_*^2^***K***;, following a multivariate normal distribution, where **K** is the covariance matrix structure (the GRM), the empirically observed relatedness of all individuals to each other, and *σ_j_*^2^ measures the drift at that variant.

*s_j_*, *σ_j_*^2^ and *⍺_j_* are independently estimated for each of 9.9 million variants. Refitting them without being constrained by the values at other variants means the methodology is robust to false-positives due to processes that vary across SNPs such as degree of background selection which increases the effective amount of random genetic drift or variation in minor allele frequency (MAF); these nuisance random effects are soaked up by allowing *σ_j_*^2^ and *⍺_j_* to vary, allowing us to test for a time-dependent influence on allele frequency fluctuations *s_j_* beyond what can be explained by the GRM. Our test for a non-zero *s_j_* is thus a test for selection above and beyond what could be explained not just by structure but also other non-time-dependent processes. The penalty we pay for estimating variance components at millions of SNPs—in contrast to the constant variance component assumption used in mixed model analysis in Genome-Wide Association Studies (GWAS)^71^—is computational load. We grouped individuals with similar ancestry and dates into 3000 clusters (Supplementary Information section 7); at this resolution, our method required ∼140,000 CPU hours.

Using the GLMM, we obtain a point estimate for the selection coefficient at each variant and its standard error, and a Z-score for the number of standard errors this is from zero, a naïve test for selection. In practice, the statistic needs recalibration as it is inflated due to unmodeled features of the data, so we empirically assess significance from enrichment of signals in independent GWAS (Supplementary Information section 2).

### Fitting the generalized linear mixed model (GLMM)

We developed PQLseq2, a faster implementation of PQLseq^74^ for fitting the GLMM to count data. Despite a 27-fold speed increase, running a GLMM on ∼15,000 individuals for ∼9.9 million variants was infeasible given our resources. To make analysis tractable, we grouped individuals into clusters of individuals with similar ancestry and coming from similar times.

To identify the T = 3000 clusters we analyze, we required there to be a maximum date gap G = 500 years between any two individuals in each cluster. We initialized the interval I = (l=2, r=T) with midpoint m. We applied hierarchical clustering on the top 30 principal components (PCs) using the sklearn.cluster.AgglomerativeClustering function in Python with default parameters and n_clusters = m. For each of the S clusters from the previous step, we performed hierarchical clustering on the dates with distance_threshold = G and n_clusters = None. If the resulting number of clusters was larger than T + 1, we repeated the process with I = (l, m). If it was less than T-1, we updated I = (m, r). We repeated these steps until the final number of clusters was within T-1 to T+1. Across 3,000 clusters, the individuals per cluster has a first quartile of 1, a median of 3, a third quartile of 6, and a maximum of 46.

We use the same GLMM model as for the single variant analysis. However, the cluster can include more than one individual. The allele counts for each cluster *i* are drawn from a binomial distribution *B(2n_i_, p_ij_)*, where *n_i_* is the number of diploid individuals in the cluster, and *p_ij_* unknown frequency of allele *j* in the population where individuals in cluster *i* reside.

### Proportion of variance explained by directional selection

The proportion of variance in allele frequency on the logit scale for each SNP *j* is:

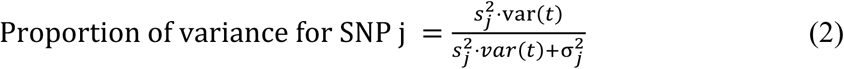

We used 1000 independent SNPs, randomly selected across the genome with pairwise LD (r^2^) less than 0.05, to estimate that directional selection explains an average of 2.35% of the variance in allele frequency, with a standard error of 0.13% based on jackknife estimation. The GLMM used for this analysis is based on the full sample size, rather than clustering individuals according to their ancestry and date.

### Covariance structure for the GLMM

The covariance structure matrix **K** for clusters m and n is defined as:

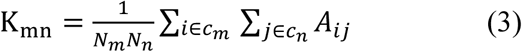

Where *c*_*m*_ is the set of individuals in cluster *m*, *N*_*m*_ is the number of individuals in cluster m, and A_ij_ is the genetic relationship matrix (GRM) between individuals i and j and defined as^17^:

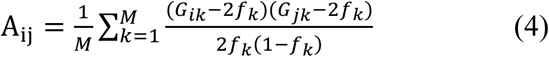

Here *G_i_*_*k*_ is the genotype for SNP k of individual i, *f*_*k*_ is the allele frequency of SNP k, and M is the number of SNPs. We created a GRM using all autosomal SNPs and applied a leave-one-chromosome-out (LOCO) scheme to prevent proximal contamination^75,76^, creating a separate GRM for each chromosome.

### Polygenic score computation

The polygenic score (PGS) is a weighted average of genotypes for M independent variants.

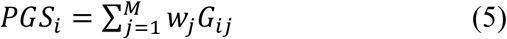

Here, *G_ij_* is the genotype for SNP *j* of individual *i* and *w_j_* is the SNP weight. We generate four variations of the PGS score by including or excluding the HLA region, and utilizing the GWAS effect values (*β_i_*) or only the sign of the effects, sign(*β_i_*), as weights. For each phenotype, we generate an independent set of SNPs using a two-step clumping and thresholding proces s. Initially, we clump SNPs with PLINK using a P-value <10^−3^, r^2^<0.05, and a 500 kb window. Then, we select the SNP with the smallest P-value as the index SNP, remove SNPs with D’ > 0.2 within 500 kb, and repeat until no SNP remains. Consequently, all remaining SNPs have P<0.001, and if two SNPs are within 500 kb, their r^2^ < 0.05 and D’ < 0.2. To minimize residual population structure, we use the linear mixed model (LMM),

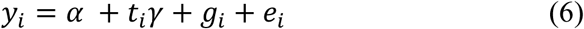

Here, *y_i_* is the polygenic score of the sample *i*, centered at zero and scaled by the standard error of PGS of the modern samples; *t_i_* is the date scaled down by −10000 (so it is in units of ten millennia); *⍺* is the intercept; 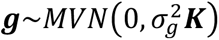 is a vector of random effects where the covariance structure matrix **K** is the genetic relationship matrix; and 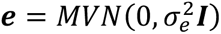 is a vector of residual errors where **I** is the identity matrix. The coefficient *γ* is the change of the polygenic score over 10000 years in unit of standard deviation from the zero-centered PGS of the modern samples. We use the coefficient *γ* as a proxy for directional polygenic selection.

### Fitting the linear mixed model (LMM)

We used GEMMA (v0.98.5)^77^ to fit the LMM and estimate the polygenic selection coefficient (*γ*). The running time was tractable, so we did not apply the clustering algorithm used in the GLMM analysis. We used the genetic relationship matrix as the covariance structure matrix **K**. Here, PGS is calculated over all autosomes, and we could not use the LOCO approach from single-variant GLMM to avoid influence from neighboring positions. Instead, we used 80,085 high-quality, independent SNPs generated by the ‘indep-pairwise 1000 1 0.05’ option of PLINK2 to calculate a GRM, using this as a covariance structure in the LMM to handle population structure and reduce proximal contamination.

### Analyzing correlation between GWAS summary statistics and selection coefficients

We use LD score regression (LDSC) version 1.0.1^23,29,59^ to calculate the genetic correlation between GWAS summary statistics and the estimated selection coefficient. We use the pre-calculated LD scores computed using individuals of European ancestry from the 1000 Genomes Project, which are provided with the LDSC software. To compute trans-ethnic genetic correlation, we used S-LDXR software^78^. We used the pre-calculated reference files for European and East Asian populations that are provided with this software.

### Studying heritability enrichment and computing standardized effect size (*r*^∗^)

We utilized stratified LD score regression (S-LDSC)^29^ to estimate the contribution of each annotation to the heritability of polygenic traits. The set of annotations of interest was combined with the baseline-LD model (v2.2), which includes 97 annotations modeling minor allele frequency (MAF), linkage disequilibrium (LD), and functional architectures including coding regions, promoters, enhancers, and conserved elements^29,79,80^. Heritability enrichment quantifies the effects of the annotation. It is defined as the proportion of heritability explained by SNPs in the annotation divided by the proportion of SNPs in the annotation. The standardized effect size (*r*^∗^) measures the effects unique to the focal annotation after conditioning on all the other annotations in the baseline-LD model^67^.

### Adjusting for residual inflation in directional polygenic analysis

To adjust for residual inflation in the estimated ***Z***_***γ***_ for each trait, we carried out 100 randomizations for each trait of interest, using the same SNPs employed for calculating the PGS of that trait and randomly assigning a weight of +1 or −1 to each SNPs for each simulation. The simulated PGS is not expected to show a signal of selection, as the weights are randomly flipped and should cancel for polygenic traits. Therefore, for each trait, we define an inflation factor by calculating the ratio of the median 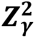 for the simulation to the median of the chi-square distribution with 1 degree of freedom (0.455). If the inflation factor exceeds the median of 3.13 across all traits, we apply the median value as the correction factor for the test statistics. This allows us to carry out a valid analysis of polygenic signals driven by only a few SNPs under strong selection, which can cause a large inflation factor.

### Simulation of genotypes

To simulate the genotypes of individuals for a variant with a selection coefficient *s_j_*, we used a random sample drawn from a Gaussian distribution with a covariance matrix of *σ_j_*^2^*K*. We estimated the genetic relationship matrix A using real data, and randomly selected *σ_j_*^2^ from an empirical distribution. This distribution was derived by applying a GLMM to real data, specifically for 1000 randomly chosen SNPs, without clustering. We employed equation 1 to simulate different selection coefficients and determined the initial allele frequency by drawing from an empirical distribution of allele frequency in modern samples. We used this value as a constraint to define the constant α*_j_*. To sample genotypes, we drew from a binomial distribution, with the probability of the alternative allele calculated using the standard logistic function applied to both sides of equation 1.

### Sources of data for 8433 ancient individuals

We restricted to 8433 ancient individuals living between longitude 25W and 60E and latitude 35N to 80N (Online Table 1). For 3644 ancient individuals, the sequences we analyze are published in other papers^11,81–196^ and are reanalyzed here. For 244 ancient individuals, we newly publish shotgun sequencing data obtained on Illumina instruments on libraries for which either in-solution enrichment data from the same ancient DNA samples, extracts, or libraries was previously published; the present study serves as the formal report of these new sequences, and reanalysis of the data presented here should cite both the present study and the study that originally reported data from these individuals. Online Table 2 lists these samples along with newly reported shotgun data for an additional 56 anonymized newly reported individuals (for a total of 300 newly reported shotgun genomes which have a median of 4.87-fold coverage and of which 40 have at least 17-fold coverage).

For 74 ancient individuals, we publish higher coverage in-solution enrichment data based on additional extracts, libraries and sometimes recaptures of libraries for which smaller amounts of in-solution enrichment data from the same samples were previous published, obtained by adding data from 155 newly reported ancient DNA libraries (Online Table 3). The present study serves as the formal report of these merges of previously published data with the newly generated data. Reanalysis of the data presented here should cite both the present study and the study that originally reported data from these individuals.

For 4471 never-before-reported ancient individuals obtained by sequencing 5227 newly reported ancient DNA libraries (Online Table 3), we release raw ancient DNA data with permission of sample custodians. The individuals are anonymized, with the only information provided about them being point estimates of their dates and broad geographic categorization into five regions of West Eurasia. Analyses of population history and presentation of full archaeological information will be provided in subsequent studies and we request that the research community respects “Fort Lauderdale principles”^15^, allowing the generators of the data to report the first population history analyses. Any researchers are welcome to analyze the full dataset for studies of natural selection.

### Sources of data for contemporary individuals

We analyzed data from 6,510 contemporary individuals, comprising 5,935 from the UK Biobank^19^, 503 from the 1000 Genomes Project^18^, and 72 from published studies^174,197–201^.

For the UK Biobank data, we selected individuals genotyped on the UK Biobank Axiom array, excluding those sequenced on the UK BiLEVE array to minimize batch effects. To ensure broad representation across Western Eurasia, we subsampled the UK Biobank, limiting the selection to at most 300 people per “country of birth” within Western Eurasia, focusing on countries with ancient DNA in this study. This yielded 6,088 individuals.

For the remaining individuals, we calculated the Mahalanobis distance P-value based on the top 20 principal components, assuming the squared Mahalanobis distance follows a χ^2^ distribution with 20 degrees of freedom. Samples with P-values below the Bonferroni-corrected threshold of 8.2e-6 were removed, resulting in a final set of 5,935 individuals from 58 countries, with a median of 55 and a mean of 102 individuals per country. These principal components were derived from the full set of UK Biobank samples.

### Ancient DNA data generation

The great majority of wet laboratory work was performed in the ancient DNA laboratory at Harvard Medical School in Boston, USA, following established protocols that evolved over time from mostly manual processing (sample preparation, DNA extraction with silica columns^202,203^ and partial UDG treated double-stranded library preparation^204,205^; capture was automated using a Perkin Elmer EP3 or Agilent Bravo NGS Workstations^11,206,207^) to mostly automated processing (DNA extraction^208^, double- and single-stranded library preparation^209^, capture, pooling for sequencing). New libraries (if not deeply shotgun sequenced) were enriched with the Twist Ancient DNA panel^193^, whereas older libraries were enriched with the 1240k reagent (or its predecessor, 390k and 840k). We sequenced on an Illumina NextSeq500 instrument until 2019, when we switched to an Illumina HiSeq X10 instrument, and finally to an Illumina NovaSeq X instrument in 2022. Archaeologists or collaborators from other ancient DNA laboratories in some cases provided sample powder, DNA extracts, or libraries, which we continued to process. Online Table 3 provides summary statistics based on in-solution enrichment for 5382 ancient DNA libraries for which we newly report data.

### Ancient DNA bioinformatic processing

Most of the newly reported data come from sequencing the products of in-solution enrichment targeting a set of more than a million known polymorphisms^193,207^. In-solution enrichment extracts more information by enriching sequenced molecules to overlap sites that are polymorphic in humans (which also helps to greatly reduce the proportion of non-endogenous bacterial/microbial sources that colonized the samples post-mortem). The great majority of ancient DNA libraries we analyzed are marked with identification tags (barcodes and indices) before sequencing in pools. We merged paired-end sequences, requiring that there is no more than one mismatch in the overlap between paired sequences where the base quality is at least 20 or three mismatches if the base quality is <20. We did not analyzed sequences we could not merge. We stripped adapters and identification tags to prepare molecules for alignment. A custom toolkit (https://github.com/DReichLab/ADNA-Tools) was used for all these steps. We aligned merged sequences to the hg19 version of the human reference genome with decoy sequences (hs37d5) using the single-ended aligner, BWA SAMSE v.0.7.15^210^ with typical ancient DNA alignment parameters -n 0.01 -o 2 and -l 16500 which disables pre-alignment seeding. Duplicate reads were marked using Picard MarkDuplicates (v.2.17.10)^211^. In addition, merged sequences are also mapped with the same parameters as the Reconstructed Sapiens Reference Sequence (RSRS)^212^, which enables mitochondrial-specific metrics. Our bioinformatic processing produces data and key metrics, including estimates of authenticity based on elevated damage rates at the end of sequences (indicative of ancient DNA), contamination rates, and endogenous rates. A subset of libraries that had a very high proportion of human DNA were additionally shotgun sequenced to generate coverage throughout the genome and underwent the same bioinformatics processing.

### Imputation

To carry out imputation, we used as input either data from ancient individuals (mapped sequences) or modern individuals (SNP array genotypes), and then used allelic correlation patterns in a haplotype reference panel^18,213^ to predict genotypes at millions of sites.

In detail, for each sample we used bcftools mpileup (v1.13)^214^ to generate genotype likelihoods for all variants (SNPs and indels) in the panel. We used the high coverage (30x) 1000 Genomes Project^18^ phase 3 sequences as the reference panel and converted the assembly version to GRCh37/hg19 using CrossMap (v0.5.2)^215^. We kept 2504 unrelated samples and biallelic variants that pass all the quality control filters reported by gnomAD (v2.1.1)^216^. We used GLIMPSE (v1.0.0)^20^ with the reference panel to impute and phase each sample individually. Due to higher reference bias for indels, we ignored their genotype likelihood, set them to missing, and passed this to GLIMPSE to impute all biallelic autosomal SNPs and indels based on genotype likelihood of SNPs and haplotype information for both SNPs and indels in the reference panel. This means we only use reference panel information to impute indels even where we have sequences overlapping the indels. After imputation is done, we add the genotype caller information of all variants (SNPs and indels) to the final bcf file.

To minimize discrepancies between imputation of ancient DNA and UK Biobank data, we re-imputed the UK Biobank genotyping data. We utilized Affymetrix confidence files to simulate genotype likelihoods and processed these through the same imputation pipeline employed for ancient DNA.

### Sample quality control

For each imputed sample, we define imputation quality score *IQS* = *mean*(*GP*_1_|*GT* = 1), where *GT* is the most likely genotype based on the imputed genotype posterior ***GP*** = (*GP*_0_, *GP*_1_, *GP*_2_) and 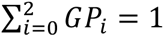. We only kept samples with high imputation quality score IQS>0.9. We used KING to detect duplicates and related samples up to the second degree. We prioritize samples by their IQS and drop relatives up to the second degree until there are no two samples that are second-degree related or closer. We also fit a linear regression model to the top 100 PCs as explanatory variables and used the reported date of samples as the response variable to remove outliers where reported and predicted date are very different. Sample quality control is described in detail in Supplementary Information section 1.

### Variant quality control

The data analyzed in this study come from multiple sources and sequencing technologies: imputed ancient DNA sequences (shotgun sequences and enrichment for more than a million SNPs), European ancestry individuals largely from the 1000 Genomes Project, and imputed individuals of Western Eurasian ancestry from the UK Biobank genotyped using the UK Biobank Axiom Array. Variant quality control involved a two-step procedure. Initially, we applied brute-force filtering to compute principal components, allowing for the identification of ancestry-matching samples across datasets with similar allele frequencies. We filtered out variants if their allele frequencies differed strongly between sample sets, with the goal of minimizing batch effects from combining samples from different sources. This results in 9,926,484 variants, including 8,212,921SNPs and 1,713,563 indels, passing the final variant QC out of 52,382,872 imputed variants. The step-by-step variant quality control process is detailed in Supplementary Information section 1.

### Allele frequency trajectories

We computed allele frequency trajectories using all individuals in the time series. We used a moving average sliding window, with a window size of 1000 years and a step size of 100. We used a binomial likelihood function to estimate the mean, confidence intervals, and standard error. We smoothed the mean and standard error using the GaussianProcessRegressor function from the Scikit-learn library in Python. We parameterized this function with alpha = 1e-4 and a 1*RationalQuadratic kernel, with length_scale_bounds set to (10, 1e6). We clipped the resulting values to remain within the range of 0 and 1.

### Assembly of GWAS data to which we correlated selection coefficients

We processed 6,951 phenotypes with European ancestry from the Pan-UK Biobank^24^, of which 452 passed quality control with the flag phenotype_qc_EUR being PASS. We also analyzed 107 curated sets of independent GWAS studies^55,217^ with European ancestry for meta-analysis. For the trans-ethnic analysis, we analyzed 31 phenotypes in East Asians: 30 phenotypes from the Biobank of Japan (BBJ)^218^ and the GWAS summary statistics from the study of years of schooling GWAS by Chen et al. 2024^219^. We then co-analyzed these GWAS results with that of the corresponding phenotypes in the Pan-UK Biobank.

## Supporting information

Supplementary Information sections 1-7

## Data availability

The aligned sequences for newly reported data are available through the European Nucleotide Archive under an accession number that will be made available upon final publication. Imputed genomes for all ancient and modern individuals are available at the permanent Dataverse repository at a link that will made available upon final publication.

## Software and code availability

An interactive web application for this study is available at https://reich-ages.rc.hms.harvard.edu. The PQLseq2 software is available from https://github.com/zhengli09/PQLseq2.

## Acknowledgments

We thank Brian Browning, Shai Carmi, Evan Koch, Mark Lipson, Iain Mathieson, Vagheesh Narasimhan, Benjamin Neale, Simone Rubinacci, Guy Sella, and Shamil Sunyaev for discussions during the development of this project. We are grateful to Iosif Lazaridis, Heng Li, Adam Micco, Mariam Nawaz, Zhao Zhang, and Mengyao Zhao for bioinformatic support; and to Nicole Adamski, Rebecca Bernardos, Nasreen Broomandkhoshbacht, Kim Callan, Alex Claxton, Olivia Cheronet, Elizabeth Curtis, Matthew Ferry, Trudi Frost, Ilana Greenslade, Eadaoin Harney, Lora Iliev, Aisling Kearns, Jack Kellogg, Ann Marie Lawson, Megan Michel, Jonas Oppenheimer, Iris Patterson, Susanne Nordenfelt, Lijun Qiu, Kristin Stewardson, Anna Szécsényi-Nagy, Noah Workman and Fatma Zalzala for wet laboratory support. We are grateful to the archaeologists and anthropologists who supported early release of raw data for samples they shared with us for the purposes of studies of selection, decoupled from the archaeological contextual information which will be presented in papers on which they are co-authors and is necessary for any studies of the population history of these samples. This research was conducted by using the UK Biobank Resource under Application 16549. We acknowledge the Research Computing Group at Harvard Medical School, for their support with the computational analyses in this paper and the AGES web application; support from NIH grant HG012287; from the Allen Discovery Center program, a Paul G. Allen Frontiers Group advised program of the Paul G. Allen Family Foundation; from John Templeton Foundation grant 61220; and from the Howard Hughes Medical Institute (HHMI). This article is subject to HHMI’s Open Access to Publications policy. HHMI lab heads have previously granted a nonexclusive CC BY 4.0 license to the public and a sublicensable license to HHMI in their research articles. Pursuant to those licenses, the author-accepted manuscript of this article can be made freely available under a CCBY 4.0 license immediately upon publication.

## Extended Data Figure Legends

**Extended Data Figure 1:**
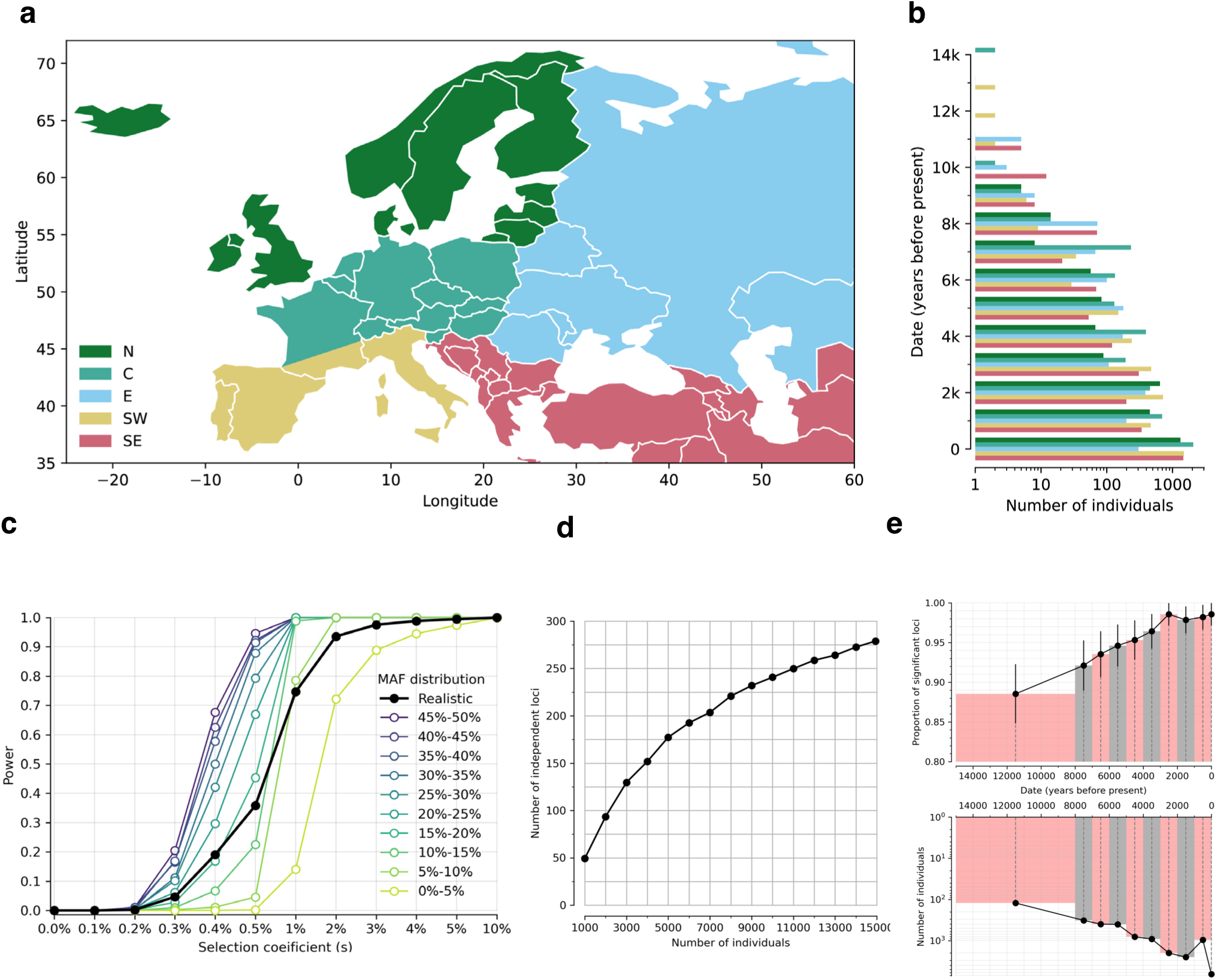
Spatiotemporal distribution of individuals and effect on power. **(a)** Geographic origin: North (N), Central (C), East (E), Southwest (SW) and Southeast (SE). **(b)** Temporal distribution (x-axis on a logarithmic scale). **(c)** Power analysis based on simulations. Sample size, dates, and pattern of genetic relatedness are matched to real data. Power is defined as proportion of true positives expected at p<5×10^−8^. We ran 20000 simulations for each selection coefficient, with minor allele frequency (MAF) at present (time=0) randomly drawn from the MAF distribution in modern Europeans. **(d)** Number of independent and significant loci as function of sample size (from downsampling). **(e)** Effect of age on power. Data are divided into 10 non-overlapping periods; modern individuals are a separate bin. In top panel, y-axis is proportion of loci that remain significant after excluding 100 random individuals from that bin (bottom is number of individuals in the same bin).

**Extended Data Figure 2:**
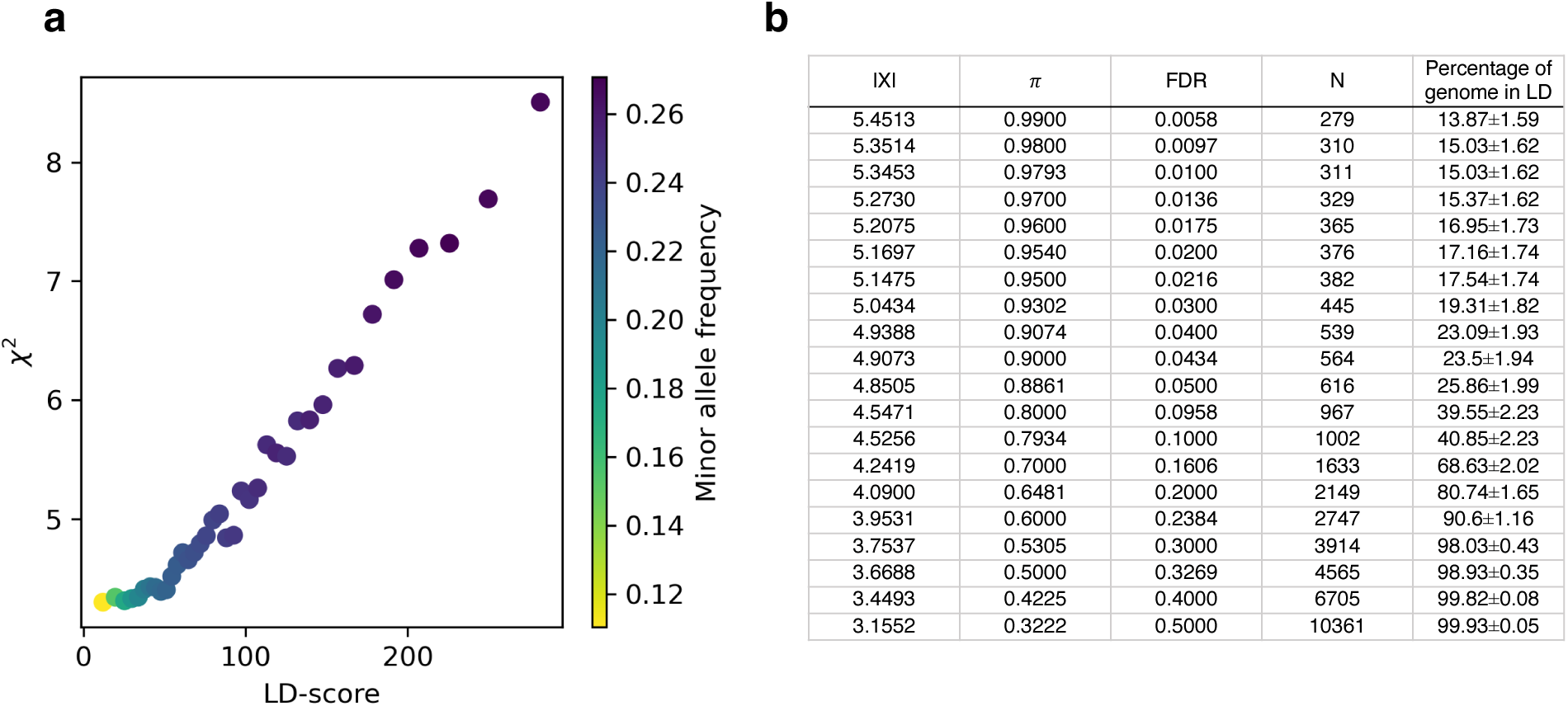
High proportion of genome affected by directional selection. **(a)** LD score plot for nominal χ² statistics, with each point representing an LD score quantile. Values are averaged across each bin. **(b)** Mapping X-score to posterior probability (π), False Discovery Rate (FDR), number of independent loci excluding the HLA region (N), and the percentage of the genome in LD (r² > 0.05) with tag SNPs representing these loci.

**Extended Data Figure 3:**
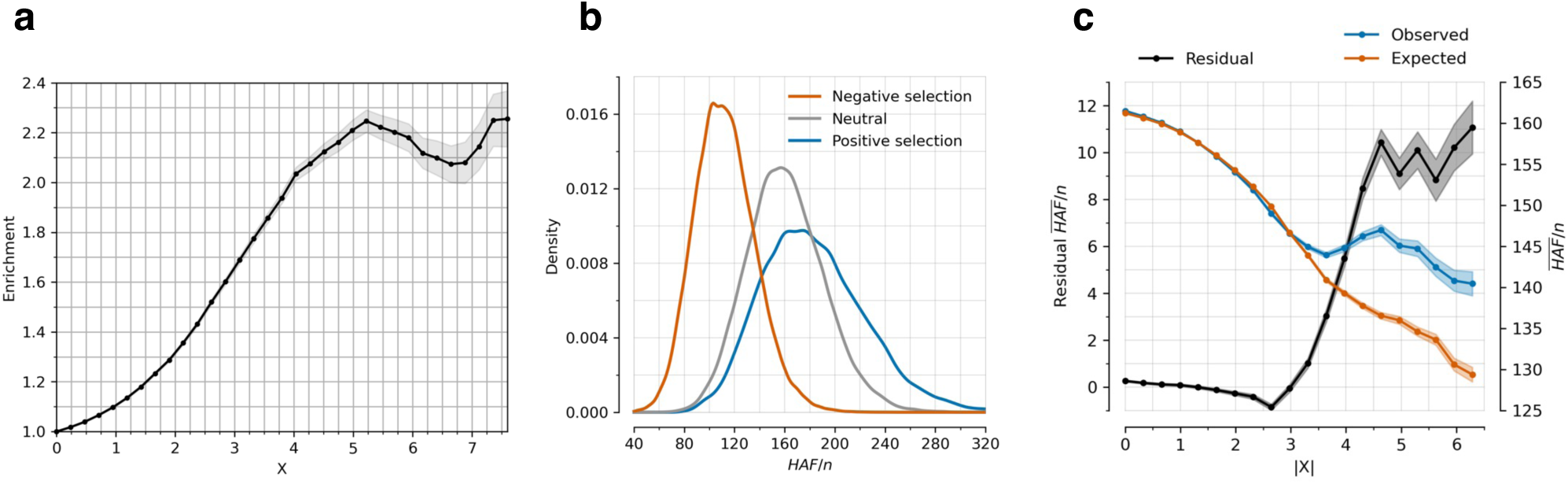
Robustness of directional selection signals (related to Figure 1a,b). **(a)** Proportion of SNPs significant in any of 452 pan-UK Biobank GWAS studies for X-statistics with magnitudes larger than the threshold on the x-axis, adjusted for minor allele frequency and measures of linked purifying selection (McVicker-B, Murphy-phastCons, and Murphy-CADD). Background selection tends to be higher in functional genomic regions, so SNPs with higher |X| are more penalized than in Figure 1a hence the lower plateau. **(b)** Simulating neutral, negative, and positive selection for a 200 kb window around a focal SNP, with derived allele frequency drawn uniformly from [0,1]. The focal SNP has *s*=0.01, population size is constant at 20000 diploid individuals, mutation rate per base pair per generation is 2×10^−8^, and recombination rate is 1 cM per 1 Mb. **(c)** Residual mean (HAF)/n for a haploid sample size n over 200 bp windows is observed minus expected value. Expected value is determined using a linear regression model with McVicker-B, Murphy-phastCons, and Murphy-CADD as variables, providing the expected mean (HAF)/n conditioned on them.

**Extended Data Figure 4:**
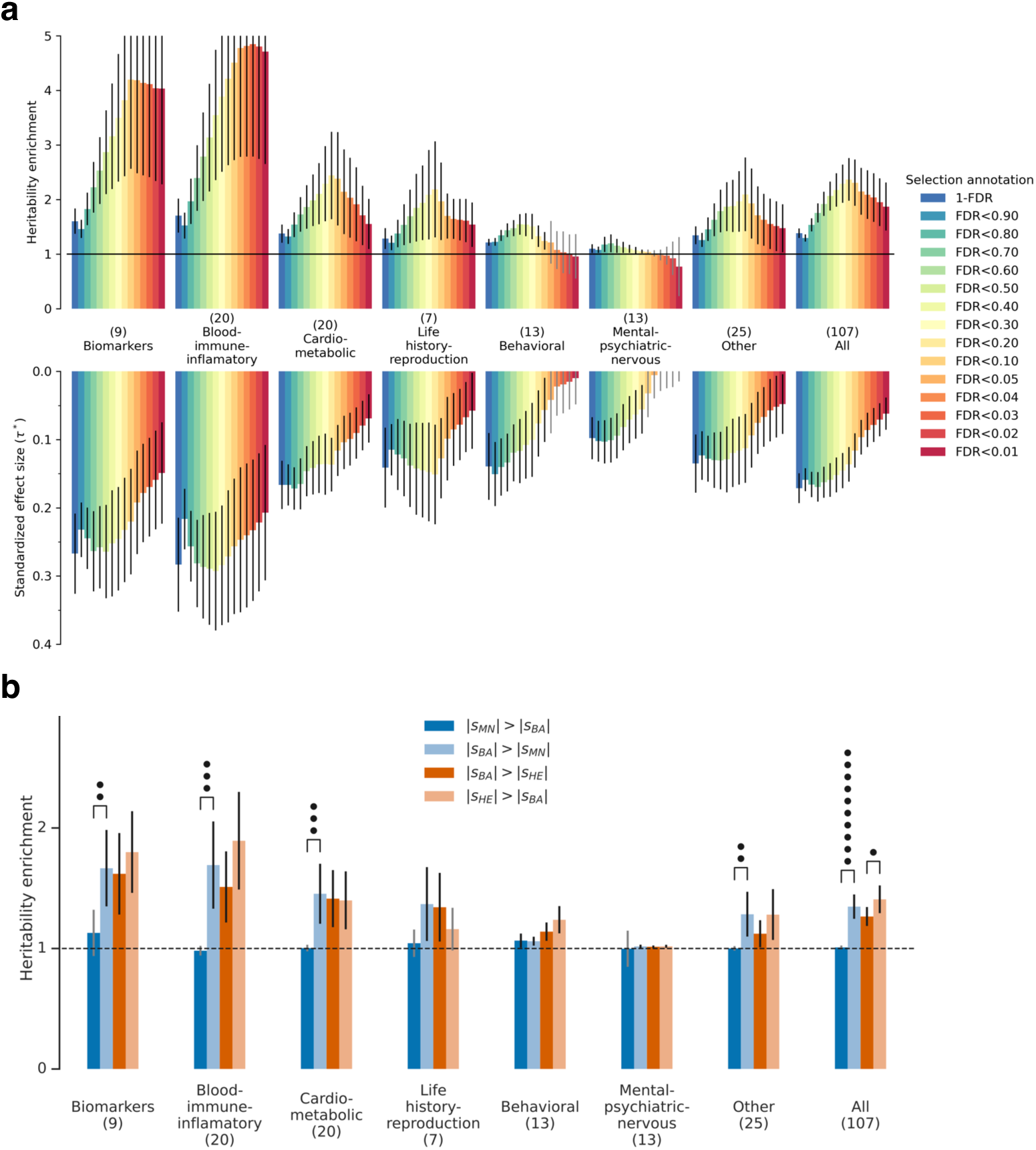
Stratified LD Score Regression shows that alleles affecting blood-immune-inflammatory and cardio-metabolic traits were unusually affected by selection, and that selection intensity increased in the Bronze Age (related to Figure 1c). **(a)** We annotated sites based on their inferred strength of selection—based on their FDR being above a specified threshold, or 1-FDR as a continuous annotation—and used Stratified LD Score Regression (S-LDSC) to study enrichment of GWAS signals and standardized effect sizes (*τ*\*) for traits in different functional categories. Our analysis adjusts for 97 annotations that are known to affect heritability and are part of the standard correction in S-LDSC; dots represent significance of elevation above the baseline of 1 expected for random variants. **(b)** Tests for changes in selection intensity during different cultural transitions: Mesolithic-Neolithic (MN) to Bronze Age (BA); and Bronze Age (BA) to Historical Era (HE). Each annotation is binary, identifying SNPs among the top 5% with the highest probability of experiencing stronger selection during one time period compared to another. This is determined using the estimated selection coefficient and standard error from models separately fit to each cultural period. Error bars are 95% confidence intervals.

**Extended Data Figure 5:**
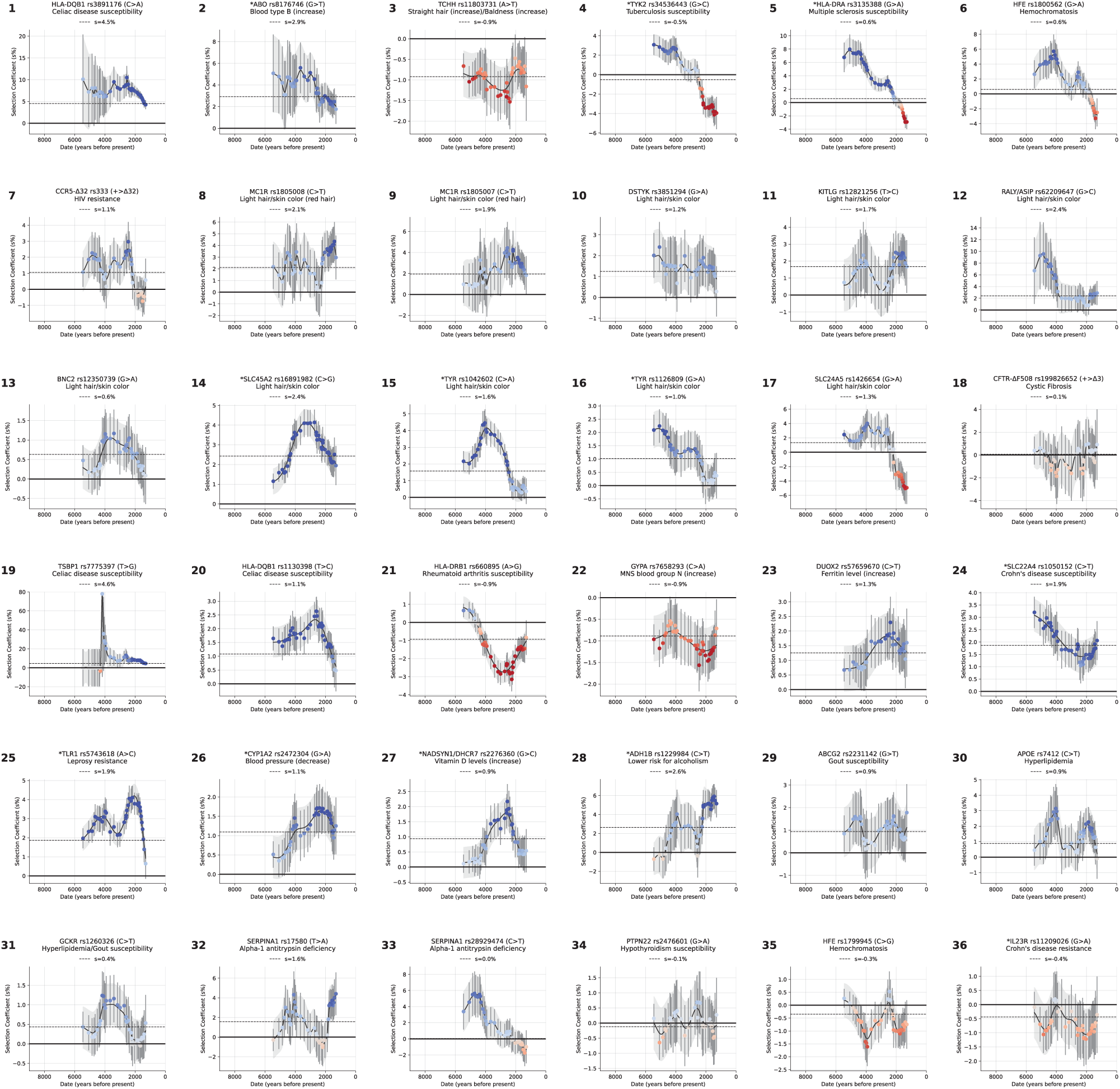
How selection coefficients on single variants changed in intensity over time (for the gallery of 36 loci also highlighted in Figure 3). Time-variant selection coefficients are estimated by refitting our model in sliding windows of 2000 years, with a step size of 100 years, and a minimum of 500 people per window. The present-day is excluded. Color map represents the Z-score for the selection coefficient being non-zero in that window, ranging from −5 (dark red) to 5 (dark blue).

**Extended Data Figure 6:**
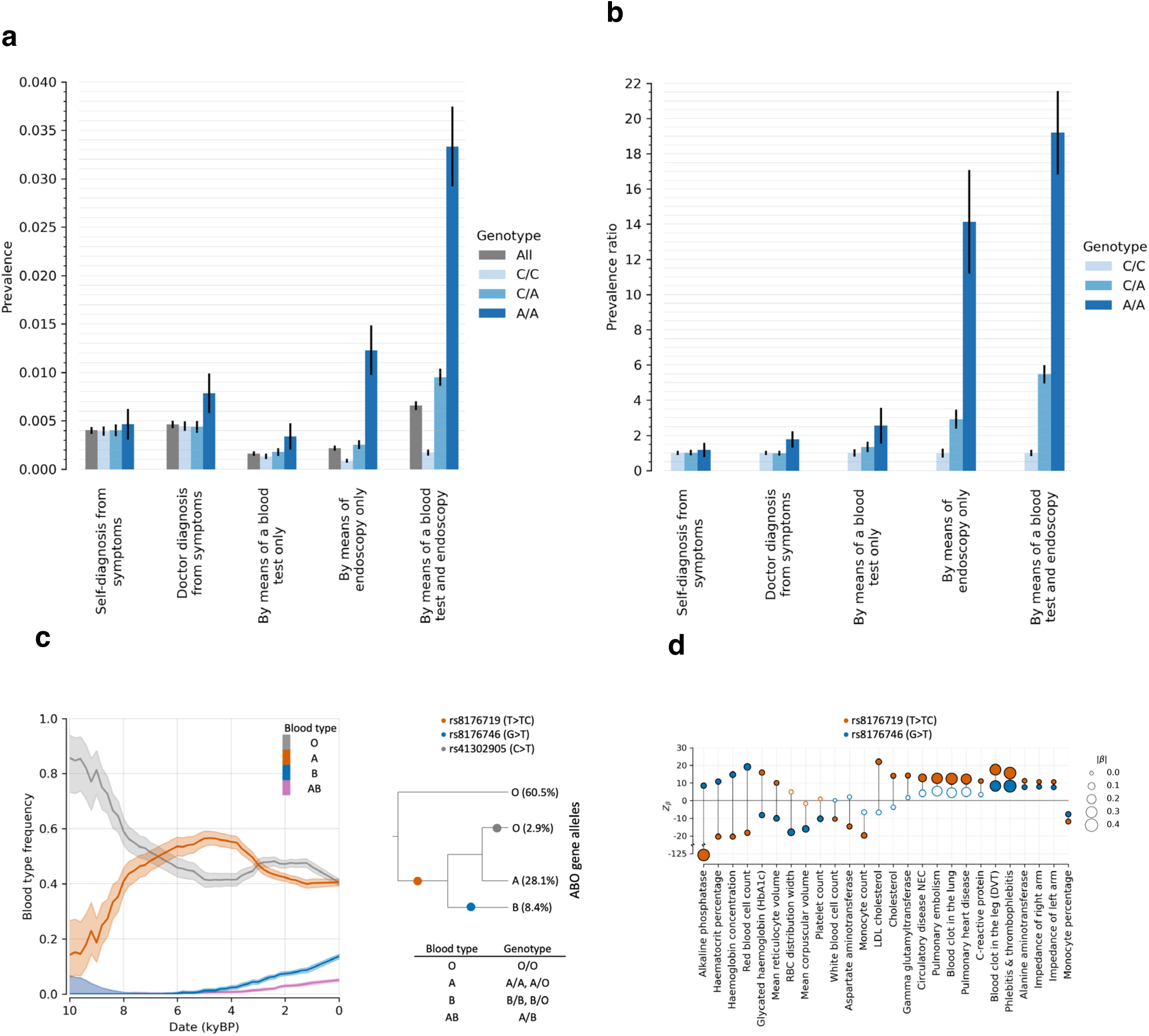
Genotype-phenotype correlations for the signals of selection for Celiac disease at HLA and the ABO blood group locus. **(a)** Prevalence and **(b)** prevalence ratio of individuals with celiac disease or gluten sensitivity (data field 21068) in the UK Biobank, conditioned on the genotype of rs3891176 (C>A). The prevalence ratio compared to the A/A genotype as a baseline; bars are 95% confidence intervals. **(c)** Left: Blood type frequency trajectories for O, A, B, and AB estimated from our aDNA time series. Right: Genealogy of the ABO alleles approximated by Shelton et al. 2021^220^. The allele frequencies are estimated from Europeans in the 1000 Genomes Project; shading gives 95% confidence interval. **(d)** Significant association to traits in Pan-UKBB for the two base pair insertion rs8176719 (T>TC) and SNP rs8176746 (G>T), approximating the alleles A and B.

**Extended Data Figure 7:**
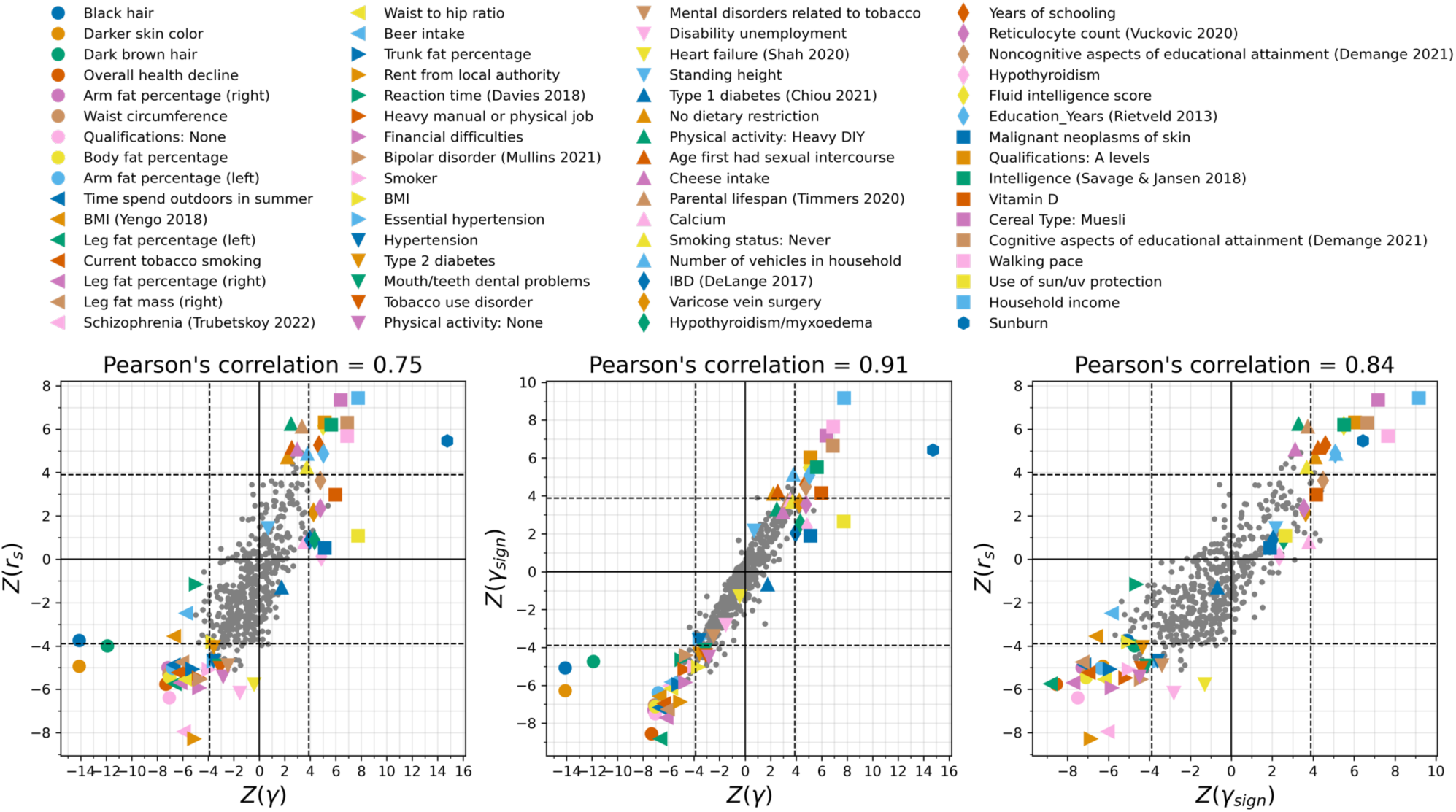
High correlation of 3 tests for polygenic selection (*γ*, *γ*_sign_, r_s_). Each dot represents a phenotype, some annotated by colors. Pearson’s correlation for x and y axes at top; dashed line is the P<0.0001 significance threshold (correcting for 500 tests).

**Extended Data Figure 8:**
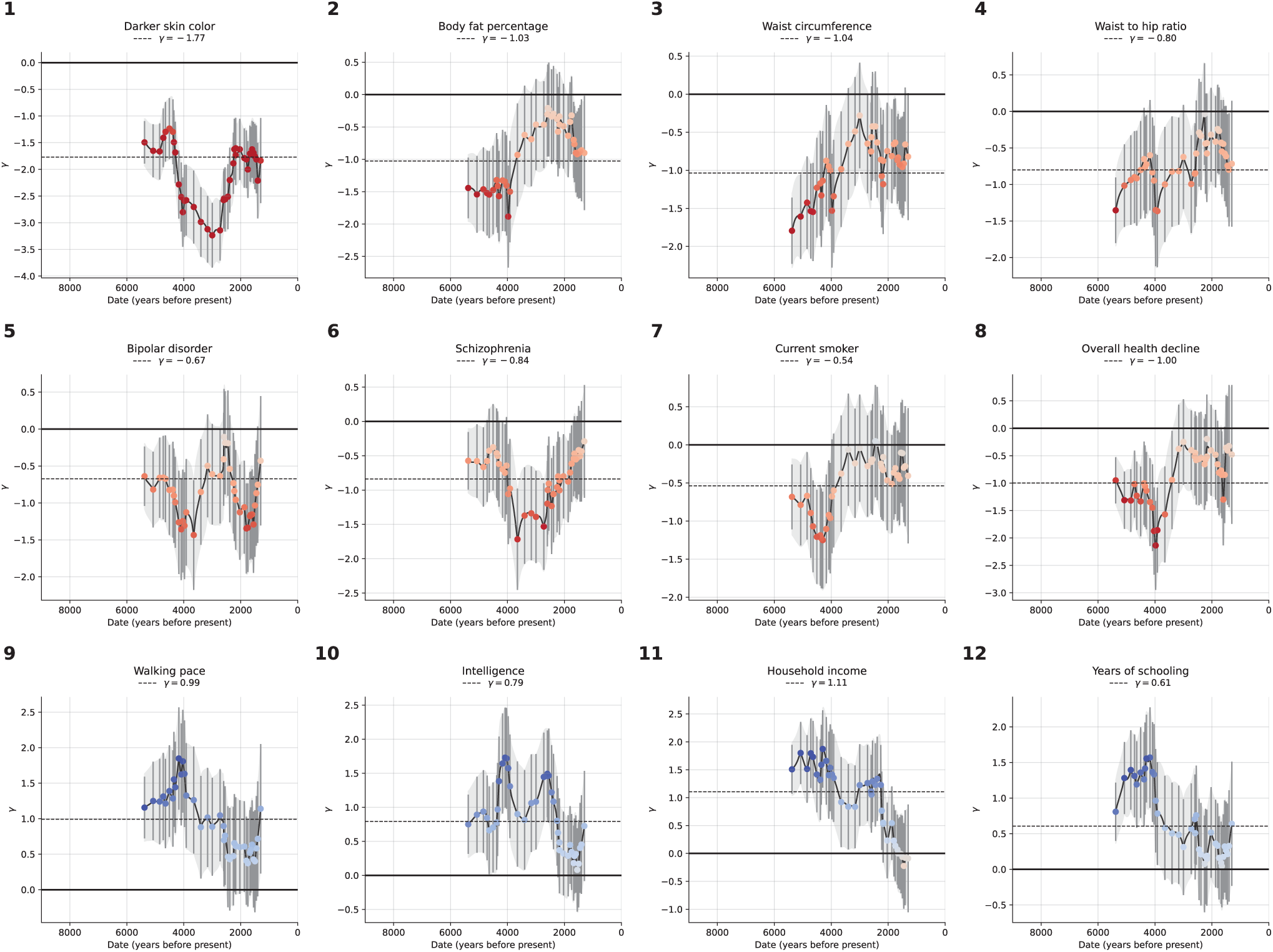
How coordinated selection on alleles affecting the same traits changed in intensity over time (gallery of 12 complex traits also highlighted in Figure 4). We estimate time-variant polygenic selection intensity *γ* by refitting our model in sliding windows of 2000 years, with a step size of 100 years, and a minimum of 500 people per window. The present-day is excluded. Color map represents the Z-score for the selection coefficient being non-zero in that window, ranging from −5 (dark red) to 5 (dark blue).

**Extended Data Figure 9:**
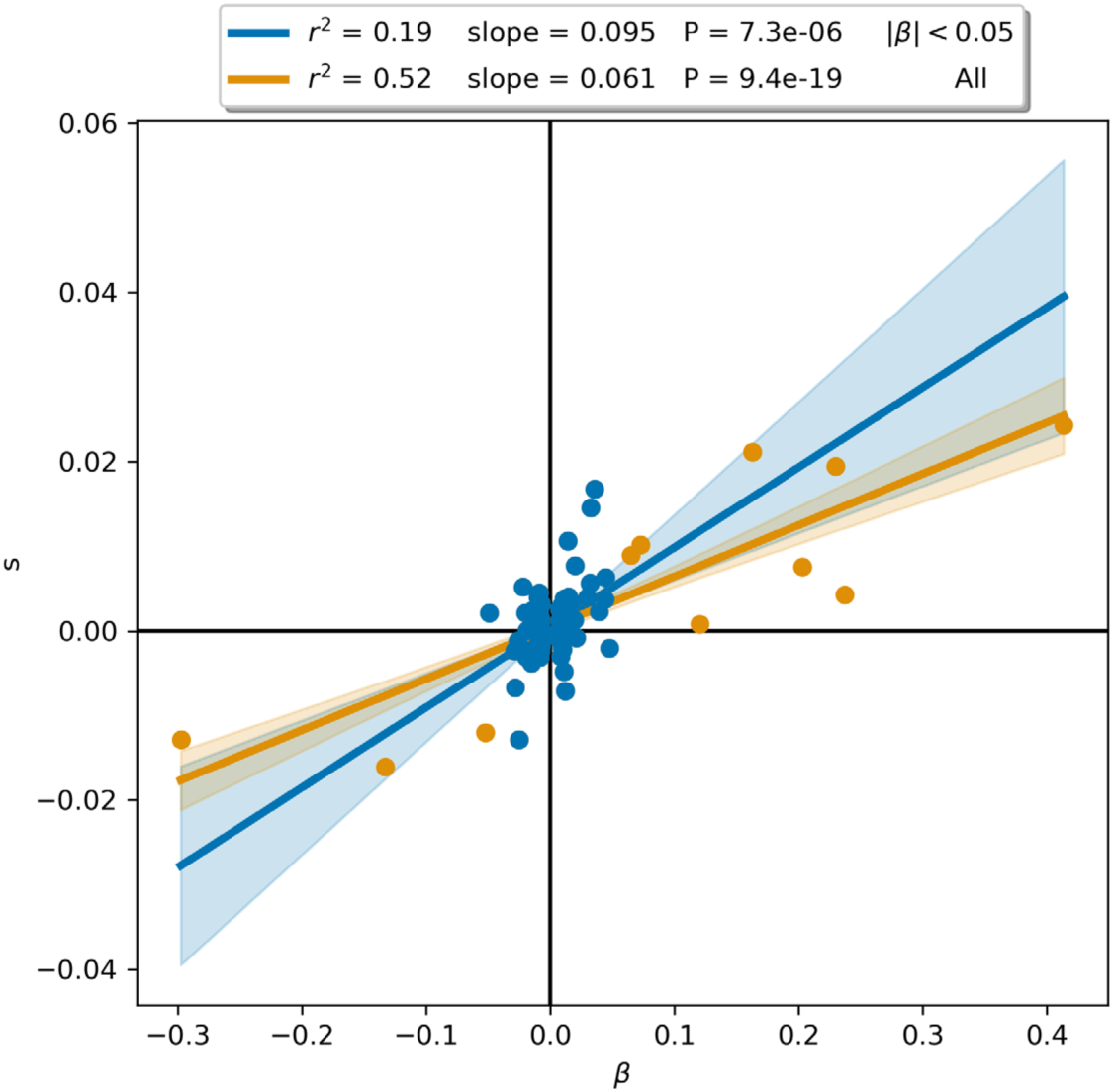
Pigmentation is oligogenic but selection on it was polygenic. Selection coefficient (s) and effect size (β) from the pan-UKBB skin color phenotype for 110 independent SNPs passing the GWAS P-value threshold of p<5×10^−8^. Following ^57^, the orange line is a linear regression on all SNPs (99 blue and 11 orange markers), while the blue line includes only SNPs with |β| < 0.05 (99 blue markers). Although the correlation appears different (with the difference between Fisher Z-transformed Pearson r showing a P-value of 0.001), the slopes are not significantly different (P = 0.10), consistent with a model in which selection for pigmentation had an equal impact on all variants in proportion to effect size.

**Extended Data Figure 10:**
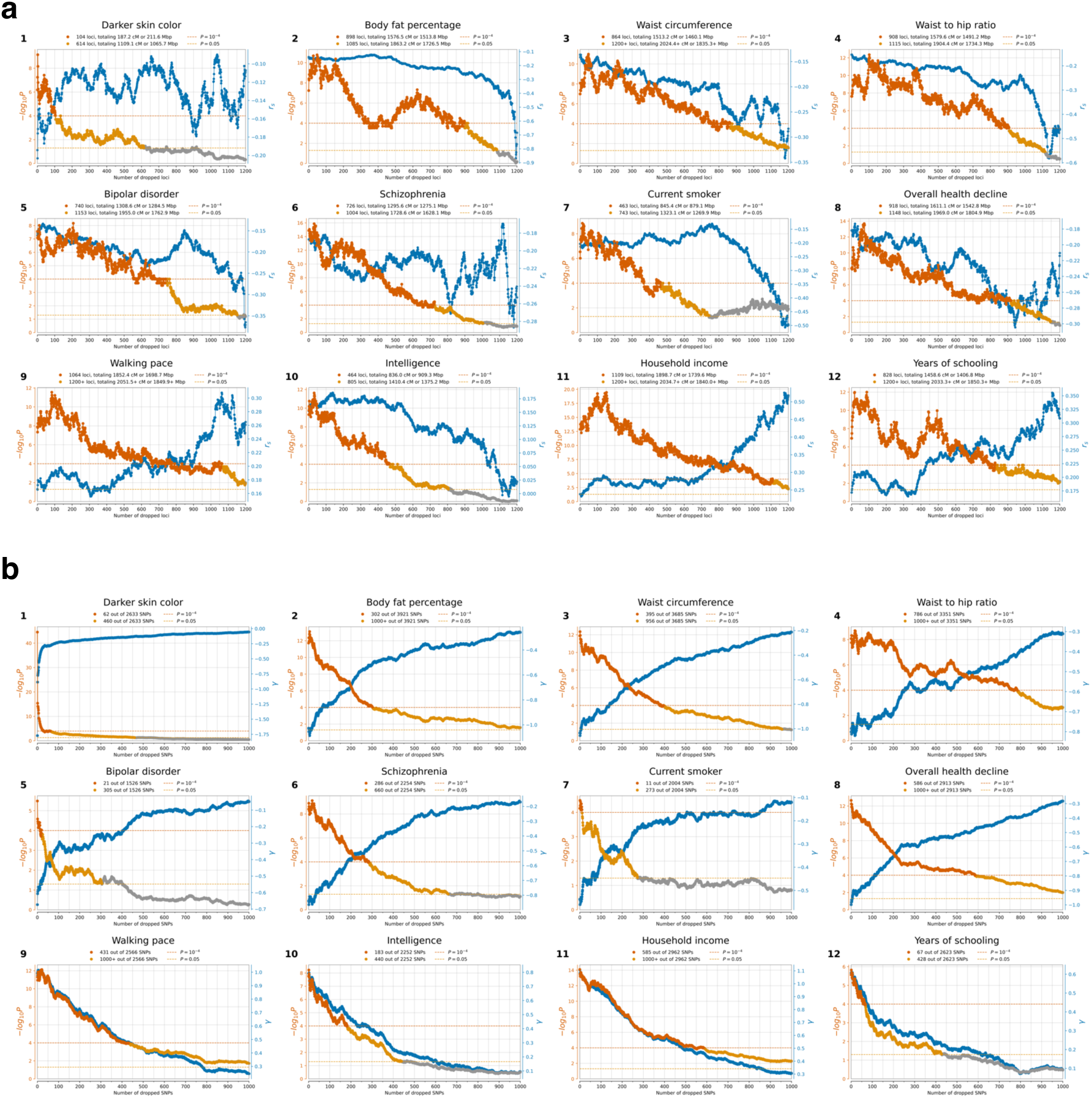
Estimating the minimum number of SNPs affected by selection for each trait (gallery of 12 traits also highlighted in Figure 4). Each panel shows the correlation of a trait with selection summary statistics (*r_s_*) as a function of number of dropped loci. The right axis displays *r_s_* in blue; P-value on the left axis in orange. For each SNP, we define a priority score | β × s × f × (1-f) |, where β is the GWAS effect size, *s* the selection coefficient, and *f* allele frequency. SNPs are sorted by priority score, and in each iteration, a 2cM region around the highest priority SNP is dropped, *r_s_* is recalculated for the remaining genome, and this continues until no SNPs are left. **(b)** We similarly show *γ* estimates at right as a function of number of dropped SNPs (blue), and P-value for polygenic selection at left with dark orange P<0.0001, light orange P<0.05, and gray otherwise.

**Extended Data Figure 11:**
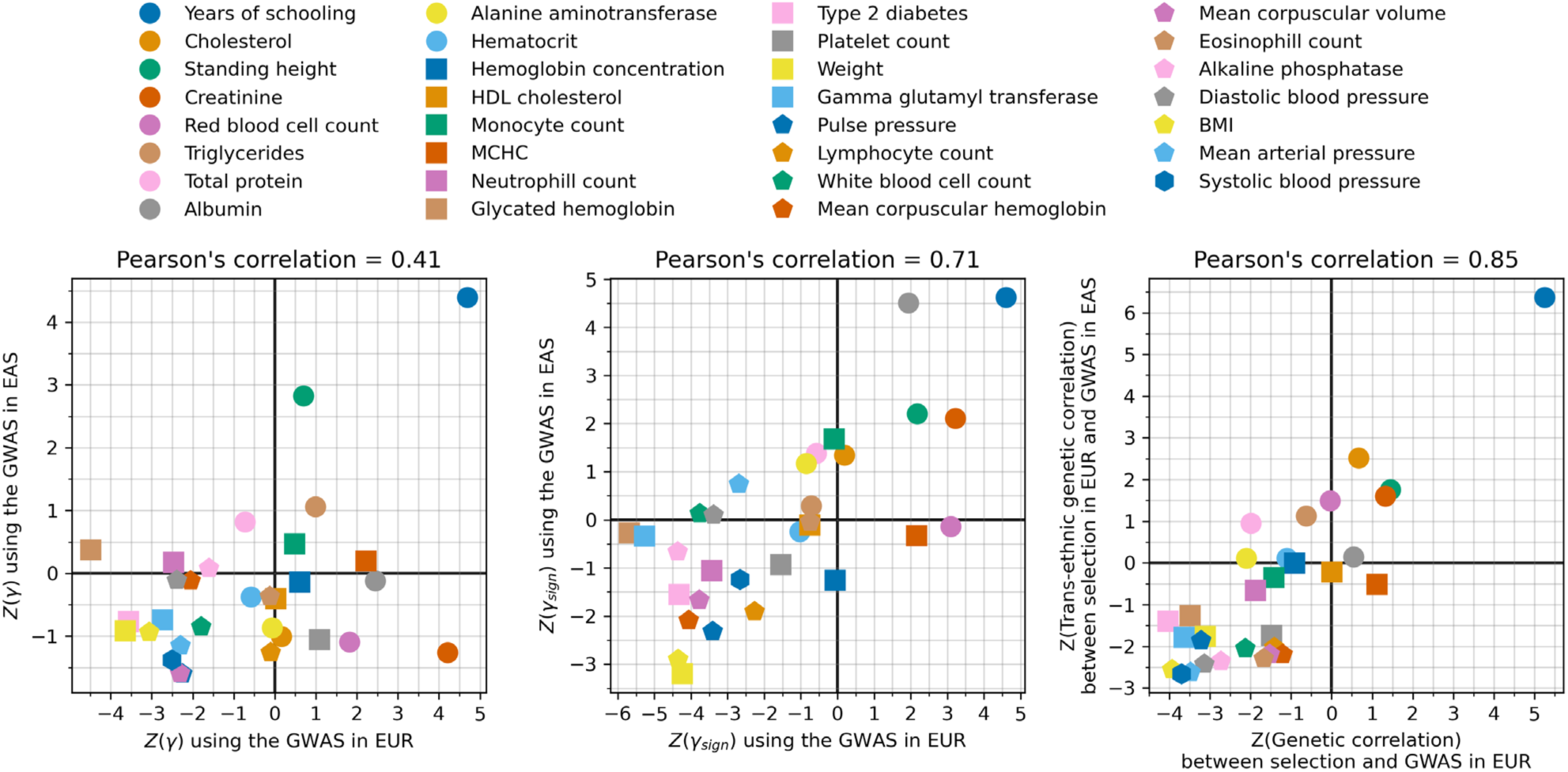
Replication of signals of polygenic selection using effect size estimates in East Asians whose population structure is uncorrelated to West Eurasians. We applied our polygenic selection test to 31 traits using pairs of GWAS studies for the trait, one from Europe and one from East Asia. We assessed if PGS (*γ*), PGS-sign (*γ*_sign_), and genetic correlation tests (*r_s_)* were consistent in these two analyses.

**Extended Data Figure 12:**
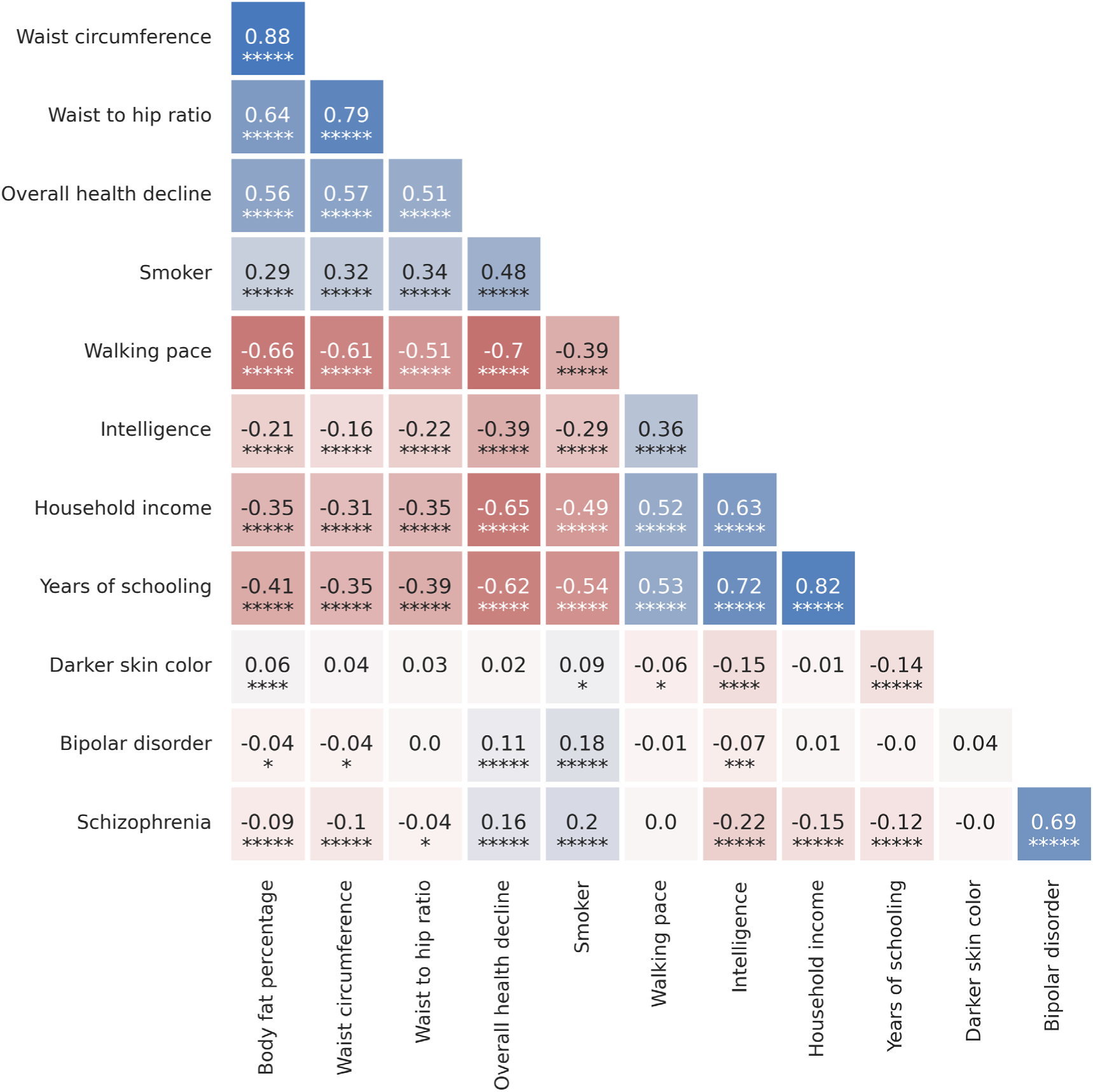
Correlations of polygenic scores for complex traits with strong evidence of coordinated selection (the same 12 traits highlighted in Figure 4). Genetic correlations of traits were computed using LDSC. Asterisks indicate significance level (n asterisks represent a jackknife estimated P<0.5×10^−n^).

**Extended Data Figure 13:**
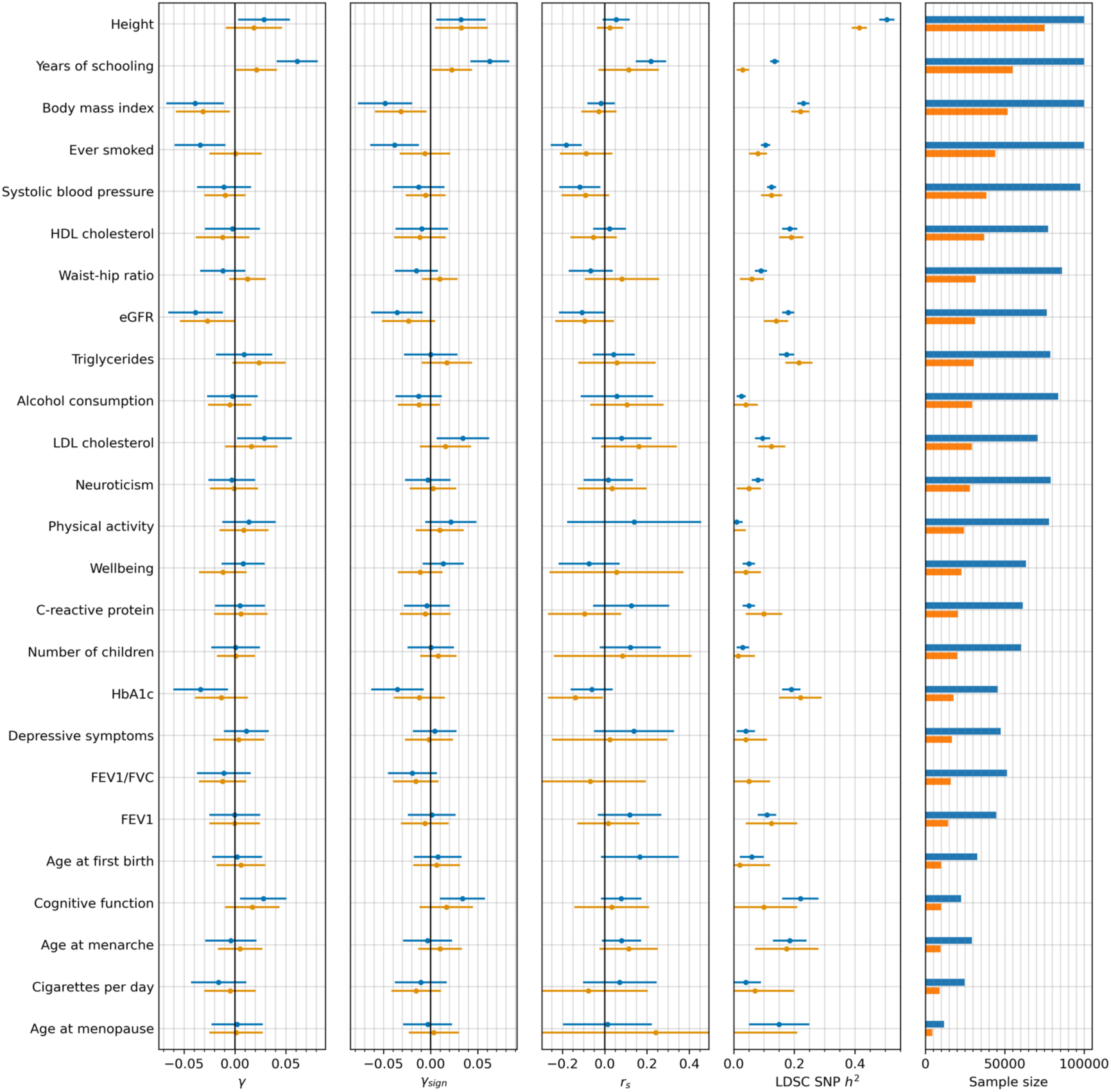
Consistency of polygenic selection signals using effect sizes estimated from both GWAS of unrelated people, and sibling-based GWAS^66^. The first three columns show estimates for each of the three polygenic tests of selection. The fourth column replicates Figure 5 from^66^, and shows the estimated SNP heritability h^2^ by LDSC. The fifth column shows the sample sizes for both GWAS of unrelated people (blue) and sibling-based GWAS (orange). Error bars indicate the 95% confidence interval, which is often larger for the sibling-based GWAS due to limited sample size.

**Extended Data Figure 14:**
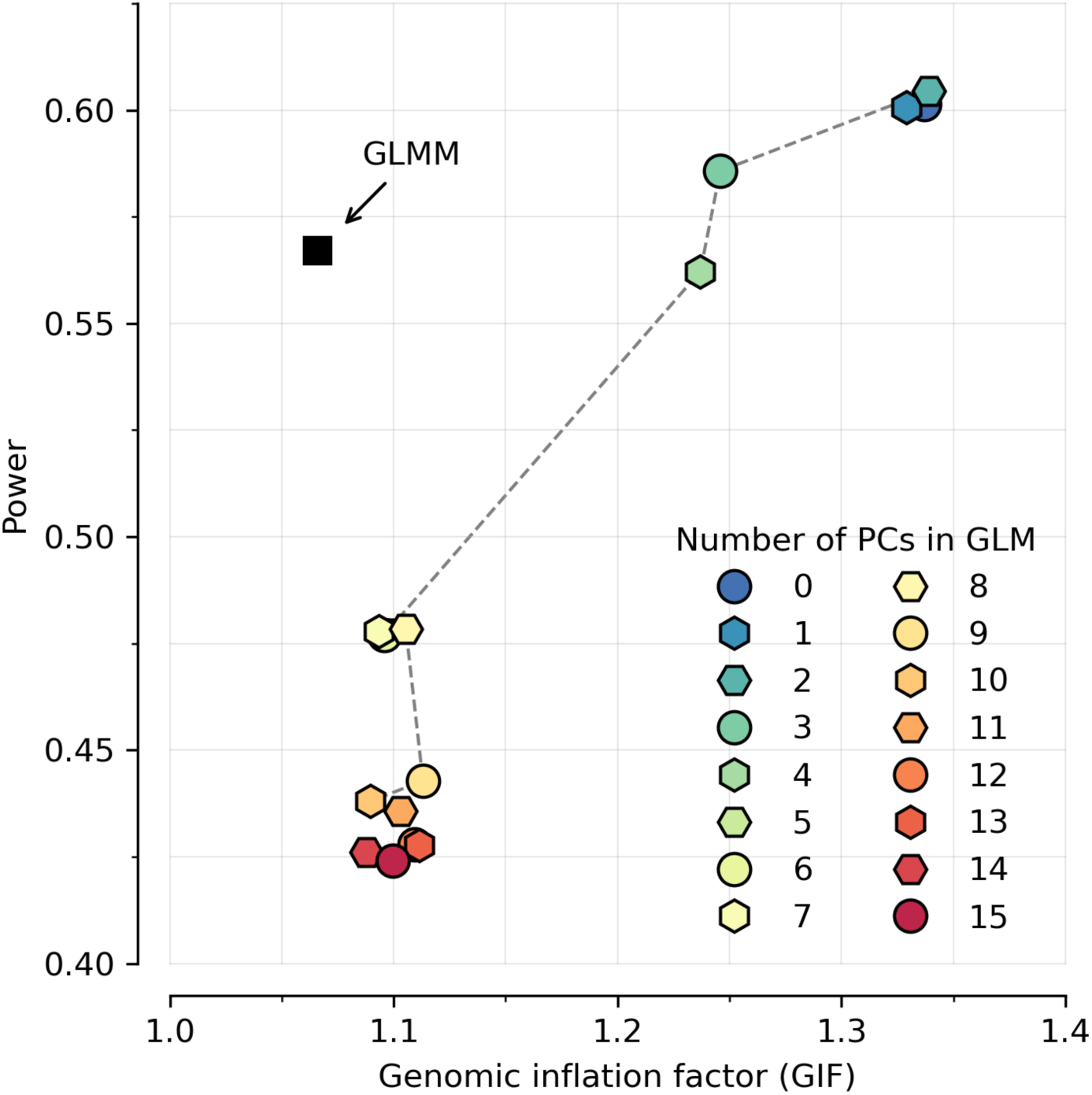
Our generalized linear mixed model (GLMM) method is far more powerful than a generalized linear model (GLM) with PC covariates. To compute the inflation factor for different approaches, we ran 10000 simulations of neutral evolution for a scenario of population structure, sample size, and temporal distribution of samples matching real data. For the power calculation, we ran 20000 simulations of selective sweeps for a range of selection coefficients (power is the proportion of true positives at P<5×10^−8^). Because of co-linearity of time and population structure, correcting for PCs greatly weakens power to detect selection, but the GLMM methodology does not suffer from this.

## Supplementary Data Sets

**Online Table 1: List of 8433 ancient individuals analyzed in our time transect.** Data Source 1 is 3644 individuals whose previously published sequences we reanalyze. Data Source 2 is 244 individuals with previously published in-solution enrichment data for which we report and analyze whole genome shotgun data. Data Source 3 is 74 individuals with previously published in-solution enrichment data for which we report and analyze additional in-solution enrichment data. Data Source 4 is 4471 never-before-reported individuals for which we report and analyze data that is anonymized except for a point estimate of the data of origin and information about broad region in West Eurasia. Available as an Excel table.

**Online Table 2: List of 300 newly reported shotgun ancient genomes.** Most are from individuals with previously reported in-solution enrichment data (n=244; the remainder are from samples reported for the first time (n=56). Available as an Excel table.

**Online Table 3: List of 5382 newly reported ancient DNA libraries.** The majority (n=5227) are from 4471 never-before-reported ancient individuals; the rest (n=155) are from 74 individuals for which we increase data quality. Available as an Excel table.

**Online Table 4: Summary statistics for selection in 9.9 million variants.** The data are provided as a tab-delimited text file, compressed using gzip. Available at Harvard Dataverse: https://doi.org/10.7910/DVN/7RVV9N.

**Online Table 5: Summary statistics for tests of polygenic selection.** The data includes: 452 European GWAS from Pan-UKBB^24^, 107 curated European GWAS used for S-LDSC meta-analysis^55,217^, 50 European GWAS with 25 pairs of sibling and population GWAS from Howe et al. 2022^66^, 30 East Asian GWAS from Biobank Japan, and one from Chen et al. 2024^219^. Available as an Excel table.

## Notes

https://reich-ages.rc.hms.harvard.edu

https://doi.org/10.7910/DVN/7RVV9N

### Competing Interest Statement

The authors have declared no competing interest.

## References

1. Racimo, F., Sikora, M., Vander Linden, M., Schroeder, H. & Lalueza-Fox, C. Beyond broad strokes: sociocultural insights from the study of ancient genomes. Nat. Rev. Genet. 21, 355–366 (2020).

2. Kerner, G., Choin, J. & Quintana-Murci, L. Ancient DNA as a tool for medical research. Nat. Med. 29, 1048–1051 (2023).

3. Bennett, E. A. & Fu, Q. Ancient genomes and the evolutionary path of modern humans. Cell 187, 1042–1046 (2024).

4. Chen, H., Patterson, N. & Reich, D. Population differentiation as a test for selective sweeps. Genome Res. 20, 393–402 (2010).

5. Yi, X. et al. Sequencing of 50 human exomes reveals adaptation to high altitude. Science 329, 75–78 (2010).

6. Sabeti, P. C. et al. Detecting recent positive selection in the human genome from haplotype structure. Nature 419, 832–837 (2002).

7. Voight, B. F., Kudaravalli, S., Wen, X. & Pritchard, J. K. A map of recent positive selection in the human genome. PLoS Biol. 4, e72 (2006).

8. Field, Y. et al. Detection of human adaptation during the past 2000 years. Science 354, 760–764 (2016).

9. Speidel, L., Forest, M., Shi, S. & Myers, S. R. A method for genome-wide genealogy estimation for thousands of samples. Nat. Genet. 51, 1321–1329 (2019).

10. Berg, J. J. & Coop, G. A Coalescent Model for a Sweep of a Unique Standing Variant. Genetics 201, 707–725 (2015).

11. Mathieson, I. et al. Genome-wide patterns of selection in 230 ancient Eurasians. Nature 528, 499–503 (2015).

12. Le, M. K. et al. 1,000 ancient genomes uncover 10,000 years of natural selection in Europe. BioRxiv Prepr. Serv. Biol. 2022.08.24.505188 (2022) doi:10.1101/2022.08.24.505188.

13. Kerner, G. et al. Genetic adaptation to pathogens and increased risk of inflammatory disorders in post-Neolithic Europe. Cell Genomics 3, 100248 (2023).

14. Irving-Pease, E. K. et al. The selection landscape and genetic legacy of ancient Eurasians. Nature 625, 312–320 (2024).

15. Pritchard, J. K., Pickrell, J. K. & Coop, G. The genetics of human adaptation: hard sweeps, soft sweeps, and polygenic adaptation. Curr. Biol. CB 20, R208–215 (2010).

16. Gao, Z. Unveiling recent and ongoing adaptive selection in human populations. PLoS Biol. 22, e3002469 (2024).

17. Mallick, S. et al. The Allen Ancient DNA Resource (AADR) a curated compendium of ancient human genomes. Sci. Data 11, 182 (2024).

18. Byrska-Bishop, M. et al. High-coverage whole-genome sequencing of the expanded 1000 Genomes Project cohort including 602 trios. Cell 185, 3426–3440.e19 (2022).

19. Bycroft, C. et al. The UK Biobank resource with deep phenotyping and genomic data. Nature 562, 203–209 (2018).

20. Rubinacci, S., Ribeiro, D. M., Hofmeister, R. J. & Delaneau, O. Efficient phasing and imputation of low-coverage sequencing data using large reference panels. Nat. Genet. 53, 120–126 (2021).

21. Yang, J. et al. Genomic inflation factors under polygenic inheritance. Eur. J. Hum. Genet. EJHG 19, 807–812 (2011).

22. Devlin, B., Roeder, K. & Wasserman, L. Genomic control, a new approach to genetic-based association studies. Theor. Popul. Biol. 60, 155–166 (2001).

23. Bulik-Sullivan, B. K. et al. LD Score regression distinguishes confounding from polygenicity in genome-wide association studies. Nat. Genet. 47, 291–295 (2015).

24. Karczewski, K. J. et al. Pan-UK Biobank GWAS improves discovery, analysis of genetic architecture, and resolution into ancestry-enriched effects. 2024.03.13.24303864 Preprint at 10.1101/2024.03.13.24303864 (2024).

25. Ronen, R. et al. Predicting Carriers of Ongoing Selective Sweeps without Knowledge of the Favored Allele. PLoS Genet. 11, e1005527 (2015).

26. Akbari, A. et al. Identifying the Favored Mutation in a Positive Selective Sweep. Nat. Methods 15, 279–282 (2018).

27. McVicker, G., Gordon, D., Davis, C. & Green, P. Widespread genomic signatures of natural selection in hominid evolution. PLoS Genet. 5, e1000471 (2009).

28. Murphy, D. A., Elyashiv, E., Amster, G. & Sella, G. Broad-scale variation in human genetic diversity levels is predicted by purifying selection on coding and non-coding elements. eLife 12, e76065 (2023).

29. Finucane, H. K. et al. Partitioning heritability by functional annotation using genome-wide association summary statistics. Nat. Genet. 47, 1228–1235 (2015).

30. Abadie, V., Sollid, L. M., Barreiro, L. B. & Jabri, B. Integration of genetic and immunological insights into a model of celiac disease pathogenesis. Annu. Rev. Immunol. 29, 493–525 (2011).

31. Sams, A. & Hawks, J. Patterns of population differentiation and natural selection on the celiac disease background risk network. PloS One 8, e70564 (2013).

32. Sams, A. & Hawks, J. Celiac disease as a model for the evolution of multifactorial disease in humans. Hum. Biol. 86, 19–36 (2014).

33. Abegaz, S. B. Human ABO Blood Groups and Their Associations with Different Diseases. BioMed Res. Int. 2021, 6629060 (2021).

34. Pettenkofer, H. J., Stoess, B., Helmbold, W. & Vogel, F. Alleged causes of the present-day world distribution of the human ABO blood groups. Nature 193, 445–446 (1962).

35. Medland, S. E. et al. Common Variants in the Trichohyalin Gene Are Associated with Straight Hair in Europeans. Am. J. Hum. Genet. 85, 750–755 (2009).

36. Kerner, G. et al. Homozygosity for TYK2 P1104A underlies tuberculosis in about 1% of patients in a cohort of European ancestry. Proc. Natl. Acad. Sci. U. S. A. 116, 10430–10434 (2019).

37. Kerner, G. et al. Human ancient DNA analyses reveal the high burden of tuberculosis in Europeans over the last 2,000 years. Am. J. Hum. Genet. 108, 517–524 (2021).

38. Barrie, W. et al. Elevated genetic risk for multiple sclerosis emerged in steppe pastoralist populations. Nature 625, 321–328 (2024).

39. Moutsianas, L. et al. Class II HLA interactions modulate genetic risk for multiple sclerosis. Nat. Genet. 47, 1107–1113 (2015).

40. Moalem, S., Percy, M. E., Kruck, T. P. A. & Gelbart, R. R. Epidemic pathogenic selection: an explanation for hereditary hemochromatosis? Med. Hypotheses 59, 325–329 (2002).

41. Bos, K. I. et al. A draft genome of Yersinia pestis from victims of the Black Death. Nature 478, 506–510 (2011).

42. Wagner, D. M. et al. Yersinia pestis and the plague of Justinian 541-543 AD: a genomic analysis. Lancet Infect. Dis. 14, 319–326 (2014).

43. Samson, M. et al. Resistance to HIV-1 infection in caucasian individuals bearing mutant alleles of the CCR-5 chemokine receptor gene. Nature 382, 722–725 (1996).

44. Hütter, G. et al. Long-term control of HIV by CCR5 Delta32/Delta32 stem-cell transplantation. N. Engl. J. Med. 360, 692–698 (2009).

45. Gupta, R. K. et al. HIV-1 remission following CCR5Δ32/Δ32 haematopoietic stem-cell transplantation. Nature 568, 244–248 (2019).

46. Stephens, J. C. et al. Dating the origin of the CCR5-Delta32 AIDS-resistance allele by the coalescence of haplotypes. Am. J. Hum. Genet. 62, 1507–1515 (1998).

47. Novembre, J., Galvani, A. P. & Slatkin, M. The geographic spread of the CCR5 Delta32 HIV-resistance allele. PLoS Biol. 3, e339 (2005).

48. Sabeti, P. C. et al. The Case for Selection at CCR5-Δ32. PLoS Biol. 3, e378 (2005).

49. Seersholm, F. V. et al. Repeated plague infections across six generations of Neolithic Farmers. Nature 1–8 (2024) doi:10.1038/s41586-024-07651-2.

50. Rasmussen, S. et al. Early divergent strains of Yersinia pestis in Eurasia 5,000 years ago. Cell 163, 571–582 (2015).

51. Spyrou, M. A. et al. Analysis of 3800-year-old Yersinia pestis genomes suggests Bronze Age origin for bubonic plague. Nat. Commun. 9, 2234 (2018).

52. Kerem, B. et al. Identification of the cystic fibrosis gene: genetic analysis. Science 245, 1073–1080 (1989).

53. Cutting, G. R. Cystic fibrosis genetics: from molecular understanding to clinical application. Nat. Rev. Genet. 16, 45–56 (2015).

54. Gabriel, S. E., Brigman, K. N., Koller, B. H., Boucher, R. C. & Stutts, M. J. Cystic fibrosis heterozygote resistance to cholera toxin in the cystic fibrosis mouse model. Science 266, 107–109 (1994).

55. Kim, A. et al. Inferring causal cell types of human diseases and risk variants from candidate regulatory elements. 2024.05.17.24307556 Preprint at 10.1101/2024.05.17.24307556 (2024).

56. Chen, M. et al. Evidence of Polygenic Adaptation in Sardinia at Height-Associated Loci Ascertained from the Biobank Japan. Am. J. Hum. Genet. 107, 60–71 (2020).

57. Ju, D. & Mathieson, I. The evolution of skin pigmentation-associated variation in West Eurasia. Proc. Natl. Acad. Sci. U. S. A. 118, e2009227118 (2021).

58. Ding, Y. et al. Polygenic scoring accuracy varies across the genetic ancestry continuum. Nature 618, 774–781 (2023).

59. Bulik-Sullivan, B. et al. An atlas of genetic correlations across human diseases and traits. Nat. Genet. 47, 1236–1241 (2015).

60. Berg, J. J. et al. Reduced signal for polygenic adaptation of height in UK Biobank. eLife 8, e39725 (2019).

61. Sohail, M. et al. Polygenic adaptation on height is overestimated due to uncorrected stratification in genome-wide association studies. eLife 8, e39702 (2019).

62. O’Connor, L. J. et al. Extreme Polygenicity of Complex Traits Is Explained by Negative Selection. Am. J. Hum. Genet. 105, 456–476 (2019).

63. Beauchamp, J. P. Genetic evidence for natural selection in humans in the contemporary United States. Proc. Natl. Acad. Sci. U. S. A. 113, 7774–7779 (2016).

64. Kong, A. et al. Selection against variants in the genome associated with educational attainment. Proc. Natl. Acad. Sci. U. S. A. 114, E727–E732 (2017).

65. Sanjak, J. S., Sidorenko, J., Robinson, M. R., Thornton, K. R. & Visscher, P. M. Evidence of directional and stabilizing selection in contemporary humans. Proc. Natl. Acad. Sci. U. S. A. 115, 151–156 (2018).

66. Howe, L. J. et al. Within-sibship genome-wide association analyses decrease bias in estimates of direct genetic effects. Nat. Genet. 54, 581–592 (2022).

67. Hernandez, R. D. et al. Classic selective sweeps were rare in recent human evolution. Science 331, 920–924 (2011).

68. Coop, G. et al. The role of geography in human adaptation. PLoS Genet. 5, e1000500 (2009).

69. Simon, A. & Coop, G. The contribution of gene flow, selection, and genetic drift to five thousand years of human allele frequency change. bioRxiv 2023.07.11.548607 (2024) doi:10.1101/2023.07.11.548607.

70. Koch, E. et al. Genetic association data are broadly consistent with stabilizing selection shaping human common diseases and traits. 2024.06.19.599789 Preprint at 10.1101/2024.06.19.599789 (2024).

71. Yang, J., Zaitlen, N. A., Goddard, M. E., Visscher, P. M. & Price, A. L. Advantages and pitfalls in the application of mixed-model association methods. Nat. Genet. 46, 100–106 (2014).

72. Taus, T., Futschik, A. & Schlötterer, C. Quantifying Selection with Pool-Seq Time Series Data. Mol. Biol. Evol. 34, 3023–3034 (2017).

73. Crow, J. F. An Introduction to Population Genetics Theory. (Scientific Publishers, 2017).

74. Sun, S. et al. Heritability estimation and differential analysis of count data with generalized linear mixed models in genomic sequencing studies. Bioinforma. Oxf. Engl. 35, 487–496 (2019).

75. Loh, P.-R. et al. Efficient Bayesian mixed model analysis increases association power in large cohorts. Nat. Genet. 47, 284–290 (2015).

76. Listgarten, J. et al. Improved linear mixed models for genome-wide association studies. Nat. Methods 9, 525–526 (2012).

77. Zhou, X. & Stephens, M. Genome-wide efficient mixed-model analysis for association studies. Nat. Genet. 44, 821–824 (2012).

78. Shi, H. et al. Population-specific causal disease effect sizes in functionally important regions impacted by selection. Nat. Commun. 12, 1098 (2021).

79. Gazal, S. et al. Linkage disequilibrium-dependent architecture of human complex traits shows action of negative selection. Nat. Genet. 49, 1421–1427 (2017).

80. Gazal, S., Marquez-Luna, C., Finucane, H. K. & Price, A. L. Reconciling S-LDSC and LDAK functional enrichment estimates. Nat. Genet. 51, 1202–1204 (2019).

81. Damgaard, P. D. B. et al. 137 ancient human genomes from across the Eurasian steppes. Nature 557, 369–374 (2018).

82. Jensen, T. Z. T. et al. A 5700 year-old human genome and oral microbiome from chewed birch pitch. Nat. Commun. 10, 5520 (2019).

83. Olalde, I. et al. A Common Genetic Origin for Early Farmers from Mediterranean Cardial and Central European LBK Cultures. Mol. Biol. Evol. msv181 (2015) doi:10.1093/molbev/msv181.

84. Cassidy, L. M. et al. A dynastic elite in monumental Neolithic society. Nature 582, 384–388 (2020).

85. Moots, H. M. et al. A genetic history of continuity and mobility in the Iron Age central Mediterranean. Nat. Ecol. Evol. 7, 1515–1524 (2023).

86. Olalde, I. et al. A genetic history of the Balkans from Roman frontier to Slavic migrations. Cell 186, 5472–5485.e9 (2023).

87. Nikitin, A. G. et al. A genomic history of the North Pontic Region from the Neolithic to the Bronze Age. Preprint at 10.1101/2024.04.17.589600 (2024).

88. Fernandes, D. M. et al. A genomic Neolithic time transect of hunter-farmer admixture in central Poland. Sci. Rep. 8, 14879 (2018).

89. Fowler, C. et al. A high-resolution picture of kinship practices in an Early Neolithic tomb. Nature 601, 584–587 (2022).

90. Harney, É. et al. A minimally destructive protocol for DNA extraction from ancient teeth. Genome Res. 31, 472–483 (2021).

91. González-Fortes, G. et al. A western route of prehistoric human migration from Africa into the Iberian Peninsula. Proc. R. Soc. B Biol. Sci. 286, 20182288 (2019).

92. Marciniak, S., et al. An integrative skeletal and paleogenomic analysis of stature variation suggests relatively reduced health for early European farmers. Proc. Natl. Acad. Sci. 119, e2106743119 (2022).

93. Unterländer, M. et al. Ancestry and demography and descendants of Iron Age nomads of the Eurasian Steppe. Nat. Commun. 8, 14615 (2017).

94. Dulias, K., et al. Ancient DNA at the edge of the world: Continental immigration and the persistence of Neolithic male lineages in Bronze Age Orkney. Proc. Natl. Acad. Sci. 119, e2108001119 (2022).

95. Zalloua, P. et al. Ancient DNA of Phoenician remains indicates discontinuity in the settlement history of Ibiza. Sci. Rep. 8, 17567 (2018).

96. Skourtanioti, E. et al. Ancient DNA reveals admixture history and endogamy in the prehistoric Aegean. Nat. Ecol. Evol. 7, 290–303 (2023).

97. Lamnidis, T. C. et al. Ancient Fennoscandian genomes reveal origin and spread of Siberian ancestry in Europe. Nat. Commun. 9, 5018 (2018).

98. O’Sullivan, N. et al. Ancient genome-wide analyses infer kinship structure in an Early Medieval Alemannic graveyard. Sci. Adv. 4, eaao1262 (2018).

99. Rivollat, M. et al. Ancient genome-wide DNA from France highlights the complexity of interactions between Mesolithic hunter-gatherers and Neolithic farmers. Sci. Adv. 6, eaaz5344 (2020).

100. Ebenesersdóttir, S. S. et al. Ancient genomes from Iceland reveal the making of a human population. Science 360, 1028–1032 (2018).

101. Fregel, R. et al. Ancient genomes from North Africa evidence prehistoric migrations to the Maghreb from both the Levant and Europe. Proc. Natl. Acad. Sci. 115, 6774–6779 (2018).

102. Brunel, S. et al. Ancient genomes from present-day France unveil 7,000 years of its demographic history. Proc. Natl. Acad. Sci. 117, 12791–12798 (2020).

103. Brace, S. et al. Ancient genomes indicate population replacement in Early Neolithic Britain. Nat. Ecol. Evol. 3, 765–771 (2019).

104. Günther, T. et al. Ancient genomes link early farmers from Atapuerca in Spain to modern-day Basques. Proc. Natl. Acad. Sci. 112, 11917–11922 (2015).

105. Žegarac, A., et al. Ancient genomes provide insights into family structure and the heredity of social status in the early Bronze Age of southeastern Europe. Sci. Rep. 11, 10072 (2021).

106. Gnecchi-Ruscone, G. A. et al. Ancient genomes reveal origin and rapid trans-Eurasian migration of 7th century Avar elites. Cell 185, 1402–1413.e21 (2022).

107. Saupe, T. et al. Ancient genomes reveal structural shifts after the arrival of Steppe-related ancestry in the Italian Peninsula. Curr. Biol. 31, 2576–2591.e12 (2021).

108. Sikora, M. et al. Ancient genomes show social and reproductive behavior of early Upper Paleolithic foragers. Science 358, 659–662 (2017).

109. Krzewińska, M. et al. Ancient genomes suggest the eastern Pontic-Caspian steppe as the source of western Iron Age nomads. Sci. Adv. 4, eaat4457 (2018).

110. Gnecchi-Ruscone, G. A. et al. Ancient genomic time transect from the Central Asian Steppe unravels the history of the Scythians. Sci. Adv. 7, eabe4414 (2021).

111. Wang, C.-C. et al. Ancient human genome-wide data from a 3000-year interval in the Caucasus corresponds with eco-geographic regions. Nat. Commun. 10, 590 (2019).

112. Lazaridis, I. et al. Ancient human genomes suggest three ancestral populations for present-day Europeans. Nature 513, 409–413 (2014).

113. Ariano, B. et al. Ancient Maltese genomes and the genetic geography of Neolithic Europe. Curr. Biol. 32, 2668–2680.e6 (2022).

114. Antonio, M. L. et al. Ancient Rome: A genetic crossroads of Europe and the Mediterranean. Science 366, 708–714 (2019).

115. Scorrano, G. et al. Bioarchaeological and palaeogenomic portrait of two Pompeians that died during the eruption of Vesuvius in 79 AD. Sci. Rep. 12, 6468 (2022).

116. Srigyan, M. et al. Bioarchaeological evidence of one of the earliest Islamic burials in the Levant. Commun. Biol. 5, 554 (2022).

117. Furtwängler, A. et al. Comparison of Target Enrichment Strategies for Ancient Pathogen DNA. BioTechniques 69, 455–459 (2020).

118. Linderholm, A. et al. Corded Ware cultural complexity uncovered using genomic and isotopic analysis from south-eastern Poland. Sci. Rep. 10, 6885 (2020).

119. Gokhman, D. et al. Differential DNA methylation of vocal and facial anatomy genes in modern humans. Nat. Commun. 11, 1189 (2020).

120. Papac, L. et al. Dynamic changes in genomic and social structures in third millennium BCE central Europe. Sci. Adv. 7, eabi6941 (2021).

121. Hofmanová, Z. et al. Early farmers from across Europe directly descended from Neolithic Aegeans. Proc. Natl. Acad. Sci. 113, 6886–6891 (2016).

122. Broushaki, F. et al. Early Neolithic genomes from the eastern Fertile Crescent. Science 353, 499–503 (2016).

123. Scheib, C. L. et al. East Anglian early Neolithic monument burial linked to contemporary Megaliths. Ann. Hum. Biol. 46, 145–149 (2019).

124. Saag, L. et al. Extensive Farming in Estonia Started through a Sex-Biased Migration from the Steppe. Curr. Biol. 27, 2185–2193.e6 (2017).

125. Valdiosera, C. et al. Four millennia of Iberian biomolecular prehistory illustrate the impact of prehistoric migrations at the far end of Eurasia. Proc. Natl. Acad. Sci. 115, 3428–3433 (2018).

126. Peltola, S. et al. Genetic admixture and language shift in the medieval Volga-Oka interfluve. Curr. Biol. 33, 174–182.e10 (2023).

127. Saag, L. et al. Genetic ancestry changes in Stone to Bronze Age transition in the East European plain. Sci. Adv. 7, eabd6535 (2021).

128. Marcus, J. H. et al. Genetic history from the Middle Neolithic to present on the Mediterranean island of Sardinia. Nat. Commun. 11, 939 (2020).

129. Lazaridis, I. et al. Genetic origins of the Minoans and Mycenaeans. Nature 548, 214–218 (2017).

130. Gamba, C. et al. Genome flux and stasis in a five millennium transect of European prehistory. Nat. Commun. 5, 5257 (2014).

131. Novak, M. et al. Genome-wide analysis of nearly all the victims of a 6200 year old massacre. PLOS ONE 16, e0247332 (2021).

132. Waldman, S. et al. Genome-wide data from medieval German Jews show that the Ashkenazi founder event pre-dated the 14th century. Cell 185, 4703–4716.e16 (2022).

133. Immel, A. et al. Genome-wide study of a Neolithic Wartberg grave community reveals distinct HLA variation and hunter-gatherer ancestry. Commun. Biol. 4, 113 (2021).

134. Brace, S. et al. Genomes from a medieval mass burial show Ashkenazi-associated hereditary diseases pre-date the 12th century. Curr. Biol. 32, 4350–4359.e6 (2022).

135. Gelabert, P. et al. Genomes from Verteba cave suggest diversity within the Trypillians in Ukraine. Sci. Rep. 12, 7242 (2022).

136. Yu, H. et al. Genomic and dietary discontinuities during the Mesolithic and Neolithic in Sicily. iScience 25, 104244 (2022).

137. Krzewińska, M. et al. Genomic and Strontium Isotope Variation Reveal Immigration Patterns in a Viking Age Town. Curr. Biol. 28, 2730–2738.e10 (2018).

138. Skoglund, P. et al. Genomic Diversity and Admixture Differs for Stone-Age Scandinavian Foragers and Farmers. Science 344, 747–750 (2014).

139. Skourtanioti, E. et al. Genomic History of Neolithic to Bronze Age Anatolia, Northern Levant, and Southern Caucasus. Cell 181, 1158–1175.e28 (2020).

140. Lazaridis, I. et al. Genomic insights into the origin of farming in the ancient Near East. Nature 536, 419–424 (2016).

141. Martiniano, R. et al. Genomic signals of migration and continuity in Britain before the Anglo-Saxons. Nat. Commun. 7, 10326 (2016).

142. Villalba-Mouco, V. et al. Genomic transformation and social organization during the Copper Age–Bronze Age transition in southern Iberia. Sci. Adv. 7, eabi7038 (2021).

143. Seguin-Orlando, A. et al. Heterogeneous Hunter-Gatherer and Steppe-Related Ancestries in Late Neolithic and Bell Beaker Genomes from Present-Day France. Curr. Biol. 31, 1072–1083.e10 (2021).

144. Ingman, T. et al. Human mobility at Tell Atchana (Alalakh), Hatay, Turkey during the 2nd millennium BC: Integration of isotopic and genomic evidence. PLOS ONE 16, e0241883 (2021).

145. Schiffels, S. et al. Iron Age and Anglo-Saxon genomes from East England reveal British migration history. Nat. Commun. 7, 10408 (2016).

146. Armit, I. et al. Kinship practices in Early Iron Age South-east Europe: genetic and isotopic analysis of burials from the Dolge njive barrow cemetery, Dolenjska, Slovenia. Antiquity 97, 403–418 (2023).

147. Mittnik, A. et al. Kinship-based social inequality in Bronze Age Europe. Science 366, 731–734 (2019).

148. Patterson, N. et al. Large-scale migration into Britain during the Middle to Late Bronze Age. Nature 601, 588–594 (2022).

149. Feldman, M. et al. Late Pleistocene human genome suggests a local origin for the first farmers of central Anatolia. Nat. Commun. 10, 1218 (2019).

150. Burger, J. et al. Low Prevalence of Lactase Persistence in Bronze Age Europe Indicates Ongoing Strong Selection over the Last 3,000 Years. Curr. Biol. 30, 4307–4315.e13 (2020).

151. Sánchez-Quinto, F. et al. Megalithic tombs in western and northern Neolithic Europe were linked to a kindred society. Proc. Natl. Acad. Sci. 116, 9469–9474 (2019).

152. Cassidy, L. M. et al. Neolithic and Bronze Age migration to Ireland and establishment of the insular Atlantic genome. Proc. Natl. Acad. Sci. 113, 368–373 (2016).

153. Fischer, C.-E. et al. Origin and mobility of Iron Age Gaulish groups in present-day France revealed through archaeogenomics. iScience 25, 104094 (2022).

154. Posth, C. et al. Palaeogenomics of Upper Palaeolithic to Neolithic European hunter-gatherers. Nature 615, 117–126 (2023).

155. González-Fortes, G. et al. Paleogenomic Evidence for Multi-generational Mixing between Neolithic Farmers and Mesolithic Hunter-Gatherers in the Lower Danube Basin. Curr. Biol. 27, 1801–1810.e10 (2017).

156. Lipson, M. et al. Parallel palaeogenomic transects reveal complex genetic history of early European farmers. Nature 551, 368–372 (2017).

157. Childebayeva, A. et al. Population Genetics and Signatures of Selection in Early Neolithic European Farmers. Mol. Biol. Evol. 39, msac108 (2022).

158. Veeramah, K. R. et al. Population genomic analysis of elongated skulls reveals extensive female-biased immigration in Early Medieval Bavaria. Proc. Natl. Acad. Sci. 115, 3494–3499 (2018).

159. Allentoft, M. E. et al. Population genomics of Bronze Age Eurasia. Nature 522, 167–172 (2015).

160. Günther, T. et al. Population genomics of Mesolithic Scandinavia: Investigating early postglacial migration routes and high-latitude adaptation. PLOS Biol. 16, e2003703 (2018).

161. Margaryan, A. et al. Population genomics of the Viking world. Nature 585, 390–396 (2020).

162. Zeng, T. C. et al. Postglacial genomes from foragers across Northern Eurasia reveal prehistoric mobility associated with the spread of the Uralic and Yeniseian languages. Preprint at 10.1101/2023.10.01.560332 (2023).

163. Freilich, S. et al. Reconstructing genetic histories and social organisation in Neolithic and Bronze Age Croatia. Sci. Rep. 11, 16729 (2021).

164. Järve, M. et al. Shifts in the Genetic Landscape of the Western Eurasian Steppe Associated with the Beginning and End of the Scythian Dominance. Curr. Biol. 29, 2430–2441.e10 (2019).

165. Gelabert, P. et al. Social and genetic diversity among the first farmers of Central Europe. Preprint at 10.1101/2023.07.07.548126 (2023).

166. Koptekin, D. et al. Spatial and temporal heterogeneity in human mobility patterns in Holocene Southwest Asia and the East Mediterranean. Curr. Biol. 33, 41–57.e15 (2023).

167. Antonio, M. L. et al. Stable population structure in Europe since the Iron Age, despite high mobility. eLife 13, e79714 (2024).

168. Villalba-Mouco, V. et al. Survival of Late Pleistocene Hunter-Gatherer Ancestry in the Iberian Peninsula. Curr. Biol. 29, 1169–1177.e7 (2019).

169. Gretzinger, J. et al. The Anglo-Saxon migration and the formation of the early English gene pool. Nature 610, 112–119 (2022).

170. Saag, L. et al. The Arrival of Siberian Ancestry Connecting the Eastern Baltic to Uralic Speakers further East. Curr. Biol. 29, 1701–1711.e16 (2019).

171. Olalde, I. et al. The Beaker phenomenon and the genomic transformation of northwest Europe. Nature 555, 190–196 (2018).

172. Kılınç, G. M. et al. The Demographic Development of the First Farmers in Anatolia. Curr. Biol. 26, 2659–2666 (2016).

173. Reitsema, L. J., et al. The diverse genetic origins of a Classical period Greek army. Proc. Natl. Acad. Sci. 119, e2205272119 (2022).

174. De Barros Damgaard, P., et al. The first horse herders and the impact of early Bronze Age steppe expansions into Asia. Science 360, eaar7711 (2018).

175. Narasimhan, V. M. et al. The formation of human populations in South and Central Asia. Science 365, eaat7487 (2019).

176. Fu, Q. et al. The genetic history of Ice Age Europe. Nature 534, 200–205 (2016).

177. Rodríguez-Varela, R. et al. The genetic history of Scandinavia from the Roman Iron Age to the present. Cell 186, 32–46.e19 (2023).

178. Lazaridis, I. et al. The genetic history of the Southern Arc: A bridge between West Asia and Europe. Science 377, eabm4247 (2022).

179. Aneli, S. et al. The Genetic Origin of Daunians and the Pan-Mediterranean Southern Italian Iron Age Context. Mol. Biol. Evol. 39, msac014 (2022).

180. Maróti, Z. et al. The genetic origin of Huns, Avars, and conquering Hungarians. Curr. Biol. 32, 2858–2870.e7 (2022).

181. Lazaridis, I. et al. The Genetic Origin of the Indo-Europeans. Preprint at 10.1101/2024.04.17.589597 (2024).

182. Mittnik, A. et al. The genetic prehistory of the Baltic Sea region. Nat. Commun. 9, 442 (2018).

183. Malmström, H. et al. The genomic ancestry of the Scandinavian Battle Axe Culture people and their relation to the broader Corded Ware horizon. Proc. R. Soc. B Biol. Sci. 286, 20191528 (2019).

184. Mathieson, I. et al. The genomic history of southeastern Europe. Nature 555, 197–203 (2018).

185. Clemente, F. et al. The genomic history of the Aegean palatial civilizations. Cell 184, 2565–2586.e21 (2021).

186. Olalde, I. et al. The genomic history of the Iberian Peninsula over the past 8000 years. Science 363, 1230–1234 (2019).

187. Marchi, N. et al. The genomic origins of the world’s first farmers. Cell 185, 1842–1859.e18 (2022).

188. Coutinho, A. et al. The Neolithic Pitted Ware culture foragers were culturally but not genetically influenced by the Battle Axe culture herders. Am. J. Phys. Anthropol. 172, 638–649 (2020).

189. Jones, E. R. et al. The Neolithic Transition in the Baltic Was Not Driven by Admixture with Early European Farmers. Curr. Biol. 27, 576–582 (2017).

190. Posth, C. et al. The origin and legacy of the Etruscans through a 2000-year archeogenomic time transect. Sci. Adv. 7, eabi7673 (2021).

191. Martiniano, R. et al. The population genomics of archaeological transition in west Iberia: Investigation of ancient substructure using imputation and haplotype-based methods. PLOS Genet. 13, e1006852 (2017).

192. Fernandes, D. M. et al. The spread of steppe and Iranian-related ancestry in the islands of the western Mediterranean. Nat. Ecol. Evol. 4, 334–345 (2020).

193. Rohland, N. et al. Three assays for in-solution enrichment of ancient human DNA at more than a million SNPs. Genome Res. 32, 2068–2078 (2022).

194. Amorim, C. E. G. et al. Understanding 6th-century barbarian social organization and migration through paleogenomics. Nat. Commun. 9, 3547 (2018).

195. Schroeder, H. et al. Unraveling ancestry, kinship, and violence in a Late Neolithic mass grave. Proc. Natl. Acad. Sci. 116, 10705–10710 (2019).

196. Jones, E. R. et al. Upper Palaeolithic genomes reveal deep roots of modern Eurasians. Nat. Commun. 6, 8912 (2015).

197. Meyer, M. et al. A High-Coverage Genome Sequence from an Archaic Denisovan Individual. Science 338, 222–226 (2012).

198. Raghavan, M. et al. Genomic evidence for the Pleistocene and recent population history of Native Americans. Science 349, aab3884 (2015).

199. Prüfer, K. et al. The complete genome sequence of a Neanderthal from the Altai Mountains. Nature 505, 43–49 (2014).

200. Mallick, S. et al. The Simons Genome Diversity Project: 300 genomes from 142 diverse populations. Nature 538, 201–206 (2016).

201. Raghavan, M. et al. Upper Palaeolithic Siberian genome reveals dual ancestry of Native Americans. Nature 505, 87–91 (2014).

202. Dabney, J. et al. Complete mitochondrial genome sequence of a Middle Pleistocene cave bear reconstructed from ultrashort DNA fragments. Proc. Natl. Acad. Sci. 110, 15758–15763 (2013).

203. Korlević, P. et al. Reducing Microbial and Human Contamination in DNA Extractions from Ancient Bones and Teeth. BioTechniques 59, 87–93 (2015).

204. Rohland, N., Harney, E., Mallick, S., Nordenfelt, S. & Reich, D. Partial uracil-DNA-glycosylase treatment for screening of ancient DNA. Philos. Trans. R. Soc. Lond. B. Biol. Sci. 370, 20130624 (2015).

205. Briggs, A. W. & Heyn, P. Preparation of next-generation sequencing libraries from damaged DNA. Methods Mol. Biol. Clifton NJ 840, 143–154 (2012).

206. Fu, Q. et al. DNA analysis of an early modern human from Tianyuan Cave, China. Proc. Natl. Acad. Sci. U. S. A. 110, 2223–2227 (2013).

207. Haak, W. et al. Massive migration from the steppe was a source for Indo-European languages in Europe. Nature 522, 207–211 (2015).

208. Rohland, N., Glocke, I., Aximu-Petri, A. & Meyer, M. Extraction of highly degraded DNA from ancient bones, teeth and sediments for high-throughput sequencing. Nat. Protoc. 13, 2447–2461 (2018).

209. Gansauge, M.-T., Aximu-Petri, A., Nagel, S. & Meyer, M. Manual and automated preparation of single-stranded DNA libraries for the sequencing of DNA from ancient biological remains and other sources of highly degraded DNA. Nat. Protoc. 15, 2279–2300 (2020).

210. Li, H. & Durbin, R. Fast and accurate short read alignment with Burrows-Wheeler transform. Bioinforma. Oxf. Engl. 25, 1754–1760 (2009).

211. Picard toolkit. Broad Institute, GitHub repository (2019).

212. Behar, D. M. et al. A “Copernican” Reassessment of the Human Mitochondrial DNA Tree from its Root. Am. J. Hum. Genet. 90, 675–684 (2012).

213. 1000 Genomes Project Consortium et al. A global reference for human genetic variation. Nature 526, 68–74 (2015).

214. Danecek, P. et al. Twelve years of SAMtools and BCFtools. GigaScience 10, giab008 (2021).

215. Zhao, H. et al. CrossMap: a versatile tool for coordinate conversion between genome assemblies. Bioinforma. Oxf. Engl. 30, 1006–1007 (2014).

216. Karczewski, K. J. et al. The mutational constraint spectrum quantified from variation in 141,456 humans. Nature 581, 434–443 (2020).

217. Gazal, S. S-LDSC reference files. Zenodo 10.5281/zenodo.10515792 (2024).

218. Sakaue, S. et al. A cross-population atlas of genetic associations for 220 human phenotypes. Nat. Genet. 53, 1415–1424 (2021).

219. Chen, T.-T. et al. Shared genetic architectures of educational attainment in East Asian and European populations. Nat. Hum. Behav. 8, 562–575 (2024).

220. Shelton, J. F. et al. Trans-ancestry analysis reveals genetic and nongenetic associations with COVID-19 susceptibility and severity. Nat. Genet. 53, 801–808 (2021).

