## Supplementary Information sections 1-7 for "Pervasive findings of directional selection realize the promise of ancient DNA to elucidate human adaptation"

### Supplementary Information section 1
### Quality Control (QC)

The data analyzed in this study come from multiple sources: imputed ancient DNA sequences (shotgun sequences and 1240k enrichment reagent), sequences of people of European ancestry from the 1000 Genomes Project, and imputed SNP array data from the UK Biobank (genotyped using the UK Biobank Axiom Array). To generate datasets useful for filtering out variants that do not have reliable genotyping properties, we used principal components to identify groups of individuals with similar ancestry across different datasets. We then filtered out variants whose allele frequencies differed significantly between sample sets to minimize batch effects due to combining samples from different sources. Variant quality control involves a three-step procedure.

#### Variant QC: Step 1

We performed a first step QC that restricted to the 1240k sites for which we have particularly rich genotyping data. A variant passes provisional QC if it meets all the following criteria:

1. It belongs to the 1240k SNP set.
2. Chi2-test(aDNA_SG, aDNA_1240k) < 5
3. Chi2-test(UKBB_UK, GBR_CEU) < chi2_thr
4. Chi2-test(UKBB_EUR, GNOMAD_EUR) < chi2_thr
5. INFO_WEA_1240k > 0.6
6. INFO_WEA_SG > 0.6

We define sample sets in Supplementary Table S1.1 and apply a chi-square test (chi2-test) to compare the allele counts of each variant across three pairs of sample sets. INFO_WEA_1240k and INFO_WEA_SG are IMPUTE2's INFO scores for each variant, calculated for high-quality imputed individuals from western Eurasia with 1240k and shotgun sequences, respectively. We use the Bonferroni-corrected threshold (chi2_thr = 32.03) to filter variants. We select 336 pairs of high-quality imputed sequences from ancient individuals with both shotgun (SG) and 1240k enrichment reagent data, naming them aDNA_SG and aDNA_1240k. These two sets represent different types of sequences for the same individuals, ideally having identical allele frequencies. Allowing for a small error, we choose a chi-square test threshold of 5, resulting in 935,237 SNPs for the provisional variant QC step (Supplementary Figure S1.1).

**Supplementary Table S1.1:** Sample set used in the quality control of variants.

| Sample set name | Number of samples | Description |
|---|---|---|
| aDNA_SG | 336 | Shotgun sequences of ancient individuals with both shotgun and 1240k enrichment reagent data |
| aDNA_1240k | 336 | 1240k sequences of ancient individuals with both shotgun and 1240k enrichment reagent data |
| UKBB_UK | 283 | Sample from the UK Biobank used in this study with the country of birth being the UK. |
| GBR_CEU | 190 | 1000 GP samples from GBR and CEU populations |
| UKBB_EUR | 458,937 | Allele frequency of the European subset of UKBB is presented in the af_EUR column of the variant manifest of pan-ukbb (2022-04-11). |
| GNOMAD_EUR | 4299 | Allele frequency of the European subset of GNOMAD is presented in the gnomad_genomes_an_EUR column of the variant manifest of pan-ukbb (2022-04-11). |
| WEA0 | 79 | Western Eurasian individuals in our dataset with Date=0 years BP but not part of UKBB and 1KG. |
| WEA1 | 959 | Western Eurasian individuals in our dataset with Date<1k years BP but not part of UKBB and 1KG |
| UKBB | 5935 | Sample from the UK Biobank used in this study. |
| 1KG | 503 | 1000 GP samples from EUR populations after dropping up to 2nd-degree relatives. |
| 1KG_ALL | 2504 | All the samples from 1000GP |
| GBR_CEU | 190 | 1000 GP samples from GBR and CEU populations |
| GNOMAD_NFE | 15414 | Allele frequency in samples of non-Finnish European ancestry is represented by the AF_nfe field in the INFO columns of the GNOMAD v2.1.1 publicly available variant VCF files. |
| 1KG_match_UKBB | 478 | 1KG samples that are ancestry matched with samples from UKBB |
| UKBB_match_1kg | 478 | UKBB samples that are ancestry matched with samples from 1KG |
| 1KG_match_WEA1 | 314 | 1KG samples that are ancestry matched with samples from WEA1 |
| WEA1_match_1KG | 314 | WEA1 samples that are ancestry matched with samples from 1KG |
| 1KG_match_WEA0 | 40 | 1KG samples that are ancestry matched with samples from WEA0 |
| WEA0_match_1KG | 40 | WEA0 samples that are ancestry matched with samples from 1KG |
| UKBB_match_WEA1 | 778 | UKBB samples that are ancestry matched with samples from WEA1 |
| WEA1_match_UKBB | 778 | WEA1 samples that are ancestry matched with samples from UKBB |
| UKBB_match_WEA0 | 61 | UKBB samples that are ancestry matched with samples from WEA0 |
| WEA0_match_UKBB | 61 | WEA0 samples that are ancestry matched with samples from UKBB |

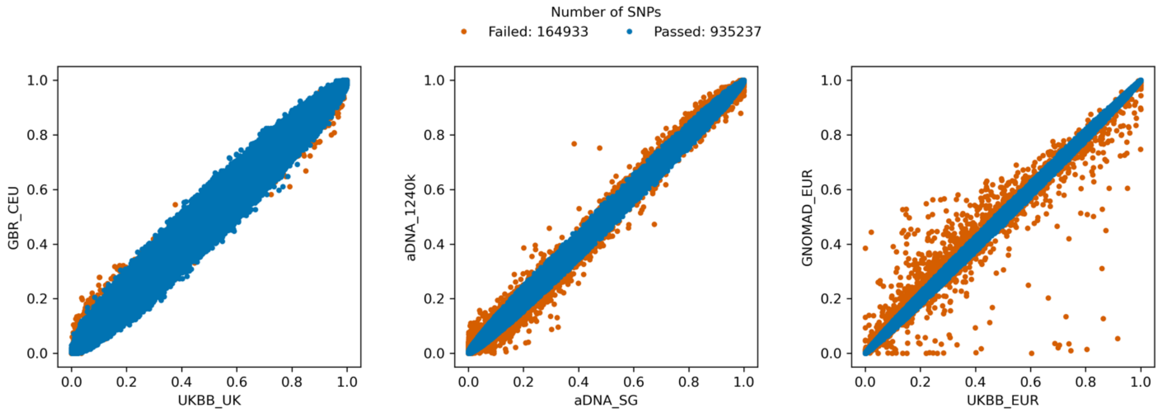

**Supplementary Figure S1.1:** Allele frequency of 1240k SNPs in different sample sets. Each dot represents a SNP; blue dots are SNPs that passed the provisional QC, while orange ones failed. Both the x and y axes show allele frequency in the subset of individuals labeled on the corresponding axis.

### Variant QC: Step 2

We next performed QC not just on the variants in the 1240k SNP set, but on all imputed variants in the 1000 Genomes Project variant set.

We use the provisional QCed SNP set and prune them using plink2 with the --indep-pairwise 1000 1 0.2 option. Then, we use the 5,935 individuals from the UKBB in our imputed dataset restricting to the pruned SNP set, using plink2 to calculate eigenvectors and project all individuals to calculate principal components (PCs). We use 4 subsets of samples to create 5 pairs of matching subsets with similar ancestry using the following procedure: For pop1 and pop2 non-overlapping subsets of individuals, we ran hierarchical clustering on their union set using the sklearn.cluster.AgglomerativeClustering package in Python on the top two PCs calculated above. The number of clusters (n_clusters) is the total number of individuals in these sets divided by 100, with a minimum of 10 clusters. Then, for each cluster, we pair individuals such that one is from pop1 and the other is from pop2 until no such pair is available. The set of all individuals that have been paired and are from pop1 is called pop1_match_pop2, and the other is called pop2_match_pop1. These two subsets consist of samples with similar ancestry based on the top two PCs. The 4 original subsets and their 5 matching pairs are detailed in Supplementary Table S1.1. These matched subsets of individuals are visualized in Supplementary Figure S1.2.

For individual $i$ with both shotgun and 1240k sequences that are in aDNA_SG and aDNA_1240k sets of sequences, we define $\Delta_i = (GT_i(\text{Shotgun}) - GT_i(\text{1240k}))/2$ for each variant such that $GT_i(Y)$ means the imputed genotype in data type Y (shotgun or 1240k) for individual i for the variant of interest. A variant passes final QC if:

1. Minor allele frequency in 1KG_ALL > 0.005
2. $\text{mean}(|\Delta|) < 0.05$
3. The p-value of the null hypothesis $\text{mean}(\Delta) = 0$ is > 1e-5

4. P-chi2-normalized(GBR_CEU, GNOMAD_NFE) > 1e-5
5. P-chi2-normalized(UKBB_match_1KG, 1KG_match_UKBB) > 1e-5
6. P-chi2-normalized(WEA1_match_1KG, 1KG_match_WEA1) > 1e-5
7. P-chi2-normalized(WEA1_match_UKBB, UKBB_match_WEA1 > 1e-5
8. P-chi2-normalized(WEA0_match_1KG, 1KG_match_WEA0) > 1e-5
9. P-chi2-normalized(WEA0_match_UKBB, UKBB_match_WEA0) > 1e-5
10. INFO_WEA_1240k > 0.6
11. INFO_WEA_SG > 0.6

Here, P-chi2-normalized(pop1, pop2) is the p-value of the chi-square statistic, divided by its mean over all variants, from the chi-square test between pop1 and pop2 given the allele counts. Allele frequency consistency between different pairs of sample sets is visualized in Supplementary Figure S1.3.

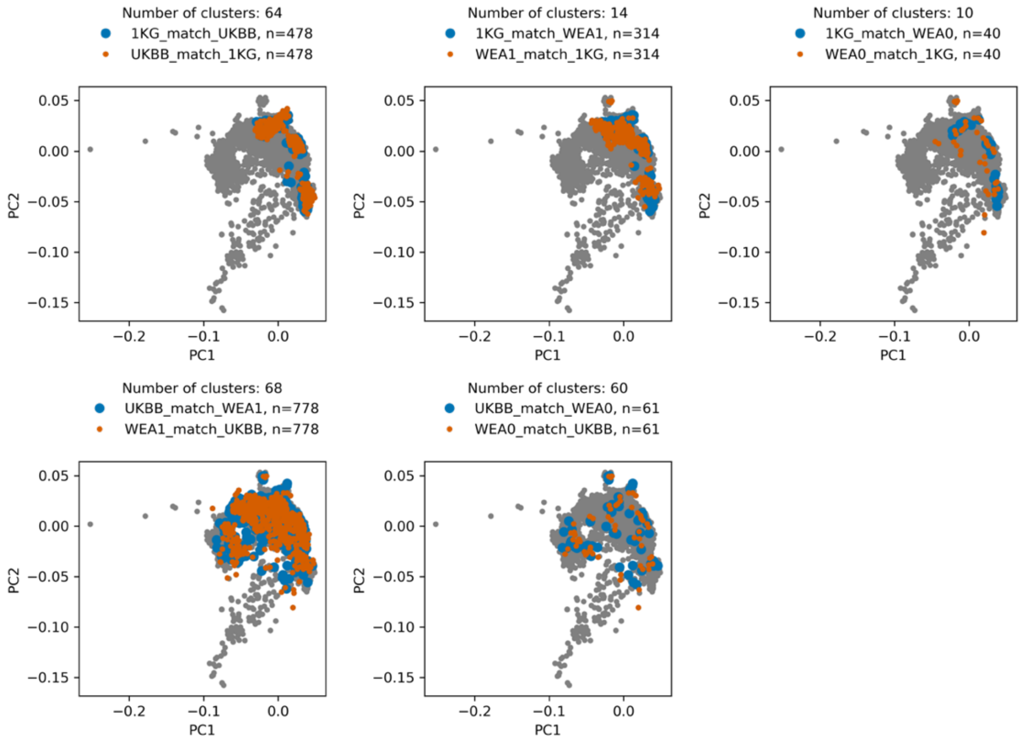

**Supplementary Figure S1.2:** Ancestry-matched samples across different datasets. Each subplot displays ancestry-matched samples. The number of clusters is the n_clusters parameter used in the hierarchical clustering procedure, and n is the final number of ancestry-matched individuals in each sample set.

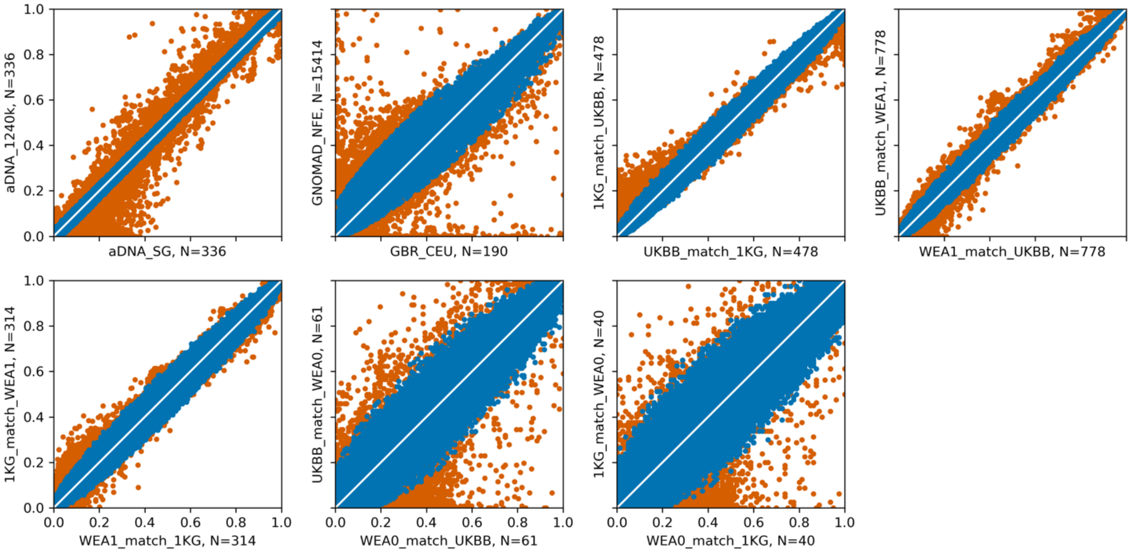

**Supplementary Figure S1.3:** Allele frequency consistency across datasets. The figure displays 1,000,000 randomly selected variants that passed the filters (blue) and 1,000,000 that did not (orange). N represents the number of individuals in each subpopulation. Both the x and y axes show allele frequency in the subset of individuals labeled on the corresponding axis.

### Variant QC: Step 3

To minimize discrepancies between the imputation of ancient DNA and UK Biobank data, we re-imputed the UK Biobank genotyping array from scratch. We utilized Affymetrix confidence files to simulate genotype likelihoods and processed these through the same imputation pipeline employed for ancient DNA. We removed variants that did not pass QC steps 1 and 2 (79.87% of all variants); the remaining 20.13% of variants passing these steps are expected to have less discrepancy across different datasets and sequencing technology (Supplementary Figure S1.3).

To further minimize the impact of batch effects from adding samples from the UK Biobank and 1000 Genomes Project (1000GP) in our selection statistics that also co-analyzed ancient individuals derived from the GLMM method, we ran our analysis both with and without modern individuals. Let Z represent the Z-score of the estimated selection coefficient calculated using all samples in our studies, and $Z_0$ represent the Z-score calculated excluding the modern samples. We employed a weighted least squares (WLS) approach on the two Z-scores and excluded all variants with Pearson residuals greater than 5.45.

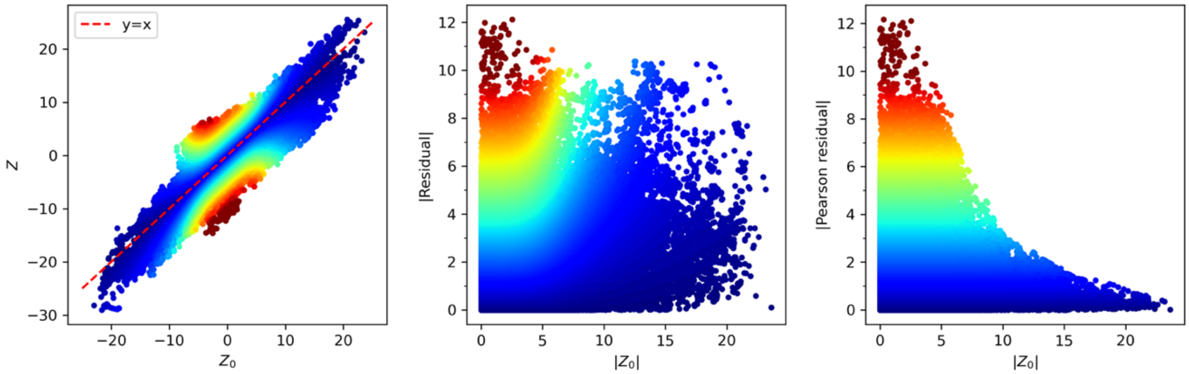

**Supplementary Figure S1.4:** Comparison of Z-scores from GLMM, with (Z) and without ($Z_0$) modern samples. Outliers, which are concerned could be due to batch effects, are identified and removed using Pearson residuals from weighted least squares. Colormap is |Pearson residual|.

We fit a WLS for $Z = a + bZ_0$ with weights = $1/(1 + cZ_0^4)$ and c = 7.41e-4. The weights of WLS should be proportional to the variance of the response variable (Supplementary Figure S1.5). Weights are inferred empirically using the following approach. First, fit an Ordinary Least Square (OLS) for $Z = a + bZ_0$. $\hat{Z}$ is the OLS prediction and $Z - \hat{Z}$ is the residual. Define variable $x = |Z_0|$, $y = \left(Z - \hat{Z}\right)^2 - 1$. Create 100 bins using percentiles of x and fit another OLS such that $\bar{y} = c\bar{x}^4$, where $\bar{x}$ and $\bar{y}$ represent mean values of x and y in each bin. We used the OLS and WLS functions of the statsmodels package in Python for this analysis.

A total of 9,926,484 variants, including 8,212,921 SNPs and 1,713,563 indels, passed all three steps of QC, representing 18.95% of the 52,382,872 imputed variants. The counts of variants passing QC for different variant types (SNP or indel) and their presence in the PAN-UKBB, 1240K, or UK Biobank axiom array SNP sets, are summarized in supplementary table S1.2.

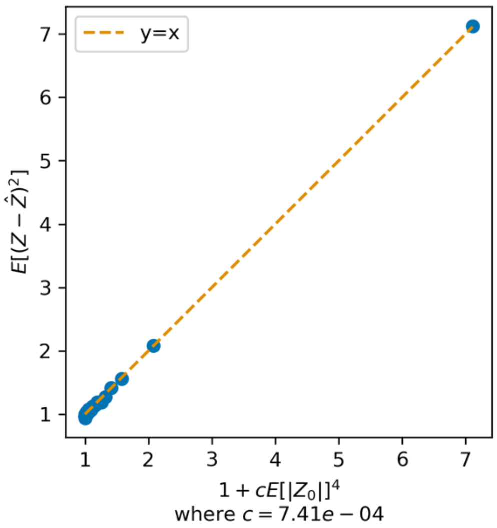

**Supplementary Figure S1.5:** Empirical inference of weights for weighted least square (WLS).

**Supplementary table S1.2:** Summary of variants (SNP and indels) that passed final quality control, categorized by their presence in different variant sets.

| Variant set | Category based on presence in different variant sets (1: present, 0: absent) | | | | | | | | Variant set count |
|---|---|---|---|---|---|---|---|---|---|
| SNP | 1 | 0 | 1 | 0 | 1 | 1 | 1 | 1 | 8,212,921 |
| PAN-UKBB | 1 | 1 | 1 | 0 | 1 | 0 | 1 | 0 | 8,976,393 |
| 1240k | 0 | 0 | 1 | 0 | 0 | 0 | 1 | 1 | 937,454 |
| UK Biobank Axiom | 0 | 0 | 0 | 0 | 1 | 0 | 1 | 0 | 486,264 |
| Category count | 6,751,912 | 929,837 | 808,380 | 783,726 | 358,296 | 165,259 | 127,968 | 1,106 | 9,926,484 |

### Sample Quality Control

For each imputed sample, we define an imputation quality score where GT is the most likely genotype based on the imputed genotype posterior **GP** = ($GP_0$, $GP_1$, $GP_2$) and $GP_0 + GP_1 + GP_2 = 1$. We only kept samples with high imputation quality score IQS>0.9. We used the KING software to capture duplicates and related samples up to the second degree. We used a greedy algorithm to prioritize samples by their IQS and drop all relatives up to the second degree until there are no two samples that are second-degree related or more.

We next carried out a filtering step to remove individuals that we found were reducing the significance of real signals of selection, either because their assigned dates were incorrect, or because they had primary ancestry that was badly captured in the time transect. We ran a generalized linear model (GLM) with the date of the sample being the explanatory variable and allele counts as the response. We use the p-values to prioritize variants based on their association with the date. We restrict to high-quality variants in the 1240k SNP set using plink2 with the option --indep-pairwise 100 1 0.2 and a fake frequency file such that p-values calculated from the GLM are mapped to a minor allele frequency (MAF) value between 0 and 0.5, where the smallest p-values are mapped to MAF=0.5 and the largest p-values are mapped to MAF=0. This way, plink2 is prioritizing by larger values of MAF, which is equivalent to smaller (more significant) p-values. This pruned SNP set is used to calculate the top 100 principal components (PCs) of individuals in the provisional set. These 100 PCs are used as explanatory variables to fit a linear regression with the date of the samples as the response variable. If the estimated date and reported date are significantly different, we exclude those samples from our analysis using the following criteria:

1. Date <15,000 years BP
2. Date_SE <500 years or the absolute value of the regression residual <500 years
3. P-value of Pearson residual > 1e-5

Here, Date is the reported date of the sample and Date_SE is the reported standard error. We use outliers in the top two PCs to remove outliers with some African admixture.

### Supplementary Information section 2
### Statistical criterion for genome-wide significance

#### Establishing the Optimal Significance Threshold

Numerous confounding factors, such as population structure, genetic drift, batch effects, the quality of genotype imputation, and the processes of sequence generation and data preprocessing, have the potential to lead to genomic inflation. We implemented extensive data cleaning and quality control measures, along with the Generalized Linear Mixed Model (GLMM) approach, to mitigate the impact of these confounding factors on our findings. However, complete elimination of these issues is not possible, and thus further correction for their effects during the final analysis phase of the summary results is essential. We tried multiple approaches to control family wise error rate (FWER). However, we found that these approaches are either inefficient or not robust. We used an alternative approach of controlling for false discovery rate (FDR) by leveraging high quality GWAS studies.

#### Controlling for family wise error rate (FWER)

We explored various approaches to identify a control factor (CF) to adjust the nominal $\chi^2$ of the selection signal that is both reliable and maximizes the utility of the data. We tried three different CF: genomic inflation factor ($\lambda_{GC}$), simulation based, and finally LD score regression intercept.

**Genomic inflation factor ($\lambda_{GC}$).** To adjust for residual confounding of our selection statistics, we tried adjusting with genomic inflation ($\lambda_{GC}$), defined as the median of the nominal $\chi^2$ of the selection coefficient divided by the median of a chi-square distribution with 1 degree of freedom (0.455). This empirical correction factor is 5.26 for our dataset. We tried using the $\lambda_{GC}$ as the control factor for the nominal $\chi^2$ . The nominal genome-wide significance threshold, corresponding to an adjusted P-value threshold of 5e-8 with CF=5.26, is p=7.2e-36, which yields 48 independent loci, excluding the HLA region. By using orthogonal information (GWAS data) to estimate the fraction of the genome affected by directional selection, however, we infer that 26% of analyzed sites are in at least weak linkage disequilibrium (LD) ($r^2 > 0.05$) with high-confidence signals of selection (<5% False Discovery Rate - FDR) (Extended Data Figure 2b). This is a much larger fraction of the genome shaped by directional selection signals than can be explained by 48 loci, and thus, relying on $\lambda_{GC}$ is too conservative.

**Simulation.** We turned to simulation, which suggests that the control factor for the nominal $\chi^2$ is 1.04. The nominal genome-wide significance threshold, corresponding to an adjusted P-value threshold of 5e-8 with CF=1.04, is p=2.7e-8, which yields 8,210 independent loci. The FDR for this threshold is estimated to be 44%, which is far more than the <1% that would be expected if the threshold was well-calibrated. Therefore, our simulation is not realistic enough and does not adequately adjust for all the artifacts contributed to false-positives in real data.

**LD score and pseudo-intercept.** "LD score regression" (LDSC) analyzes the relationship between the LD among variants and their associated test statistics to disentangle the effects of polygenicity from confounding. If it is working properly is expected to show a linear relationship between "LD score bin" and tests statistics whose y-intercept that can be used to estimate inflation and is often in fact used in this way in GWAS studies[1]. We explored using this approach to correct for inflation in our selection scan.

A potential challenge in using LDSC to correct for inflation in a selection scan is that our dataset is plausibly more affected by data artifacts than GWAS studies in living people where data sources are relatively homogeneous. The data we are analyzing here comes from much more diverse sources: imputed ancient DNA sequences (shotgun sequences and 1240k enrichment reagent), sequences of people of European ancestry from the 1000 Genomes Project, and imputed SNP array data from the UK Biobank (genotyped using the UK Biobank Axiom Array). While we employed extensive quality control and data cleaning (Supplementary Information section 1) with the goal of minimizing the impact of artifacts, data cleaning is never perfect, and some artifacts will always be present. The accuracy of imputed genotypes is also expected to be lower for variants with low levels of LD and low minor allele frequency in the reference panel[2], and for ancient individuals who are less related than modern individuals to the populations in the reference panel used for imputation[3]. This raises the possibility of higher inflation for variants with lower LD score, and indeed, the LD-score vs. nominal $\chi^2$ of selection reveals exactly what would be expected from such issues: a non-linear relationship between LD score bin and test statistics that is not expected for well-behaved LDSC (Supplementary Figure S2.1). Because of these issues, we cannot simply use the LDSC framework to determine an appropriate inflation factor correction.

We explored computing a pseudo-intercept using S-LDSC with baseline v2.2, where SNPs with LD-scores above a minimum threshold, called min(LD-score), are considered in the calculation of the intercept. The pseudo-intercept as a function of min(LD-score) decreases monotonically until min(LD-score) = 89, which gives a pseudo-intercept of 2.63, and then after that point, it fluctuates. The nominal genome-wide significance threshold, corresponding to an adjusted P-value threshold of 5e-8 with CF = 2.63, is 9.5e-19, which yields 319 independent loci, excluding the HLA region. The FDR for this threshold is estimated to be 1.2%. This CF from the pseudo-intercept is the most reasonable compared to previous approaches, assessed based on the independent FDR criterion, and in fact it is very similar to the threshold we used in practice and which we obtained by calibration from orthogonal data (enrichment in signals of association to phenotypes from GWAS). However, because of the non-linearities, we refrain from using this correction to control formally for FWER.

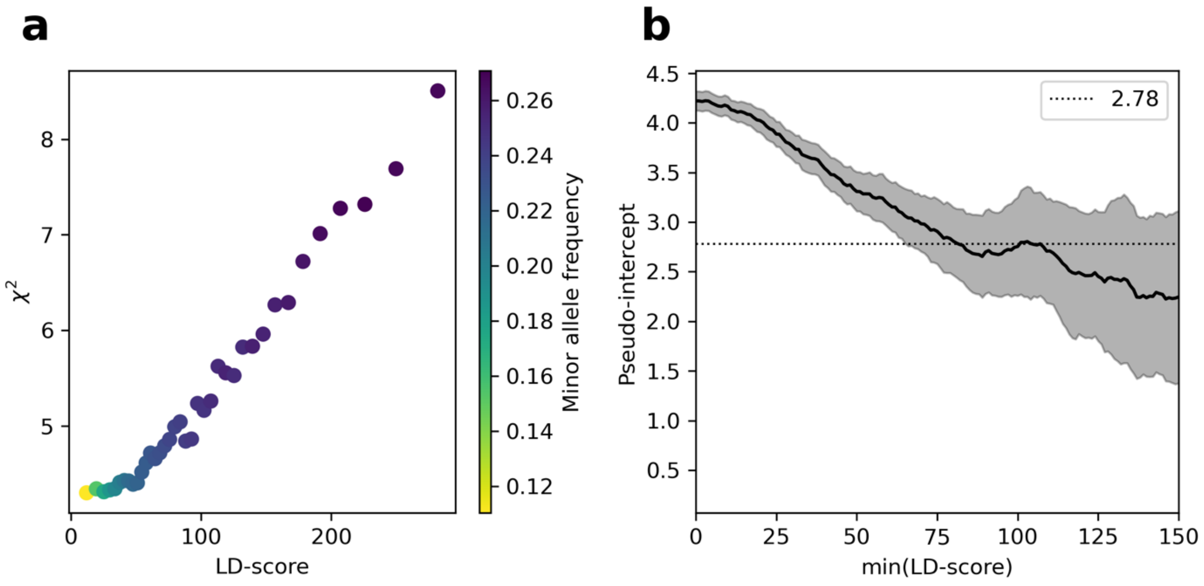

**Supplementary Figure S2.1:** (**a**) LD score plot for nominal $\chi^2$ statistics, with each point representing an LD score quintile. Values are averaged within each bin for visualization purposes only. (**b**) The pseudo-intercept is the intercept of the LD score regression after dropping all SNPs with LD scores below the minimum threshold.

#### Controlling for false discovery rate (FDR) by leveraging GWAS signals

In this study, we abandoned controlling for the family-wise error rate (FWER), which is the probability of obtain one or more false positives in a multiple testing scenario, and instead decided to take advantage of information-rich, high-quality GWAS data to estimate the false discovery rate (FDR) and posterior probability, and thereby to calibrate our summary statistics.

In Supplementary Figure S2.2, the x-axis is the nominal p-value of the selection coefficient, and the left y-axis is the enrichment in pan-UKBB GWAS studies. To estimate the enrichment values, ultimately determining FDR and the posterior probability of being a true selection signal, we followed the procedure outlined below.

First, for all SNPs that pass the quality control, we apply a pruning procedure using PLINK (version 1.9) with an $r^2$ threshold of 0.99 and a window size of 1 Mbp. This results in 3,085,793 SNPs, of which 14.5% are genome-wide significant in at least one of 454 GWAS from the Pan-UKBB that pass QC.

To adjust for minor allele frequency variation, we initially applied a logistic regression model to this set of SNPs:

$$log\left(\frac{p_i}{1-p_i}\right) = a + bf_i$$

Here, $p_i$ denotes the expected probability that SNP $i$ is a GWAS hit, given the minor allele frequency $f_i$, and $y_i$ is a binary outcome variable indicating whether SNP $i$ is identified as a GWAS hit ($y_i = 1$) or not ($y_i = 0$).

For each threshold for significance $t$, we define $R(t)$ as:

$$R(t) = \frac{1}{N(t)} \sum_{i \in S_t} \frac{y_i}{p_i}$$

Where $S_t=\{i: P_i \leq 10^{-t}\}$ and $N(t)$ is number of SNPs in set $S_t$. Subsequently, we define $\alpha_0=R(0)$, and $E(t)=R(t)/R(0)$. We rewrite E(t) in the following form:

$$E(t) = \frac{\int_{x=t}^{\infty}\left[n(x)p(\text{real}|x)\alpha_1 + \left(1-p(\text{real}|x)\right)n(x)\alpha_0\right]dx}{(\alpha_1-\alpha_0)\int_{x=t}^{\infty} n(x)dx}$$

where $n(t) = \frac{-\partial N(t)}{\partial t}$ and $p(\text{real}|t)$ is the posterior probability of being a true selection signal.

Thus,

$$p(\text{real}|t) = \frac{\partial\left(E(t)N(t)\right)}{\partial\left(N(t)\right)}$$

The false discovery rate *FDR(t)* is defined as:

$$FDR(t) = 1 - \frac{E(t) - 1}{\lim_{t \to \infty} E(t) - 1}$$

To estimate *E(t)* and *N(t)* smoothly, we utilize the polyfit function from the numpy package in Python. Our estimations for values approaching 0 or 1 tend to be less reliable. Consequently, we refrain from reporting values below 1% or above 99%.

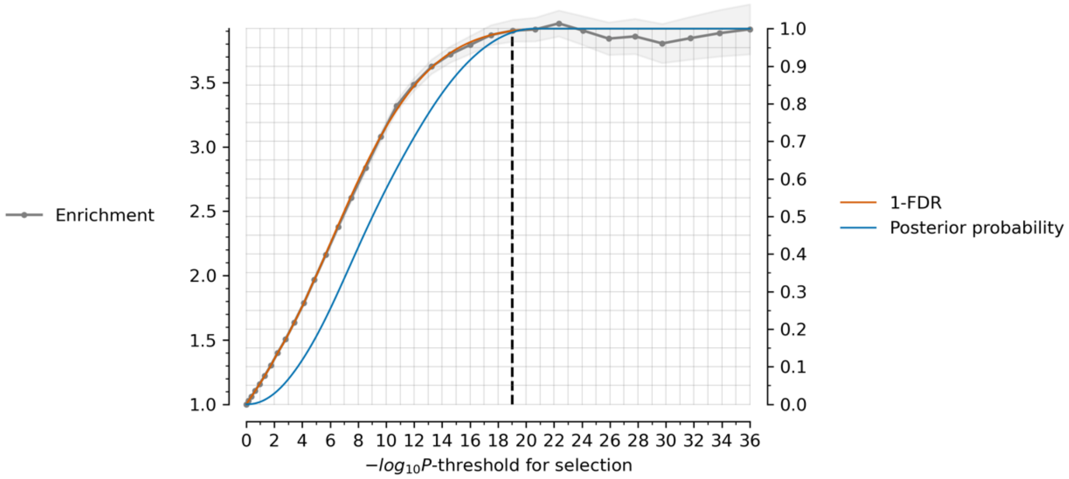

**Supplementary Figure S2.2:** Estimated FDR and posterior probability of being a true selection signal as a function of the nominal p-value threshold. The black dashed line shows the P-threshold (9.55e-20) with a posterior probability of 99%.

This analysis suggests that at a nominal p-value threshold of 9.55e-20, the posterior probability is ~99%. The appropriate threshold for controlling FWER should be somewhere between 9.5e-19 and 7.2e-36 based on the discussion above. However, the 7.2e-36 threshold seems extremely conservative, while the enrichment in GWAS studies gives us a good handle on FDR, not FWER, and makes the most out of the data, and hence we use the FDR approach.

We implement the FDR approach in practice using a newly defined statistic $X = Z/\sqrt{CF}$, where Z is the nominal test staticic for the selection signal without full correction for inflation, and CF is an empirically calibrated correction factor. We set CF=2.78 ($\sqrt{CF} = 1.67$) based on this being the point at which we estimate that the posterior probability is 99%. Encouragingly, this falls within the 95% confidence interval (2.47, 2.91) estimated using the pseudo-intercept of the LD score regression, providing a supporting line of evidence that this is a reasonable approach.

With our parameterization of the X-statistic, the threshold |X|=5.45 which if it was a normally distributed variably would imply a classically genome-wide significance threshold of $5 \times 10^{-8}$, is the threshold of genome-wide significance for our study as well. Thus, p-values obtained from interpreting our X-score as a normally distributed variable provide reasonable guidelines for whether particular SNPs are genome-wide significant.

### Supplementary Information section 3
### HAF score analysis proves directional selection

#### HAF score dynamic for positive and negative selection

The haplotype allele frequency (HAF) score for a given haplotype is calculated by summing the derived allele counts of the polymorphic sites on that haplotype[4,5]. It distinguishes carrier haplotypes from non-carriers of the favored allele in an ongoing selective sweep, without prior knowledge of the favored allele. The HAF-score is defined for a haploid population, requiring phased haplotype information and ancestral and derived allelic states to distinguish carriers from non-carriers of the favored mutation in an ongoing selective sweep. For a diploid population, however, calculating the mean HAF-score does not require phased information, and only the derived allele frequency (DAF) is needed. The mean HAF-score is given by:

$$\overline{HAF} = n \sum_{i=1}^{m} DAF_i^2$$

where $n$ is the number of haplotypes (twice the number of diploid individuals), $m$ is the number of polymorphic sites in the sample, and $DAF_i$ is the derived allele frequency for the $i$-th polymorphic site.

In a neutrally evolving population with a constant population size $N$, the expected HAF score under the coalescent model[6] is:

$$E[HAF] = \frac{\theta(n-1)}{2}$$

where $\theta = 2N\mu L$ represents the scaled mutation rate, $\mu$ is the mutation rate per base pair per generation, $L$ is the haplotype length, and $n$ is the number of sampled haplotypes.

Ronen et al. (2015) show that for strong selection ($Ns \gg 1$) without recombination, the expected HAF scores for carrier and non-carrier haplotypes of a favored allele with frequency $f$ during a hard selective sweep are given by:

$$E[HAF^{car}] \approx \theta n \left(\frac{f+1}{2} - \frac{1}{(1-f)n+1}\right)$$

$$E[HAF^{non}] \approx \theta n \left(\frac{1}{2} + \frac{1}{2n} - \frac{1}{(1-f)n+1}\right)$$

Thus, the expected HAF score for $n$ haplotypes undergoing a hard selective sweep is:

$$E[HAF] \approx \theta n\left(\frac{f^2+1}{2}+\frac{1-f}{2n}-\frac{1}{(1-f)n+1}\right)$$

When $n(1-f)f > (2-f)$, E[HAF] is greater than $\theta(n-1)/2$, indicating that the expected HAF score is larger than the neutral case if the sweep is not near fixation (Supplementary Figure S3.1). Thus, a positive deviation from expectation provides evidence of a partial sweep.

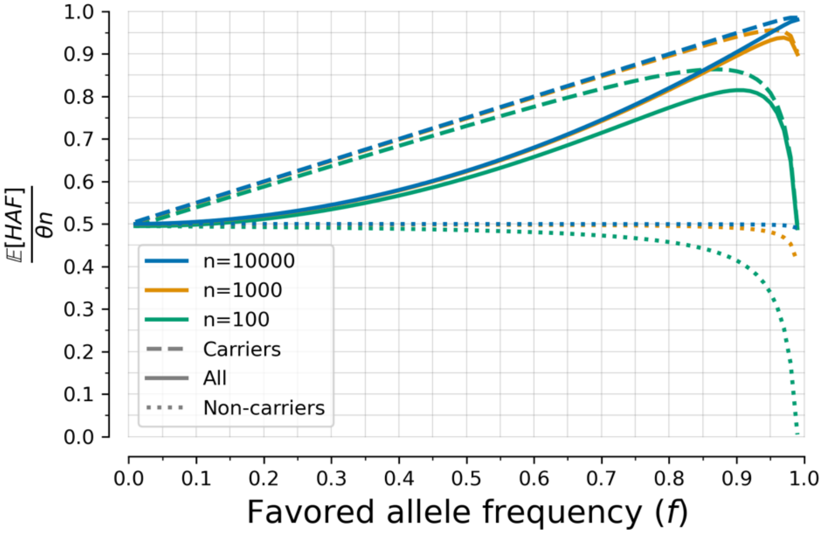

**Supplementary Figure S3.1:** Haplotype allele frequency (HAF) score as a function of favored allele frequency during an ongoing hard selective sweep without recombination.

For negative selection, the effective population size (*N*) is reduced due to linked background selection. This leads to a decrease in $\theta$ and the expected HAF score, making it lower than in the neutral case[7]. Thus, in principle, observation of a rise in the HAF score can provide unambiguous evidence of positive directional selection associated with our X-statistic.

A challenge in testing for evidence of direction selection based on a rise in the HAF score is that alleles in the genome that are subject to positive selection are also expected to be regions rich in functional effects and, thus, are expected to be more subject to purifying selection. When we compute how HAF score changes as a function of our test statistic for natural selection, we observe a nominal decrease in HAF score—not the increase expected for positive selection—which could be due to this phenomenon (Extended Data Figure 3c). To distinguish between scenarios in which the correlation of our X-statistic to functionally more important regions is due at least in part to the signal we are interested in (that is, directional selection), and cannot be trivially explained as an artifact of purifying selection at alleles in functionally more important regions leading to false-positives, we controlled for the effects of linked negative selection (B-statistic) genome-wide. The resulting residual HAF-score increases as a function of X-statistic, plateauing around |X|=5.45, the same plateau as we observe for the enrichment of signals in independent GWAS data (Figure 1b, Extended Data Figure 3c)

### Supplementary Information Section 4

### Allele frequency trajectory and selection coefficient over time for 347 independent loci with >99% probability of selection

In this section, we visualize allele frequency trajectories (Supplementary Figure S4.1-S4.29) and selection coefficient over time (Supplementary Figure S4.30-S4.58) for 347 independent loci (279 outside the HLA region and an additional 68 in HLA) with |X|>5.45 corresponding to a $\pi$>99% probability of selection. To produce this list, we identified the strongest signal in the genome and considered all SNPs in LD with it in modern Europeans from the 1000 Genomes Project ($r^2$>0.05) to potentially reflect the same signal. We then found the second-strongest signal excluding these positions, and so on, until no more SNPs pass this threshold (Extended Data Figure 2b). The SNPs are listed in genomic order.

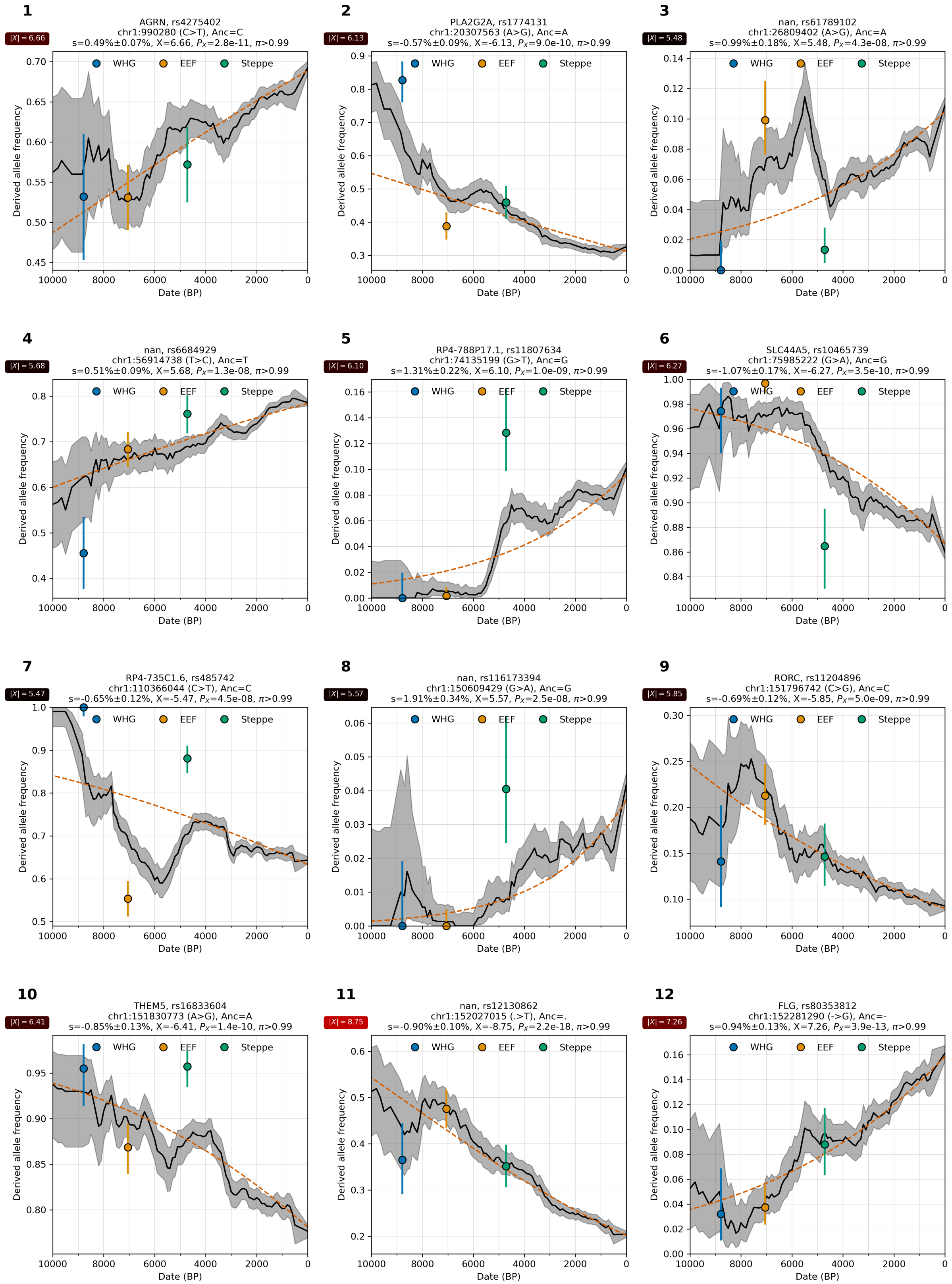

**Supplementary Figure S4.1:** Allele frequency over time. From chr 1.

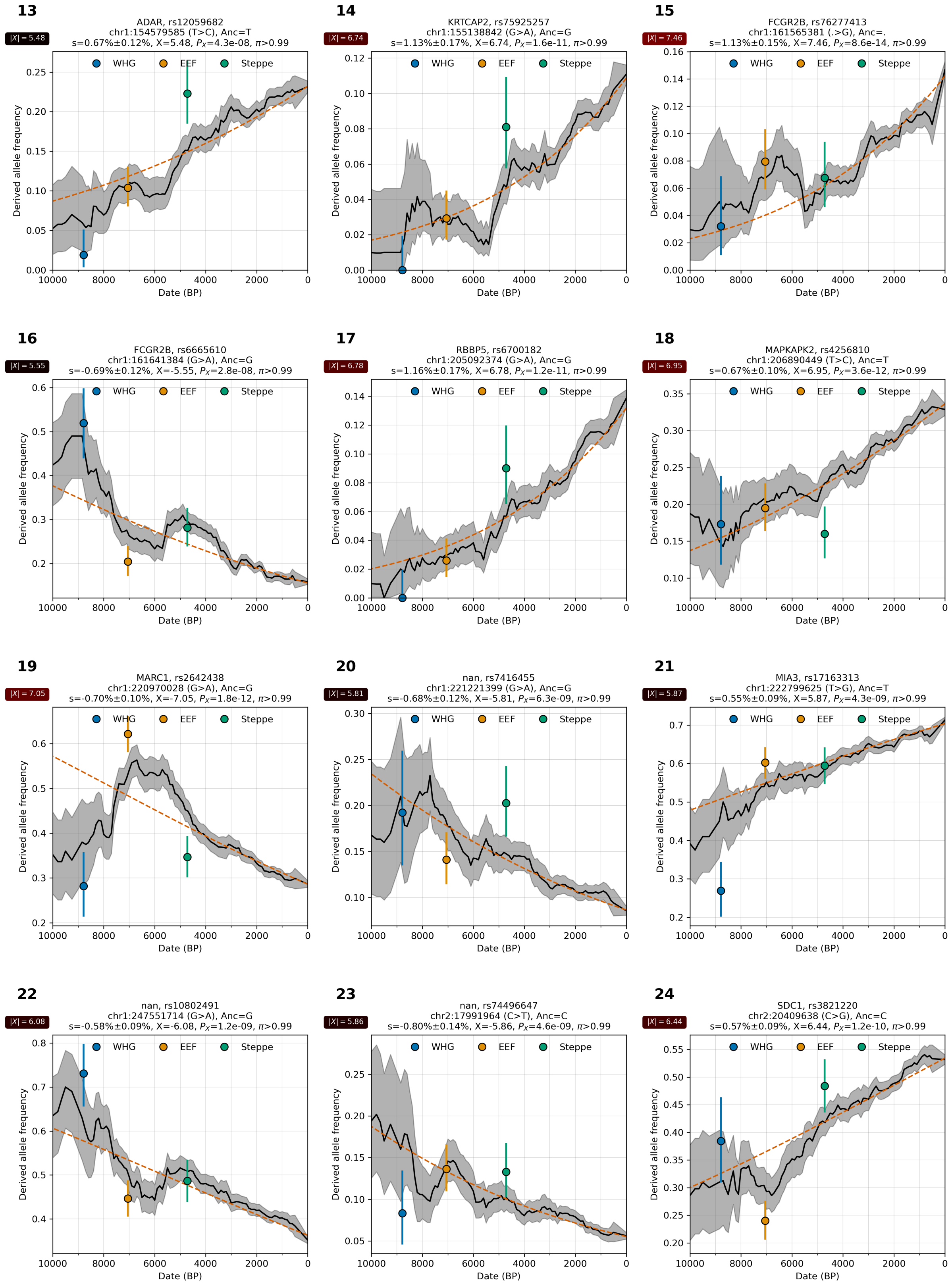

**Supplementary Figure S4.2:** Allele frequency over time. From chr 1 and 2.

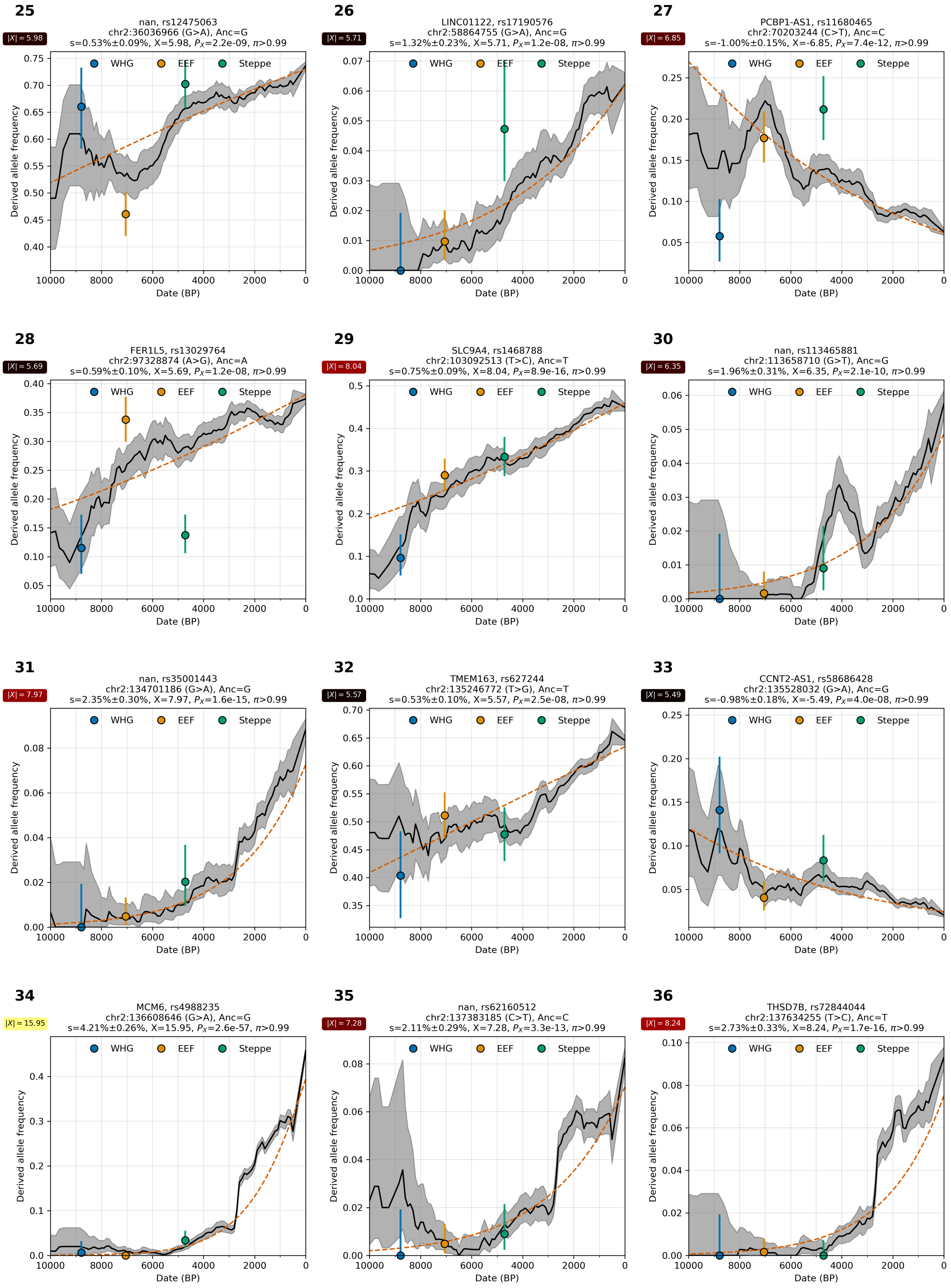

**Supplementary Figure S4.3:** Allele frequency over time. From chr 2.

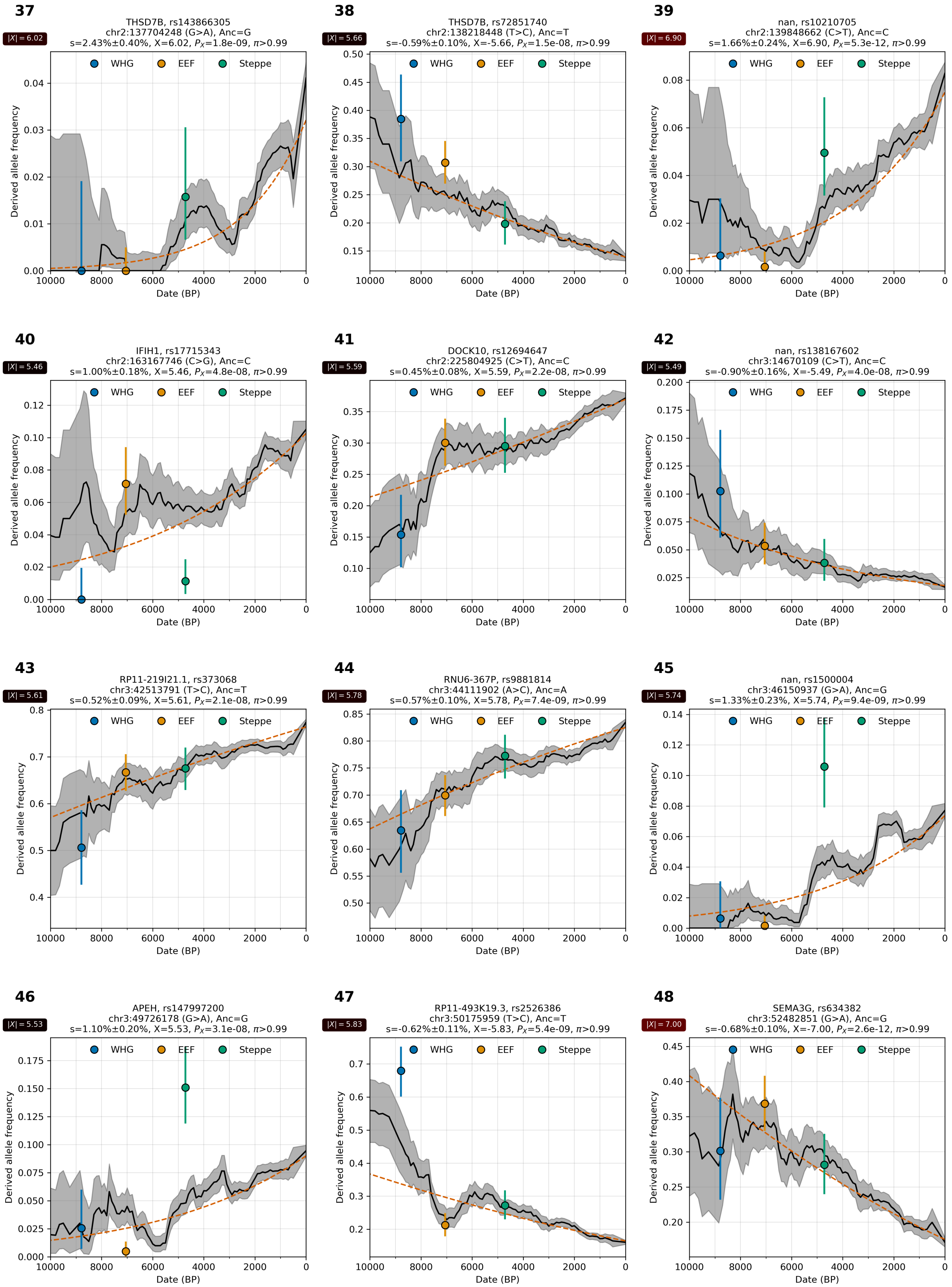

**Supplementary Figure S4.4:** Allele frequency over time. From chr 2 and 3.

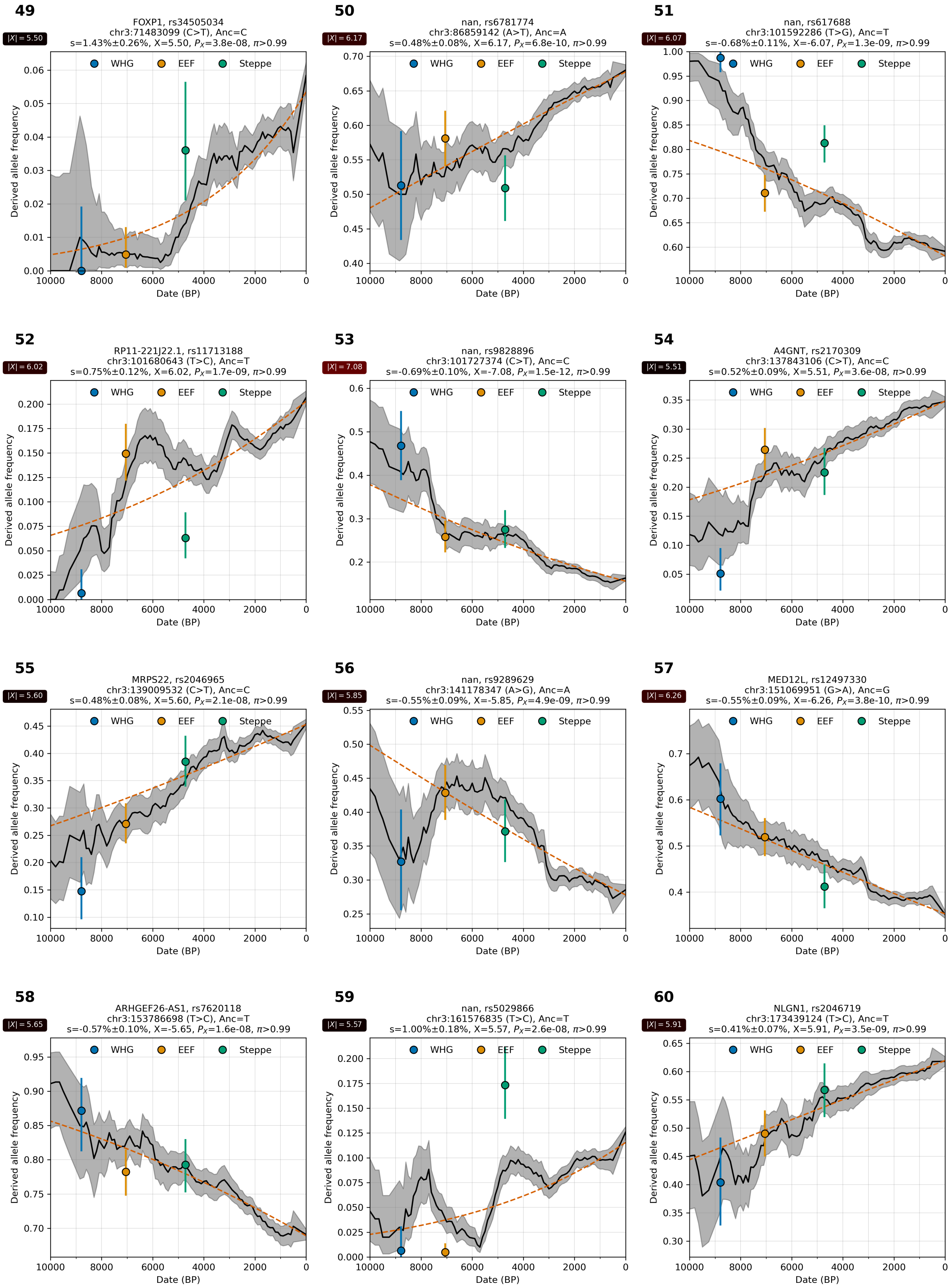

**Supplementary Figure S4.5:** Allele frequency over time. From chr 3.

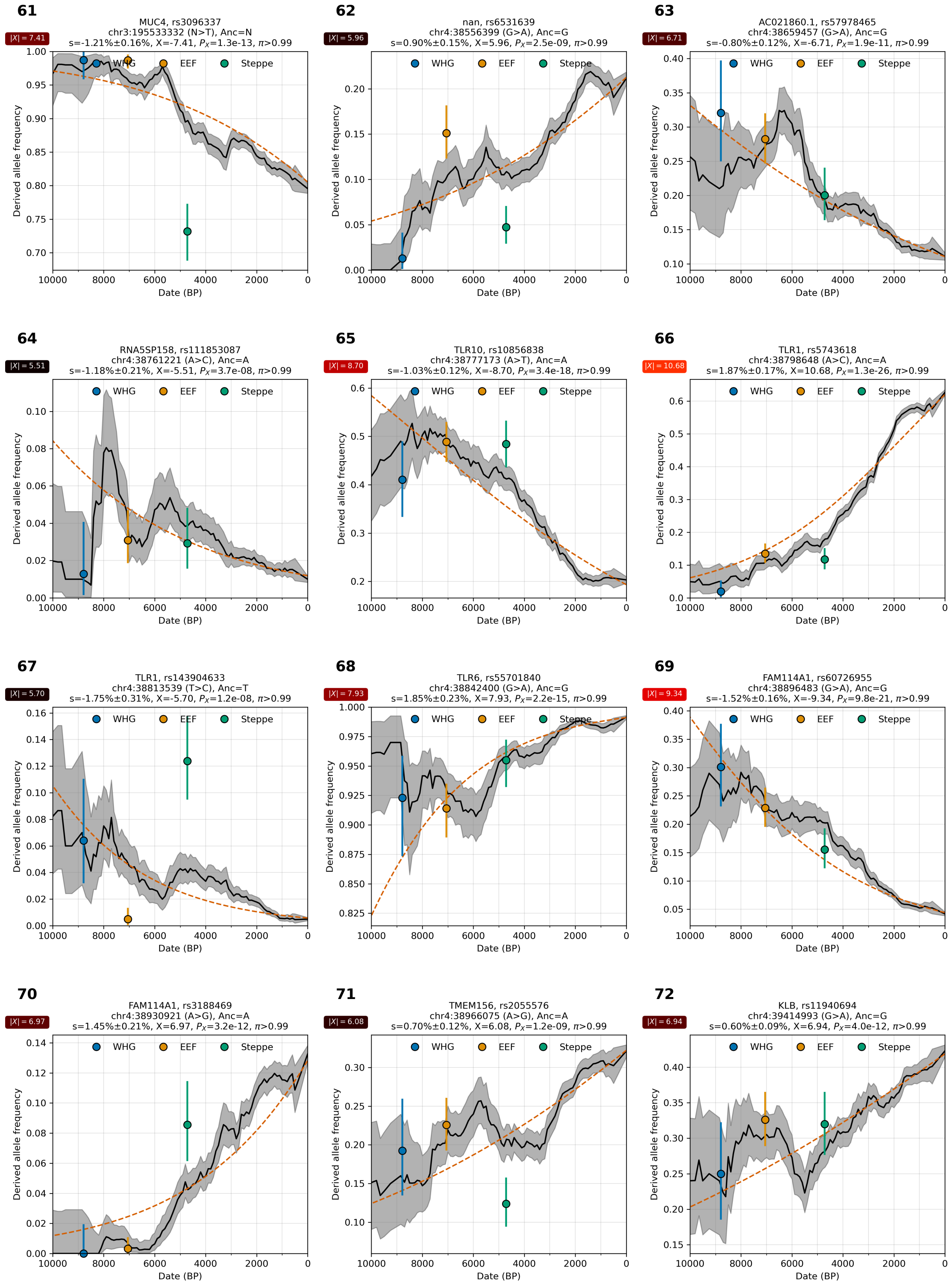

**Supplementary Figure S4.6:** Allele frequency over time. From chr 3 and 4.

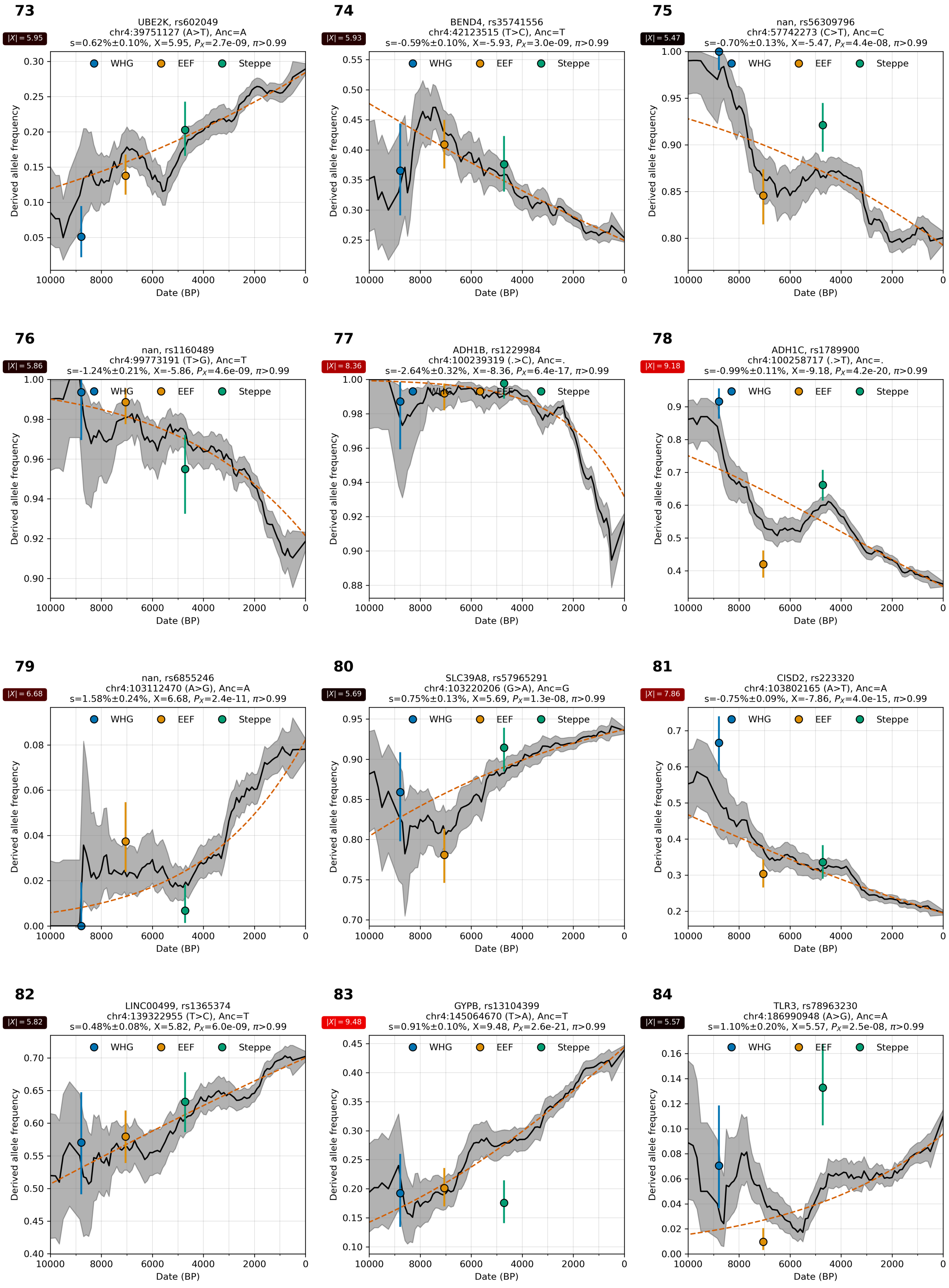

**Supplementary Figure S4.7:** Allele frequency over time. From chr 4.

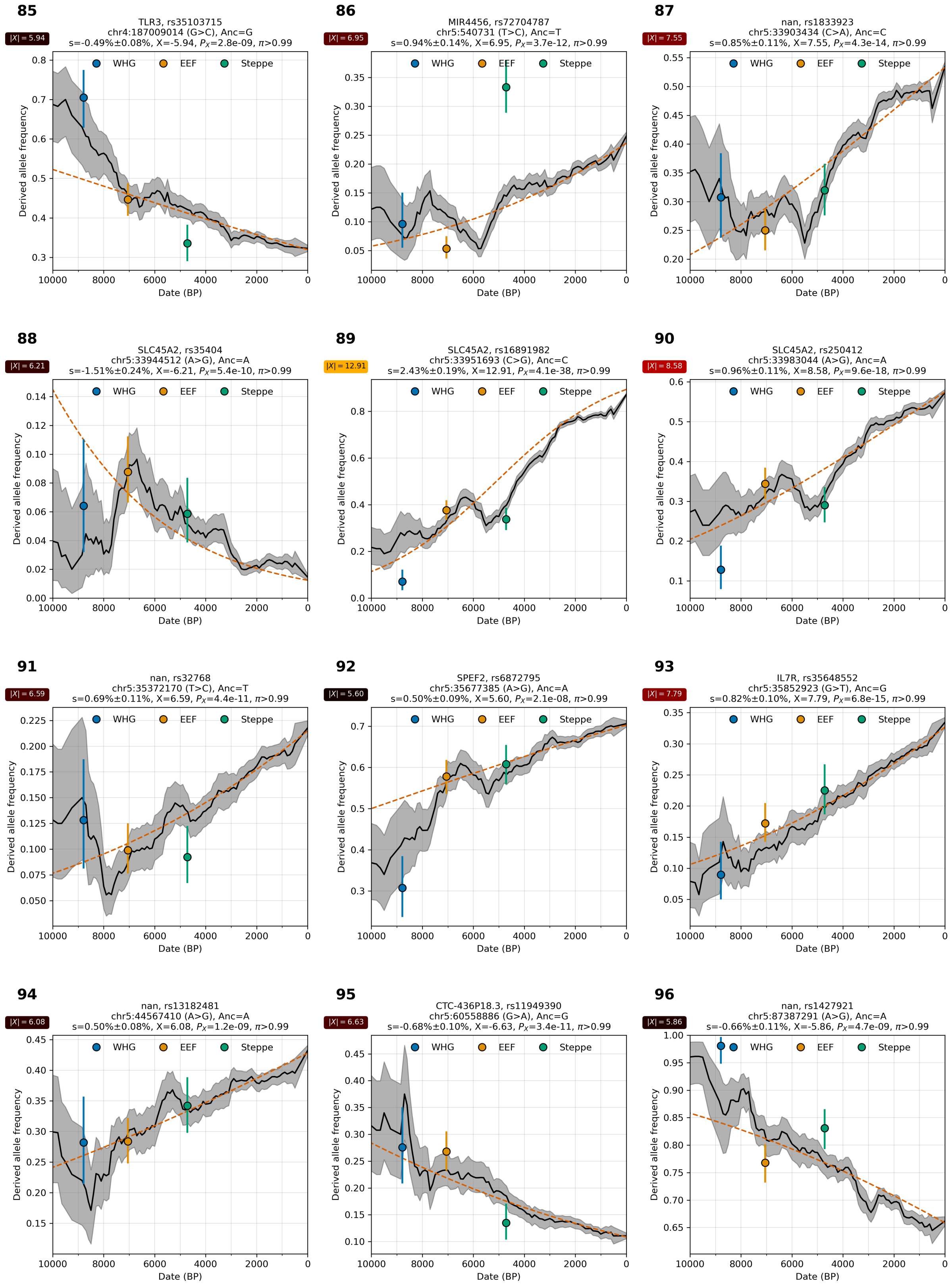

**Supplementary Figure S4.8:** Allele frequency over time. From chr 4 and 5.

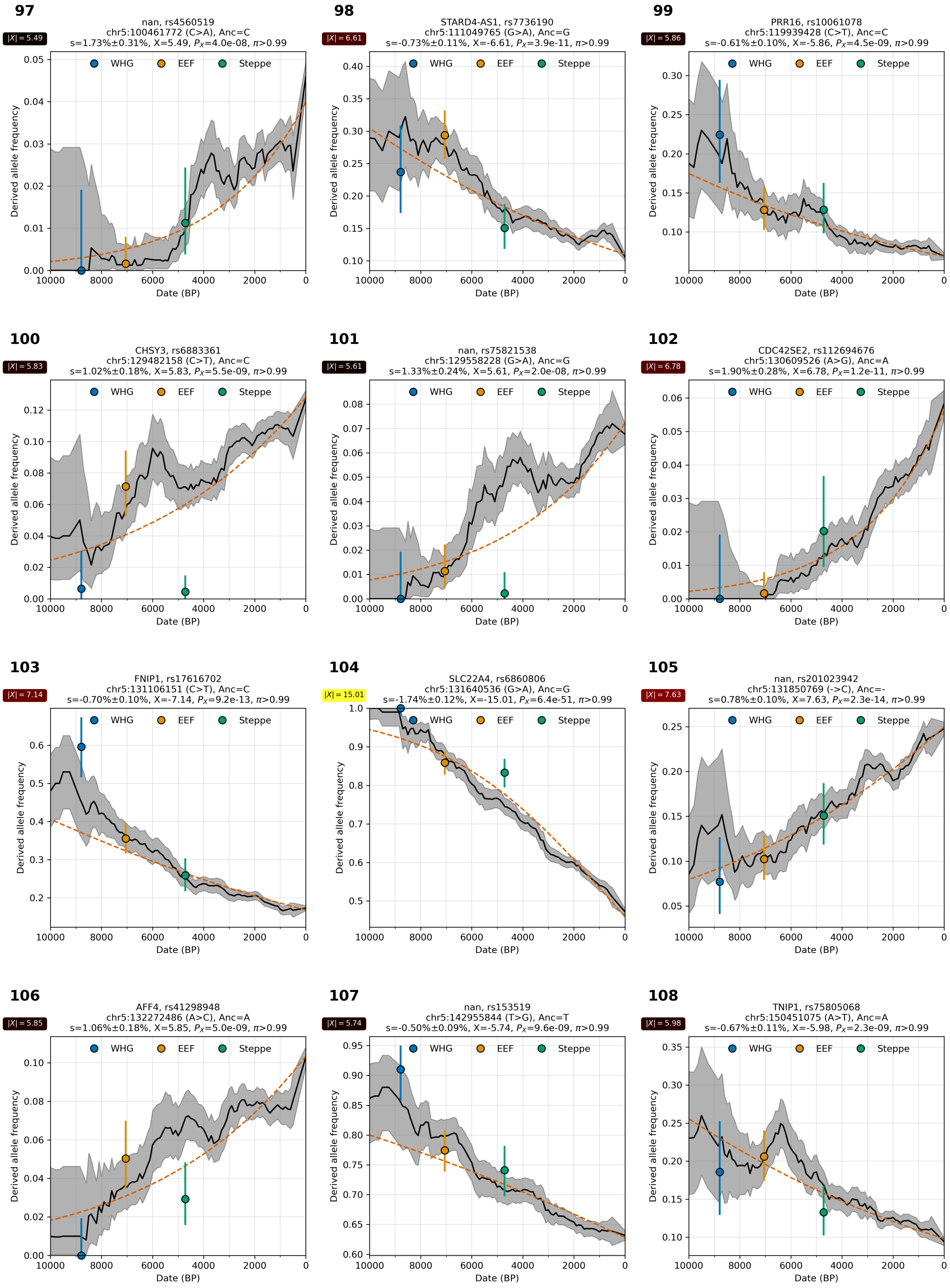

**Supplementary Figure S4.9:** Allele frequency over time. From chr 5.

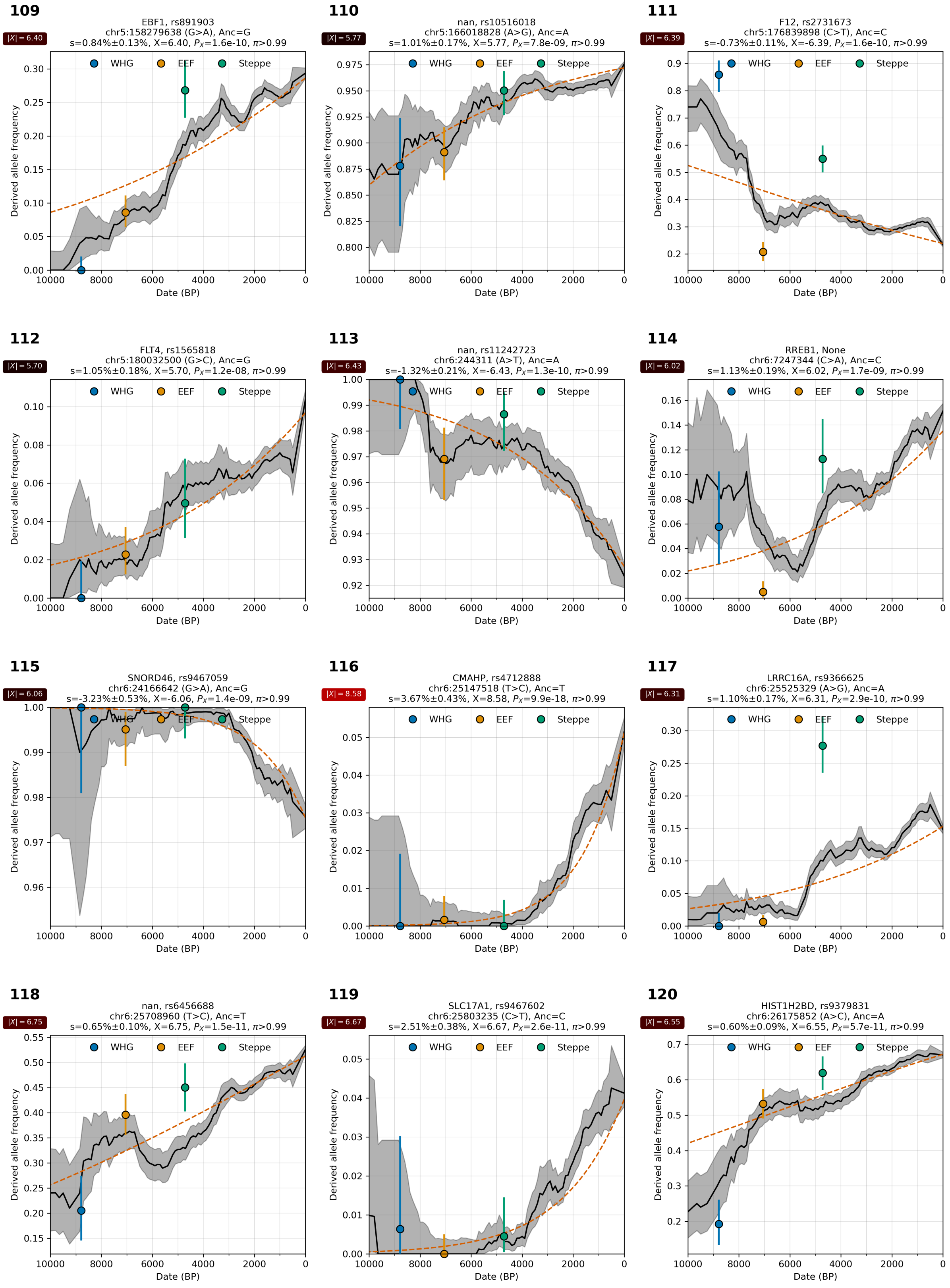

**Supplementary Figure S4.10:** Allele frequency over time. From chr 5 and 6.

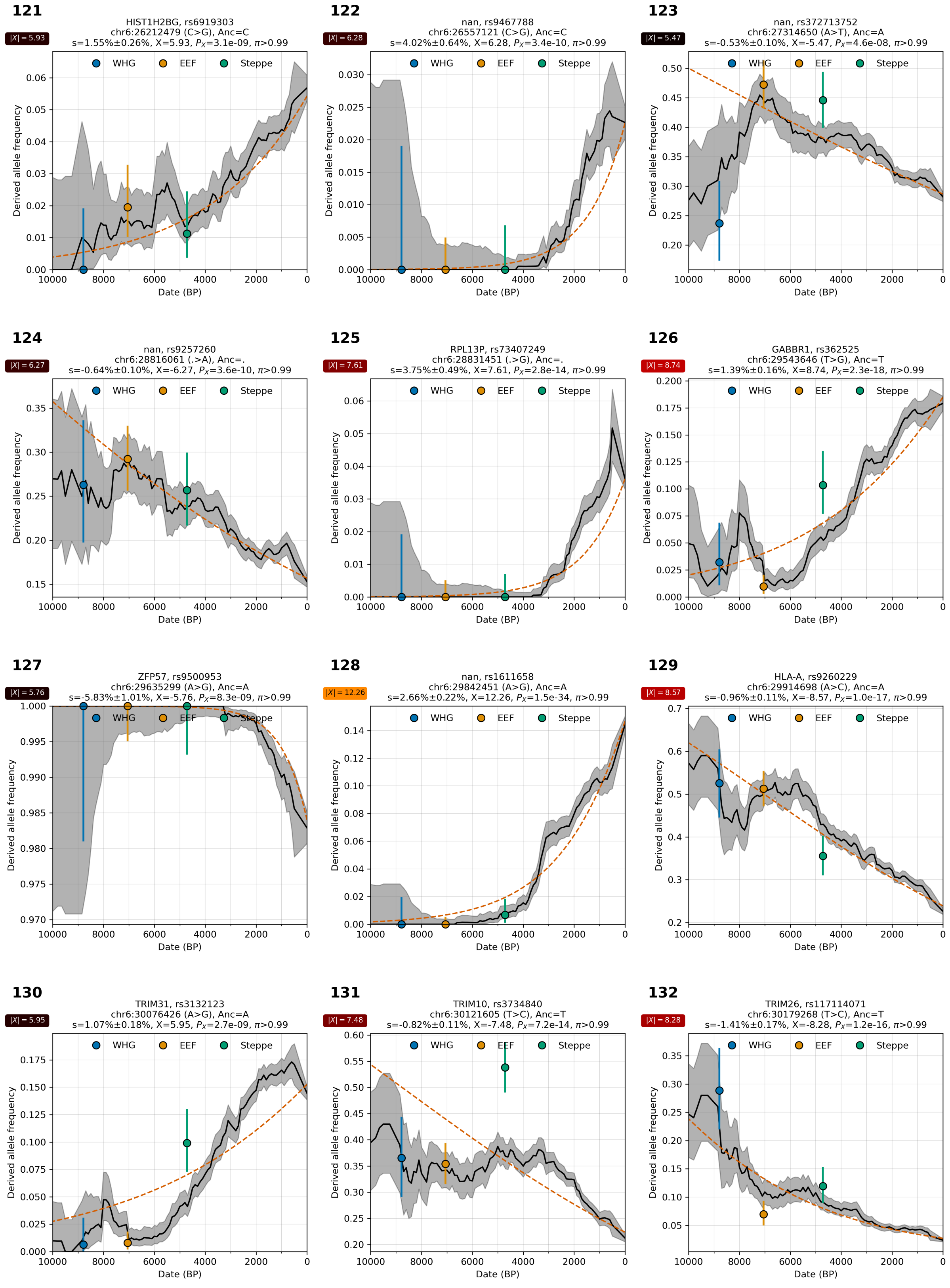

**Supplementary Figure S4.11:** Allele frequency over time. From chr 6.

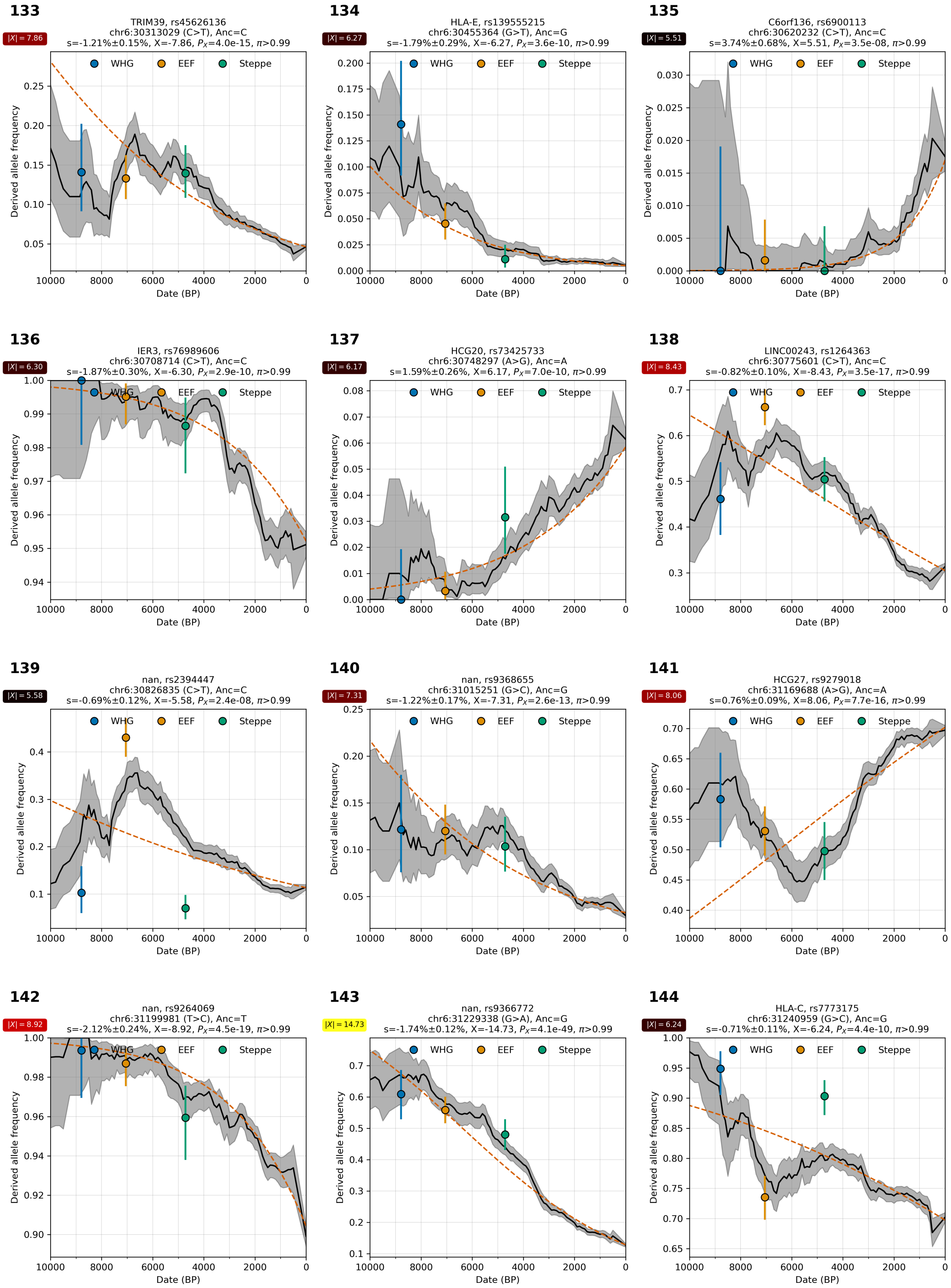

**Supplementary Figure S4.12:** Allele frequency over time. From chr 6.

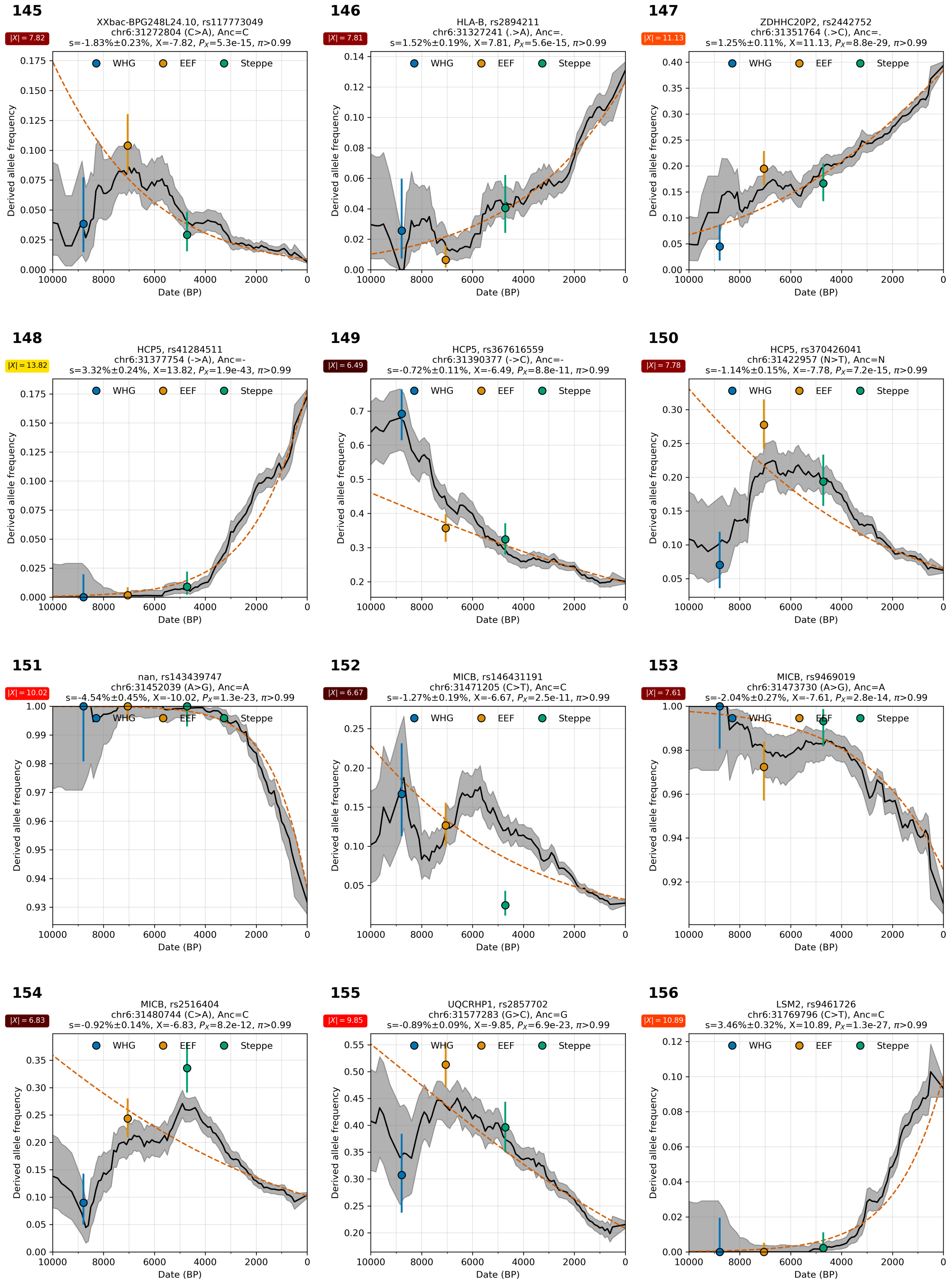

**Supplementary Figure S4.13:** Allele frequency over time. From chr 6.

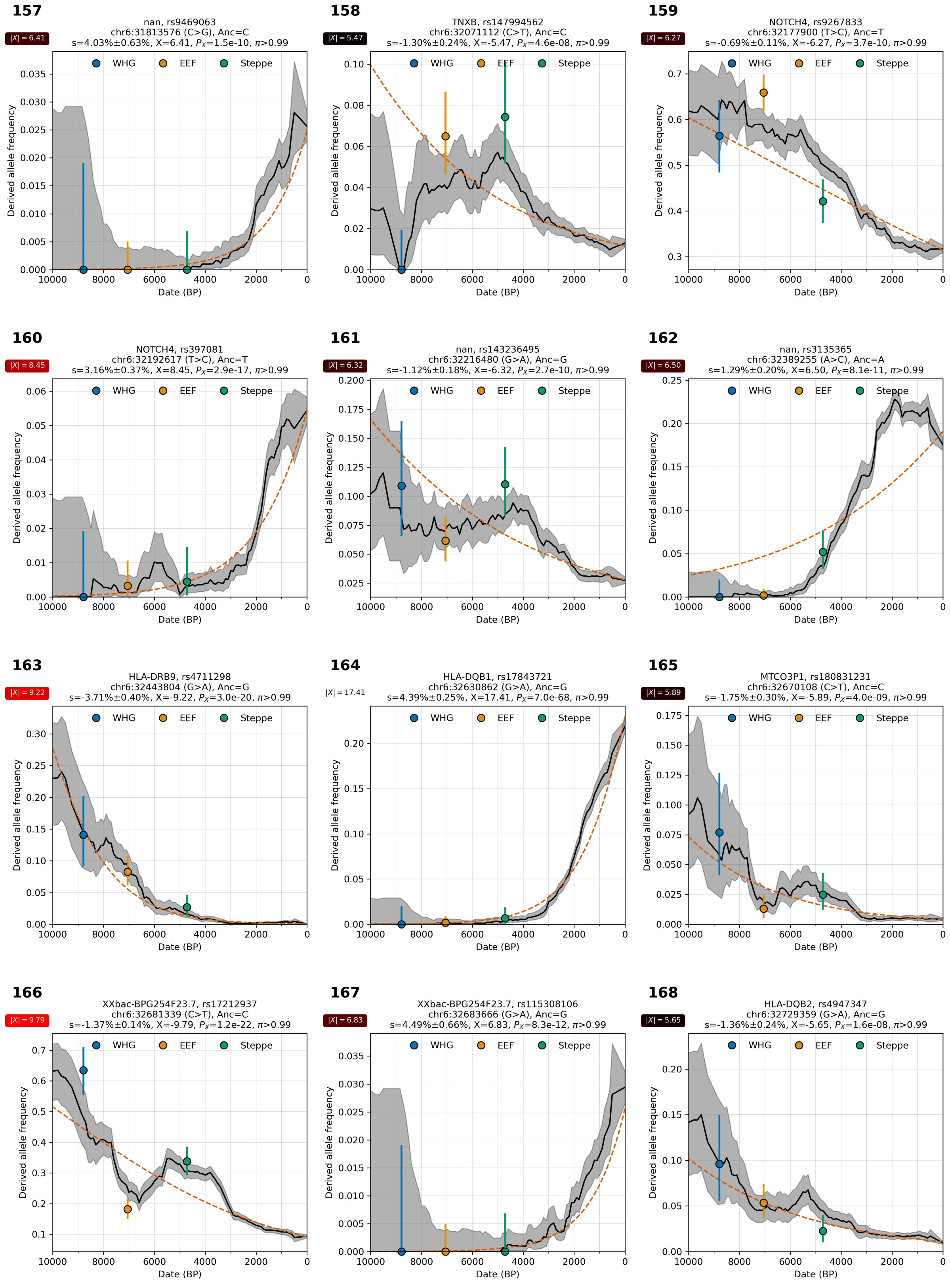

**Supplementary Figure S4.14:** Allele frequency over time. From chr 6.

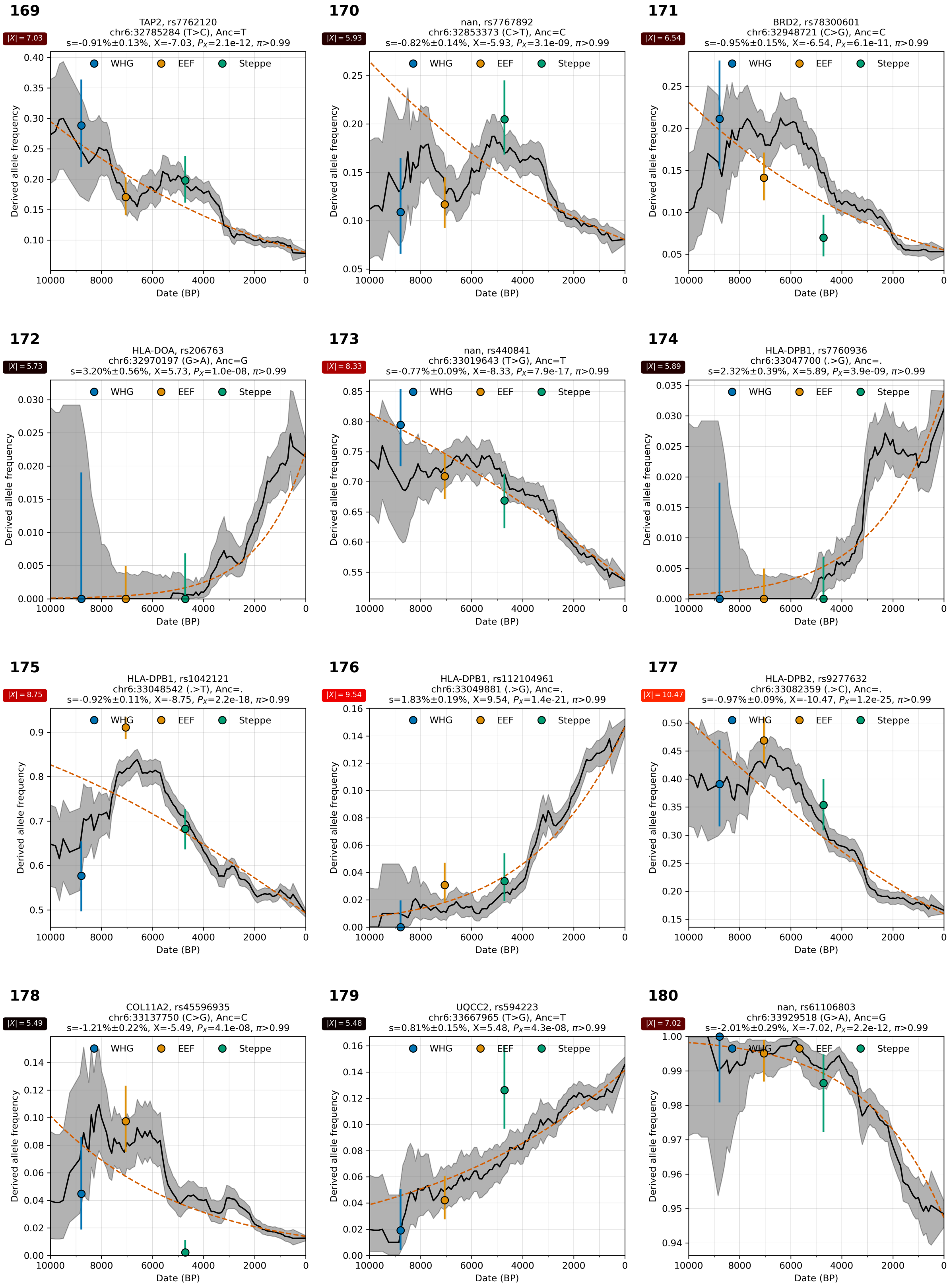

**Supplementary Figure S4.15:** Allele frequency over time. From chr 6.

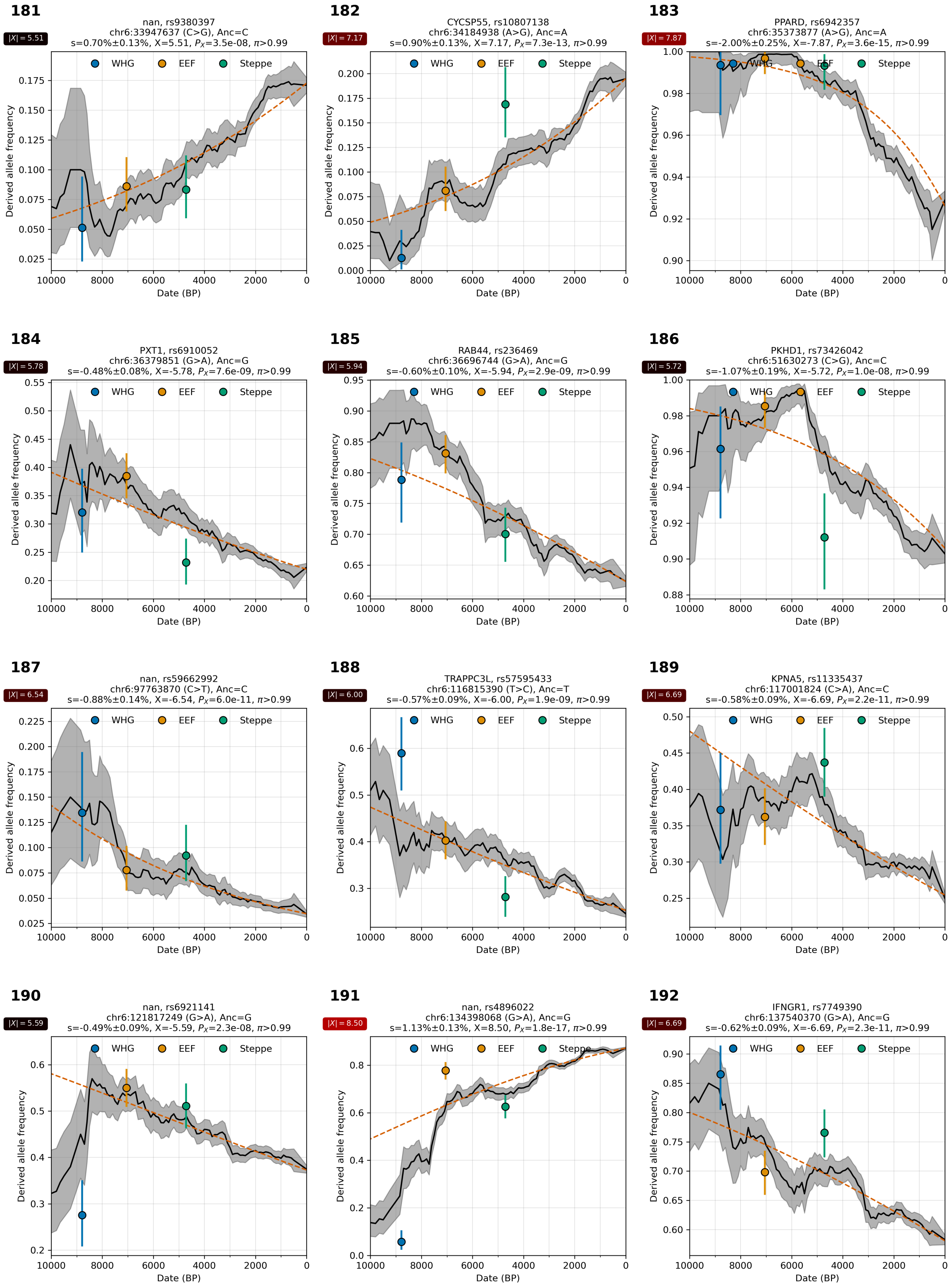

**Supplementary Figure S4.16:** Allele frequency over time. From chr 6.

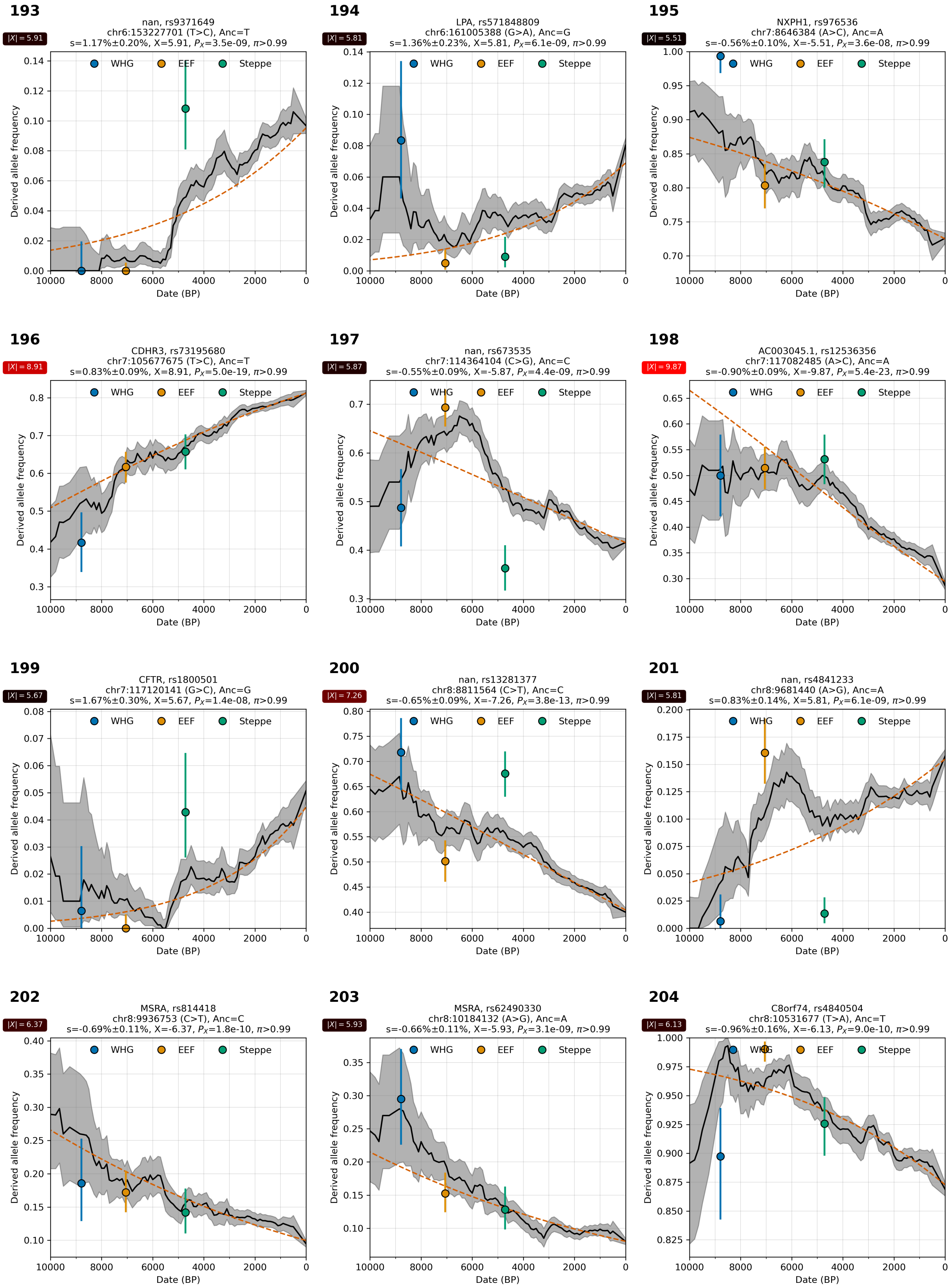

**Supplementary Figure S4.17:** Allele frequency over time. From chr 6, 7, and 8.

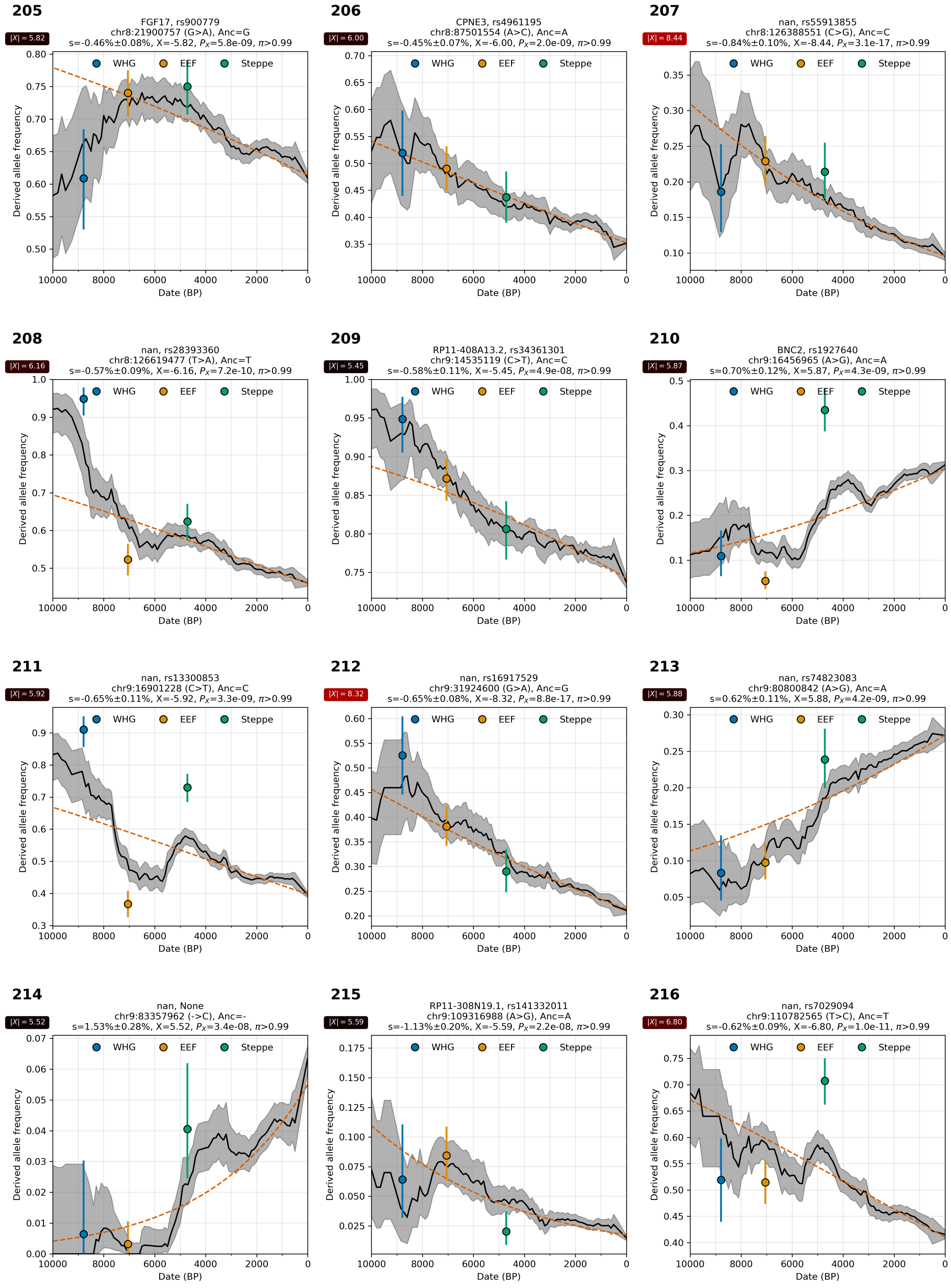

**Supplementary Figure S4.18:** Allele frequency over time. From chr 8 and 9.

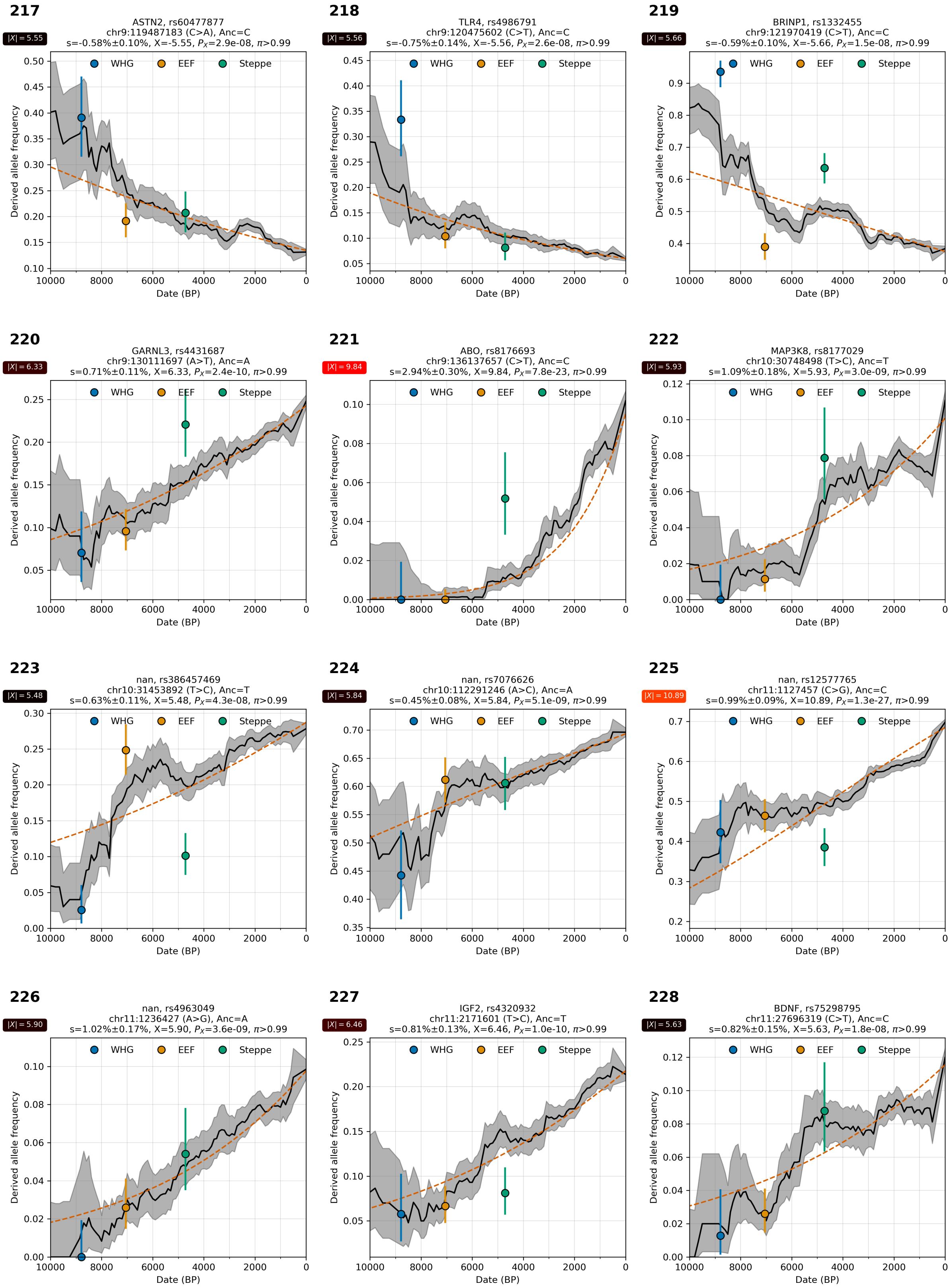

**Supplementary Figure S4.19:** Allele frequency over time. From chr 9, 10, and 11.

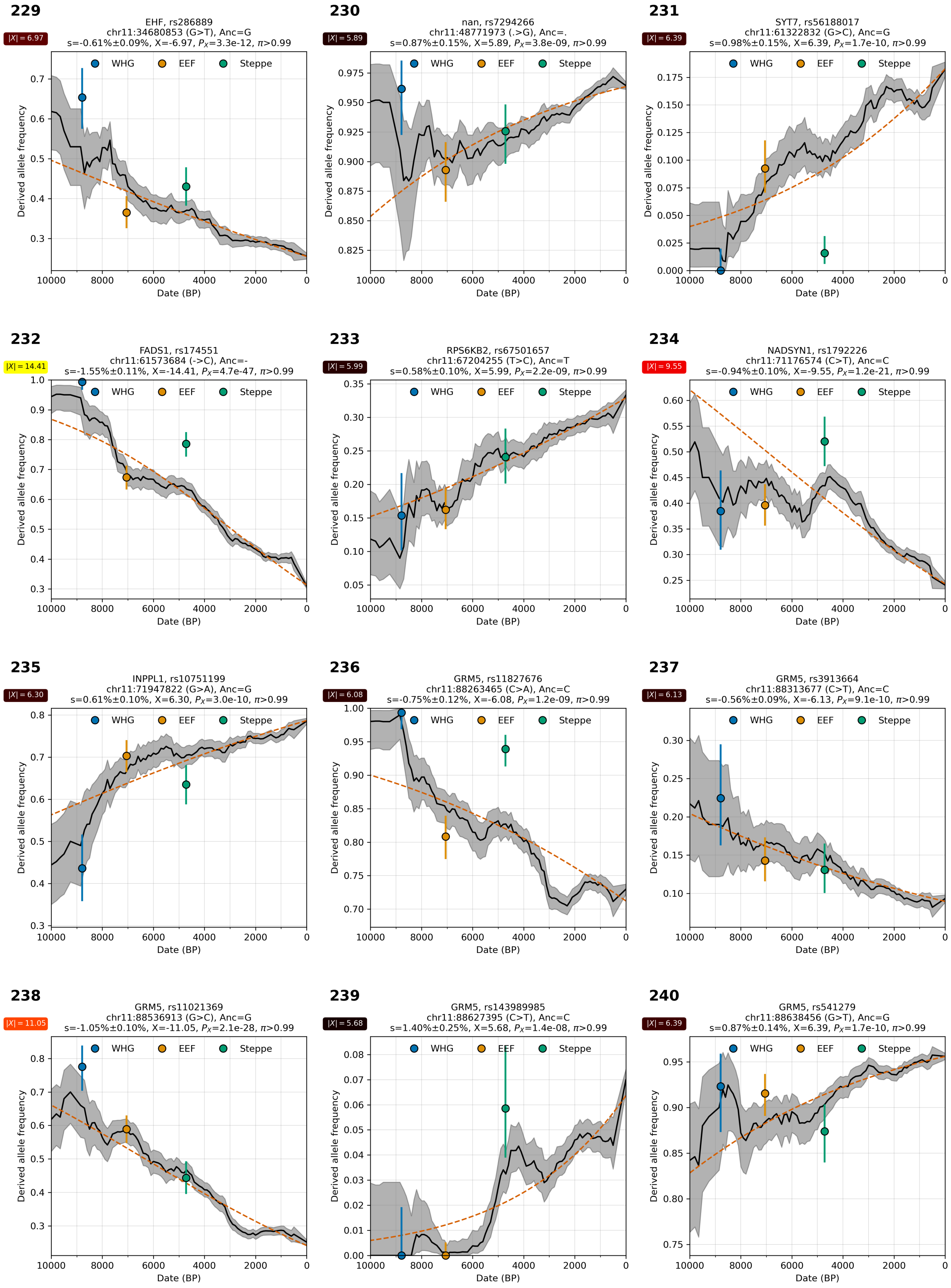

**Supplementary Figure S4.20:** Allele frequency over time. From chr 11.

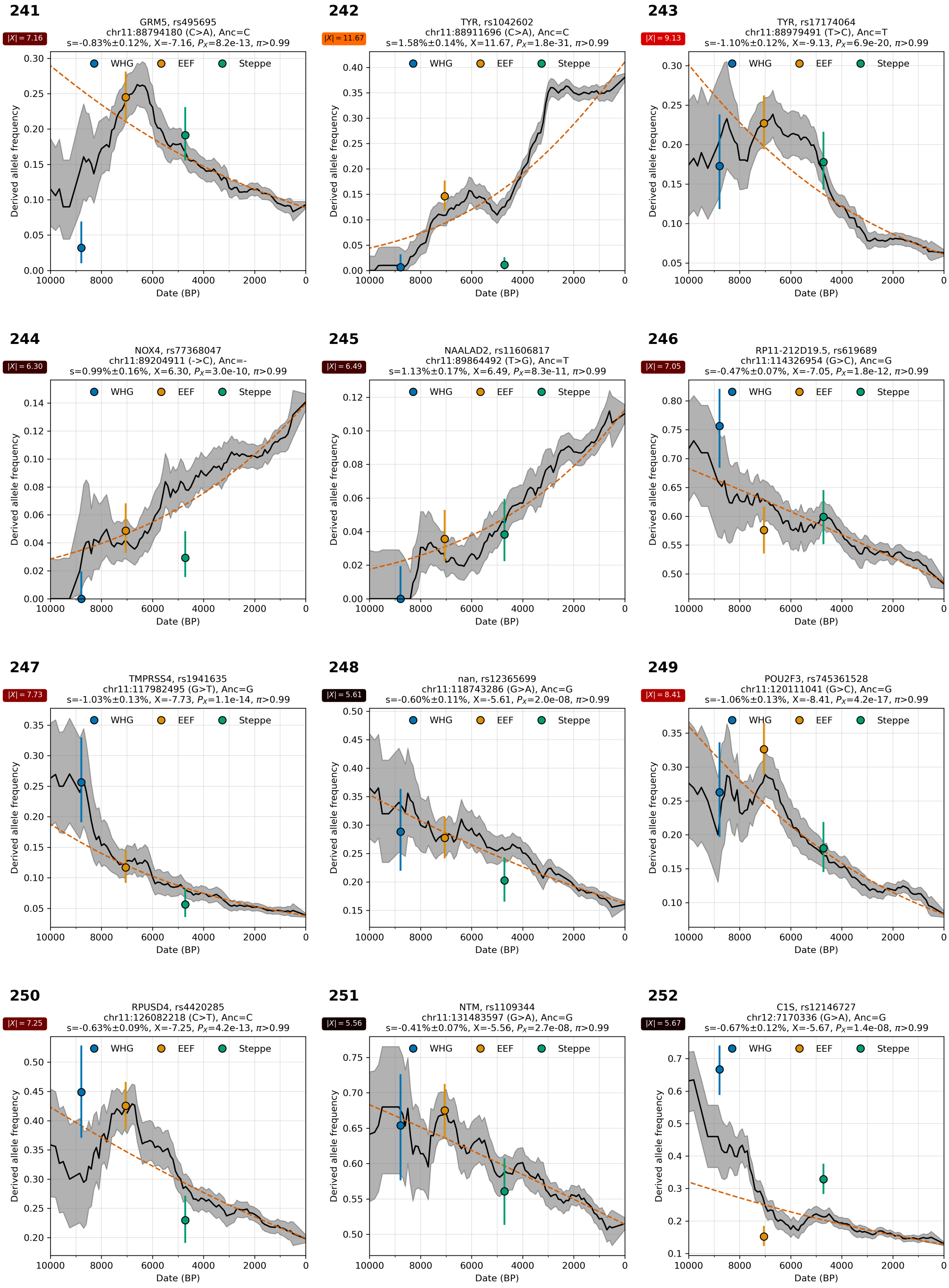

**Supplementary Figure S4.21:** Allele frequency over time. From chr 11 and 12.

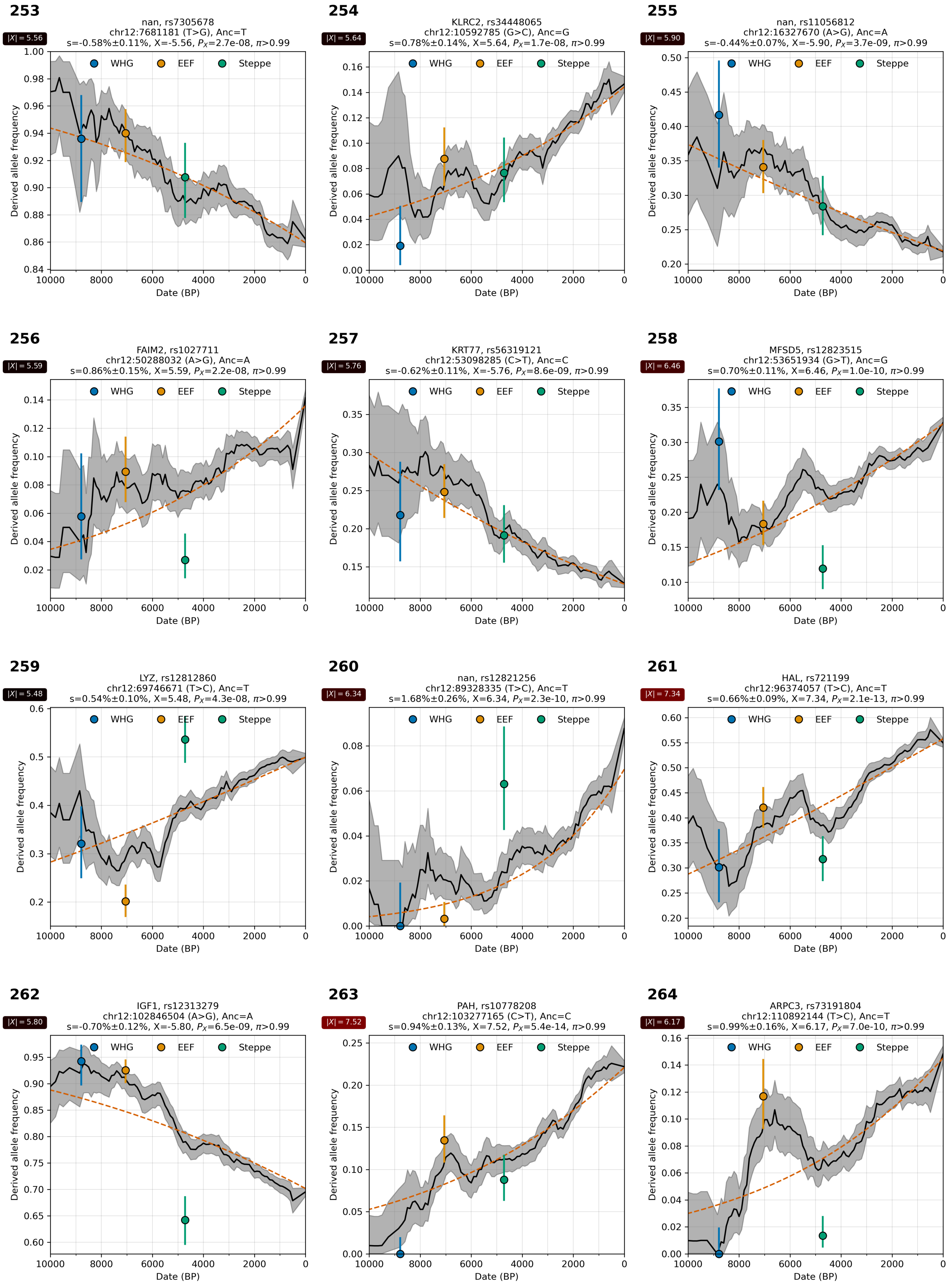

**Supplementary Figure S4.22:** Allele frequency over time. From chr 12.

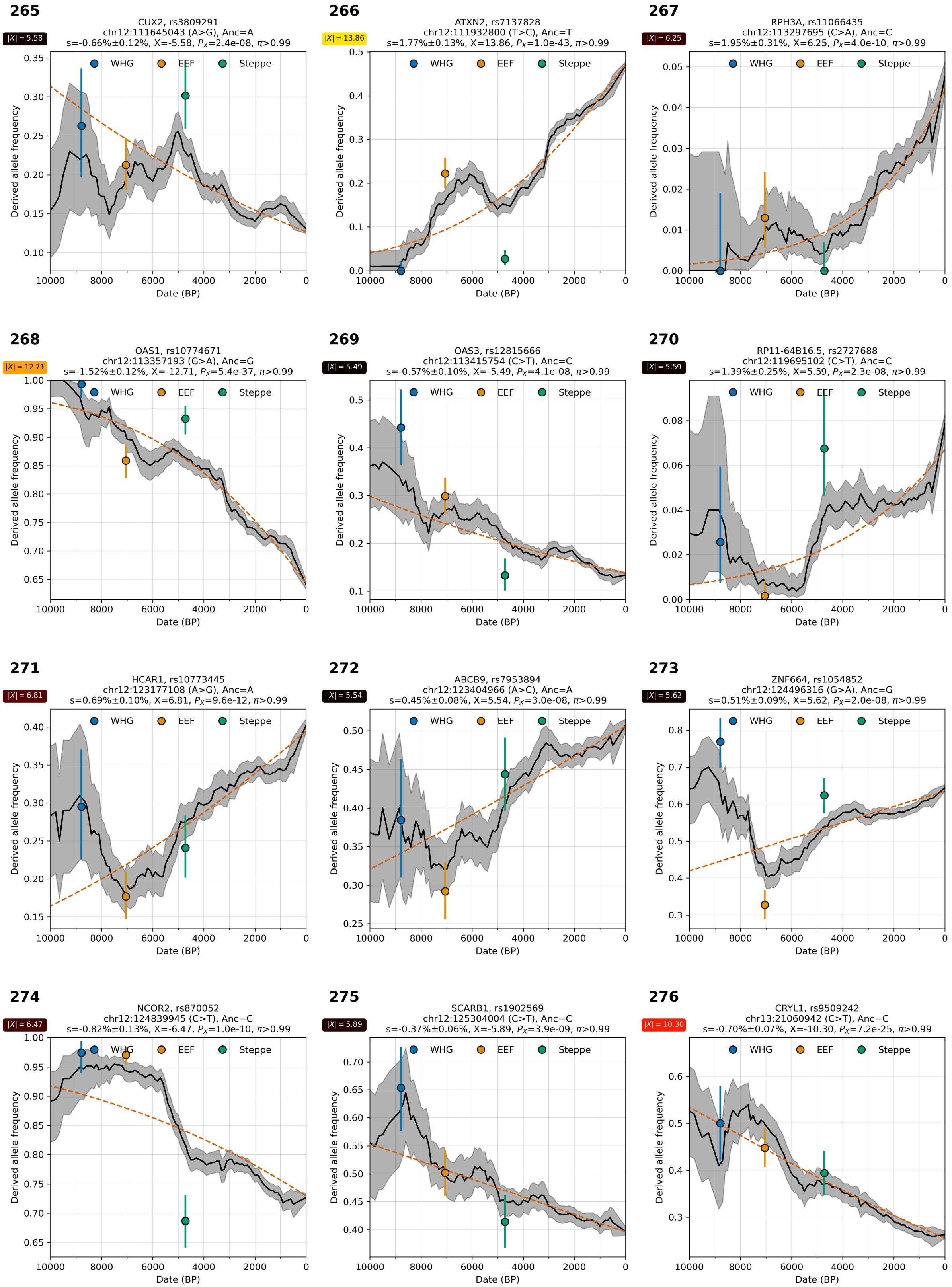

**Supplementary Figure S4.23:** Allele frequency over time. From chr 12 and 13.

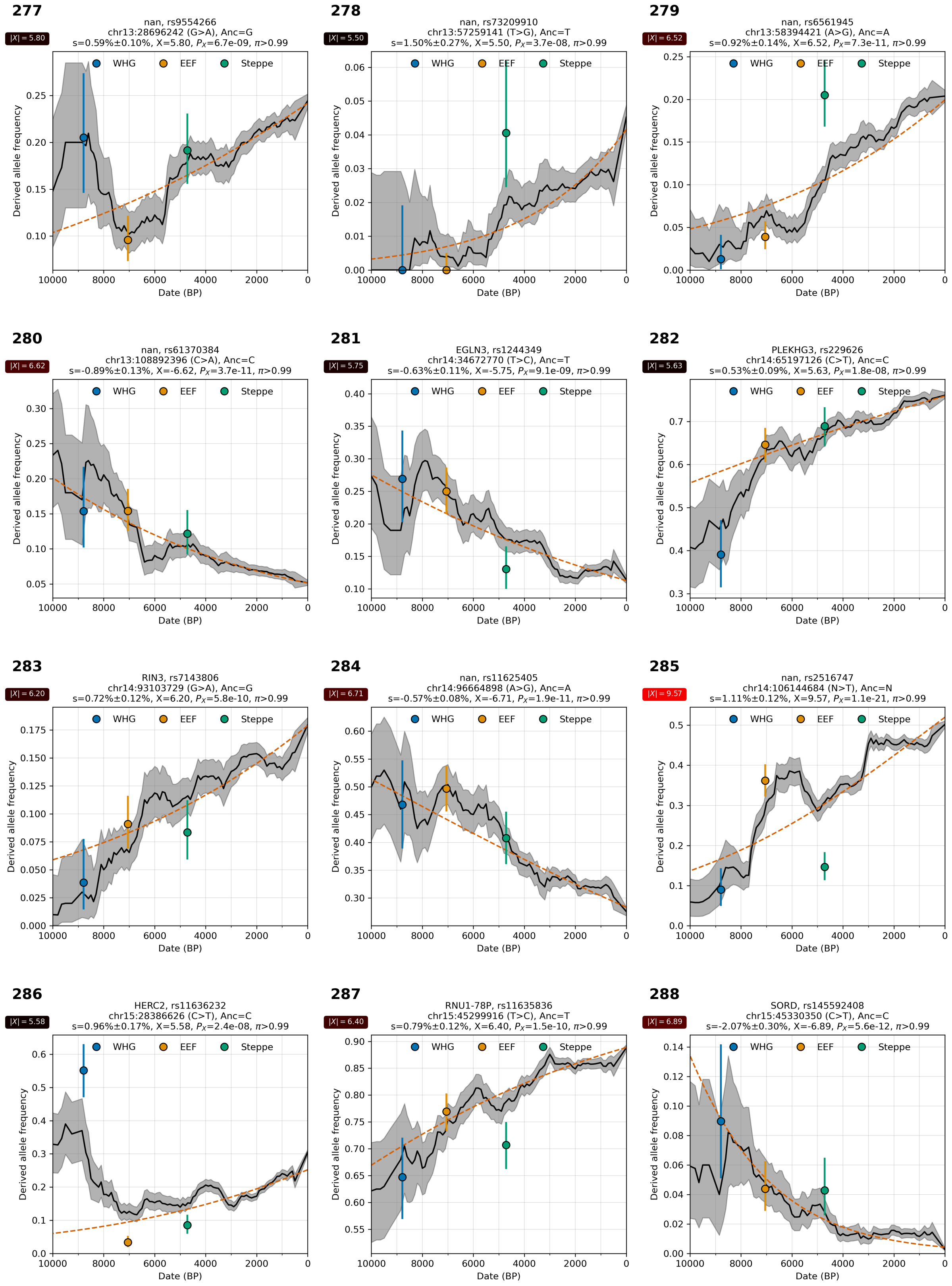

**Supplementary Figure S4.24:** Allele frequency over time. From chr 13, 14, and 15.

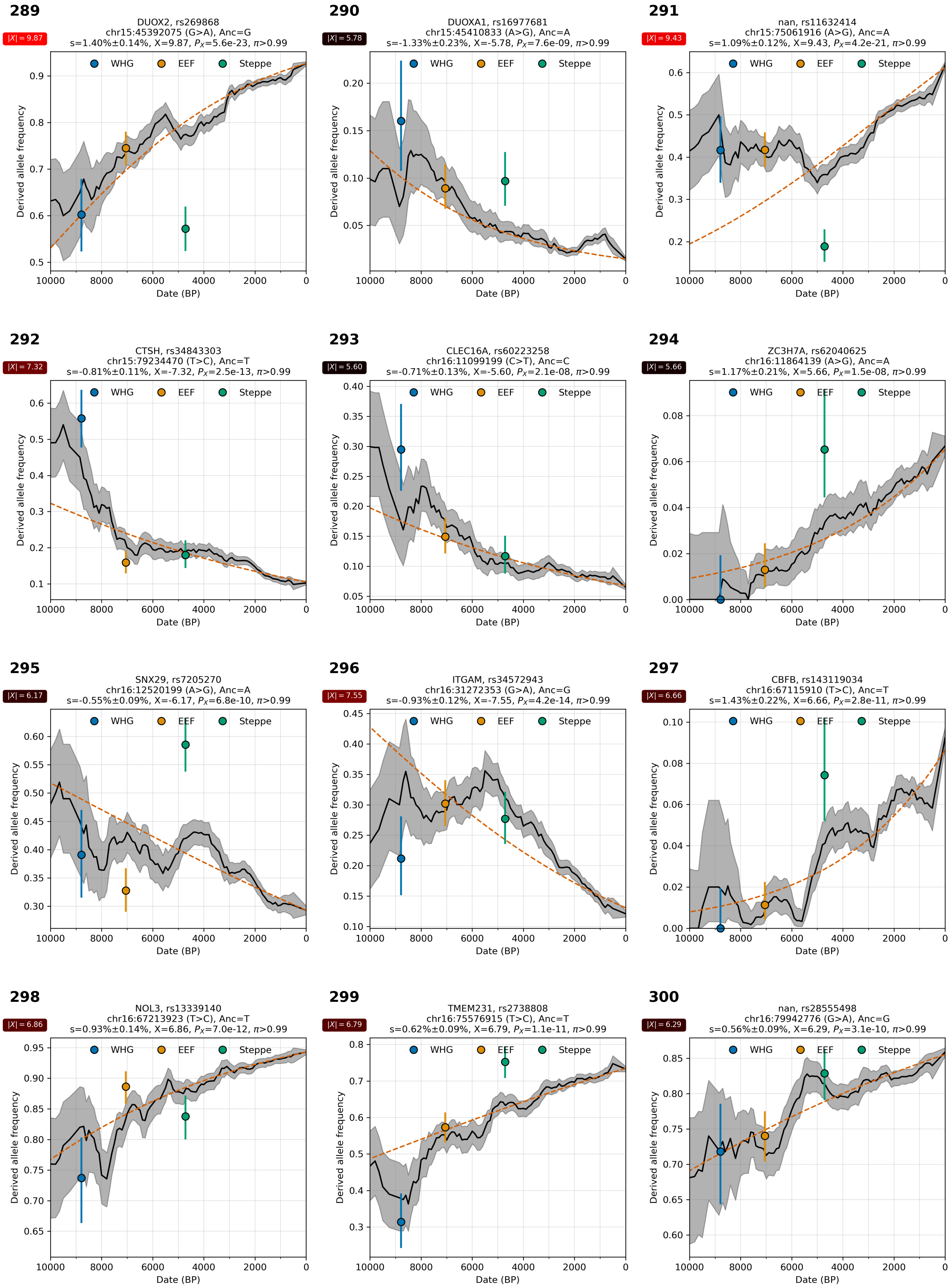

**Supplementary Figure S4.25:** Allele frequency over time. From chr 15 and 16.

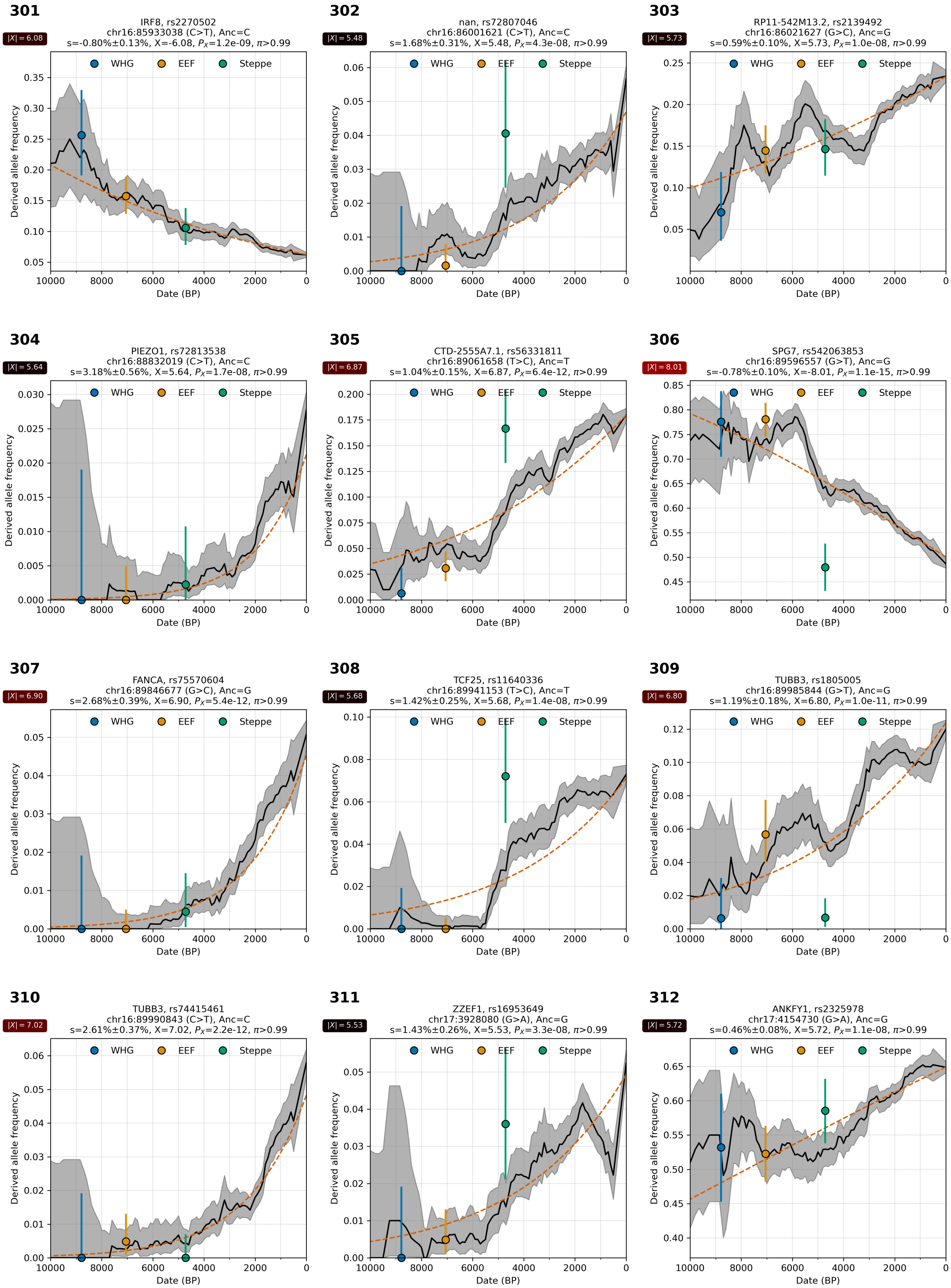

**Supplementary Figure S4.26:** Allele frequency over time. From chr 16 and 17.

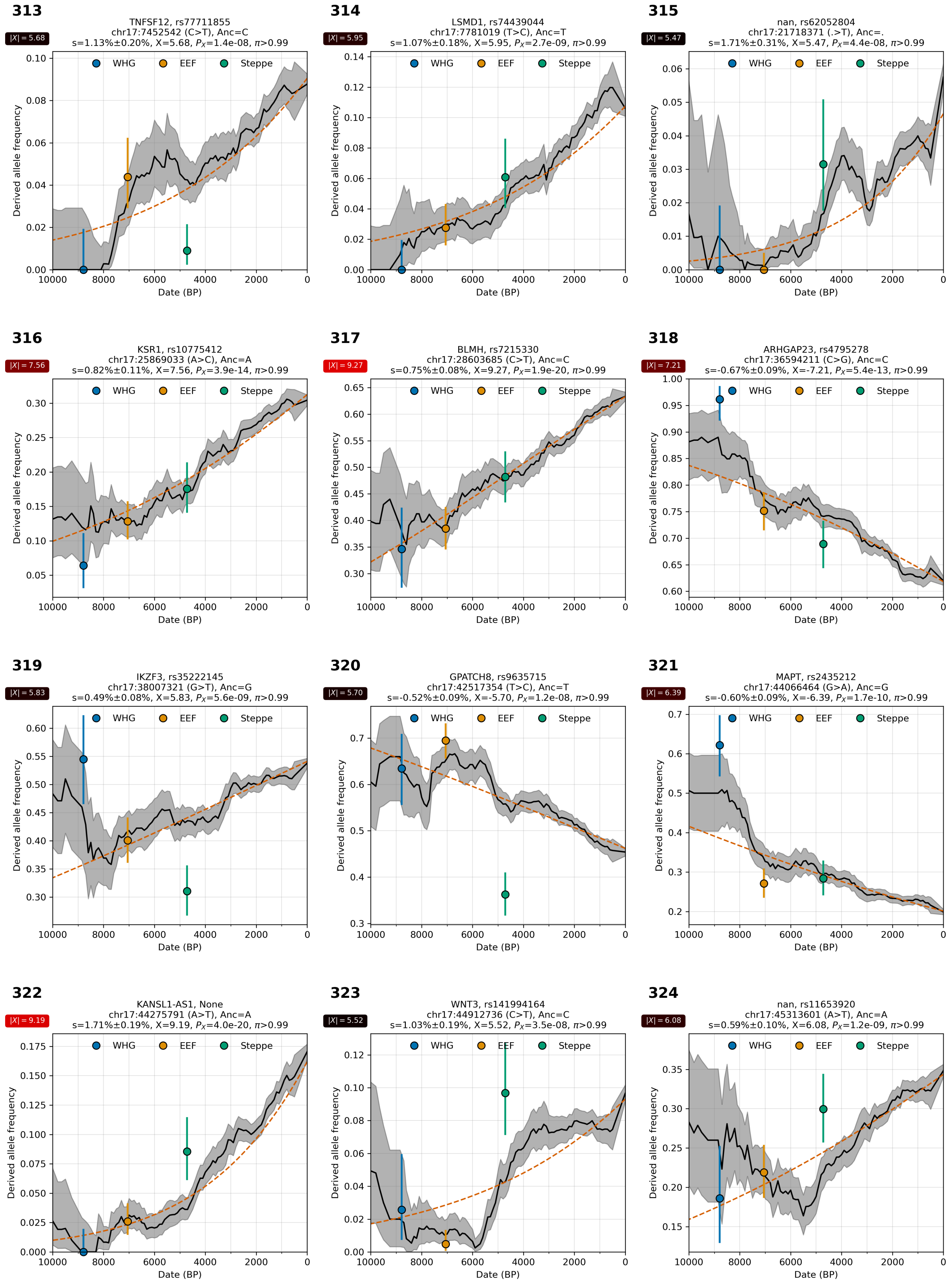

**Supplementary Figure S4.27:** Allele frequency over time. From chr 17.

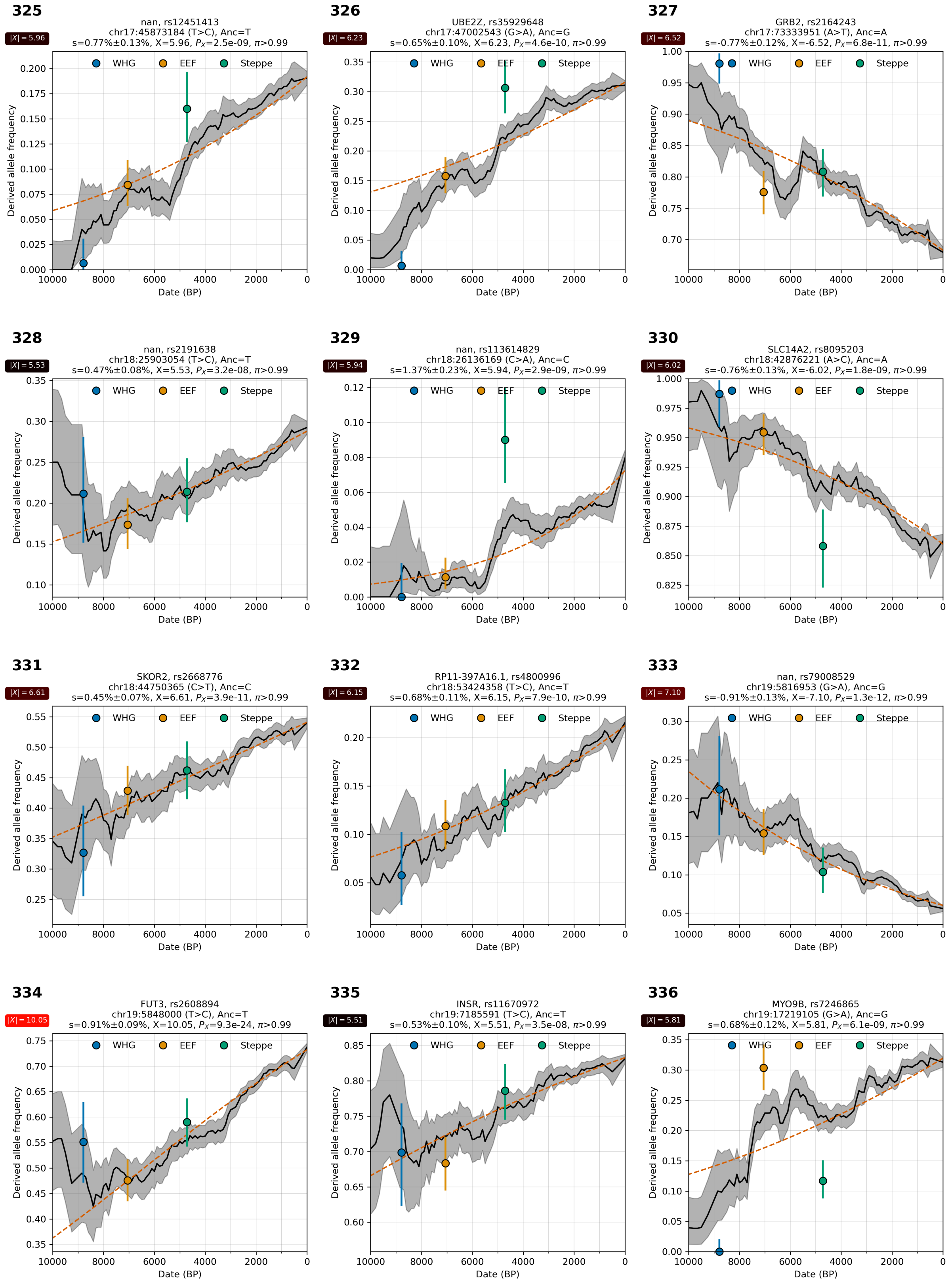

**Supplementary Figure S4.28:** Allele frequency over time. From chr 17, 18, and 19.

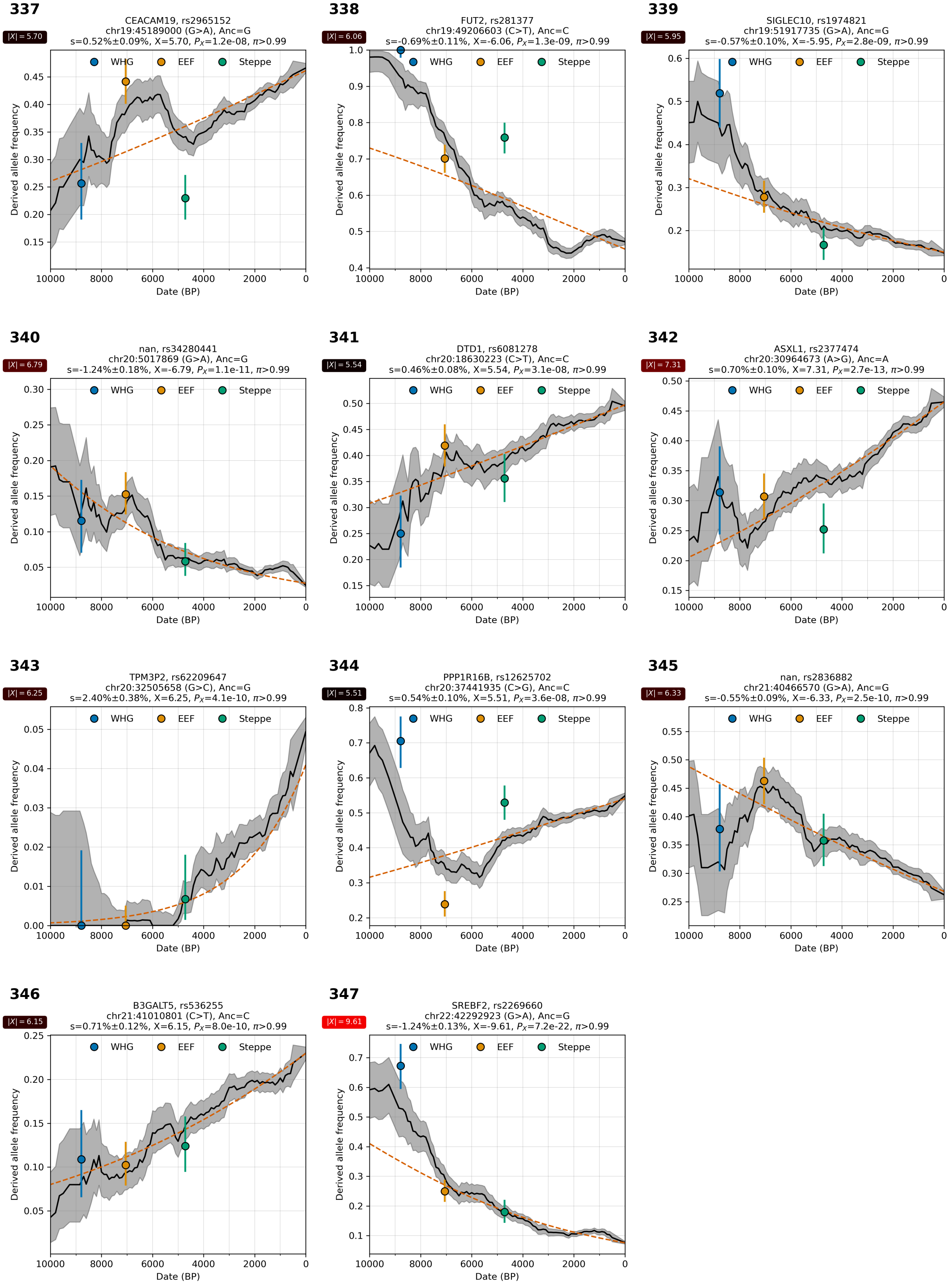

**Supplementary Figure S4.29:** Allele frequency over time. From chr 19, 20, 21, 22.

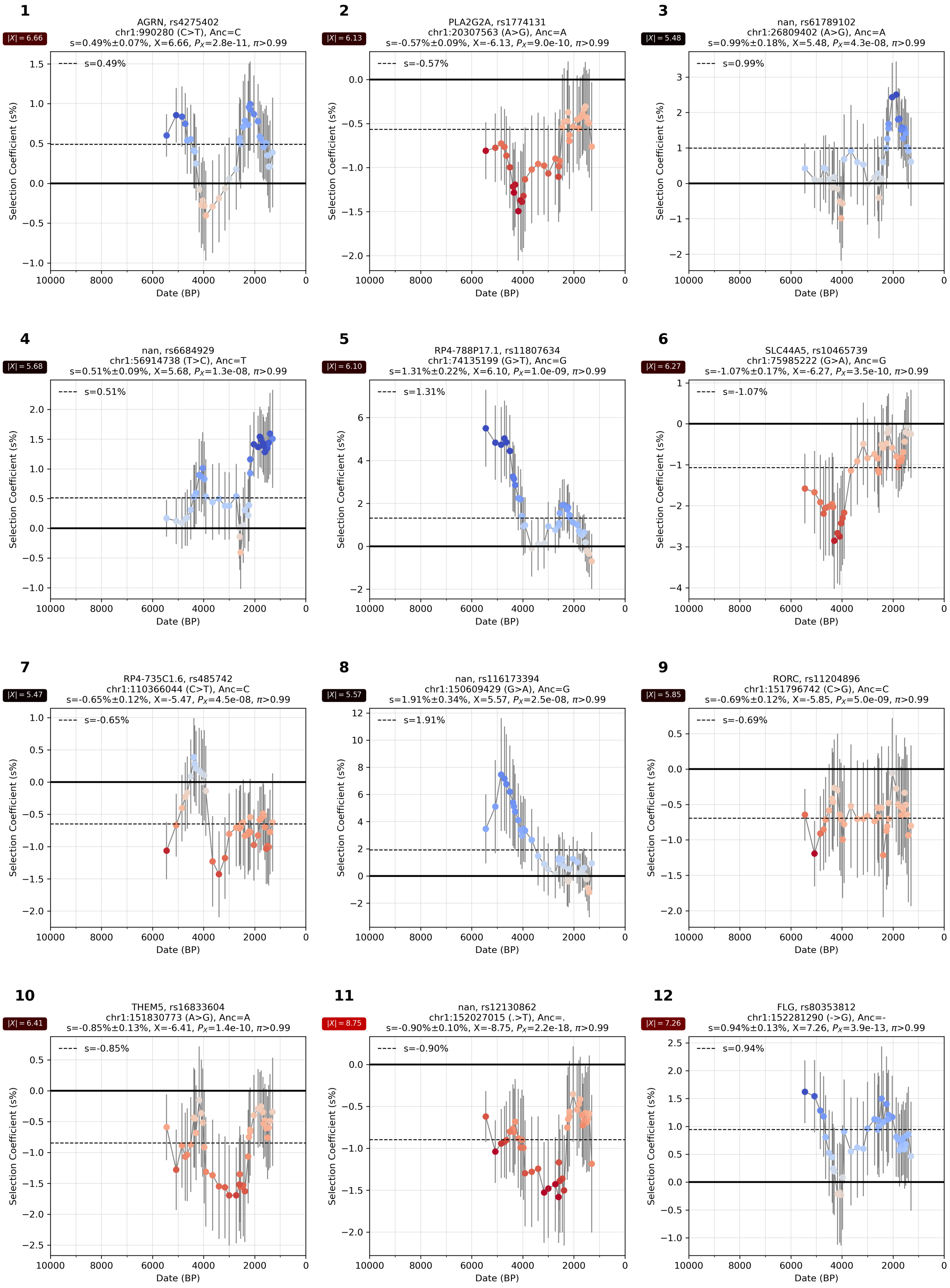

**Supplementary Figure S4.30:** Selection coefficient over time. From chr 1.

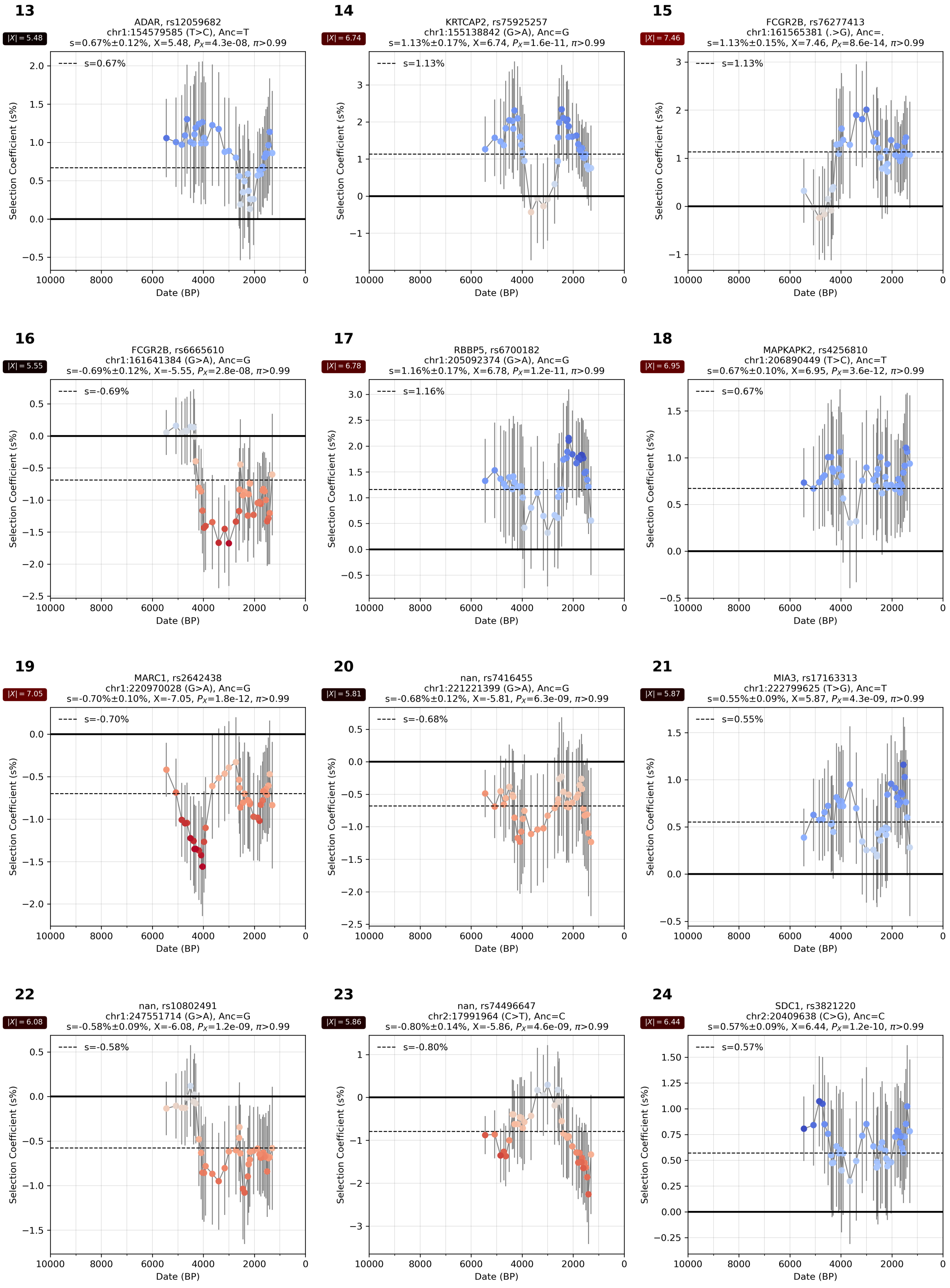

**Supplementary Figure S4.31:** Selection coefficient over time. From chr 1 and 2.

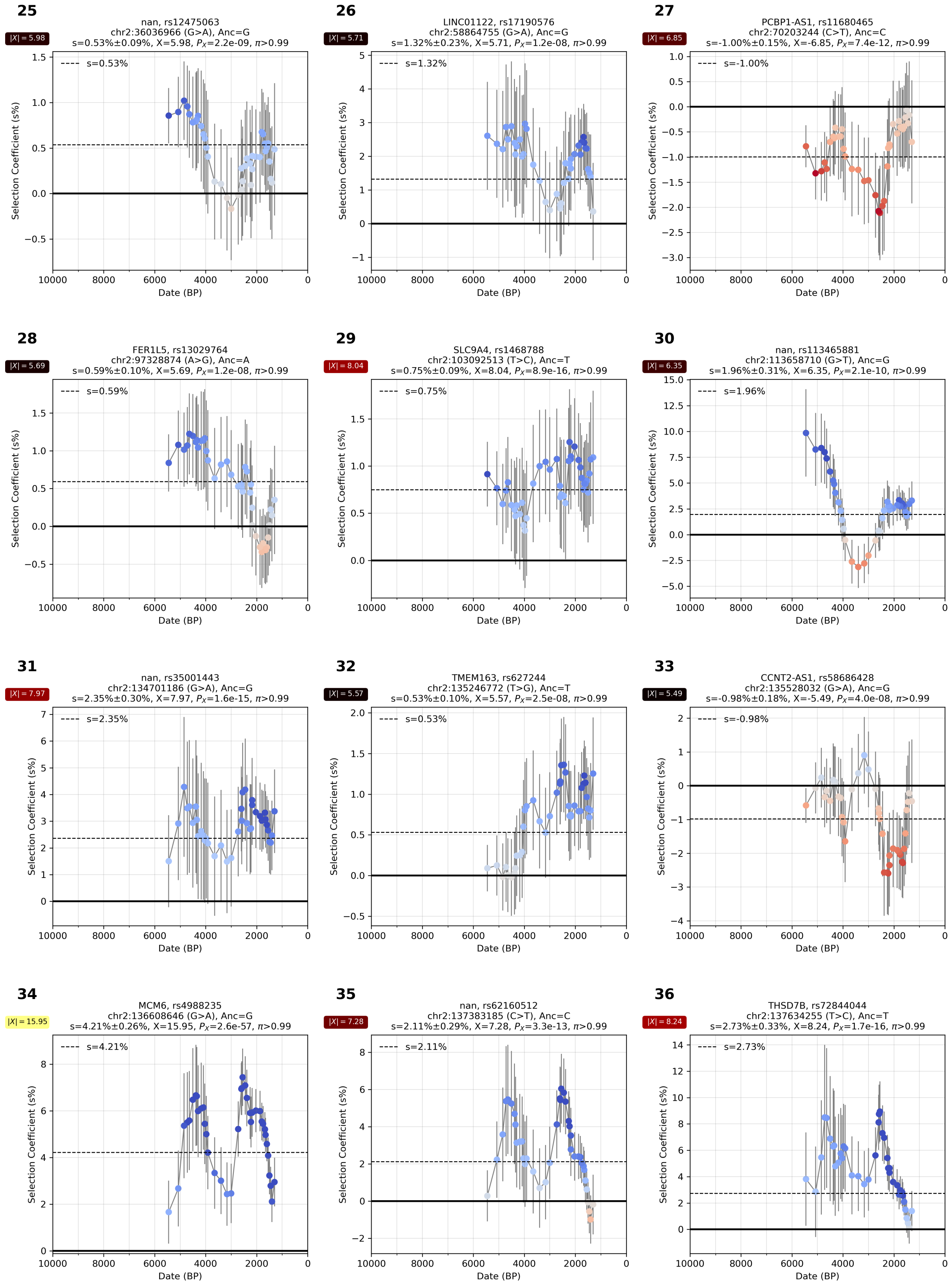

**Supplementary Figure S4.32:** Selection coefficient over time. From chr 2.

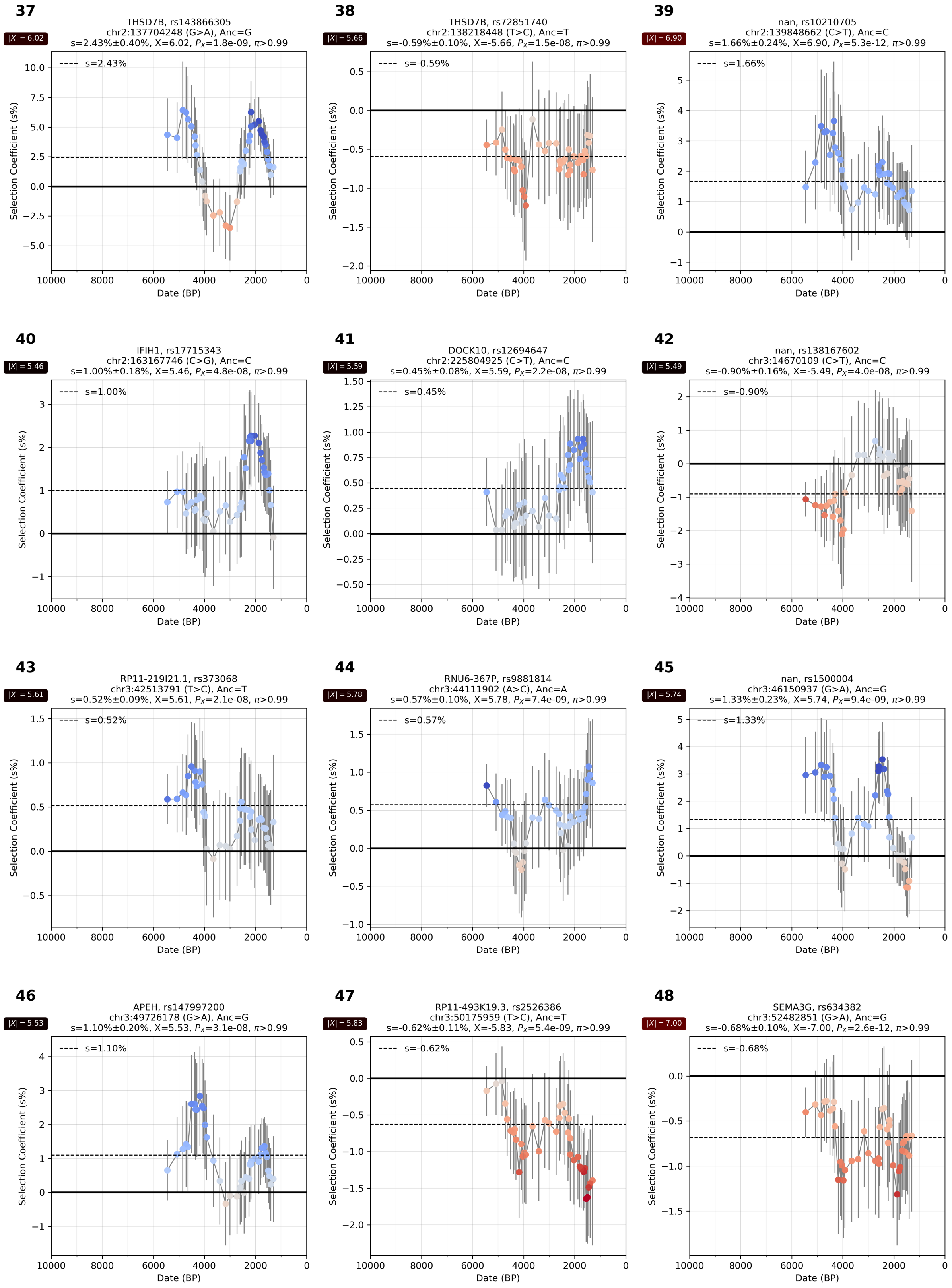

**Supplementary Figure S4.33:** Selection coefficient over time. From chr 2 and 3.

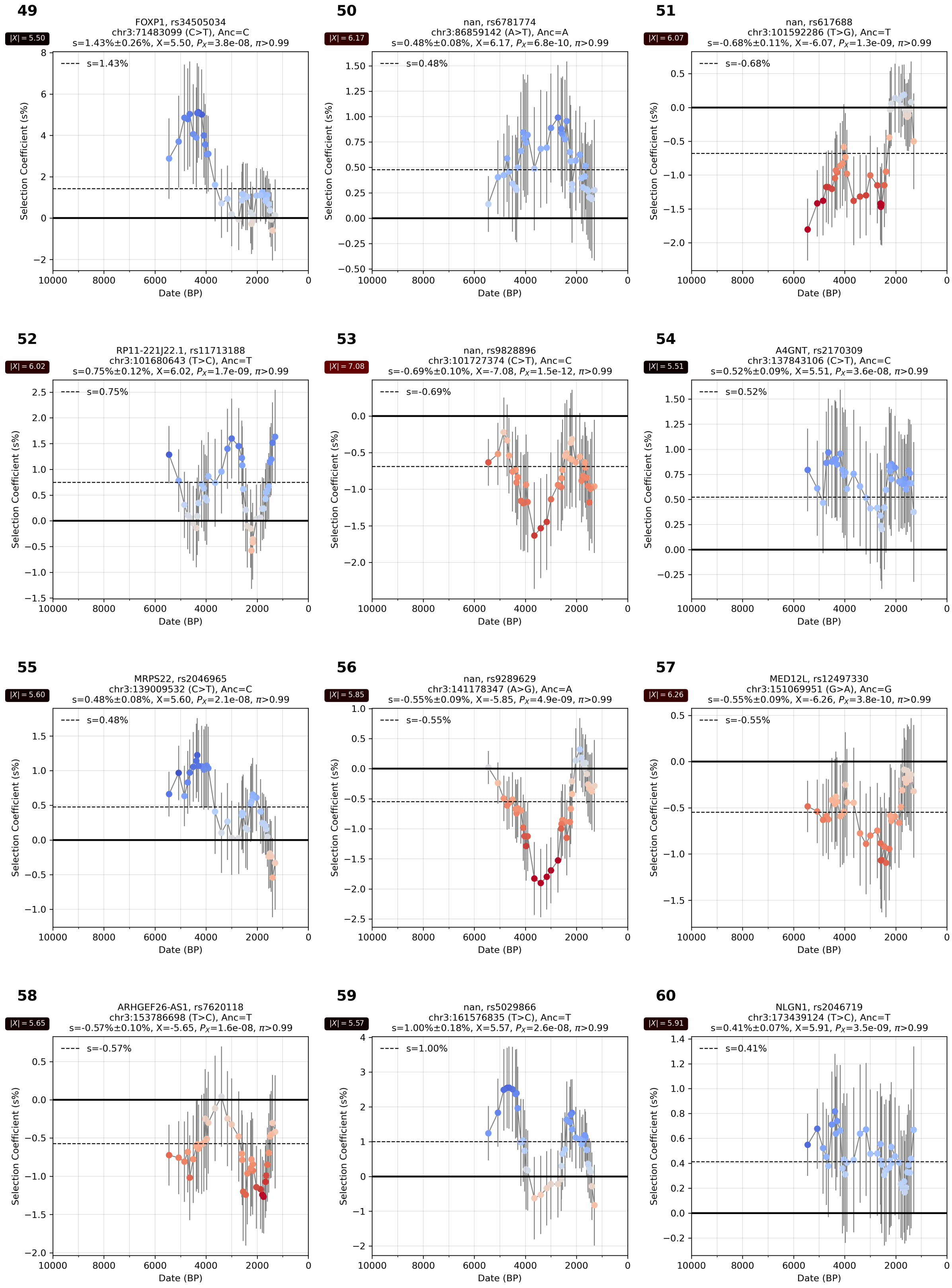

**Supplementary Figure S4.34:** Selection coefficient over time. From chr 3.

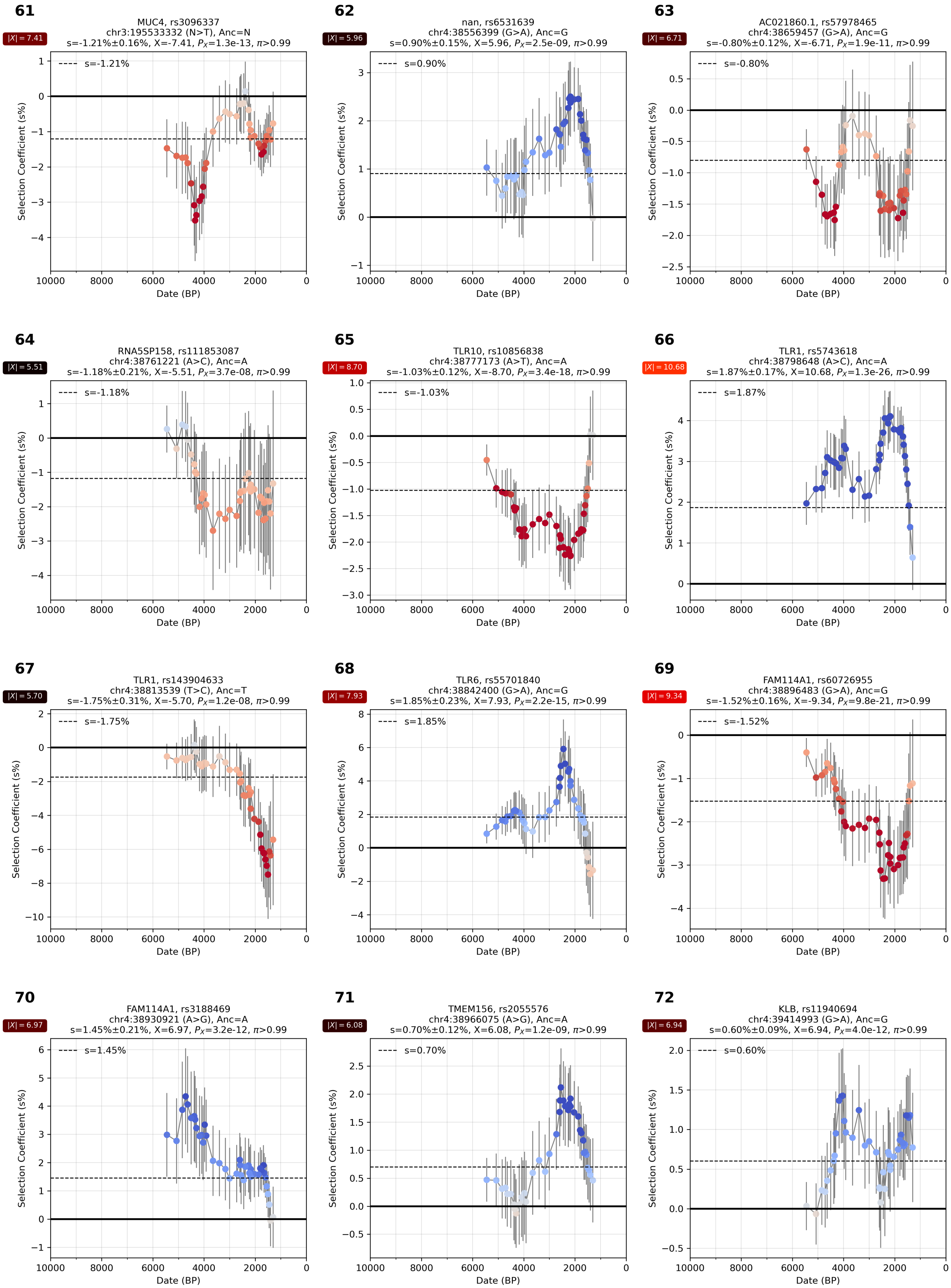

**Supplementary Figure S4.35:** Selection coefficient over time. From chr 3 and 4.

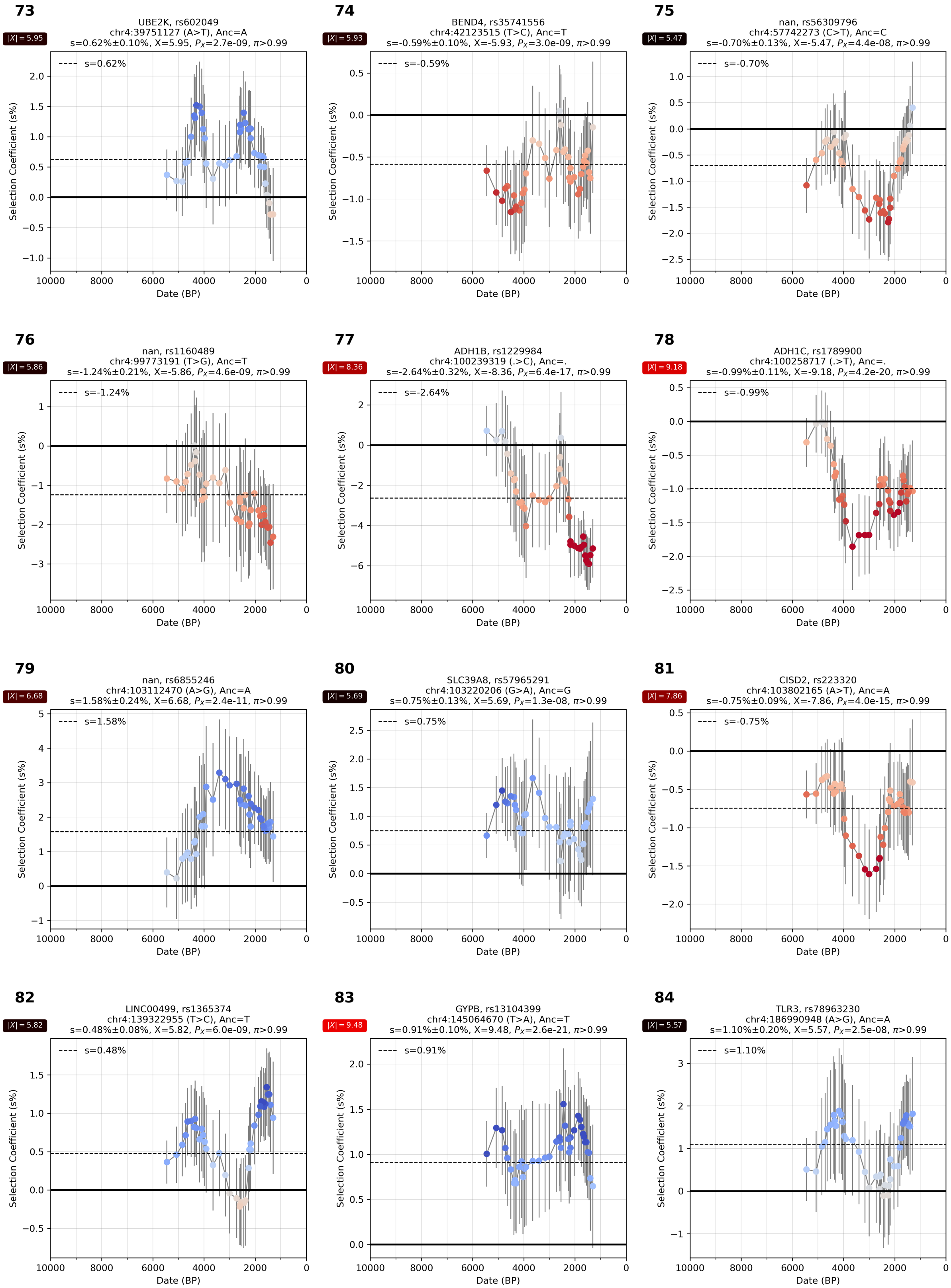

**Supplementary Figure S4.36:** Selection coefficient over time. From chr 4.

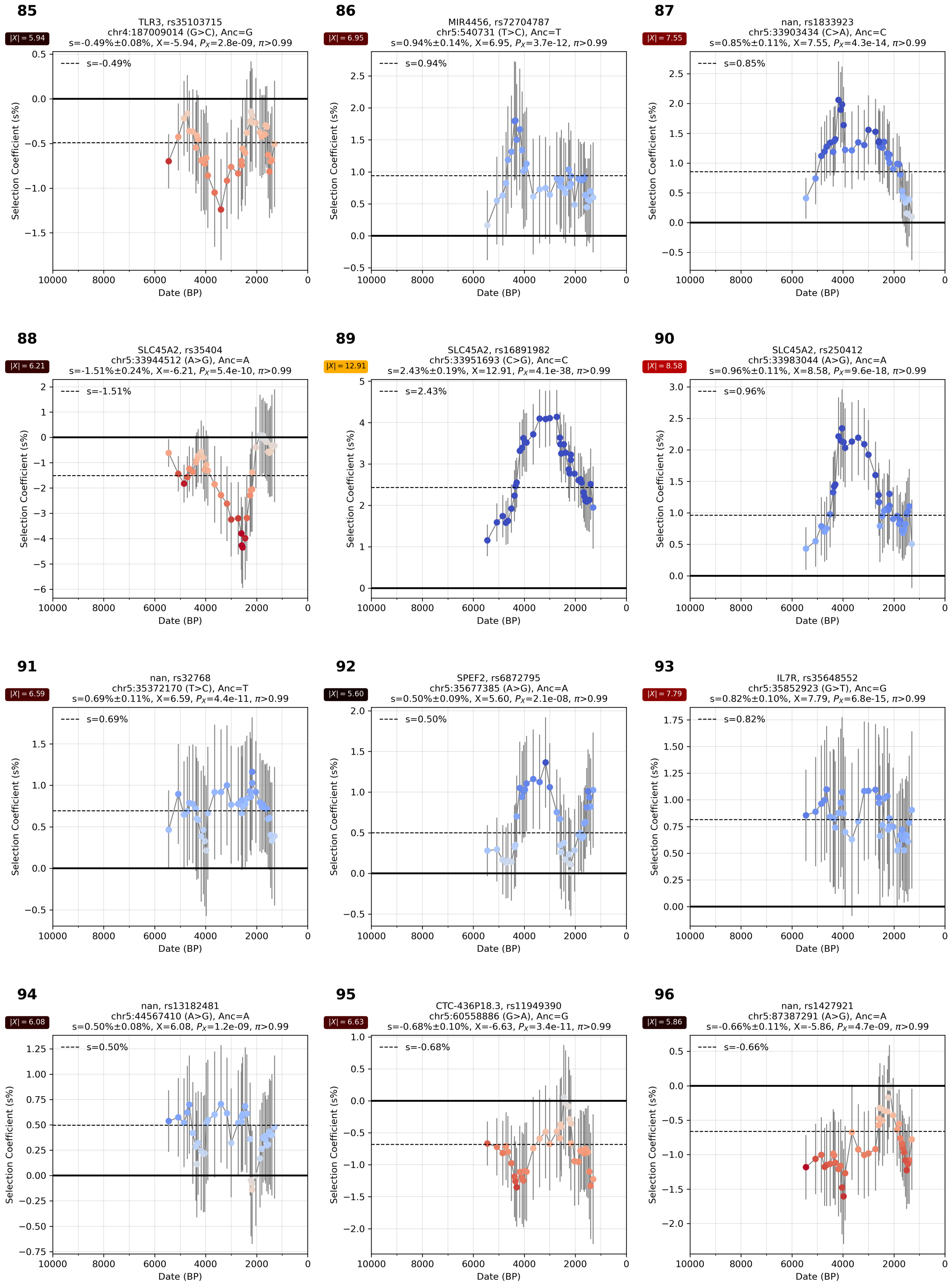

**Supplementary Figure S4.37:** Selection coefficient over time. From chr 4 and 5.

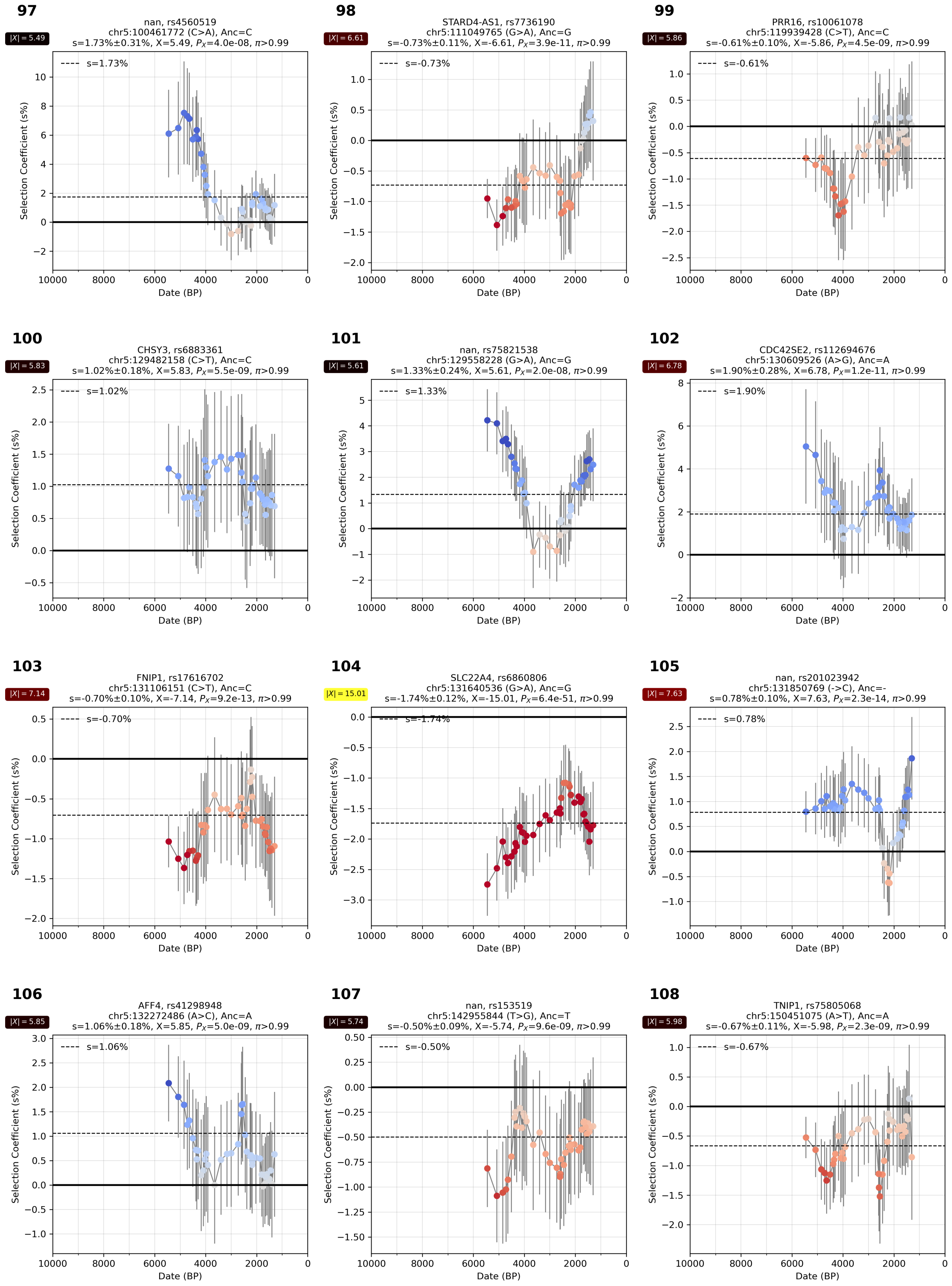

**Supplementary Figure S4.38:** Selection coefficient over time. From chr 5.

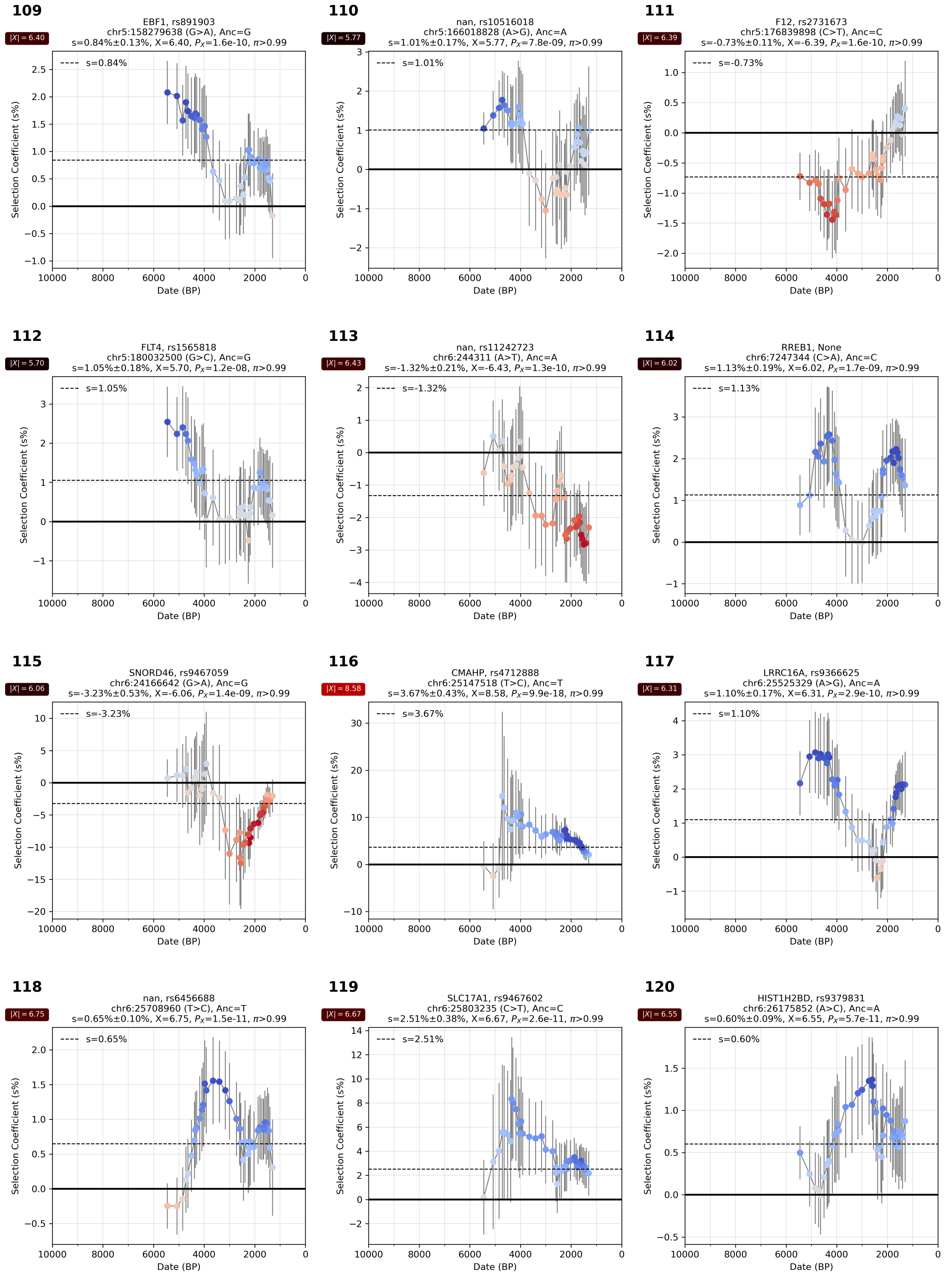

**Supplementary Figure S4.39:** Selection coefficient over time. From chr 5 and 6.

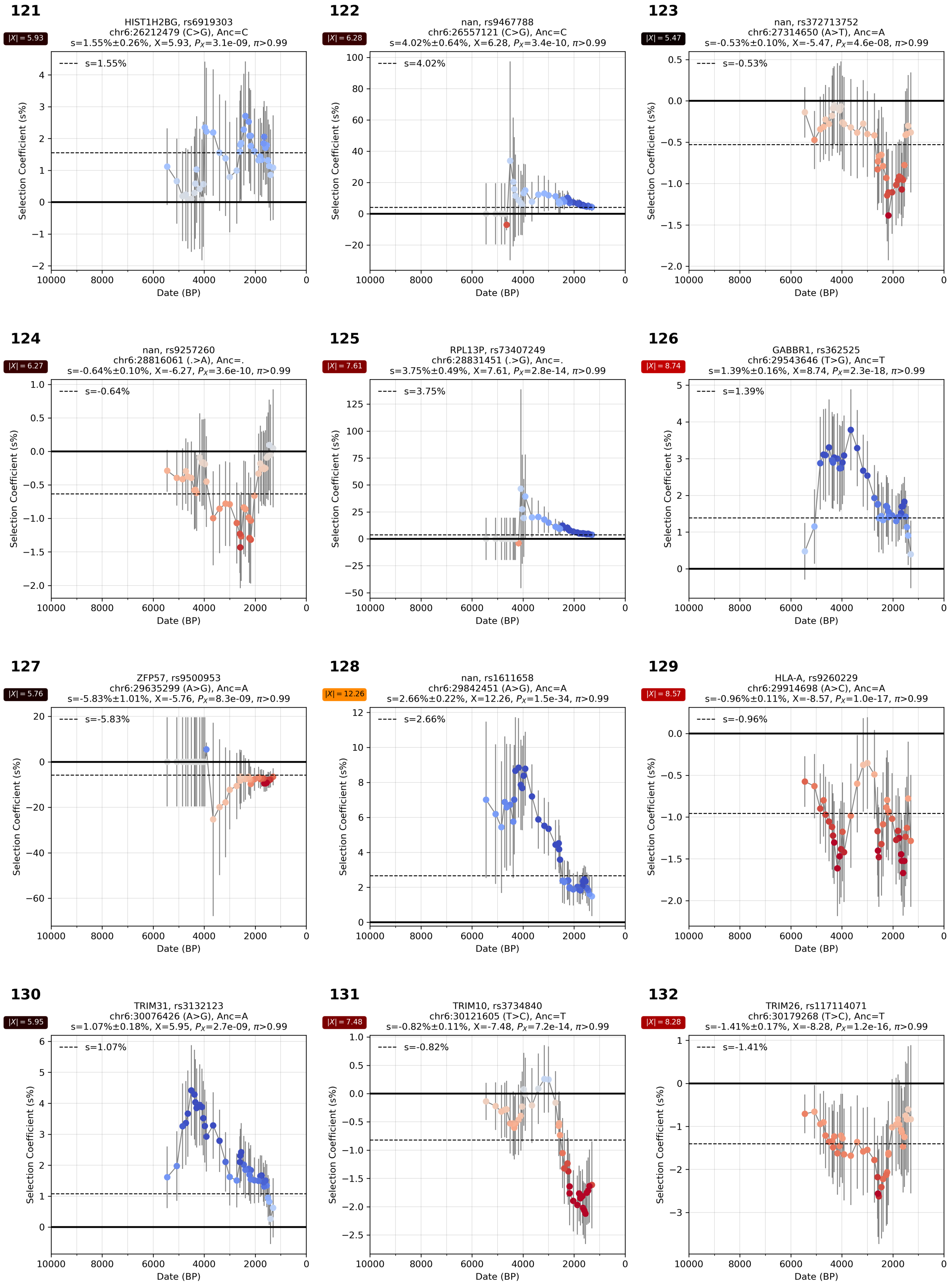

**Supplementary Figure S4.40:** Selection coefficient over time. From chr 6.

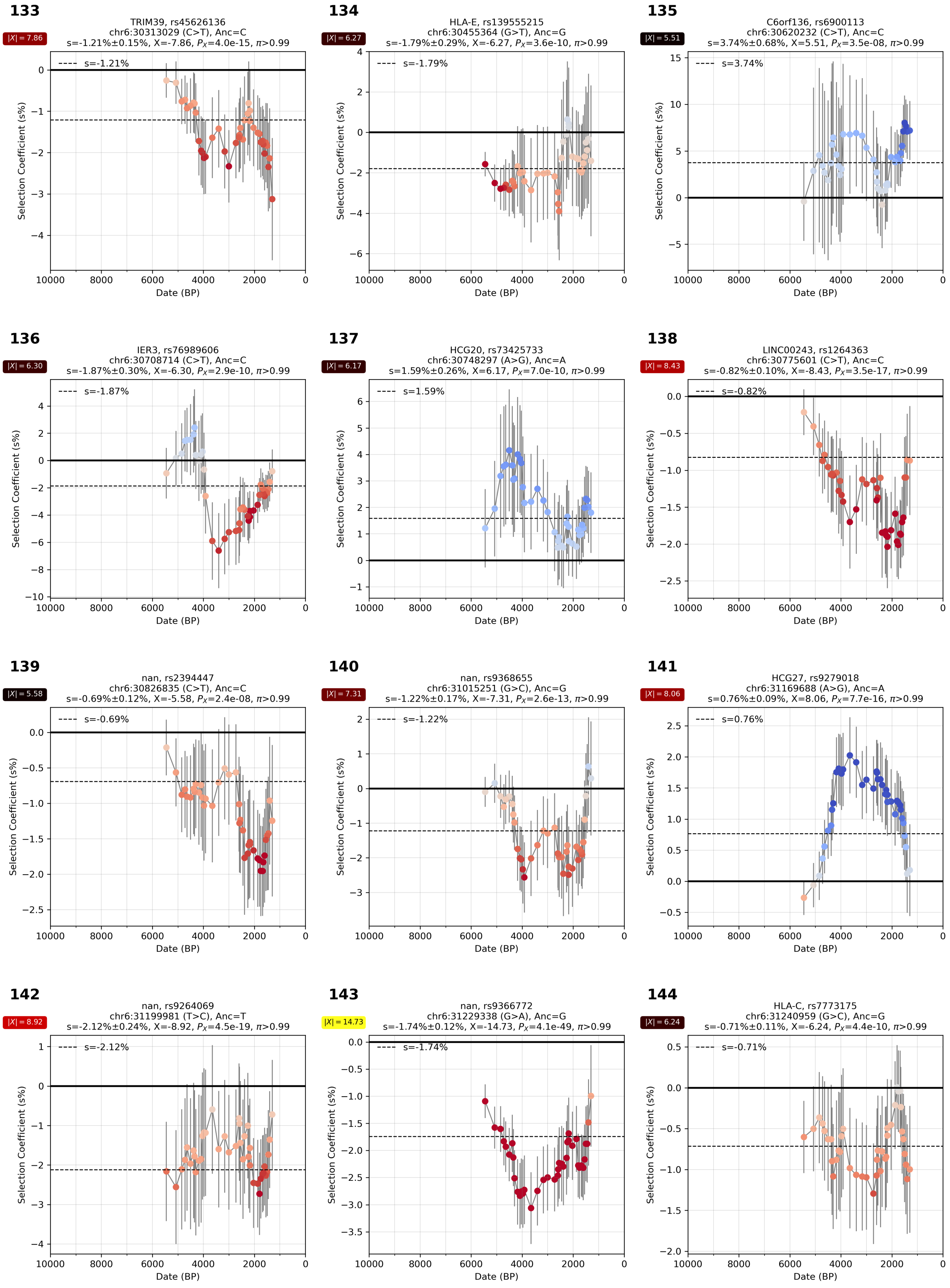

**Supplementary Figure S4.41:** Selection coefficient over time. From chr 6.

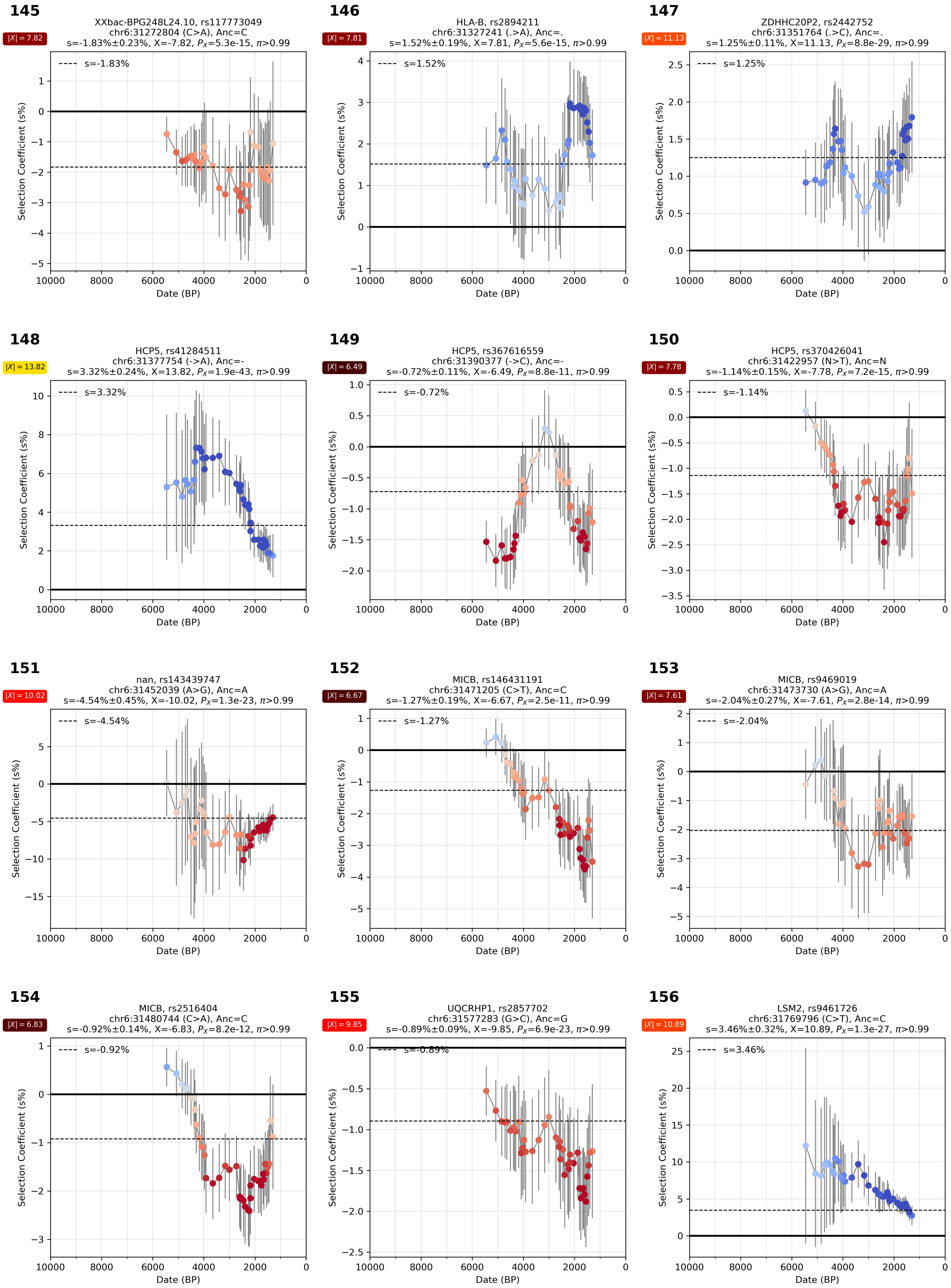

**Supplementary Figure S4.42:** Selection coefficient over time. From chr 6.

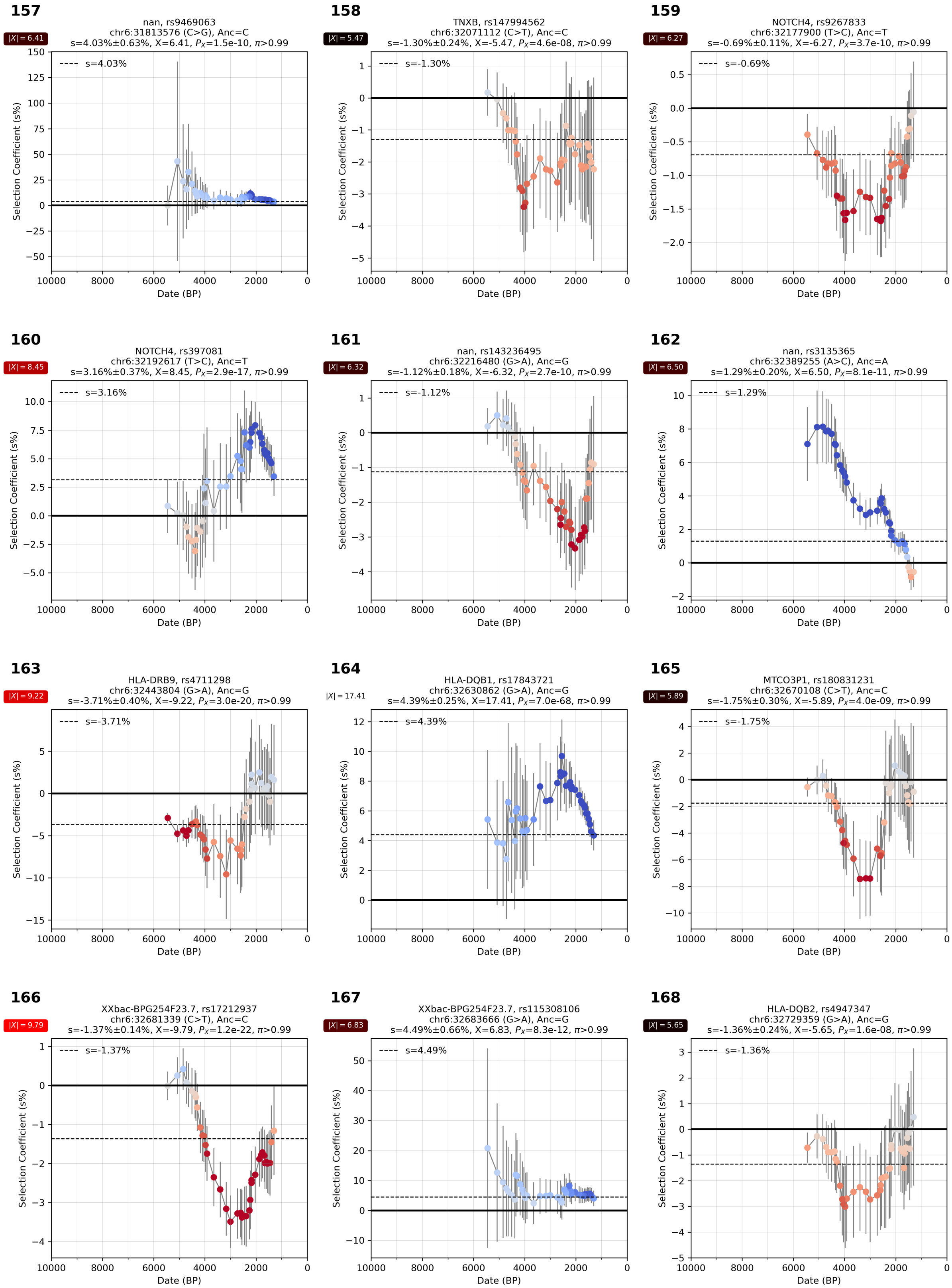

**Supplementary Figure S4.43:** Selection coefficient over time. From chr 6.

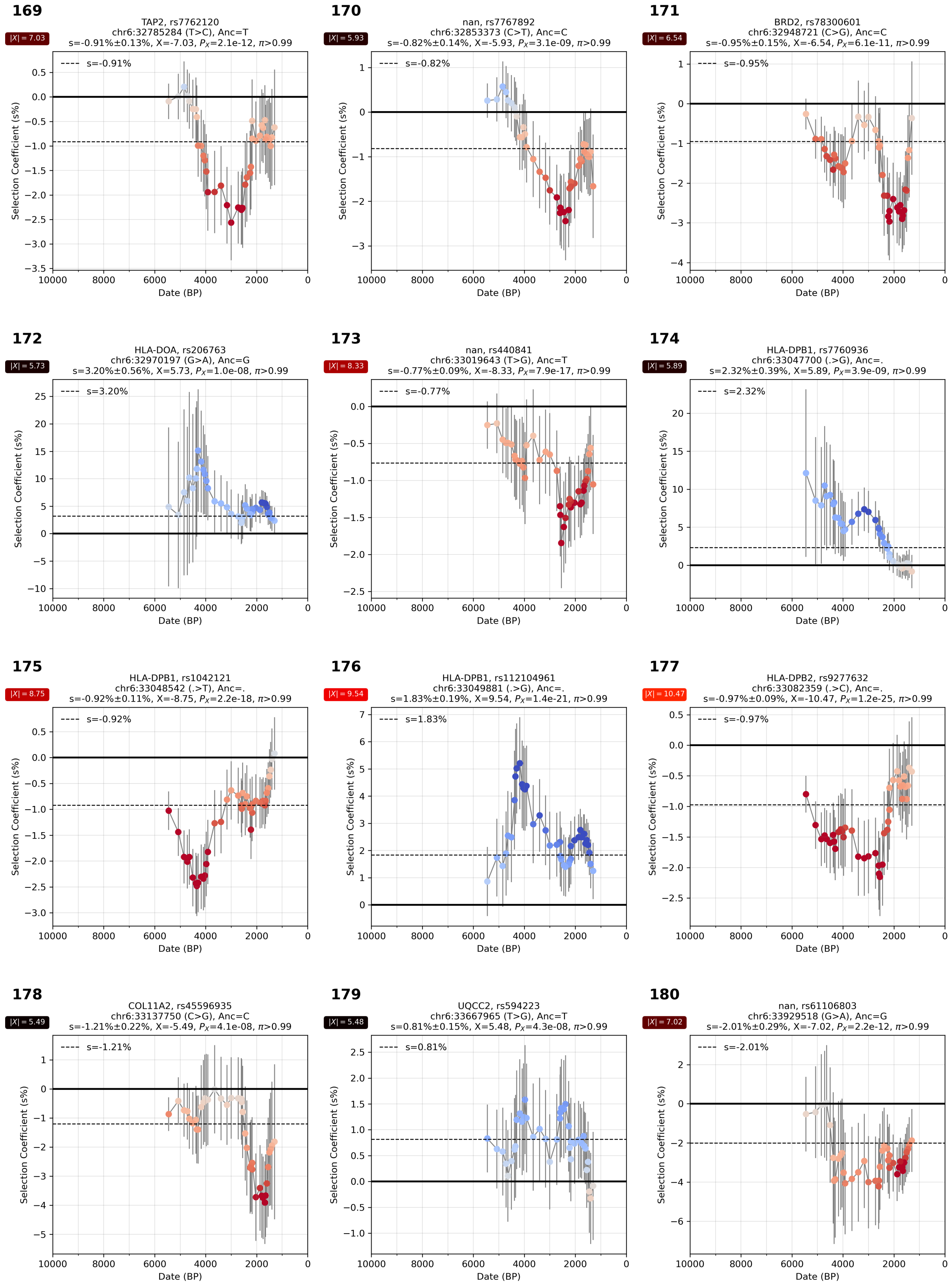

**Supplementary Figure S4.44:** Selection coefficient over time. From chr 6.

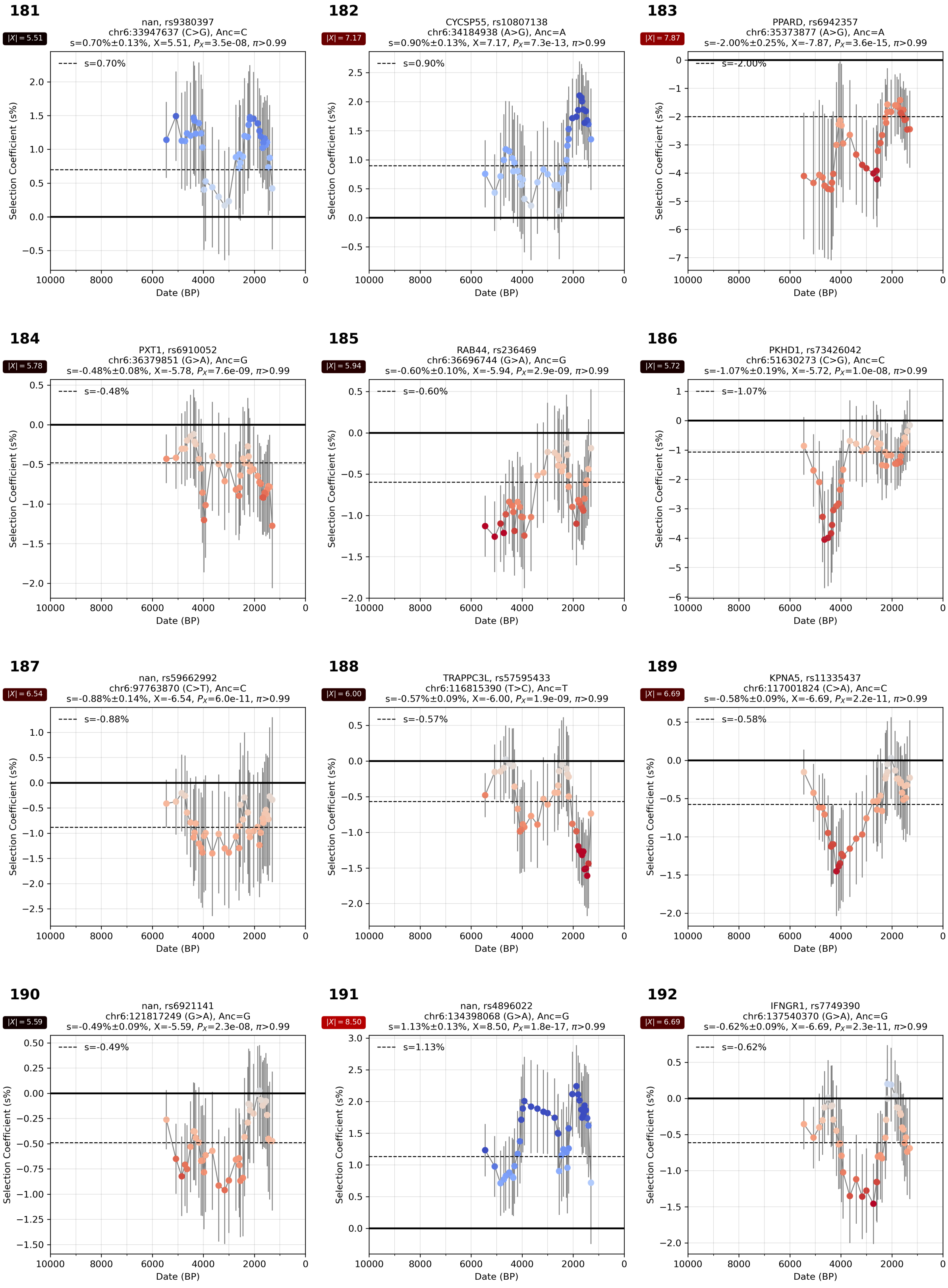

**Supplementary Figure S4.45:** Selection coefficient over time. From chr 6.

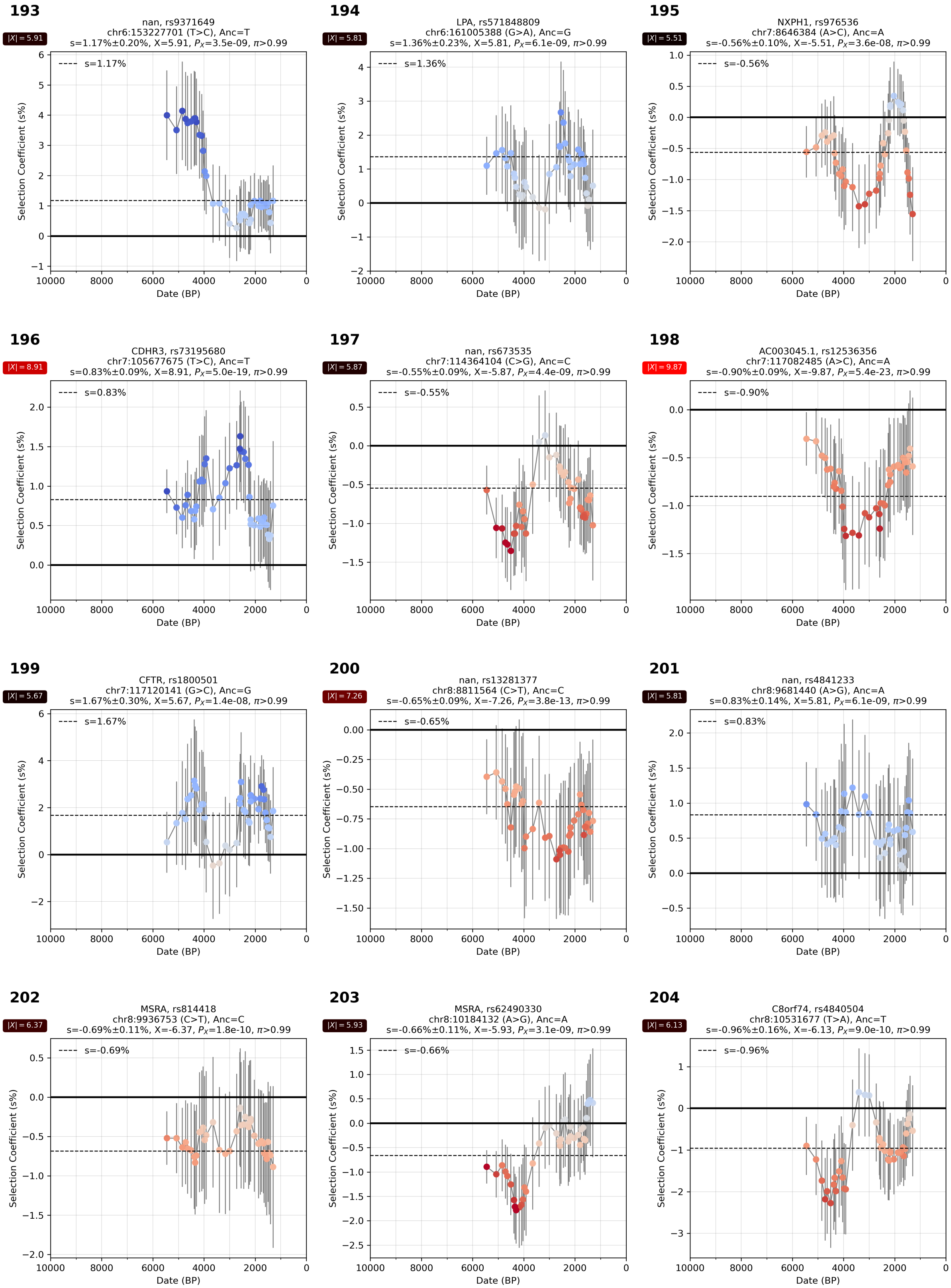

**Supplementary Figure S4.46:** Selection coefficient over time. From chr 6, 7, and 8.

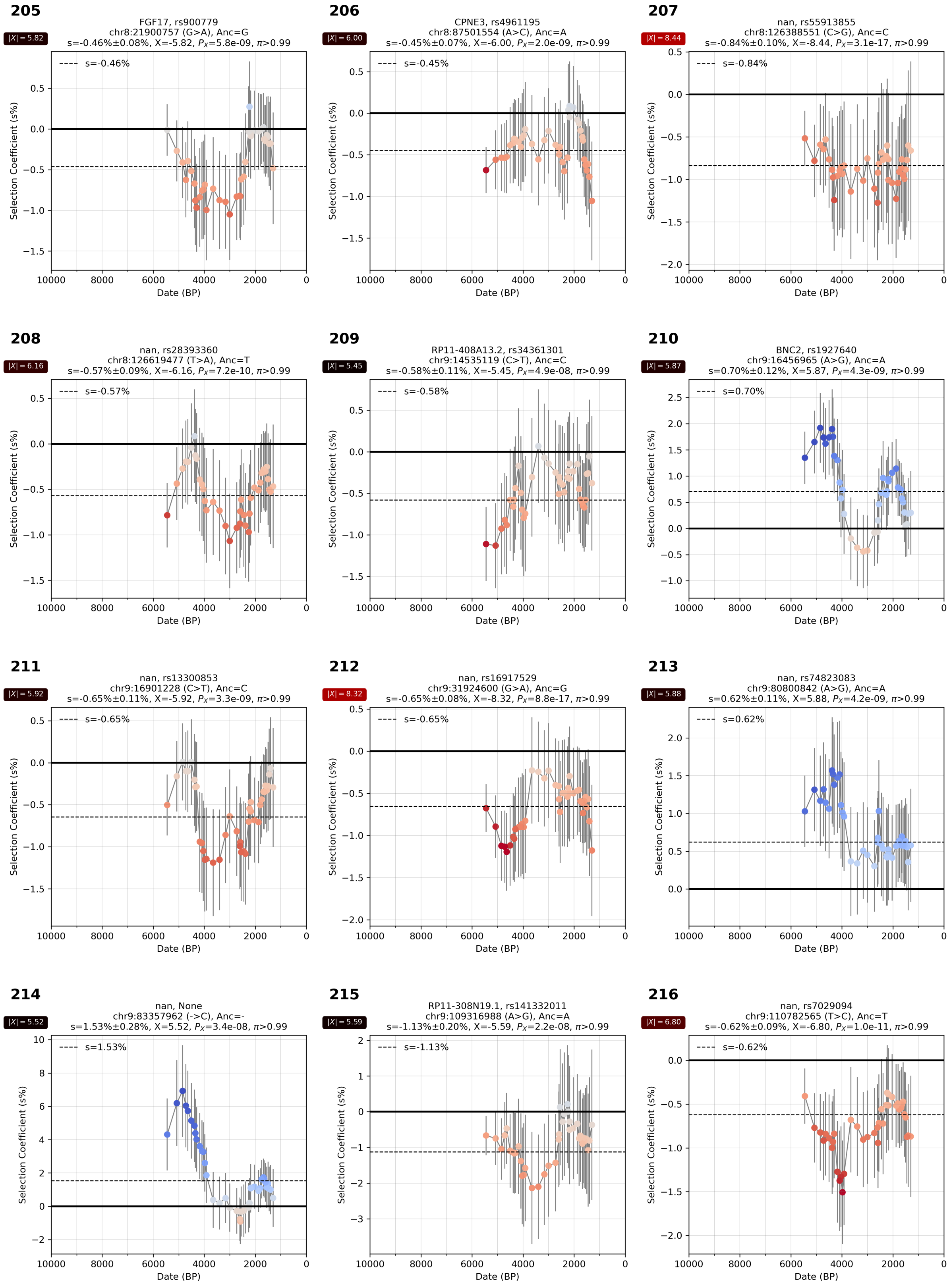

**Supplementary Figure S4.47:** Selection coefficient over time. From chr 8 and 9.

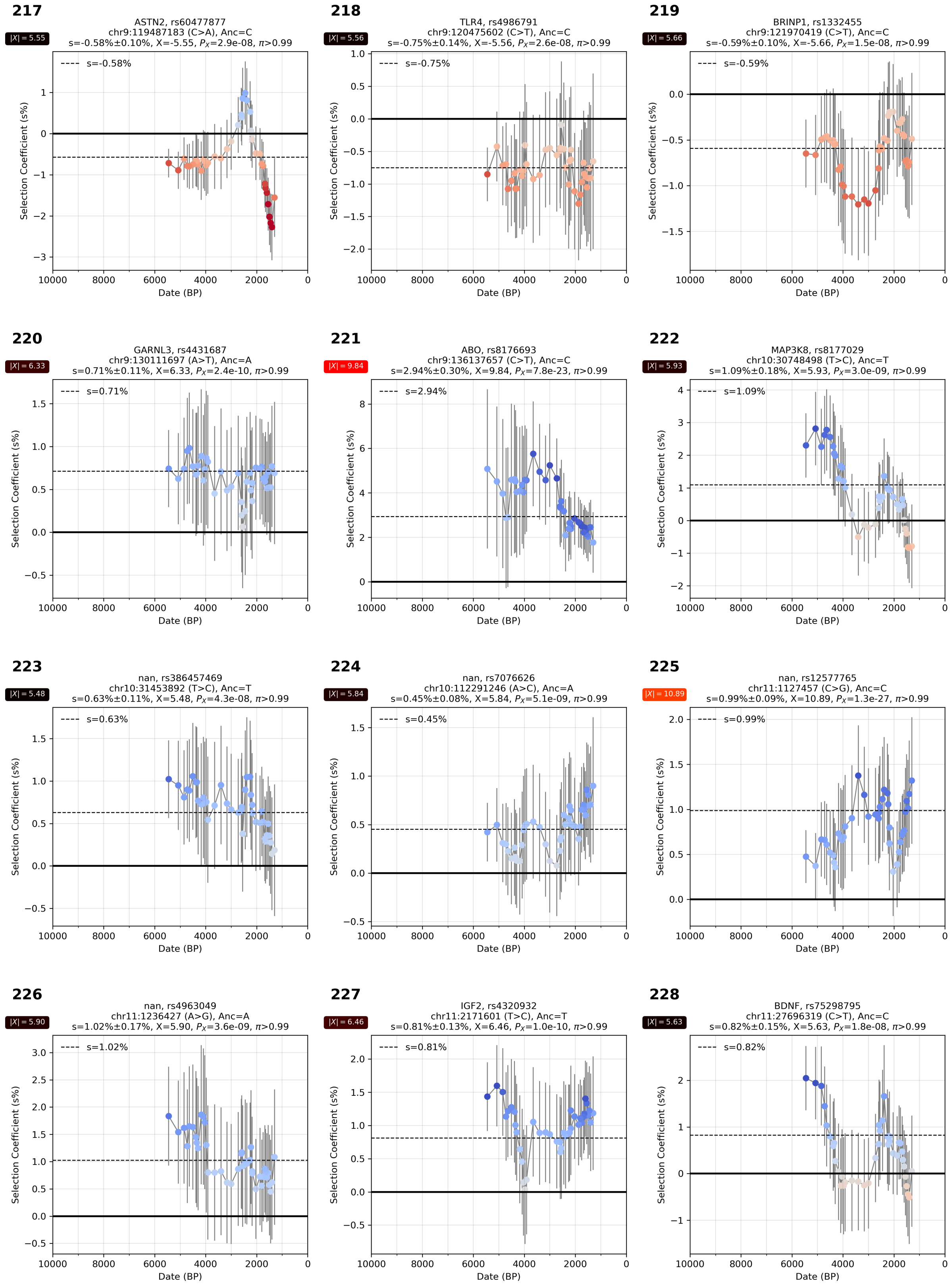

**Supplementary Figure S4.48:** Selection coefficient over time. From chr 9, 10, and 11.

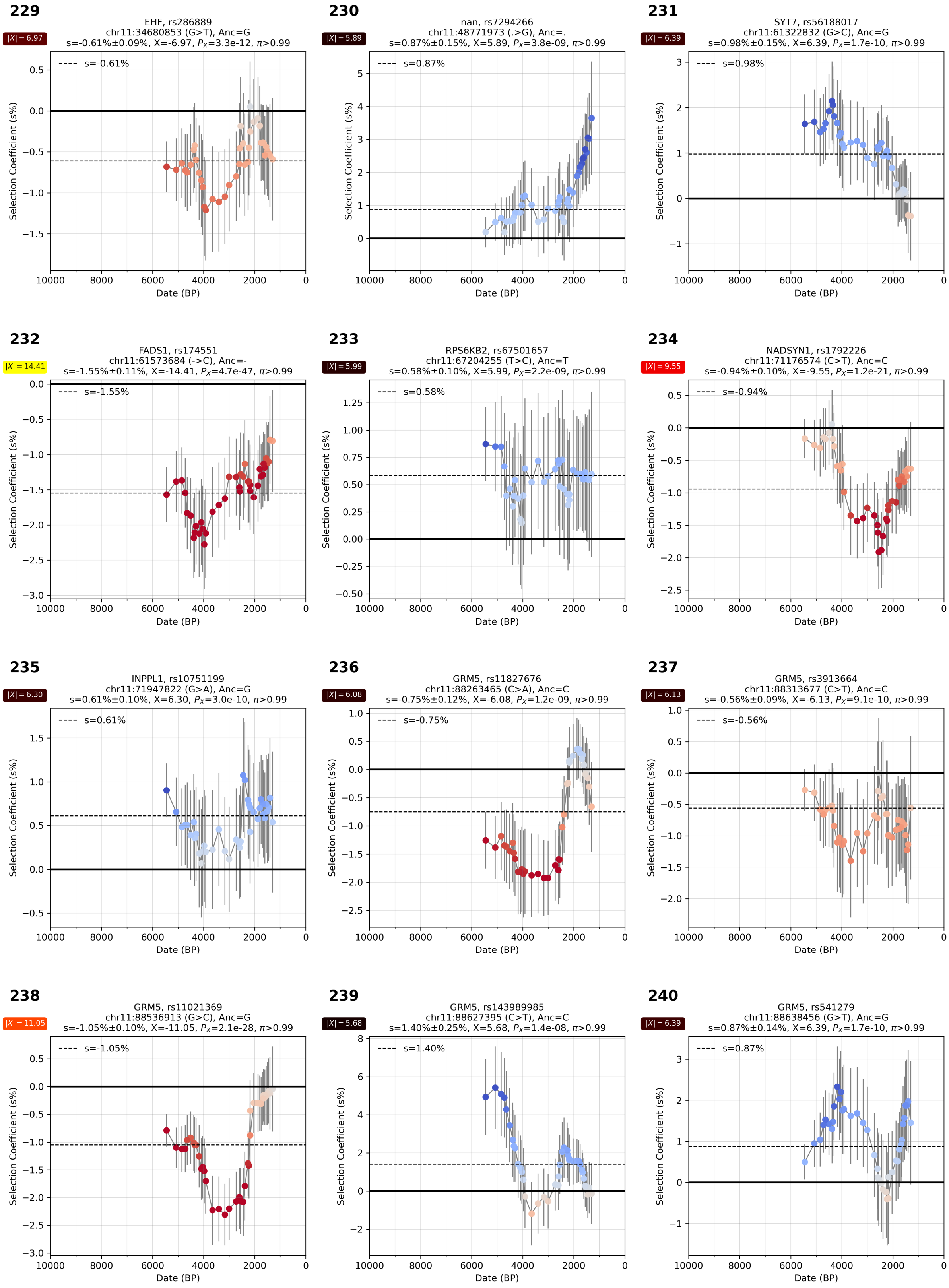

**Supplementary Figure S4.49:** Selection coefficient over time. From chr 11.

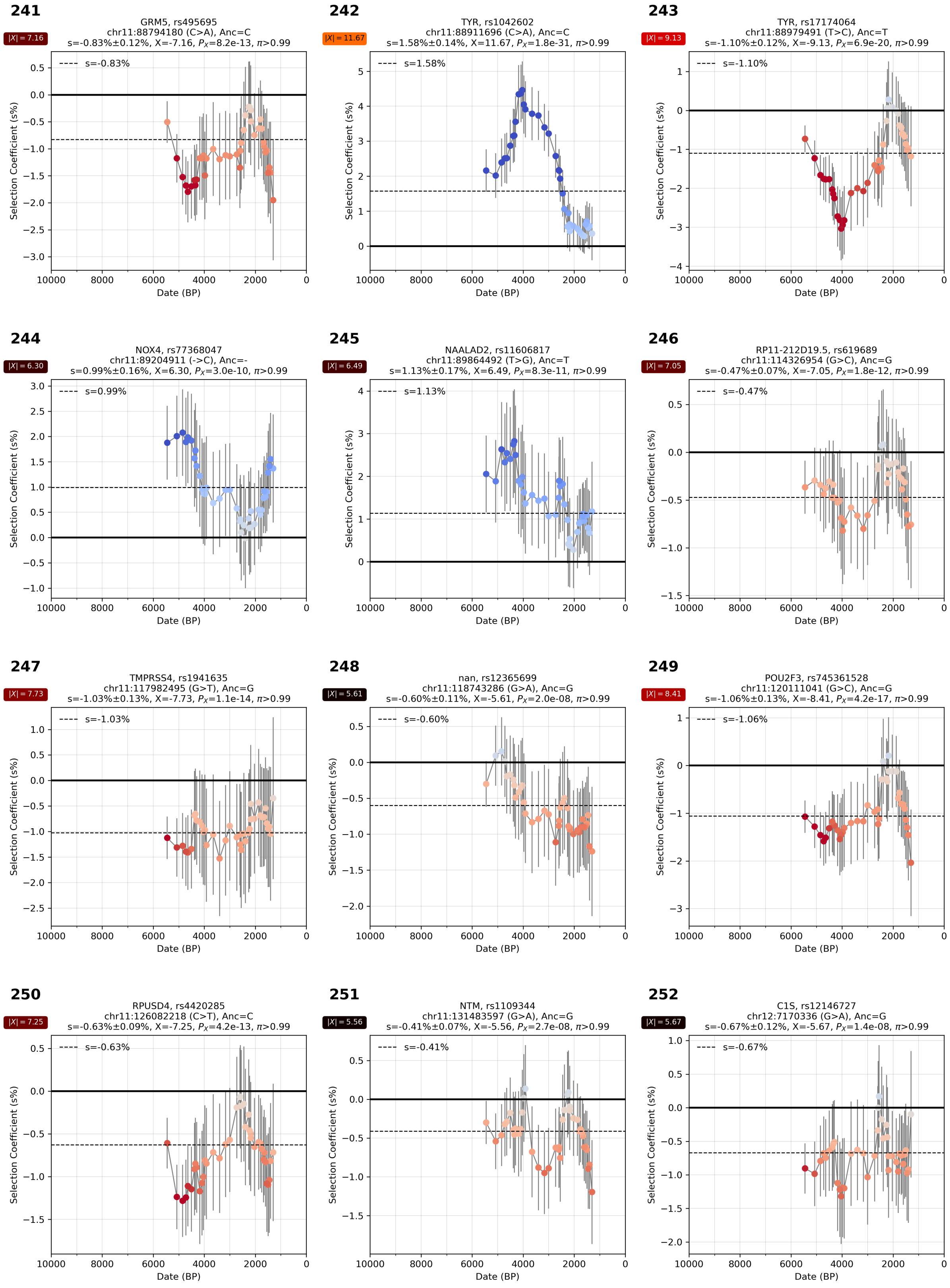

**Supplementary Figure S4.50:** Selection coefficient over time. From chr 11 and 12.

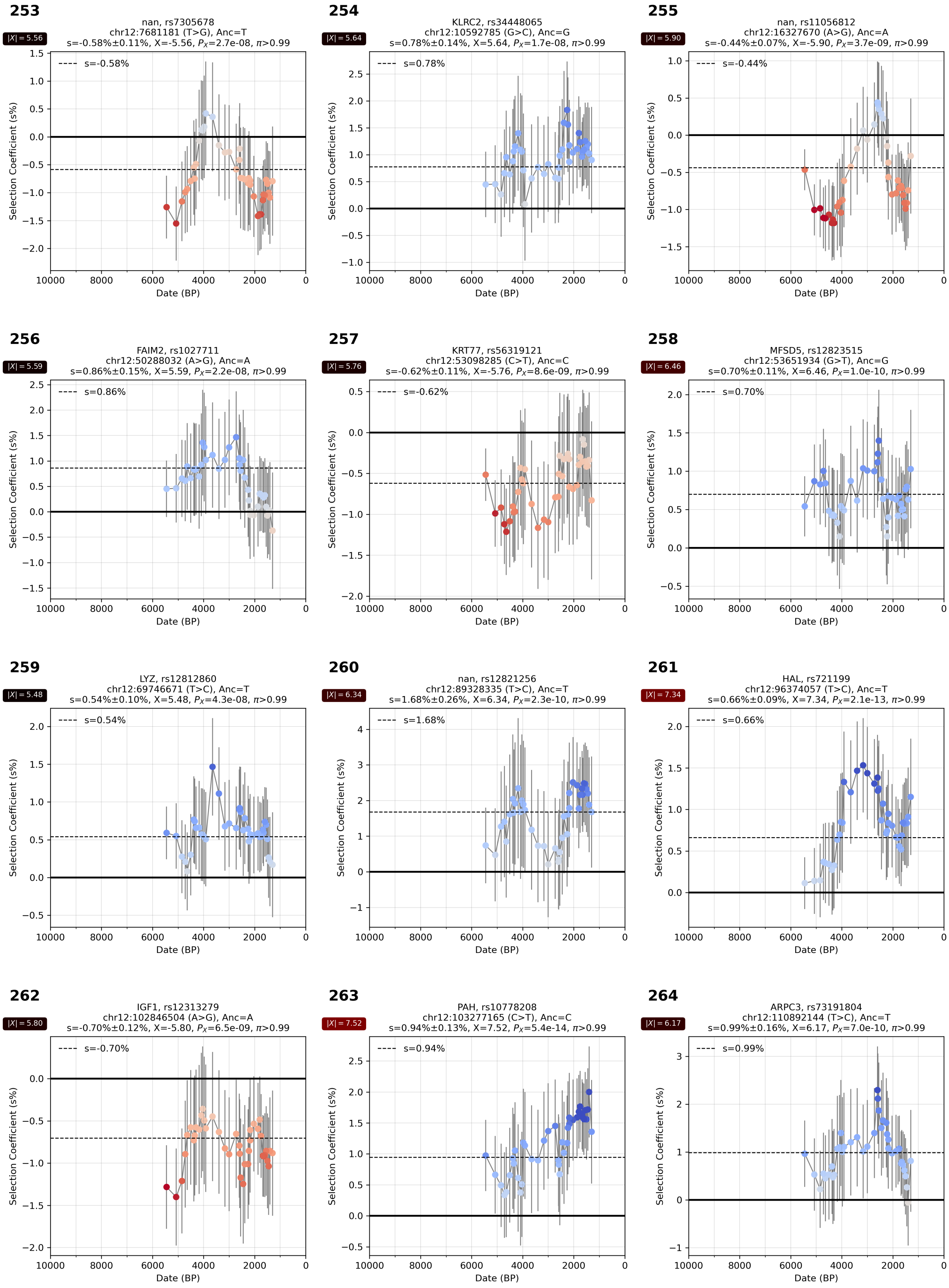

**Supplementary Figure S4.51:** Selection coefficient over time. From chr 12.

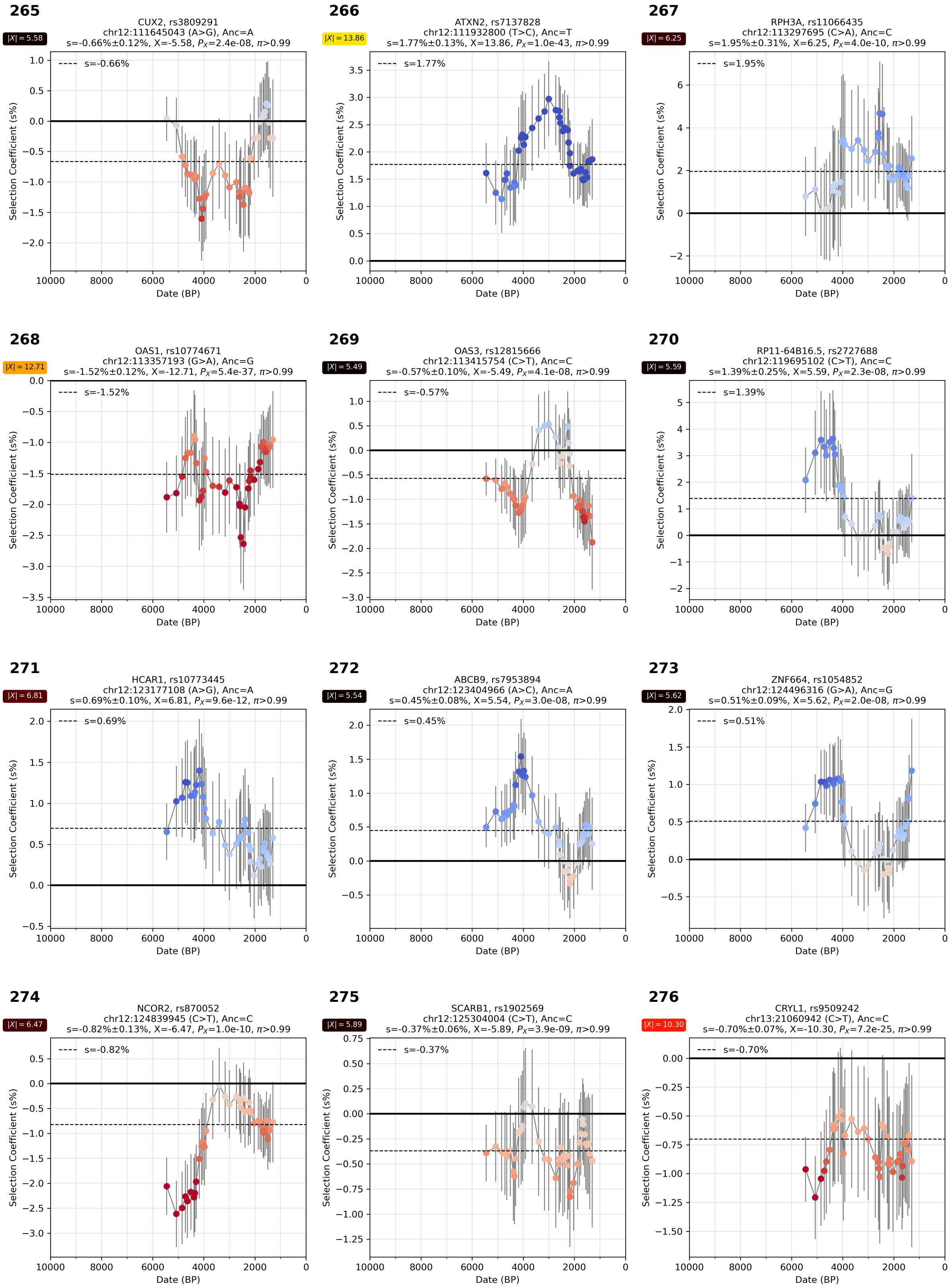

**Supplementary Figure S4.52:** Selection coefficient over time. From chr 12 and 13.

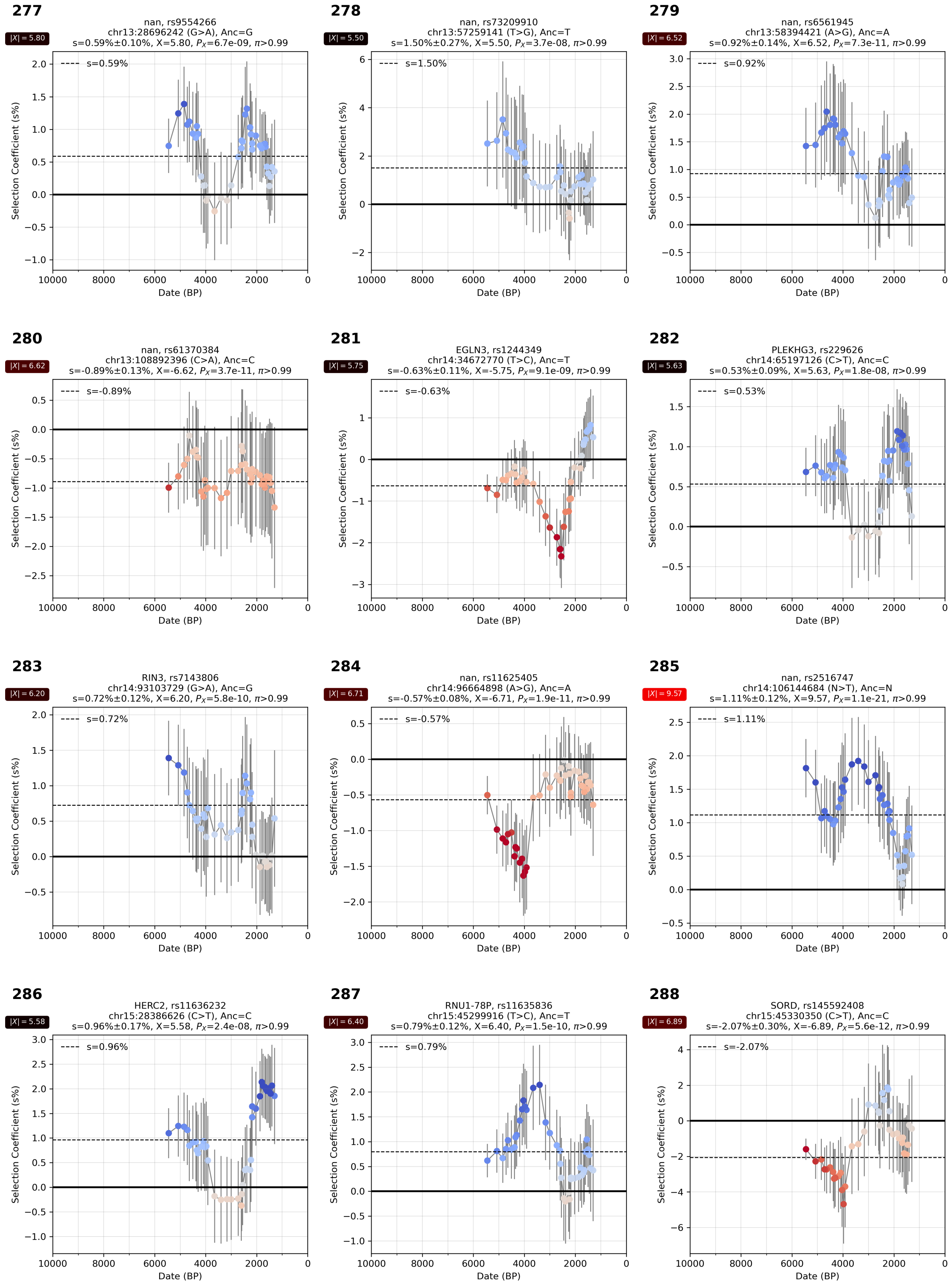

**Supplementary Figure S4.53:** Selection coefficient over time. From chr 13, 14, and 15.

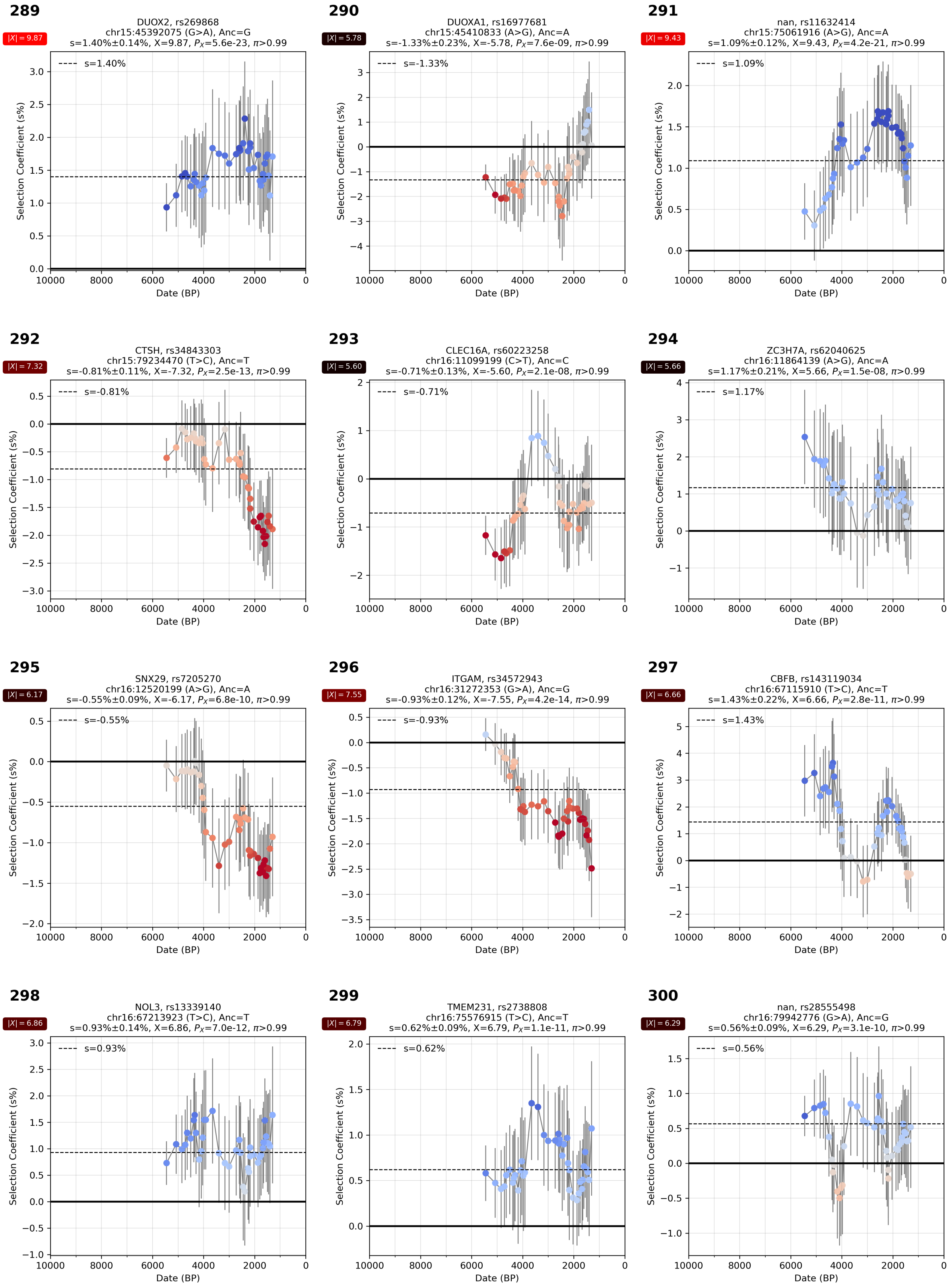

**Supplementary Figure S4.54:** Selection coefficient over time. From chr 15 and 16.

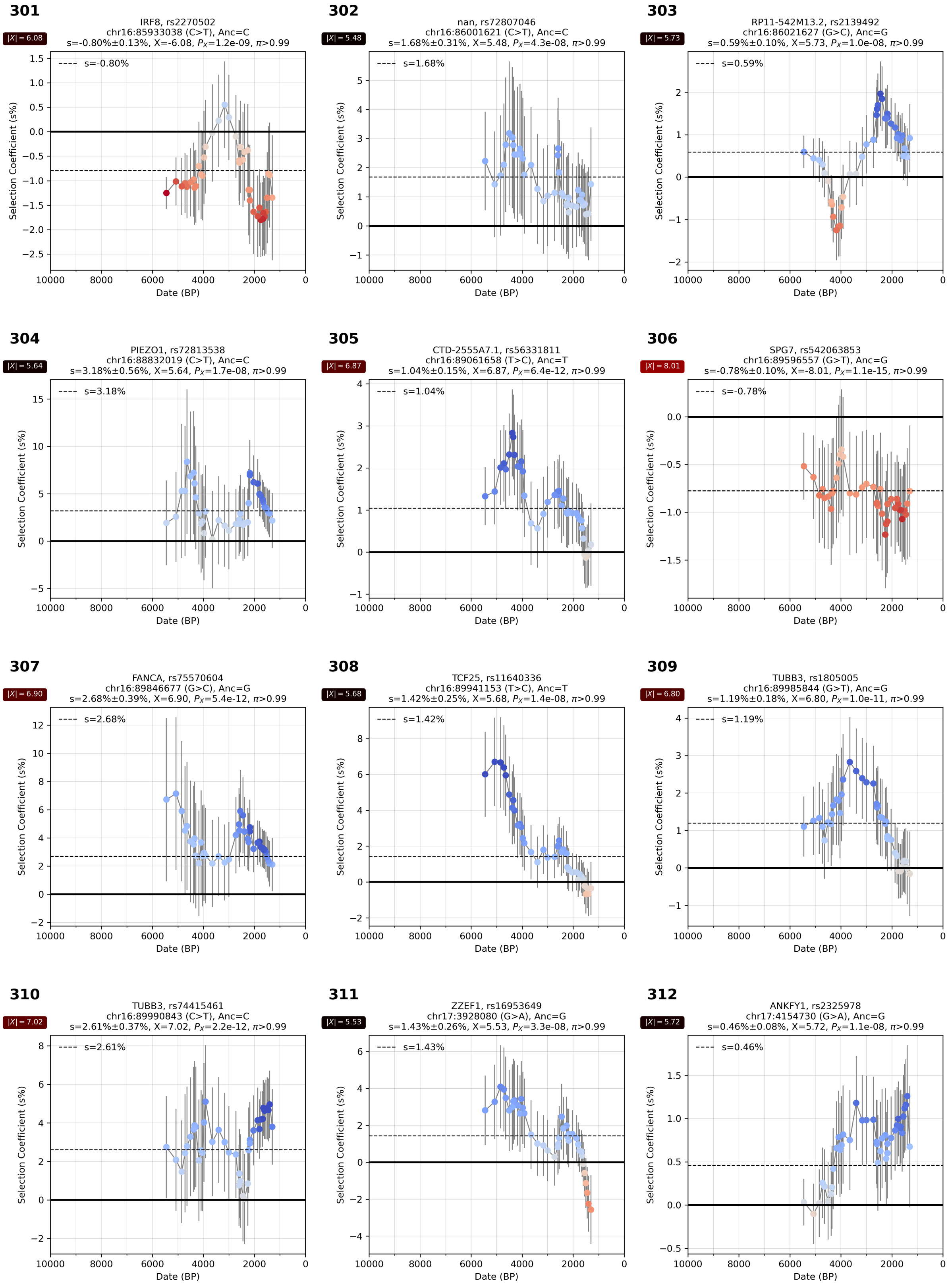

**Supplementary Figure S4.55:** Selection coefficient over time. From chr 16 and 17.

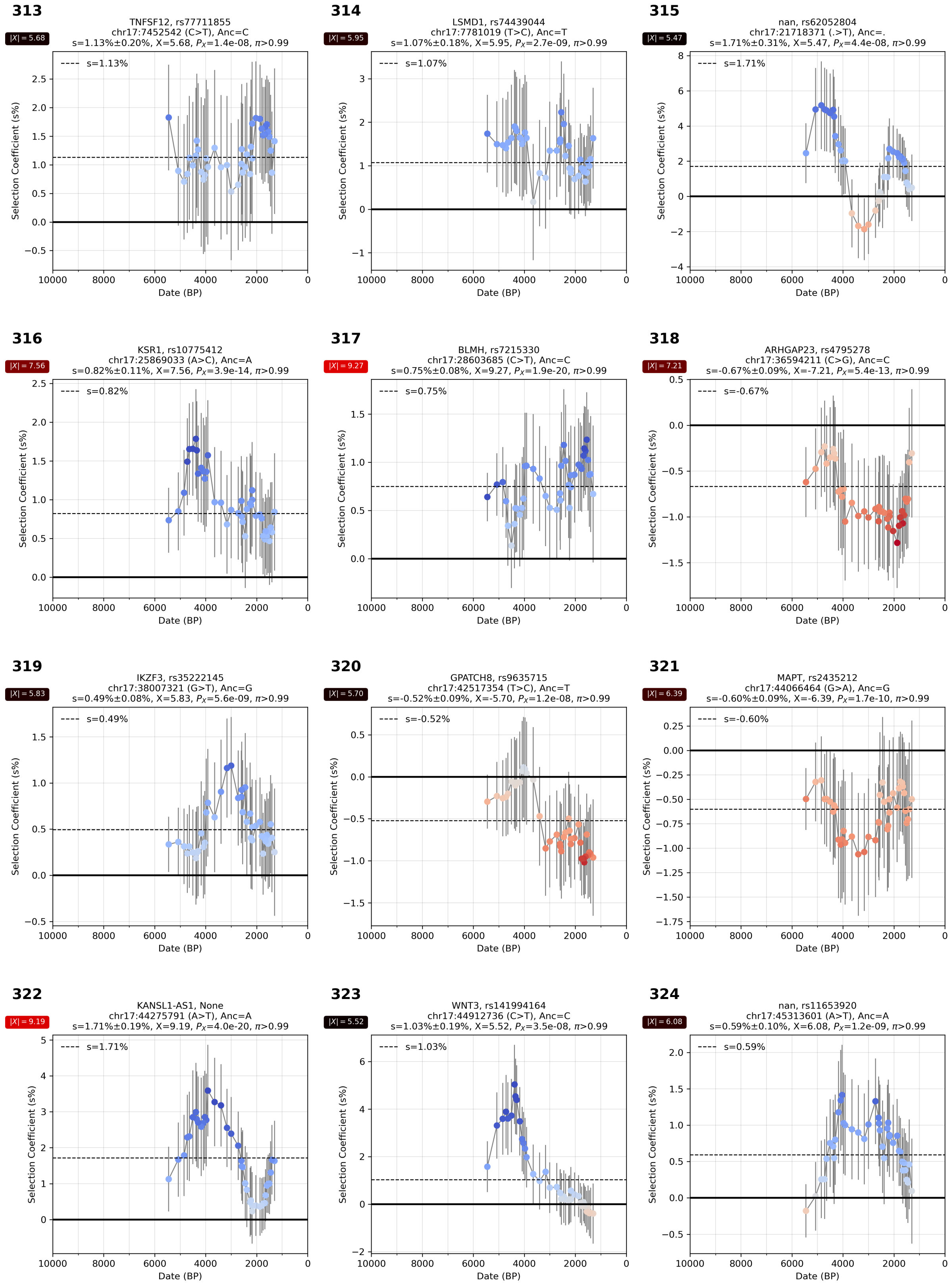

**Supplementary Figure S4.56:** Selection coefficient over time. From chr 17.

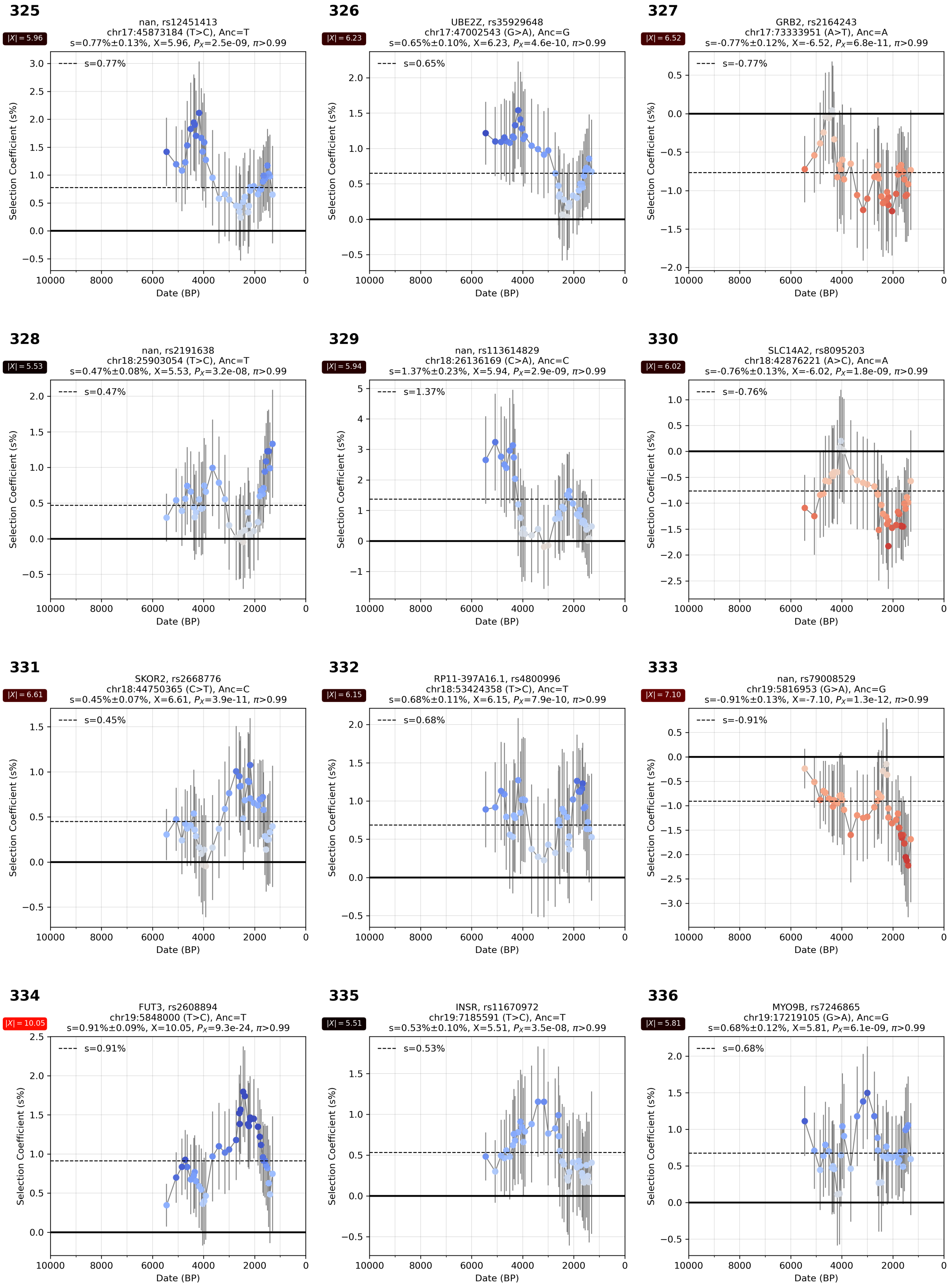

**Supplementary Figure S4.57:** Selection coefficient over time. From chr 17, 18, and 19.

**337**

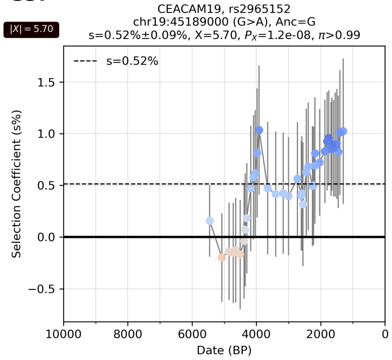

**338**

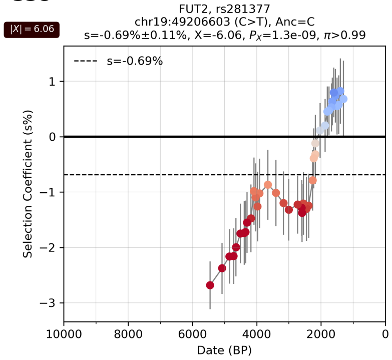

**339**

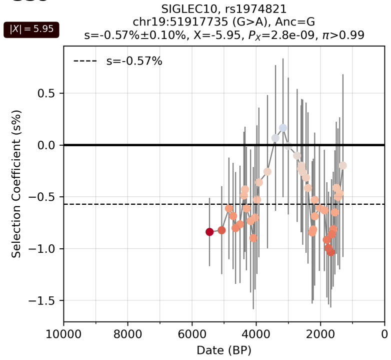

**340**

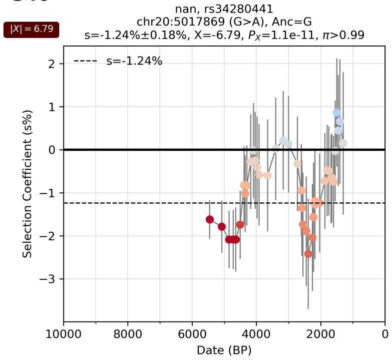

**341**

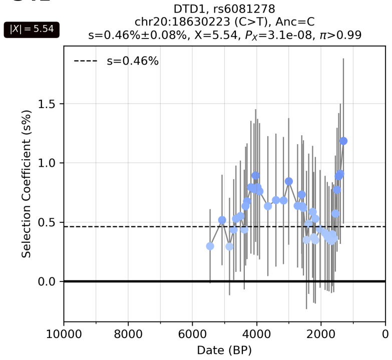

**342**

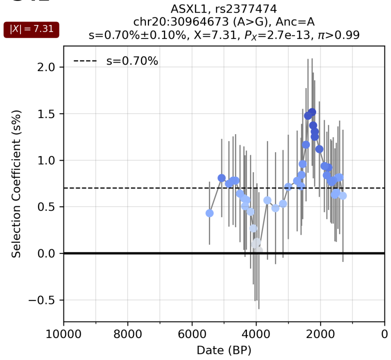

**343**

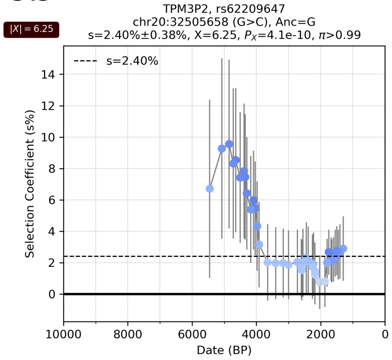

**344**

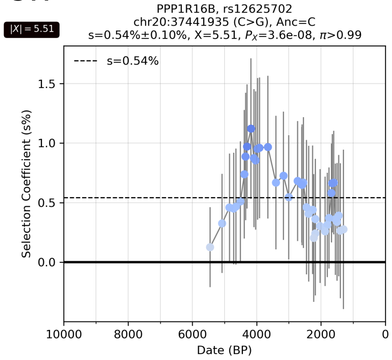

**345**

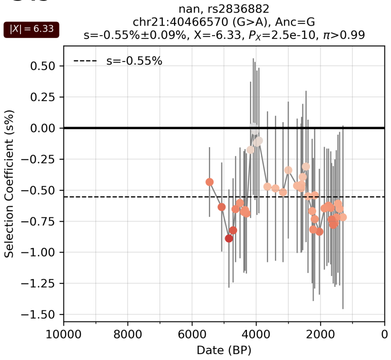

**346**

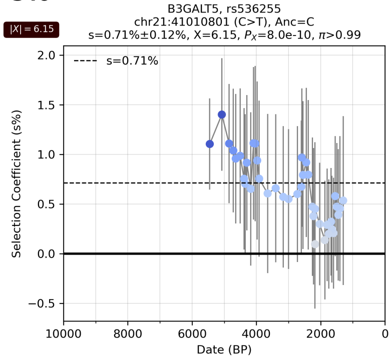

**347**

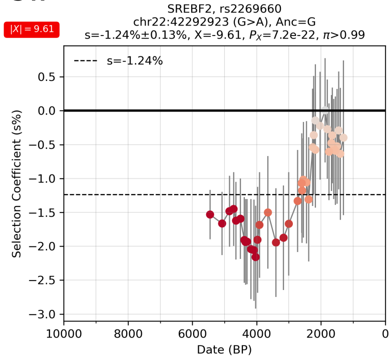

**Supplementary Figure S4.58:** Selection coefficient over time. From chr 19, 20, 21, 22.

### Supplementary Information section 5
### Re-evaluation of results from previous studies

#### Overview

We evaluated candidate loci from five different genome scans for selection: four analyzing ancient DNA time transects (Mathieson et al. 2015[8], Le et al. 2022[9], Kerner et al. 2023[10], and Irving-Pease et al. 2024[11]) and one analyzing modern variation but sensitive to signals we might expect to be replicable in our ancient DNA time transect (Field et al. 2016[12]).

To enable this comparison, we used variants from the high-coverage (30x) 1000 Genomes Project[13] mapped to GRCh38, and remapped to positions in GRCh37/hg19 using CrossMap (v0.5.2)[14]. We used data from 2,504 unrelated individuals from the phase three panel of the 1000 Genomes Project as our imputation reference panel. We retained only the variants that passed all quality control filters from gnomAD (v2.1.1)[15], indicated by a PASS value in the FILTER column of the VCF file available for all chromosome sites on the gnomAD website. This filtering resulted in 52,382,872 biallelic variants. In some cases, the SNPs reported by five studies whose results we re-examined were not included in this reference panel and were excluded from our re-analysis. Additionally, some SNPs did not pass subsequent quality control (QC), as detailed in Supplementary Information section 1. We analyzed all SNPs present in the reference panel. However, the GLMM and allele frequency trajectory results for SNPs that failed QC may be unreliable and could be influenced by the artifacts that caused QC failure.

For each study, we re-evaluated the selection signals using our GLMM approach. The cumulative number of non-HLA signals identified as genome-wide significant in these studies and confirmed in our re-analysis with a posterior probability of $\pi$>99% is 17 (6% of the 279 non-HLA loci showing $\pi$>99% in our genome-wide scan). Of these, 8 were found in Mathieson et al., Field et al. added 0, Le et al. added 3, Kerner et al. added 0, and Irving-Pease et al. added 6. (Table1, Supplementary Table S5.1). An additional 22 non-HLA loci reported as genome-wide significant in at least one of these five studies did not replicate at $\pi$>99% in our re-analysis (Table 1).

Mathieson et al. 2015 analyzed whole genome data from 230 ancient Eurasian individuals who lived between 6500 BCE and 300 BCE and compared it to data from modern Europeans from the 1000 Genomes Project. They found 12 genome-wide signals of selection with a significance exceeding a P-value threshold of 5e-8, two of which are from the HLA region. Of the 11 that pass our QC, we replicate 10 ($\pi$>99%) (the 11$^{th}$ is replicated at $\pi$>69%, and is almost certainly a real signal of selection as it is the blue eye-color variant at *OCA2*/*HERC2* which has been found to be subject to fluctuating selection which our methodology is not optimized to detect as we are explicitly testing for a scenario of a constant non-zero selection coefficient).

Field et al. 2016 analyzed 3195 contemporary individuals from the UK10K project to study signals of selection in the past 2000 years inferred from an unusually high density of singleton genetic variants associated with a tested allele, which can be evidence of a distortion of the gene as a result of selection. We applied our GLMM approach to three time transects: to all individuals in our study, to individuals who lived before 2000 BP, and to individuals who lived after 2000 BP which is the time frame where Field et al. 2016 have particularly notable statistical power. There are 3 genome-wide signals of selection in Field et al. 2016 exceeding a P value threshold of 5e-8, and two of them (*LCT* and *HLA*) are replicated in our analysis ($\pi$>99%). There is no evidence for selection during the past 2000 years for the third candidate at the *WDFY4* locus using our analysis, and we hypothesize that this signal may be an artifact of incompletely corrected population structure due to ancestry derived from steppe pastoralists, an issue known to have caused Field et al. 2016 to find false-positive signals of polygenic selection[16,17]. Using a less stringent threshold, out of 37 independent loci highlighted in Field et al. 2016 as passing the significance threshold of 1e-5, 35 pass our QC, and only three (*LCT*, HLA, *KITLG*) show a strong signal of selection.

Le et al. 2022 analyzed 1291 European individuals from the past 10,000 years to identify selection signals in the Neolithic, Bronze Age, and Historical periods. They found 25 selection signals across 24 loci, of which 22 loci could be retested in our analysis (two were not present in the imputation reference panel), and we replicated 9 in our full time transect ($\pi$>99%). We also separately analyzed each of the three epochs to mimic the approach in Le et al. 2022, and only replicated 7 of 23 signals for which the associated SNPs passed QC. In the Neolithic we replicated 1 out of 10. In the Bronze Age we replicated 1 out of 7 (we could not retest two SNPs that were not present in the imputation reference panel). In the Historical period we replicated 5 out of 6.

Kerner et al. 2023 analyzed 2879 Eurasian individuals from the past 10,000 years to screen for signals of selection. They highlighted a list of 139 SNPs, with 89 potentially showing evidence of positive selection and 50 showing evidence of negative selection on the derived allele. As the authors point out in the transparent peer review records published alongside their study (Supplementary Document S2 of that study), only 3 of these 139 signals of selection (*LCT*/*MCM6*, HLA, and *SLC45A2*) pass a formal genome-wide significant threshold with a P-value significance threshold of 5e-8; all three are replicated in our analysis. We believe that the great majority of the 139 highlighted selection signals are false positives due to not applying threshold for genome-wide statistical significance: out of 123 candidate SNPs that pass QC, only 14 are genome-wide significant in our re-analysis ($\pi$>99%); an additional 10 loci had probable evidence of selection in our re-analysis (50%<$\pi$<99%). When we visually inspect the SNPs where there is a failure to replicate, the great majority show no visual evidence for selection in our time-transect. In particular, shifts in frequency at these non-replicated variants often appear to be in the direction expected based on pre-existing allele frequency differences between source populations like steppe pastoralists, European farmers, and European hunter-gatherers (which would be expected to produce a substantial allele frequency shift following mixture of these group if not corrected).

Irving-Pease et al. 2024 analyzed 1518 West Eurasians over the past 15,000 years and used the CLUES methodology[18,19] to identify 21 genome-wide significant loci with at least one genome-wide significant signal (P-value threshold of 5e-8) across five ancestry categories: pan-ancestry (ALL), Western hunter-gatherers (WHG), Eastern hunter-gatherers (EHG), Caucasus hunter-gatherers (CHG), and Anatolian farmers (ANA). Of the 21 candidate loci identified, our analysis found that 13 had at least one SNP among five candidates per locus listed by Irving-Pease et al. 2024 with genome-wide significant posterior probability ($\pi$>99%); an additional 4 loci had probable evidence of selection in our re-analysis (50%<$\pi$<99%). Our analysis increases the chance that some or all of the remaining 8 may be false-positives.

**Supplementary Table S5.1:** Summary of genome-wide significant signals of selection across five studies. SNPs within 100 kb of each other are considered a single locus, except for HLA, which is treated as one locus regardless of distance for this reanalysis. For each study, a value of 1 indicates that the locus is reported as genome-wide significant, and 0 otherwise. The Panel ID corresponds to the identifier used in the supplementary figures below to mark each panel, starting with the first letter of the corresponding study. (In Mathieson et al. 2015, there are two signals from the HLA region, while Le et al. 2022 excluded the HLA region from their Table 1, despite its evidence of significant selection across all epochs.)

| Variant ID | RSID | Pane ID | Gene | Locus | Mathieson et al. 2015 | Field et al. 2016 | Le et al. 2022 | Kerner et al. 2023 | Irving-Pease et al. 2024 | $\pi$ |
|---|---|---|---|---|---|---|---|---|---|---|
| 1_22704191_T_G | rs11799474 | I1 | | 1 | 0 | 0 | 0 | 0 | 1 | 0.28 |
| 1_150596411_C_A | rs7517 | L3 | ENSA | 2 | 0 | 0 | 1 | 0 | 0 | 0.38 |
| 1_181018799_A_C | rs10797666 | L16 | MR1 | 3 | 0 | 0 | 1 | 0 | 0 | 0.03 |
| 1_202143512_A_G | rs12401678 | L4 | PTPRVP | 4 | 0 | 0 | 1 | 0 | 0 | 0.37 |
| 1_230854999_C_T | rs7555650 | L14 | AGT | 5 | 0 | 0 | 1 | 0 | 0 | 0.21 |
| 2_100604753_G_C | rs56127672 | I7 | AFF3 | 6 | 0 | 0 | 0 | 0 | 1 | 0.65 |
| 2_102824201_G_A | rs2310239 | L2 | IL1RL2 | 7 | 0 | 0 | 1 | 0 | 0 | >0.99 |
| 2_136608646_G_A | rs4988235 | M1 | MCM6 LCT | 8 | 1 | 0 | 0 | 0 | 0 | >0.99 |
| 2_136608646_G_A | rs4988235 | L18 | | | 0 | 0 | 1 | 0 | 0 | >0.99 |
| 2_136608646_G_A | rs4988235 | K1 | | | 0 | 0 | 0 | 1 | 0 | >0.99 |
| 2_136608646_G_A | rs4988235 | I11 | | | 0 | 0 | 0 | 0 | 1 | >0.99 |
| 2_136707982_T_C | rs6754311 | F1 | | | 0 | 1 | 0 | 0 | 0 | >0.99 |
| 3_46954614_A_C | rs201652298 | I16 | | 9 | 0 | 0 | 0 | 0 | 1 | 0.95 |
| 4_38745482_T_C | rs10008032 | I21 | TLR10 TLR1 | 10 | 0 | 0 | 0 | 0 | 1 | >0.99 |
| 4_38776107_T_G | rs11096955 | L22 | | | 0 | 0 | 1 | 0 | 0 | >0.99 |
| 4_38815502_A_C | rs4833103 | M7 | | | 1 | 0 | 0 | 0 | 0 | >0.99 |
| 4_145506871_C_T | rs6537298 | I29 | | 11 | 0 | 0 | 0 | 0 | 1 | 0.4 |
| 5_33951693_C_G | rs16891982 | M5 | SLC45A2 | 12 | 1 | 0 | 0 | 0 | 0 | >0.99 |
| 5_33951693_C_G | rs16891982 | L11 | | | 0 | 0 | 1 | 0 | 0 | >0.99 |
| 5_33952106_C_T | rs185146 | K3 | | | 0 | 0 | 0 | 1 | 0 | >0.99 |
| 5_33954880_G_A | rs35389 | I33 | | | 0 | 0 | 0 | 0 | 1 | >0.99 |
| 5_131675864_A_G | rs272872 | M4 | SLC22A4 | 13 | 1 | 0 | 0 | 0 | 0 | >0.99 |
| 5_131705458_C_G | rs2631367 | I39 | SLC22A5 | | 0 | 0 | 0 | 0 | 1 | >0.99 |
| 5_176842474_T_C | rs2731672 | I45 | GRK6 | 14 | 0 | 0 | 0 | 0 | 1 | >0.99 |
| 6_28322296_A_G | rs6903823 | M8 | HLA* | 15 | 1 | 0 | 0 | 0 | 0 | >0.99 |
| 6_28819880_A_G | rs9257267 | F2 | | | 0 | 1 | 0 | 0 | 0 | >0.99 |
| 6_30746519_G_T | rs3130673 | K4 | | | 0 | 0 | 0 | 1 | 0 | >0.99 |
| 6_31142245_C_A | rs3094188 | I50 | | | 0 | 0 | 0 | 0 | 1 | >0.99 |
| 6_32132233_G_A | rs2269424 | M10 | | | 1 | 0 | 0 | 0 | 0 | >0.99 |
| 8_9604066_T_G | rs35169606 | I54 | TNKS | 16 | 0 | 0 | 0 | 0 | 1 | >0.99 |
| 8_130981907_C_A | rs10956504 | L7 | FAM49B | 17 | 0 | 0 | 1 | 0 | 0 | 0.07 |

| | | | | | | | | | | |
|---|---|---|---|---|---|---|---|---|---|---|
| 8_132669706_A_G | rs1420488 | I60 | | 18 | 0 | 0 | 0 | 0 | 1 | 0.22 |
| 9_136137657_C_T | rs8176693 | I63 | ABO | 19 | 0 | 0 | 0 | 0 | 1 | >0.99 |
| 10_49924070_T_C | rs76203261 | F3 | WDFY4 | 20 | 0 | 1 | 0 | 0 | 0 | 0.36 |
| 11_18167630_A_G | rs4256954 | L17 | | 21 | 0 | 0 | 1 | 0 | 0 | <0.01 |
| 11_27679916_C_T | rs6265 | L9 | BDNF | 22 | 0 | 0 | 1 | 0 | 0 | <0.01 |
| 11_61569830_C_T | rs174546 | M2 | | | 1 | 0 | 0 | 0 | 0 | >0.99 |
| 11_61569830_C_T | rs174546 | I69 | FADS1 | 23 | 0 | 0 | 0 | 0 | 1 | >0.99 |
| 11_61571478_T_C | rs174550 | L19 | | | 0 | 0 | 1 | 0 | 0 | >0.99 |
| 11_71153459_C_A | rs11603330 | L21 | DHCR7 | 24 | 0 | 0 | 1 | 0 | 0 | >0.99 |
| 11_71165625_A_G | rs7944926 | M9 | NADSYN1 | | 1 | 0 | 0 | 0 | 0 | >0.99 |
| 11_88515022_A_G | rs7119749 | M6 | GRM5 | 25 | 1 | 0 | 0 | 0 | 0 | >0.99 |
| 12_112007756_C_T | rs653178 | M3 | | | 1 | 0 | 0 | 0 | 0 | >0.99 |
| 12_112007756_C_T | rs653178 | L20 | ATXN2 | 26 | 0 | 0 | 1 | 0 | 0 | >0.99 |
| 12_112007756_C_T | rs653178 | I74 | | | 0 | 0 | 0 | 0 | 1 | >0.99 |
| 14_103867320_A_G | rs4906319 | L13 | MARK3 | 27 | 0 | 0 | 1 | 0 | 0 | 0.51 |
| 15_28365618_A_G | rs12913832 | M11 | HERC2 | 28 | 1 | 0 | 0 | 0 | 0 | 0.69 |
| 15_28386626_C_T | rs11636232 | L12 | | | 0 | 0 | 1 | 0 | 0 | >0.99 |
| 15_75077367_C_A | rs1378942 | I80 | CSK | 29 | 0 | 0 | 0 | 0 | 1 | >0.99 |
| 16_54231250_A_G | rs2010410 | L8 | | 30 | 0 | 0 | 1 | 0 | 0 | 0.02 |
| 16_69969299_C_G | rs62053262 | I82 | WWP2 | 31 | 0 | 0 | 0 | 0 | 1 | 0.72 |
| 16_80036594_G_T | rs4073089 | L5 | | 32 | 0 | 0 | 1 | 0 | 0 | 0.09 |
| 17_37355093_C_T | rs57944517 | L6 | RPL19 | 33 | 0 | 0 | 1 | 0 | 0 | 0.09 |
| 17_38150946_C_T | rs4795413 | I87 | PSMD3 | 34 | 0 | 0 | 0 | 0 | 1 | 0.76 |
| 17_44086267_A_G | rs4792897 | I93 | MAPT | 35 | 0 | 0 | 0 | 0 | 1 | >0.99 |
| 19_49206603_C_T | rs281377 | L1 | FUT2 | 36 | 0 | 0 | 1 | 0 | 0 | >0.99 |
| 19_57499627_G_A | rs62132568 | I99 | | 37 | 0 | 0 | 0 | 0 | 1 | 0.02 |
| 21_43679554_C_T | rs915843 | L15 | ABCG1 | 38 | 0 | 0 | 1 | 0 | 0 | 0.11 |
| 22_22027348_C_T | rs1669125 | L10 | PPIL2 | 39 | 0 | 0 | 1 | 0 | 0 | <0.01 |
| 22_42340844_T_C | rs1023500 | I104 | CENPM | 40 | 0 | 0 | 0 | 0 | 1 | >0.99 |

#### Re-evaluation of results from Mathieson et al. 2015

We evaluated 12 SNPs identified as candidates for selection from Extended Data Table 3 of Mathieson et al. 2015[8]. In our re-evaluation of these 12 SNPs, 10 SNPs (rs4988235, rs16891982, rs2269424, rs174546, rs4833103, rs653178, rs7944926, rs7119749, rs272872, rs6903823) showed a compelling signal of selection with posterior probability $\pi$>99% in our analysis. One SNP, rs12913832 at the *OCA2*/*HERC2* locus, showed a signal of selection with posterior probability $\pi$=69%, and is likely a real signal of selection as this is the blue eye color variant which other work has shown has been subject to fluctuating selection over space and time, a scenario not tested for in our methodology which explicitly assumes a constant selection coefficient over space and time. The last SNP, rs1979866, did not pass quality control in our analysis but had a posterior probability $\pi$<1%, so we believe it may be a false-positive due to data artifact (Supplementary Figure S5.1). Two SNPs, rs6903823 and rs2269424, are from the HLA region.

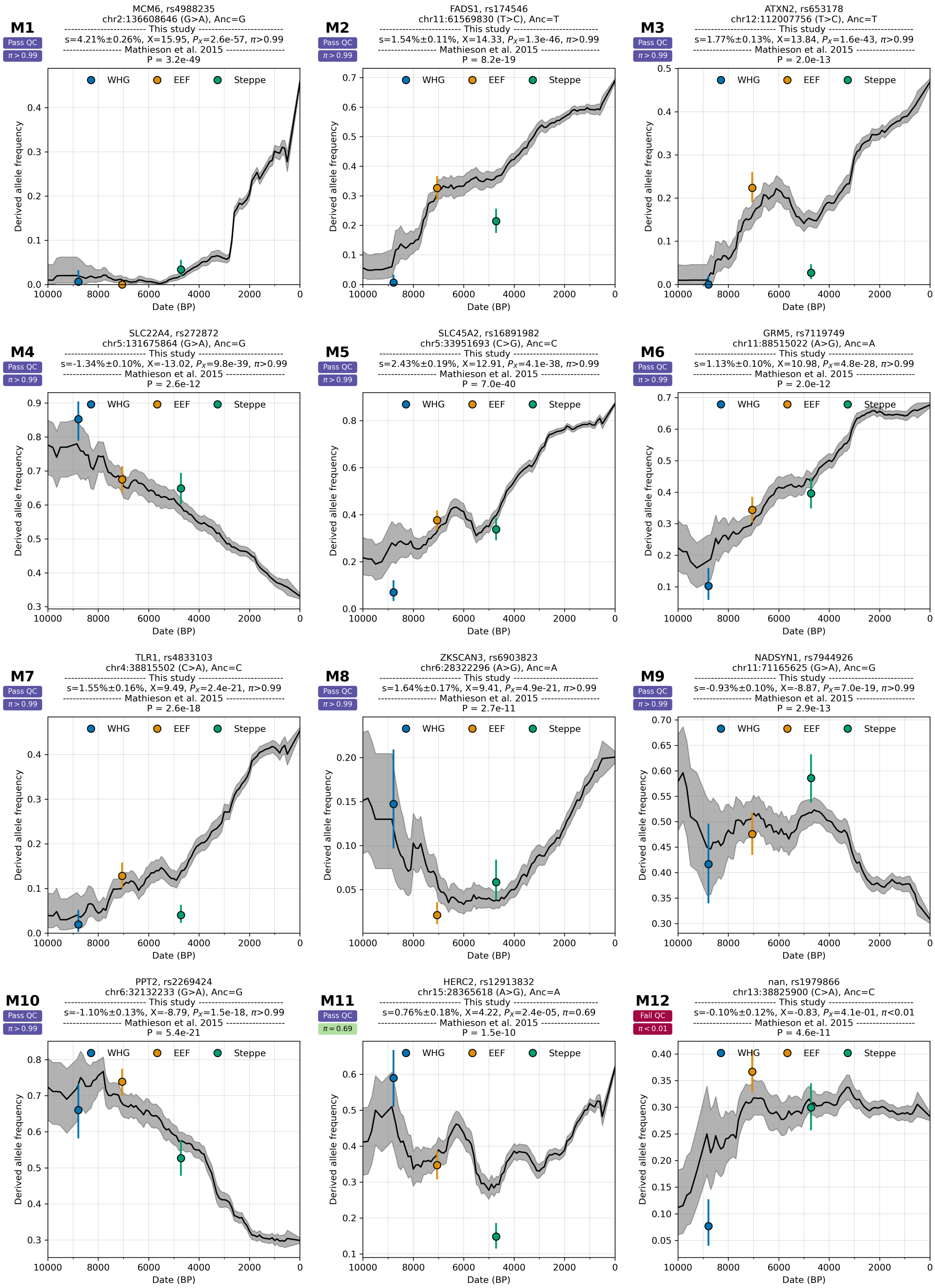

**Supplementary Figure S5.1:** Re-evaluating 12 signals of selection from Mathieson et al. 2015.

### Re-evaluation of results from Field et al. 2016

The singleton density score (SDS) from Field et al. 2016[12] was devised to capture signals of recent selection with particular statistical power during the past 2000 years. This method does not use ancient DNA and is entirely based on patterns of variation in contemporary populations. Here, we evaluated 37 independent tagging SNPs with a p-value of SDS ($P_{SDS}$) less than 1e-5. We used the PLINK clumping option, prioritizing SNPs by $P_{SDS}$ and using clump_kb = 10 Mbp and clump_r2 = 0.05. SNPs that do not exist in the imputation reference panel are dropped. These independent SNPs are shown in a Manhattan plot of the SDS score (Supplementary Figure S5.2).

Out of 37 SNPs tagging independent loci with $P_{SDS}$<1e-5, those at only three loci (*LCT* (F1), HLA (F2), *KITLG* (F17)) produced a strong signal in our ancient DNA time transect analysis ($\pi$>99%). The *LCT* and *HLA* signals are 2 of the 3 loci with $P_{SDS}$<5e-8 in Field et al. 2016. The third SDS genome-wide significant locus, *WDFY4* (F3), has a posterior probability of $\pi$=36% in our evaluation, with time transect analysis suggesting a greater probability selection before 2000 years BP ($\pi_B$=58%) than after ($\pi_A$<1%). The frequency of the tagging variant for *WDFY4* (F3) is around 5% in Steppe pastoralists and near zero in Western Hunter-Gatherers (WHG) and Early European Farmers (EEF). Its allele frequency increased rapidly with the arrival of Steppe in Europe and remained stable afterward. This allele frequency trajectory, along with our formal GLMM analysis, suggests that this signal may be an artifact of Steppe admixture, and that the significant SDS signal may be due to unresolved population structure (Supplementary Figures S5.3-S5.6)

The methodology in Field et al. 2016 is profoundly different from ours and uses a different type of data (not ancient DNA). Thus, while the two scans are maximally powered in the same time period, it is possible and even likely that the Field et al. 2016 methodology is sensitive to some genuine signals that our ancient DNA time transect study misses.

However, the far lower replication rates in the list of highlighted SNPs from Field et al. with less compelling P-values (5e-8 < $P_{SDS}$ < 1e-5) (1 of 34) than for the SNPs with strong P-values ($P_{SDS}$ < 5e-8) raises the possibility that most of the list of 37 SNPs highlighted from Field et al. 2016 were false-positives likely due unresolved population structure and the threshold for including a SNPs in the list was not stringent enough.

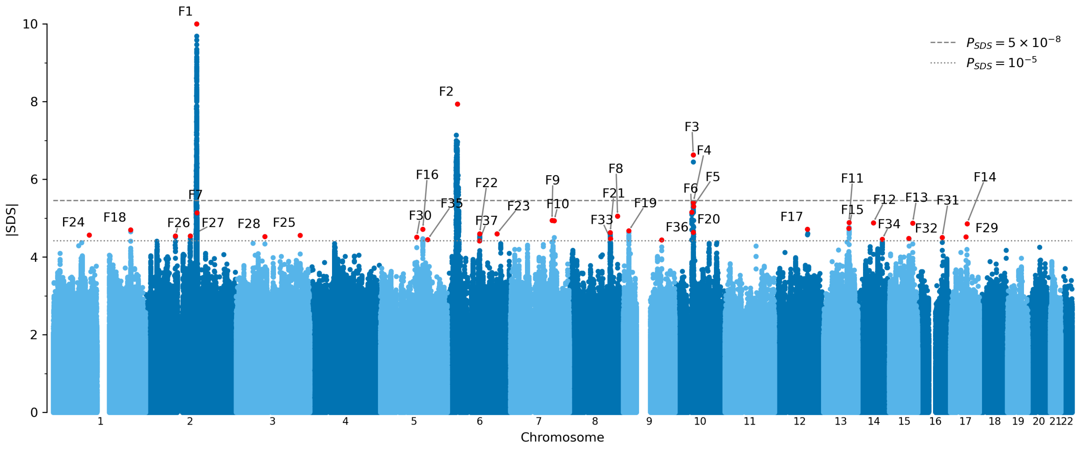

**Supplementary Figure S5.2:** Manhattan plot of SDS score from Field et al. 2016. Red circles indicate 37 independent tagging SNPs with $P_{SDS}$ < 1e-5. Each SNP is annotated with the panel name in the following Supplementary Figures 5.3-5.6.

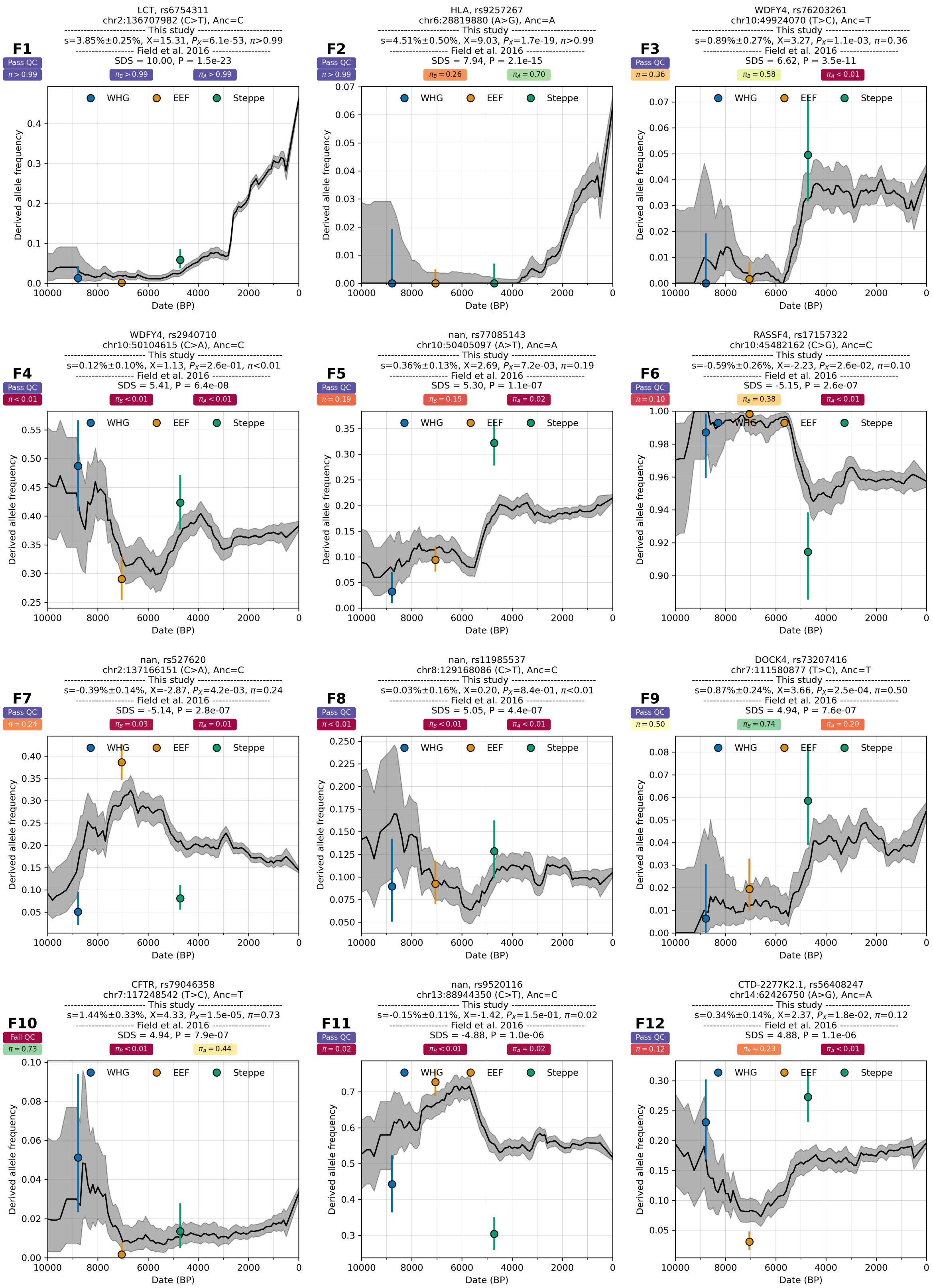

**Supplementary Figure S5.3:** Re-evaluating selection signals from Field et al. 2016. A and B refer to After and Before 2000 BP.

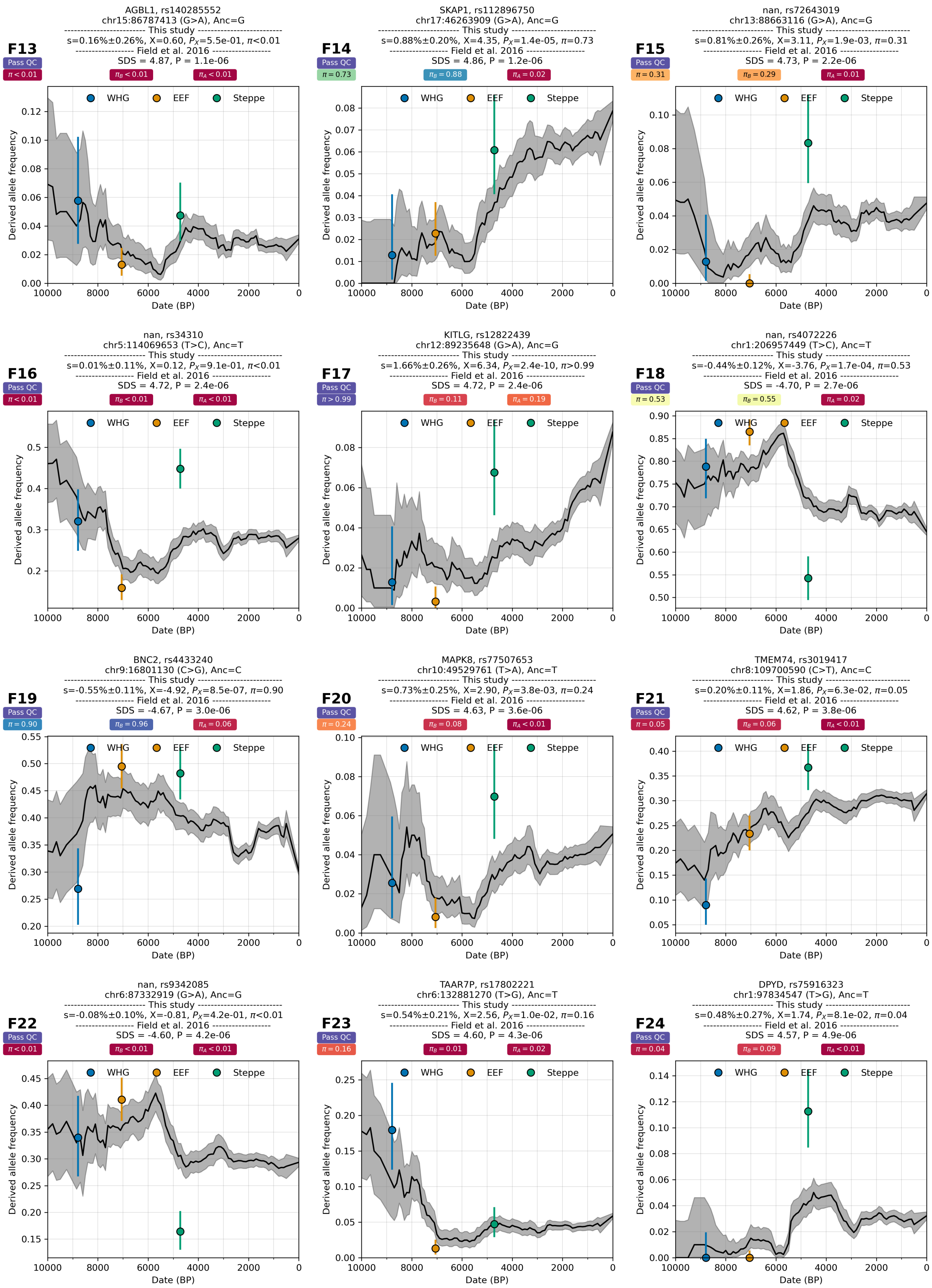

**Supplementary Figure S5.4:** Re-evaluating selection signals from Field et al. 2016. A and B refer to After and Before 2000 BP.

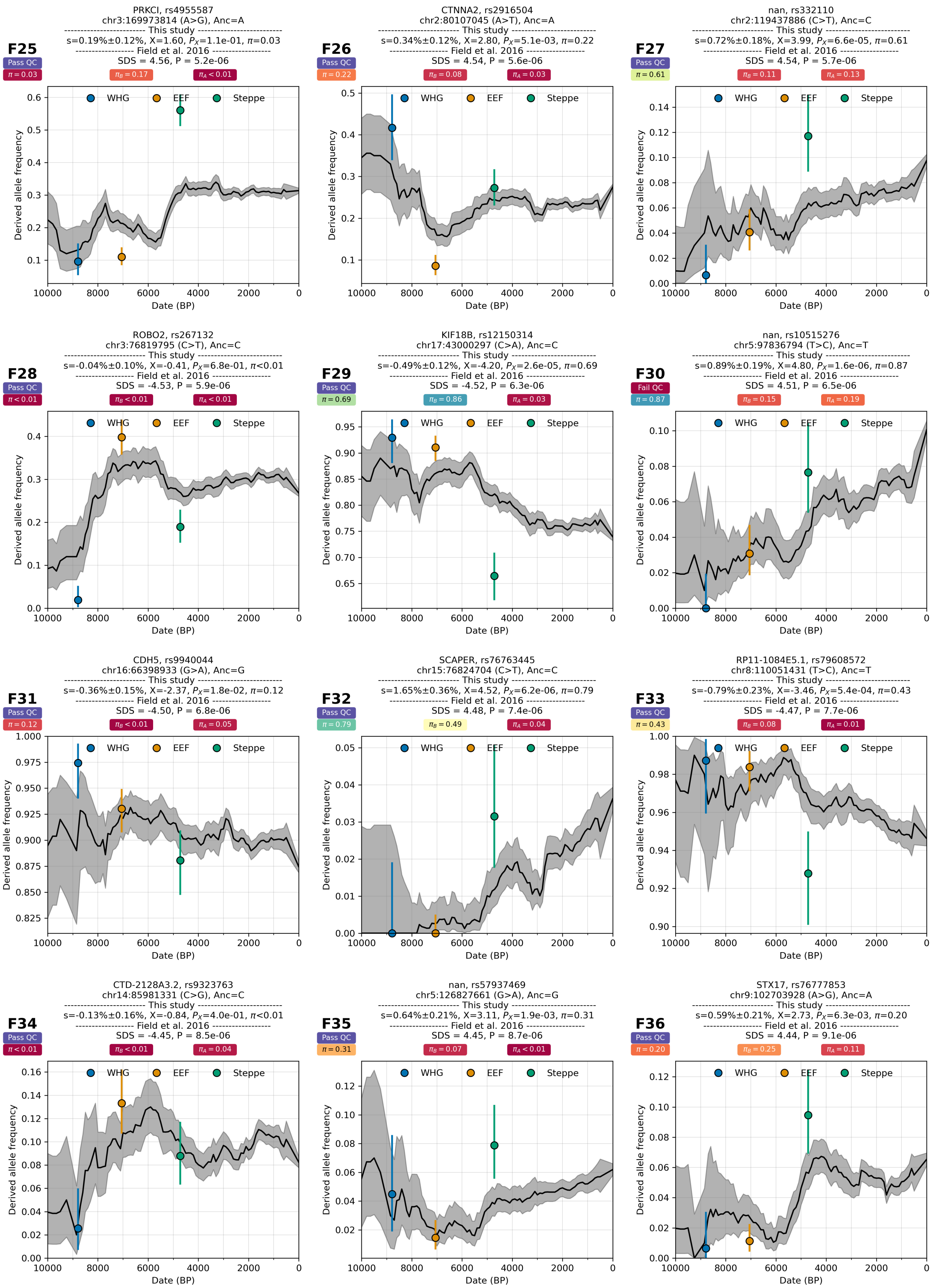

**Supplementary Figure S5.5:** Re-evaluating selection signals from Field et al. 2016. A and B refer to After and Before 2000 BP.

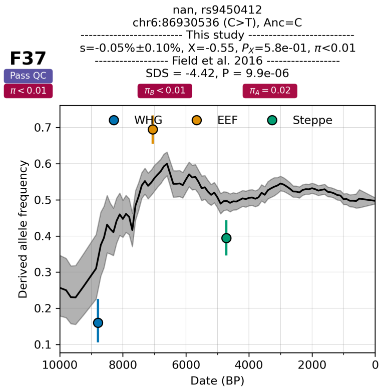

**Supplementary Figure S5.6:** Re-evaluating selection signals from Field et al. 2016. A and B refer to After and Before 2000 BP.

### Re-evaluation of results from Le et al. 2022

We evaluated 25 signals of selection at 24 loci detected during the Neolithic (N), Bronze Age (B), and Historical (H) periods, from Table 1 of Le et al. 2022[9]. Two of these signals are from SNP rs16891982 at the *SLC45A2* locus, which appear as a significant signal in both the Bronze Age and Historical periods. Two of these 24 SNPs are not present in the imputation reference panel and, therefore, are not re-evaluated here. This leaves 22 SNPs to re-evaluate, of which 9 produce strong signals ($\pi$>99%) in our full time transect analysis. We also analyzed three different time transects separately using the GLMM approach to mimic the three time periods of Le et al. 2022.

From the Neolithic period, we evaluated 10 candidate SNPs (rs281377, rs2310239, rs7517, rs12401678, rs4073089, rs57944517, rs10956504, rs2010410, rs6265, rs1669125). Only rs281377 at *FUT2* showed a compelling signal of selection ($\pi_N$>99%) during the Neolithic period, while rs57944517 at *RPL19* had a posterior probability ($\pi_N$=50%) for selection (Supplementary Figure S5.7).

From the Bronze Age period, two candidate SNPs (rs143482314, rs117124595) from Le et al. 2022 with signals of selection are not in the 1000 Genomes Project SNP set and were not analyzed here. Of the remaining 7 candidate SNPs (rs16891982, rs11636232, rs4906319, rs7555650, rs915843, rs10797666, rs4256954), only rs16891982 at *SLC45A2* showed a strong signal of selection ($\pi_B$>99%) during the Bronze Age period, while rs7555650 at *AGT* had a posterior probability ($\pi_B$=58%) for selection (Supplementary Figure S5.8).

From the Historical period, we evaluated 6 candidate SNPs (rs16891982, rs4988235, rs174550, rs653178, rs11603330, rs11096955) from Le et al. 2022. Five showed strong signals of selection ($\pi_H$>99%), with only rs11096955 having a lower posterior probability ($\pi_H$=10%) for selection (Supplementary Figure S5.9). All these 6 showed strong signals of selection ($\pi_B$>99%) during the earlier Bronze Age period as well, with only rs16891982 at the *SLC45A2* locus (Supplementary Figure S5.8, panel L11) being reported as a Bronze Age-specific signal.

Some co-authors of this study are also co-authors of Le et al. 2022. In their updated analysis, which includes additional measures to control for uncertainty in admixture proportions, allele frequency uncertainty in the source and target populations, and stochasticity in sampling, Le et al. now report 22 genome-wide significant hits at 21 loci. Of these, 17 validate with >99% posterior probability in our analysis (2 of 4 in the Neolithic, 3 of 6 in the Bronze Age, and 12 of 12 in the Historical Period) (Vagheesh Narasimhan, personal communication).

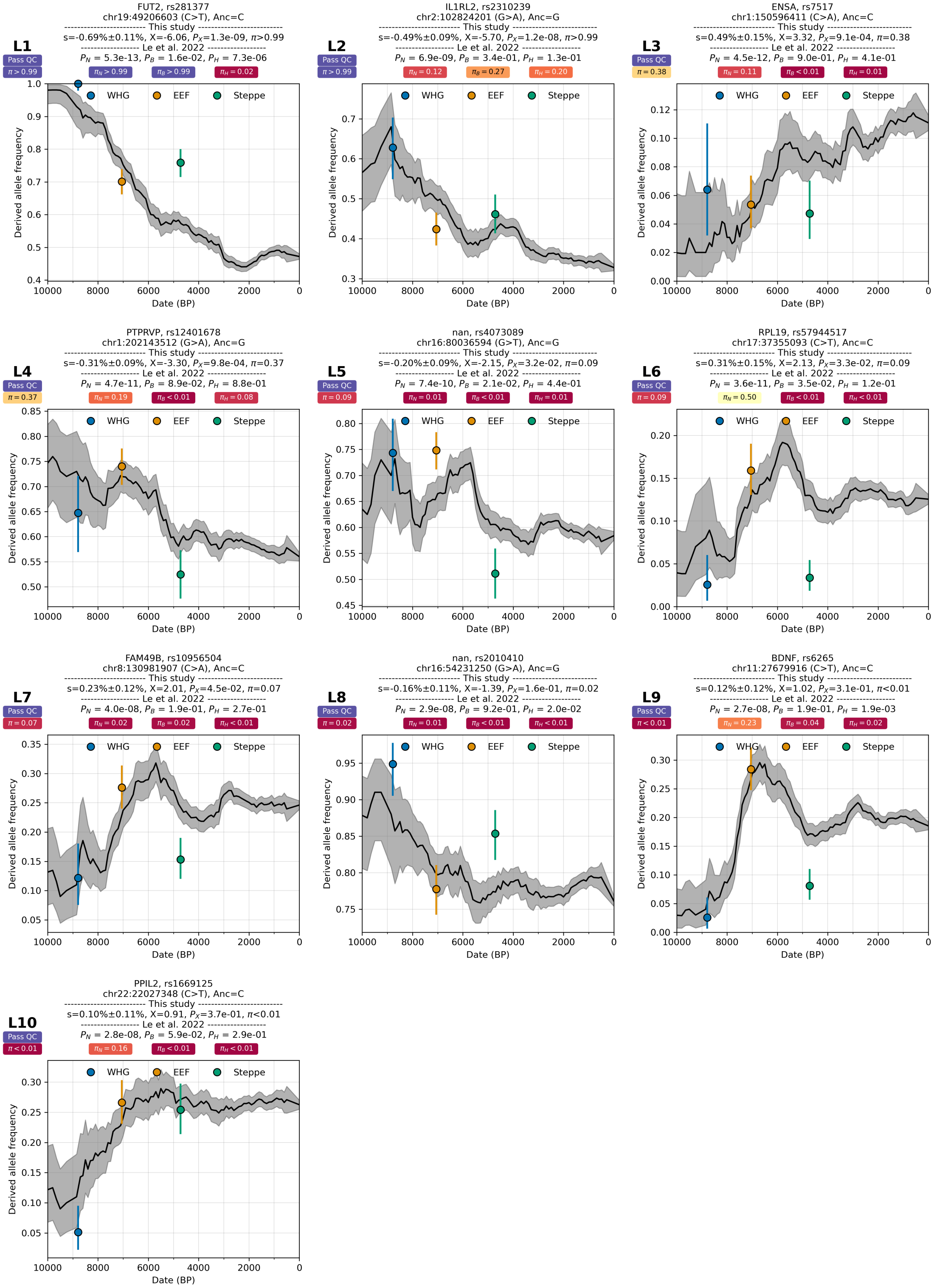

**Supplementary Figure S5.7:** Re-evaluating signals of selection from Le et al. 2022 from the Neolithic (N) period.

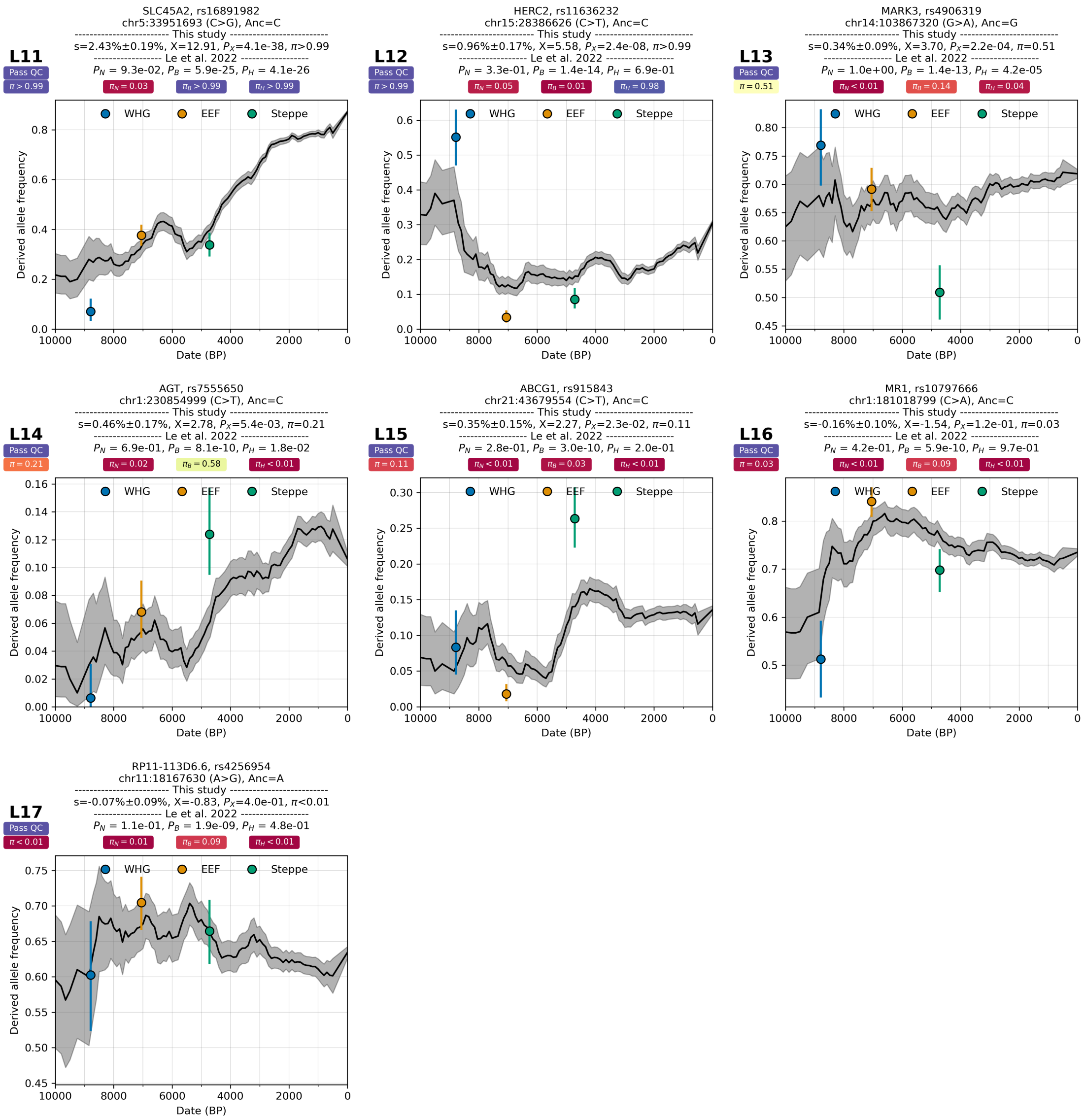

**Supplementary Figure S5.8:** Re-evaluating signals of selection from Le et al. 2022 from the Bronze Age (B).

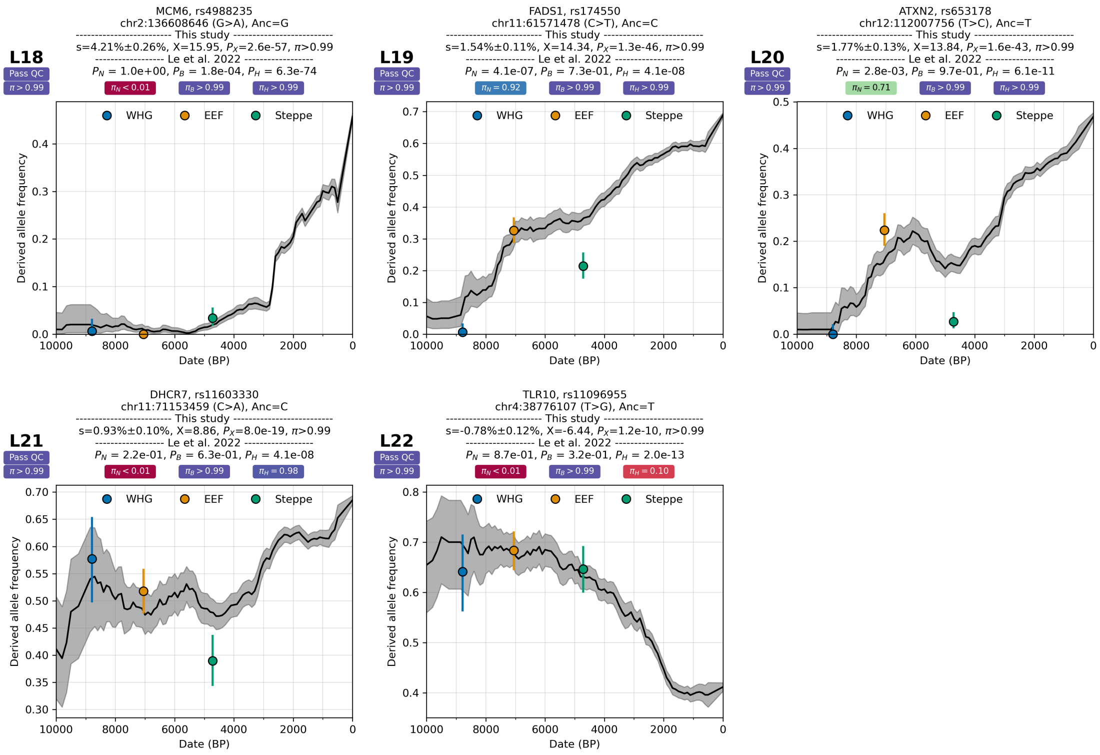

**Supplementary Figure S5.9:** Re-evaluating signals of selection from Le et al. 2022 for the Historical (H) period. Panel L11 from Supplementary Figure S5.8 is a candidate signal of selection for both the Bronze Age and the Historical period.

### Re-evaluation of results from Kerner et al. 2023

We evaluated 89 SNPs identified as candidates for positive selection on the derived allele from supplementary Table S2 of Kerner et al. 2023[10], along with 50 other candidate missense SNPs for negative selection from supplementary Table S7 of the same study. Only three candidate positive selection SNPs (rs4988235 at the *LCT* locus, rs185146 at the *SLC45A2* locus, and rs3130673 at the HLA locus) meet the genome-wide significant P-value threshold of 5e-8 for 'pbeta' test statistics of Kerner et al. 2023, and all three are replicated in our analysis with posterior probability ($\pi$>99%; Supplementary Figure S5.10). In our analysis, only 14 of 123 candidates passing QC show strong evidence for selection ($\pi$>99%), while an additional 10 are probable signals of selection in our re-analysis (50%<$\pi$<99%) This suggests most of the candidates are false positives, likely due to unresolved population structure and data artifacts (Supplementary Figures S5.11-S5.23).

These 139 SNPs are listed in Supplementary Table S5.2. Three positive selection candidate SNPs (rs11125238, rs34969536, rs4717903) are not in the 1000 Genomes Project reference panel, and therefore we did not analyze them. Three positive selection and ten negative selection candidate SNPs did not pass quality control (QC) in our study.

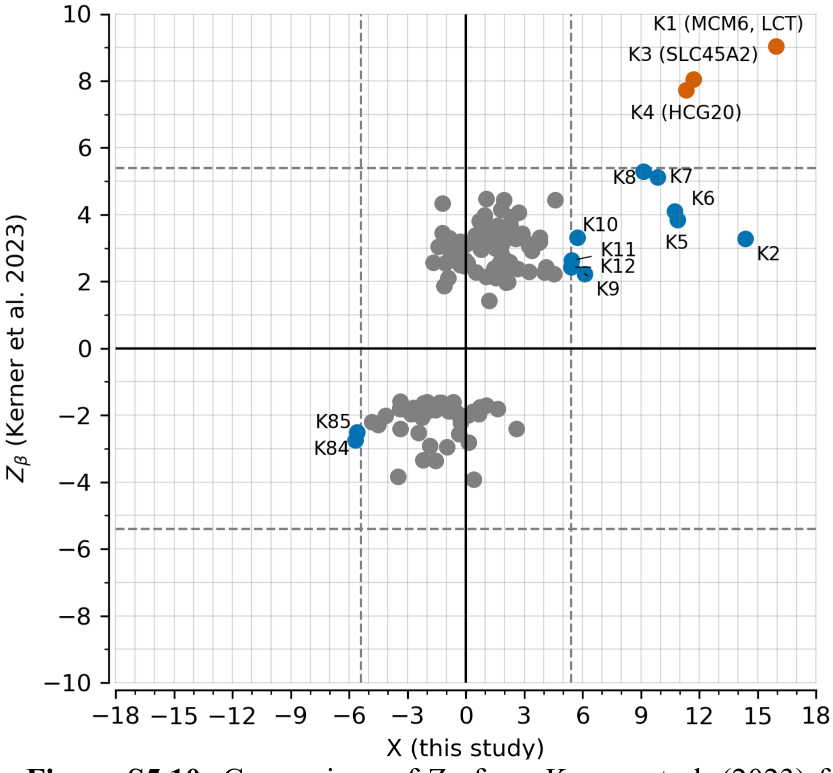

**Supplementary Figure S5.10:** Comparison of $Z_{\beta}$ from Kerner et al. (2023) for positive and negative selection candidates with the X score from this study. Labels indicate the panel ID for frequency trajectory plots in the supplementary figures below. The dashed gray line represents ±5.45, marking the classic genome-wide p-value threshold of 5e-8.

**Supplementary Table S5.2:** List of SNPs from Kerner et al. 2023 analyzed in this study. Panel ID is the identifier used in the supplementary figures below to mark each panel. The Variant ID is defined as the CHROM_POS_REF_ALT using the human genome reference assembly version hg19/GRCh37.

| RSID | Panel ID | Variant ID | Ancestral Allele | Gene name | Selection Type | QC | Kerner et al. 2023 | | $\pi$ |
|---|---|---|---|---|---|---|---|---|---|
| | | | | | | | $P_{sel}$ | $P_{\beta}$ | |
| rs4988235 | K1 | 2_136608646_G_A | G | MCM6 | Positive | Pass | <2.0e-6 | 1.58E-19 | >0.99 |
| rs174537 | K2 | 11_61552680_G_T | T | TMEM258 | Positive | Pass | 2.70E-05 | 0.001 | >0.99 |
| rs185146 | K3 | 5_33952106_C_T | C | SLC45A2 | Positive | Pass | <2.0e-6 | 8.71E-16 | >0.99 |
| rs3130673 | K4 | 6_30746519_G_T | G | HCG20 | Positive | Pass | <2.0e-6 | 1.13E-14 | >0.99 |
| rs10765770 | K5 | 11_88513636_G_A | G | GRM5 | Positive | Pass | 2.00E-06 | 0.000123 | >0.99 |
| rs4705844 | K6 | 5_131365784_A_C | A | AC034228.2-IL3 | Positive | Pass | <2.0e-6 | 4.26E-05 | >0.99 |
| rs10008492 | K7 | 4_38765720_C_T | T | RNA5SP158-TLR10 | Positive | Pass | 1.20E-05 | 3.14E-07 | >0.99 |
| rs4792831 | K8 | 17_44206665_G_A | G | RNU7-101P | Positive | Pass | 1.10E-05 | 1.16E-07 | >0.99 |
| rs7141996 | K9 | 14_93094298_G_A | G | RIN3 | Positive | Pass | 0.000849 | 0.0256 | >0.99 |
| rs11674302 | K10 | 2_102887128_T_C | T | AC007248.6-IL1RL1 | Positive | Pass | 2.80E-05 | 0.000935 | >0.99 |
| rs36083353 | K11 | 4_42041692_A_T | A | SLC30A9 | Positive | Pass | 0.000209 | 0.0085 | 0.99 |
| rs1229758 | K12 | 7_114229139_G_A | G | FOXP2 | Positive | Pass | 0.000394 | 0.0152 | 0.99 |
| rs10095927 | K13 | 8_10729177_G_T | G | RP11-177H2.2-XKR6 | Positive | Pass | <2.0e-6 | 8.67E-06 | 0.82 |
| rs11042594 | K14 | 11_2117403_A_G | G | H19-IGF2 | Positive | Pass | 0.000853 | 0.026 | 0.79 |
| rs10513801 | K15 | 3_185822353_T_G | T | ETV5 | Positive | Pass | 0.000511 | 0.0148 | 0.66 |
| rs17605165 | K16 | 16_11297562_C_T | C | RP11-396B14.2 | Positive | Pass | 0.000797 | 0.0227 | 0.63 |
| rs8176635 | K17 | 9_136152009_G_A | G | ABO | Positive | Pass | 5.40E-05 | 0.00135 | 0.55 |
| rs11057805 | K18 | 12_125246993_G_A | G | NCOR2-SCARB1 | Positive | Pass | 2.60E-05 | 0.000892 | 0.54 |
| rs389663 | K19 | 6_117868051_T_C | T | DCBLD1 | Positive | Pass | 4.00E-05 | 0.00152 | 0.53 |
| rs8022676 | K20 | 14_77825746_A_C | C | RP11-493G17.4 | Positive | Pass | 9.10E-05 | 0.00352 | 0.4 |
| rs72700902 | K21 | 1_150633347_C_T | C | GOLPH3L | Positive | Pass | 5.80E-05 | 0.00216 | 0.36 |
| rs10507766 | K22 | 13_69940025_A_G | A | LINC00401-SRSF1P1 | Positive | Pass | 0.000563 | 0.0213 | 0.35 |
| rs8033465 | K23 | 15_69773781_G_T | G | RP11-279F6.1 | Positive | Pass | 6.90E-05 | 0.00173 | 0.34 |
| rs13307276 | K24 | 7_150061525_T_C | T | REPIN1 | Positive | Pass | 2.00E-05 | 0.000597 | 0.25 |
| rs7958347 | K25 | 12_113325061_C_T | C | RPH3A | Positive | Pass | <2.0e-6 | 4.76E-05 | 0.2 |
| rs55846849 | K26 | 5_460710_G_A | G | EXOC3 | Positive | Pass | 0.000618 | 0.0173 | 0.19 |
| rs10496379 | K27 | 2_104602547_A_T | A | RP11-76I14.1 | Positive | Pass | 3.00E-05 | 0.00102 | 0.16 |
| rs721992 | K28 | 10_61539402_G_T | G | LINC00948-CCDC6 | Positive | Pass | 1.70E-05 | 0.000517 | 0.15 |
| rs6694101 | K29 | 1_2020489_C_T | C | PRKCZ | Positive | Pass | 0.000485 | 0.0172 | 0.12 |
| rs3858036 | K30 | 9_2968107_G_A | G | CARM1P1 | Positive | Pass | <2.0e-6 | 8.57E-05 | 0.12 |
| rs1055472 | K31 | 12_125510104_G_A | A | BRI3BP | Positive | Pass | 7.00E-06 | 0.000197 | 0.11 |
| rs1531212 | K32 | 19_13951830_G_A | G | MIR24-2 | Positive | Pass | 0.000303 | 0.00981 | 0.1 |
| rs10841899 | K33 | 12_8716873_G_A | G | RP11-561P12.5 | Positive | Pass | 0.001341 | 0.0465 | 0.09 |
| rs1047616 | K34 | 17_25642522_G_A | A | WSB1 | Positive | Pass | 0.001428 | 0.0486 | 0.08 |
| rs713217 | K35 | 11_69579070_T_C | T | RP11-300I6.5-AP001888.1 | Positive | Pass | 3.00E-05 | 0.00101 | 0.08 |
| rs586554 | K36 | 9_22796540_T_C | C | RP11-399D6.2 | Positive | Pass | 0.000883 | 0.0287 | 0.08 |
| rs10515671 | K37 | 5_151845342_A_G | A | NMUR2-CTC-550M4.1 | Positive | Pass | 1.60E-05 | 8.95E-06 | 0.07 |
| rs4922511 | K38 | 10_48402818_T_C | C | RBP3-GDF2 | Positive | Pass | 0.00037 | 0.0116 | 0.06 |
| rs2974658 | K39 | 5_2960267_G_T | G | RP11-35O7.1 | Positive | Pass | 0.000175 | 0.00625 | 0.06 |
| rs62054458 | K40 | 17_13672024_C_T | C | COX10-AS1 | Positive | Pass | 2.70E-05 | 0.000981 | 0.05 |
| rs3780710 | K41 | 9_132942794_G_A | G | NCS1 | Positive | Pass | 0.000352 | 0.0109 | 0.05 |
| rs1076005 | K42 | 18_5973037_C_T | C | L3MBTL4 | Positive | Pass | 5.10E-05 | 0.00188 | 0.05 |
| rs9840198 | K43 | 3_184770686_C_T | C | VPS8 | Positive | Pass | <2.0e-6 | 3.24E-05 | 0.05 |
| rs75770273 | K44 | 6_53877078_C_T | C | ERHP2 | Positive | Pass | 8.10E-05 | 0.00195 | 0.05 |
| rs7422 | K45 | 14_89884022_A_C | A | FOXN3 | Positive | Pass | 9.30E-05 | 0.00371 | 0.05 |
| rs4280687 | K46 | 4_156105105_C_T | C | RBM46-RP11-92A5.2 | Positive | Pass | 0.000238 | 0.0101 | 0.04 |
| rs13009356 | K47 | 2_8530522_G_A | G | LINC00299-AC011747.3 | Positive | Pass | 0.00024 | 0.0103 | 0.04 |
| rs8079769 | K48 | 17_9588455_G_A | G | USP43 | Positive | Pass | 2.00E-05 | 0.000238 | 0.03 |
| rs12666876 | K49 | 7_155417249_G_A | G | AC009403.2 | Positive | Pass | 0.000492 | 0.0345 | 0.03 |
| rs11884056 | K50 | 2_85339839_C_T | C | LSM3P3-TCF7L1 | Positive | Pass | 0.000105 | 0.00274 | 0.03 |
| rs34404720 | K51 | 2_86879419_A_G | A | CHMP3 | Positive | Pass | 0.000601 | 0.0169 | 0.02 |
| rs3003615 | K52 | 9_130997892_G_A | G | DNM1 | Positive | Pass | 6.50E-05 | 0.00238 | 0.02 |
| rs12580172 | K53 | 12_1623989_G_A | G | LINC00942-WNT5B | Positive | Pass | 0.006145 | 0.155 | 0.01 |
| rs6900553 | K54 | 6_10231925_C_T | C | RNU6ATAC21P-RP1-290I10.2 | Positive | Pass | <2.0e-6 | 0.000468 | 0.01 |
| rs10188894 | K55 | 2_6643733_A_G | A | AC021021.2 | Positive | Pass | <2.0e-6 | 1.40E-05 | 0.01 |
| rs12033048 | K56 | 1_246429681_T_C | T | SMYD3 | Positive | Pass | 2.10E-05 | 0.000555 | 0.01 |
| rs62436708 | K57 | 7_1323037_C_T | C | AC073094.4-MICALL2 | Positive | Pass | 0.001715 | 0.0619 | <0.01 |
| rs8031453 | K58 | 15_91108674_A_C | A | CRTC3 | Positive | Pass | 2.10E-05 | 0.000483 | <0.01 |
| rs6698312 | K59 | 1_164206815_G_T | G | U3-NMNAT1P2 | Positive | Pass | 4.50E-05 | 0.00169 | <0.01 |

| | | | | | | | | | |
|---|---|---|---|---|---|---|---|---|---|
| rs62043998 | K60 | 16_81921203_C_T | C | PLCG2 | Positive | Pass | 1.60E-05 | 7.64E-06 | <0.01 |
| rs11059425 | K61 | 12_128441207_T_G | T | LINC00507-RP11-349K16.1 | Positive | Pass | 9.00E-06 | 0.0107 | <0.01 |
| rs12715075 | K62 | 3_2620454_C_T | T | CNTN4 | Positive | Pass | 0.000484 | 0.0326 | <0.01 |
| rs4745827 | K63 | 10_65659657_A_G | A | RP11-170M17.2 | Positive | Pass | <2.0e-6 | 6.80E-05 | <0.01 |
| rs2426652 | K64 | 20_55294098_A_G | A | AL133232.1 | Positive | Pass | 3.00E-06 | 0.000135 | <0.01 |
| rs1009543 | K65 | 10_3319474_A_G | A | RP11-195B3.1-RP11-482E14.1 | Positive | Pass | 0.001032 | 0.0355 | <0.01 |
| rs10501651 | K66 | 11_87527561_G_A | G | RP11-665E10.5 | Positive | Pass | 2.90E-05 | 0.000965 | <0.01 |
| rs4405041 | K67 | 9_99088478_T_G | G | SLC35D2 | Positive | Pass | 9.00E-05 | 0.00338 | <0.01 |
| rs10809509 | K68 | 9_1162013_A_C | A | RPS27AP14 | Positive | Pass | 8.40E-05 | 0.0031 | <0.01 |
| rs12711473 | K69 | 16_87224293_A_G | G | C16orf95 | Positive | Pass | 7.90E-05 | 0.00292 | <0.01 |
| rs3957465 | K70 | 5_5817703_G_A | G | KIAA0947-CTC-471C19.1 | Positive | Pass | 9.90E-05 | 0.00449 | <0.01 |
| rs4374563 | K71 | 3_31776980_G_A | G | OSBPL10 | Positive | Pass | 3.00E-06 | 0.000141 | <0.01 |
| rs176482 | K72 | 7_105730038_G_A | G | SYPL1 | Positive | Pass | 0.000612 | 0.0225 | <0.01 |
| rs2291897 | K73 | 3_33419422_C_T | C | FBXL2 | Positive | Pass | 2.20E-05 | 0.000692 | <0.01 |
| rs2619105 | K74 | 10_118977123_C_A | C | KCNK18-RP11-501J20.5 | Positive | Pass | <2.0e-6 | 0.00882 | <0.01 |
| rs11189359 | K75 | 10_99517616_C_A | C | ZFYVE27 | Positive | Pass | <2.0e-6 | 0.00121 | <0.01 |
| rs10841952 | K76 | 12_22289304_A_G | A | ST8SIA1 | Positive | Pass | 0.000216 | 0.00898 | <0.01 |
| rs35345724 | K77 | 7_136627956_A_G | A | KRT8P51 | Positive | Pass | 0.000405 | 0.0127 | <0.01 |
| rs8103030 | K78 | 19_23568661_G_A | G | CTB-175P5.1 | Positive | Pass | 5.40E-05 | 0.00135 | <0.01 |
| rs1364095 | K79 | 16_79566150_G_C | G | RP11-467I17.1-MAF | Positive | Pass | 6.00E-05 | 0.00221 | <0.01 |
| rs2631934 | K80 | 8_21052978_G_A | A | AC021613.1-RP11-24P4.1 | Positive | Pass | 0.000164 | 0.0139 | <0.01 |
| rs9314061 | K81 | 5_164760466_C_T | C | CTB-181F24.1-CTC-535M15.2 | Positive | Pass | 0.000244 | 0.0106 | <0.01 |
| rs458552 | K82 | 9_4833437_C_T | C | RCL1 | Positive | Pass | 0.000206 | 0.00829 | <0.01 |
| rs4863449 | K83 | 4_189987905_C_T | C | RP11-818C3.1-RP11-706F1.1 | Positive | Pass | 5.10E-05 | 0.00189 | <0.01 |
| rs12146727 | K84 | 12_7170336_G_A | G | C1S | Negative | Pass | 0.000497 | 0.006 | >0.99 |
| rs3775291 | K85 | 4_187004074_C_T | C | TLR3 | Negative | Pass | 0.001151 | 0.0121 | >0.99 |
| rs3814541 | K86 | 9_109689752_C_T | C | ZNF462 | Negative | Pass | 0.002527 | 0.0279 | 0.88 |
| rs2305637 | K87 | 3_47045846_C_T | C | NBEAL2 | Negative | Pass | 0.001961 | 0.0223 | 0.79 |
| rs7722711 | K88 | 5_75906851_T_C | T | IQGAP2 | Negative | Pass | 0.004925 | 0.0439 | 0.67 |
| rs948962 | K89 | 11_76919478_C_A | A | MYO7A | Negative | Pass | <2.0e-6 | 0.000121 | 0.44 |
| rs4916685 | K90 | 5_89979698_C_T | C | GPR98 | Negative | Pass | 0.005972 | 0.0692 | 0.4 |
| rs2366926 | K91 | 5_89988504_A_G | A | GPR98 | Negative | Pass | 0.009795 | 0.111 | 0.39 |
| rs17545756 | K92 | 7_150732812_C_T | C | ABCB8 | Negative | Pass | 0.001789 | 0.0162 | 0.39 |
| rs1064583 | K93 | 6_116446576_A_G | G | COL10A1 | Negative | Pass | 0.006171 | 0.0707 | 0.34 |
| rs868738 | K94 | 10_115381747_G_A | G | NRAP | Negative | Pass | 0.004358 | 0.0496 | 0.22 |
| rs2274654 | K95 | 9_98691137_T_C | T | ERCC6L2 | Negative | Pass | 0.00503 | 0.0509 | 0.19 |
| rs1799977 | K96 | 3_37053568_A_G | A | MLH1 | Negative | Pass | 0.006816 | 0.0762 | 0.19 |
| rs1051489 | K97 | 5_32400266_A_G | A | ZFR | Negative | Pass | 0.0014 | 0.016 | 0.17 |
| rs11209026 | K98 | 1_67705958_G_A | G | IL23R | Negative | Pass | 0.00726 | 0.0696 | 0.16 |
| rs2298260 | K99 | 9_21029330_T_C | T | PTPLAD2 | Negative | Pass | 0.001052 | 0.0114 | 0.13 |
| rs34536443 | K100 | 19_10463118_G_C | G | TYK2 | Negative | Pass | 0.004458 | 0.039 | 0.1 |
| rs13009282 | K101 | 2_68364478_T_C | T | WDR92 | Negative | Pass | 4.00E-05 | 0.000847 | 0.1 |
| rs34899 | K102 | 5_95091201_A_G | G | RHOBTB3 | Negative | Pass | 0.009069 | 0.102 | 0.09 |
| rs1124649 | K103 | 2_27260469_G_A | G | TMEM214 | Negative | Pass | 0.009551 | 0.108 | 0.07 |
| rs2275477 | K104 | 1_36886117_C_T | C | OSCP1 | Negative | Pass | 0.005291 | 0.0628 | 0.06 |
| rs678892 | K105 | 15_55632859_G_T | T | PIGB | Negative | Pass | 0.000276 | 0.00353 | 0.05 |
| rs77491573 | K106 | 12_124288264_G_A | G | DNAH10 | Negative | Pass | 0.007884 | 0.0708 | 0.03 |
| rs2287059 | K107 | 2_10717806_C_T | C | NOL10 | Negative | Pass | 0.005546 | 0.0648 | 0.03 |
| rs11080134 | K108 | 17_29161503_A_G | A | ATAD5 | Negative | Pass | 3.90E-05 | 0.000784 | 0.03 |
| rs16885 | K109 | 6_16306751_G_A | G | ATXN1 | Negative | Pass | 0.009295 | 0.105 | 0.02 |
| rs3829765 | K110 | 14_58605790_G_A | G | C14orf37 | Negative | Pass | 0.009228 | 0.104 | 0.01 |
| rs11568591 | K111 | 17_48761053_G_A | G | ABCC3 | Negative | Pass | 0.009769 | 0.087 | <0.01 |
| rs3803716 | K112 | 16_24802325_C_T | C | TNRC6A | Negative | Pass | 0.000263 | 0.00321 | <0.01 |
| rs2276774 | K113 | 3_122646828_A_G | A | SEMA5B | Negative | Pass | 0.005904 | 0.0598 | <0.01 |
| rs8052655 | K114 | 16_67409180_G_A | G | LRRC36 | Negative | Pass | 0.00887 | 0.0791 | <0.01 |
| rs4935502 | K115 | 10_55955444_T_G | T | PCDH15 | Negative | Pass | 0.0048 | 0.049 | <0.01 |
| rs2303291 | K116 | 2_24431184_C_T | C | ITSN2 | Negative | Pass | 0.009605 | 0.108 | <0.01 |
| rs16853333 | K117 | 2_168108032_G_A | G | XIRP2 | Negative | Pass | 0.0062 | 0.0533 | <0.01 |
| rs12609039 | K118 | 19_11348960_G_A | G | DOCK6 | Negative | Pass | 2.40E-05 | 8.58E-05 | <0.01 |
| rs2270856 | K119 | 2_234741808_C_T | C | MROH2A | Negative | Pass | 0.006914 | 0.0589 | <0.01 |
| rs11204546 | K120 | 1_248059712_T_C | C | OR2W3 | Negative | Pass | 0.000999 | 0.0105 | <0.01 |
| rs2306541 | K121 | 12_133428242_G_A | G | CHFR | Negative | Pass | 0.002304 | 0.0264 | <0.01 |
| rs10083789 | K122 | 16_23080634_C_A | C | USP31 | Negative | Pass | 0.000395 | 0.00495 | <0.01 |
| rs45559835 | K123 | 5_155935708_G_A | G | SGCD | Negative | Pass | 0.005314 | 0.0463 | <0.01 |
| rs17080528 | K124 | 3_49389842_C_T | C | GPX1 | Positive | Fail | 1.90E-05 | 0.000557 | 0.41 |
| rs4077347 | K125 | 16_29036915_G_A | G | LAT-CTB-134H23.2 | Positive | Fail | 0.005233 | 0.182 | 0.09 |
| rs11150556 | K126 | 16_83270541_T_C | T | CDH13 | Positive | Fail | 0.000517 | 0.0201 | 0.01 |
| rs61732547 | K127 | 9_113233652_C_T | C | SVEP1 | Negative | Fail | <2.0e-6 | 1.42E-07 | >0.99 |
| rs3732380 | K128 | 3_39307562_C_T | C | CX3CR1 | Negative | Fail | 1.60E-05 | 4.54E-06 | 0.95 |

| | | | | | | | | | |
|---|---|---|---|---|---|---|---|---|---|
| rs113575767 | K129 | 11_125765587_C_T | C | PUS3 | Negative | Fail | 1.70E-05 | 5.05E-06 | 0.5 |
| rs2291375 | K130 | 3_105264129_G_A | G | ALCAM | Negative | Fail | 4.00E-05 | 7.16E-05 | 0.49 |
| rs1042311 | K131 | 22_46627780_C_T | C | PPARA | Negative | Fail | 0.000289 | 0.000853 | 0.3 |
| rs7594497 | K132 | 2_55872538_T_C | T | PNPT1 | Negative | Fail | 6.80E-05 | 0.000295 | 0.12 |
| rs2298316 | K133 | 10_101147692_G_A | G | CNNM1 | Negative | Fail | 0.005556 | 0.056 | 0.09 |
| rs2271694 | K134 | 10_71874784_C_T | C | AIFM2 | Negative | Fail | 0.000299 | 0.000924 | <0.01 |
| rs16945138 | K135 | 17_11556248_C_T | C | DNAH9 | Negative | Fail | 8.40E-05 | 0.000265 | <0.01 |
| rs2232607 | K136 | 20_36993333_A_G | A | LBP | Negative | Fail | 0.003067 | 0.0245 | <0.01 |
| rs11125238 | NA | NA | G | RNU6-439P-RPL7P13 | Positive | NA | 3.10E-05 | 0.00103 | NA |
| rs34969536 | NA | NA | A | AC008984.6 | Positive | NA | 0.000104 | 0.00261 | NA |
| rs4717903 | NA | NA | C | GTF2I | Positive | NA | 0.000464 | 0.0284 | NA |

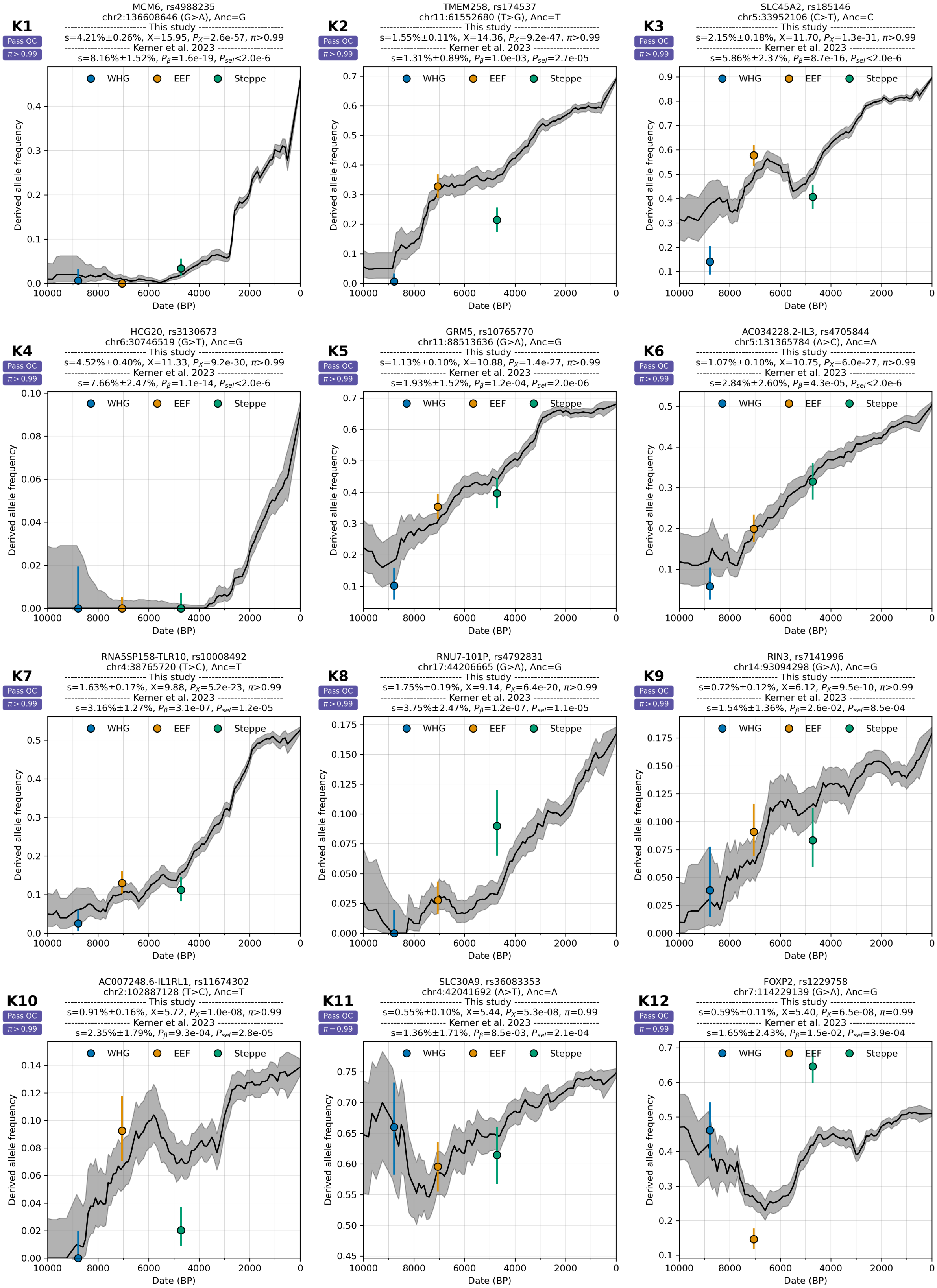

**Supplementary Figure S5.11:** Positive selection cases from Kerner et al. 2023 passing QC.

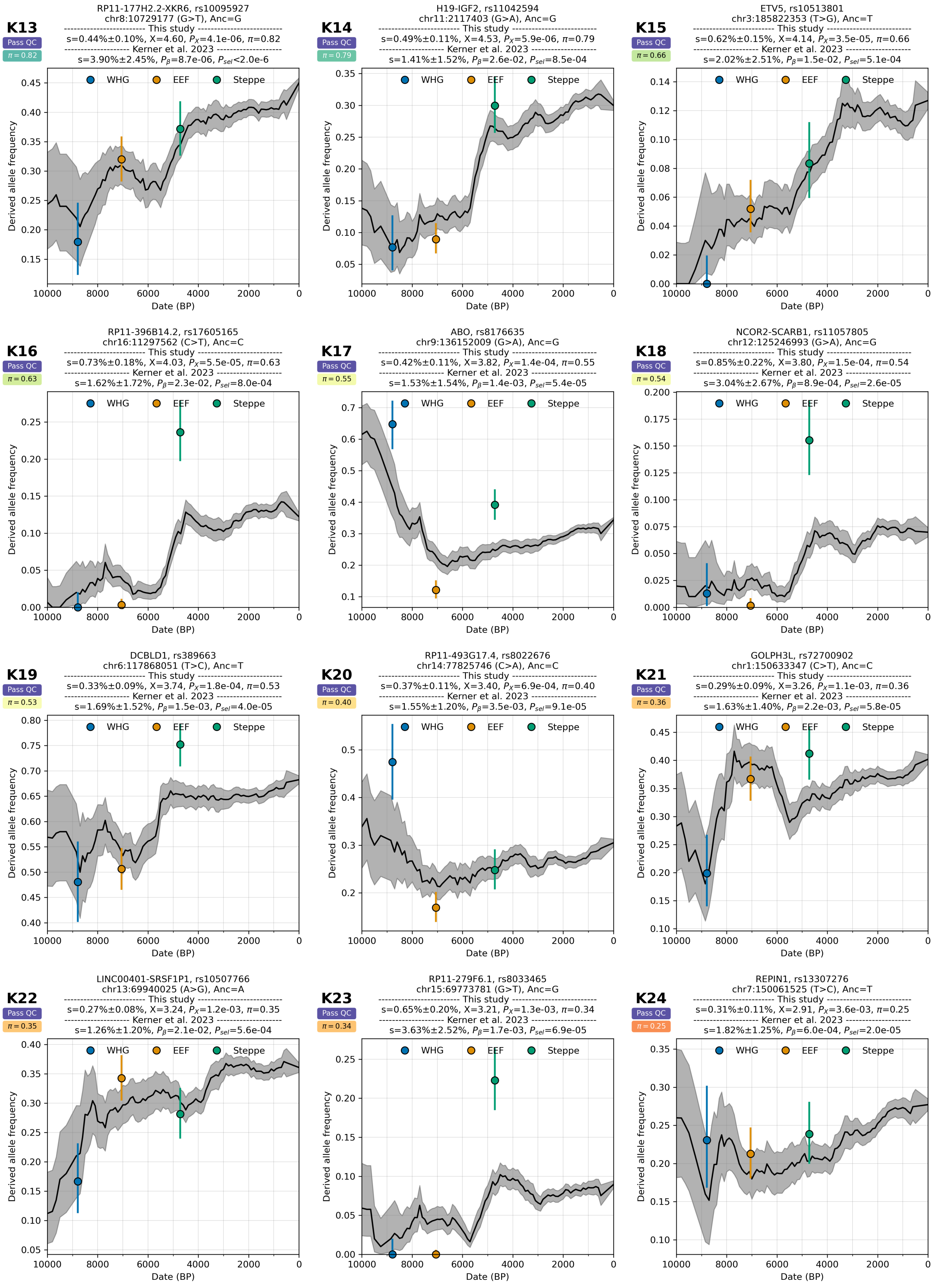

**Supplementary Figure S5.12:** Positive selection cases from Kerner et al. 2023 passing QC.

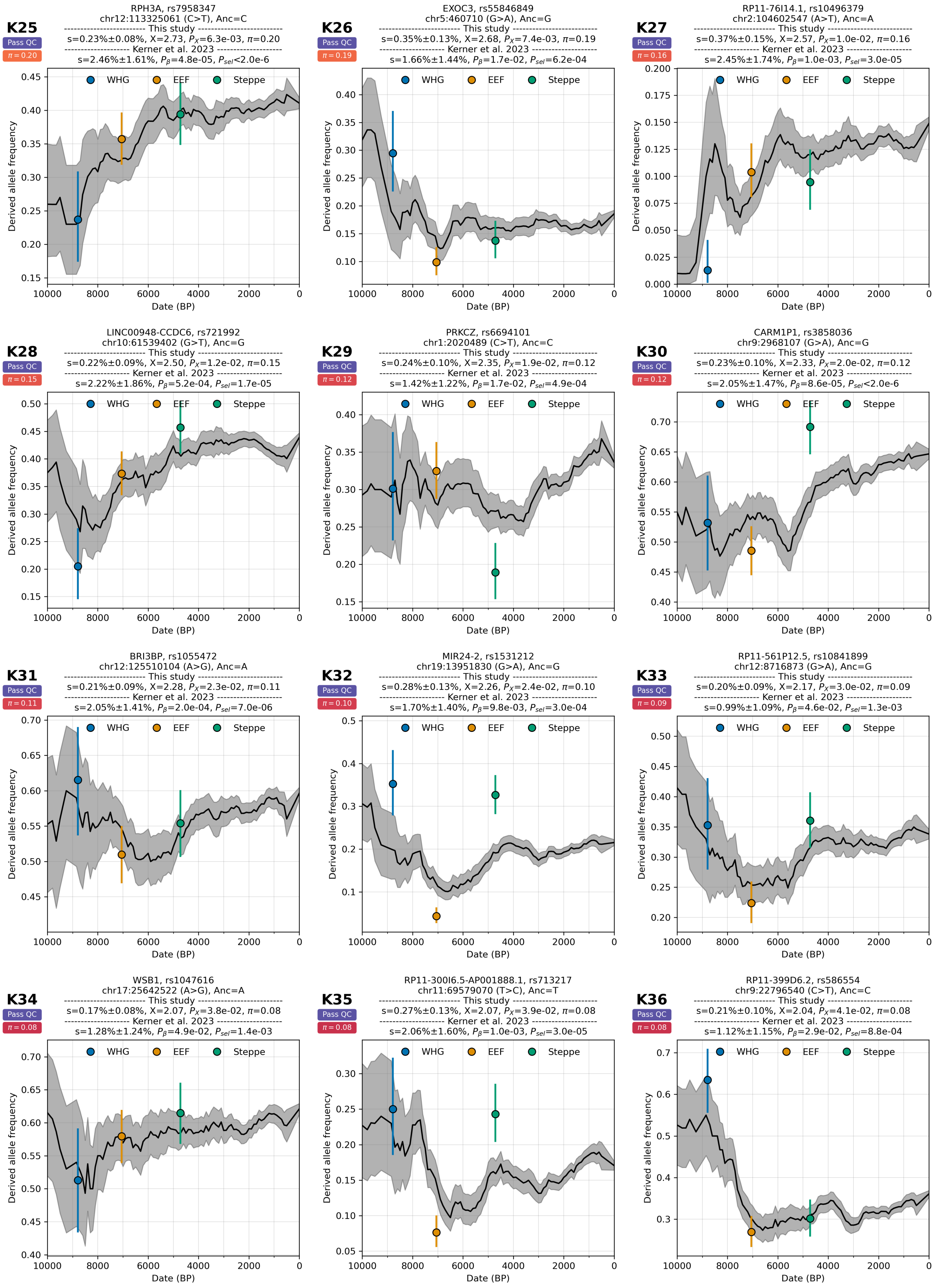

**Supplementary Figure S5.13:** Positive selection cases from Kerner et al. 2023 passing QC.

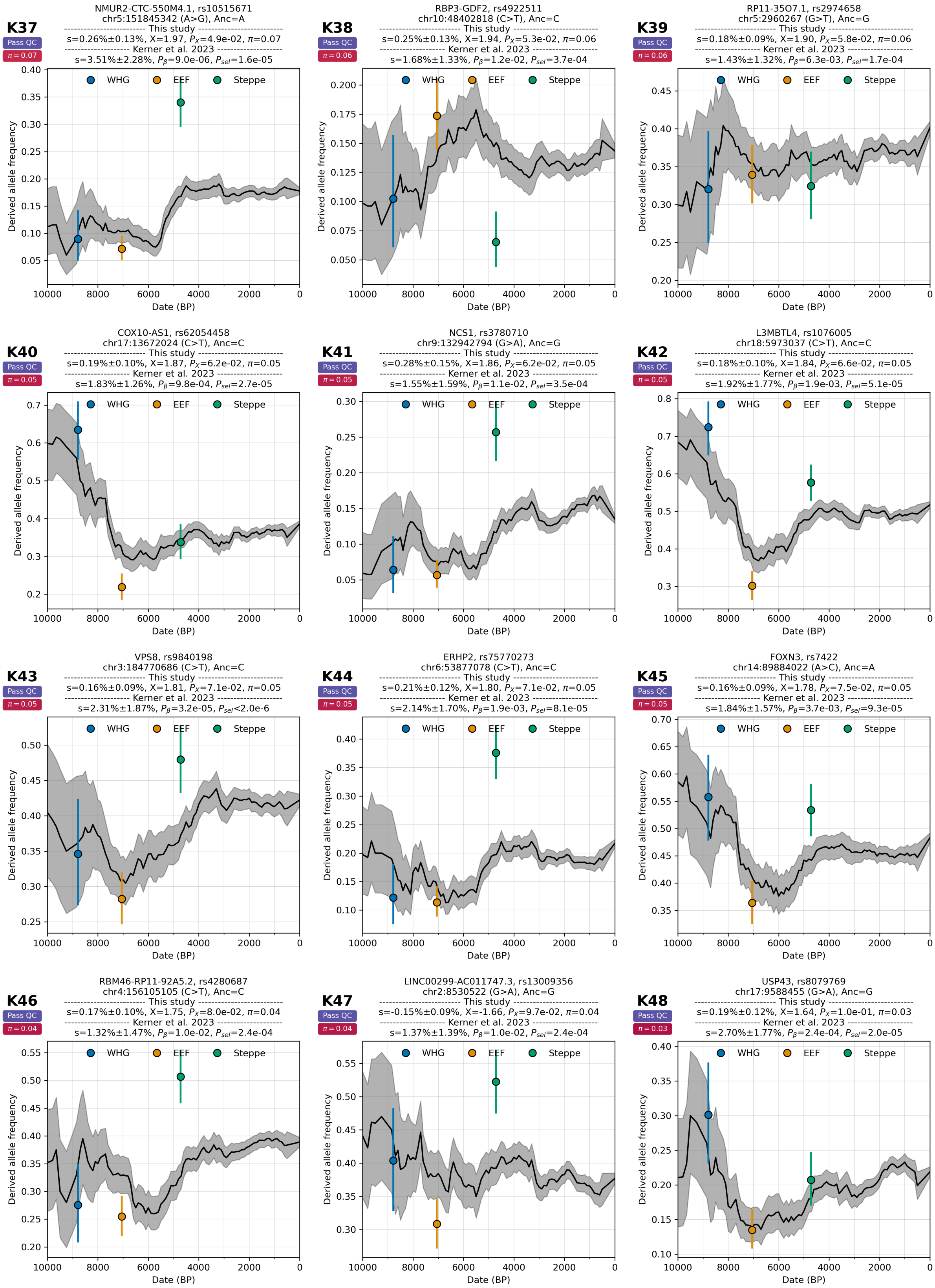

**Supplementary Figure S5.14:** Positive selection cases from Kerner et al. 2023 passing QC.

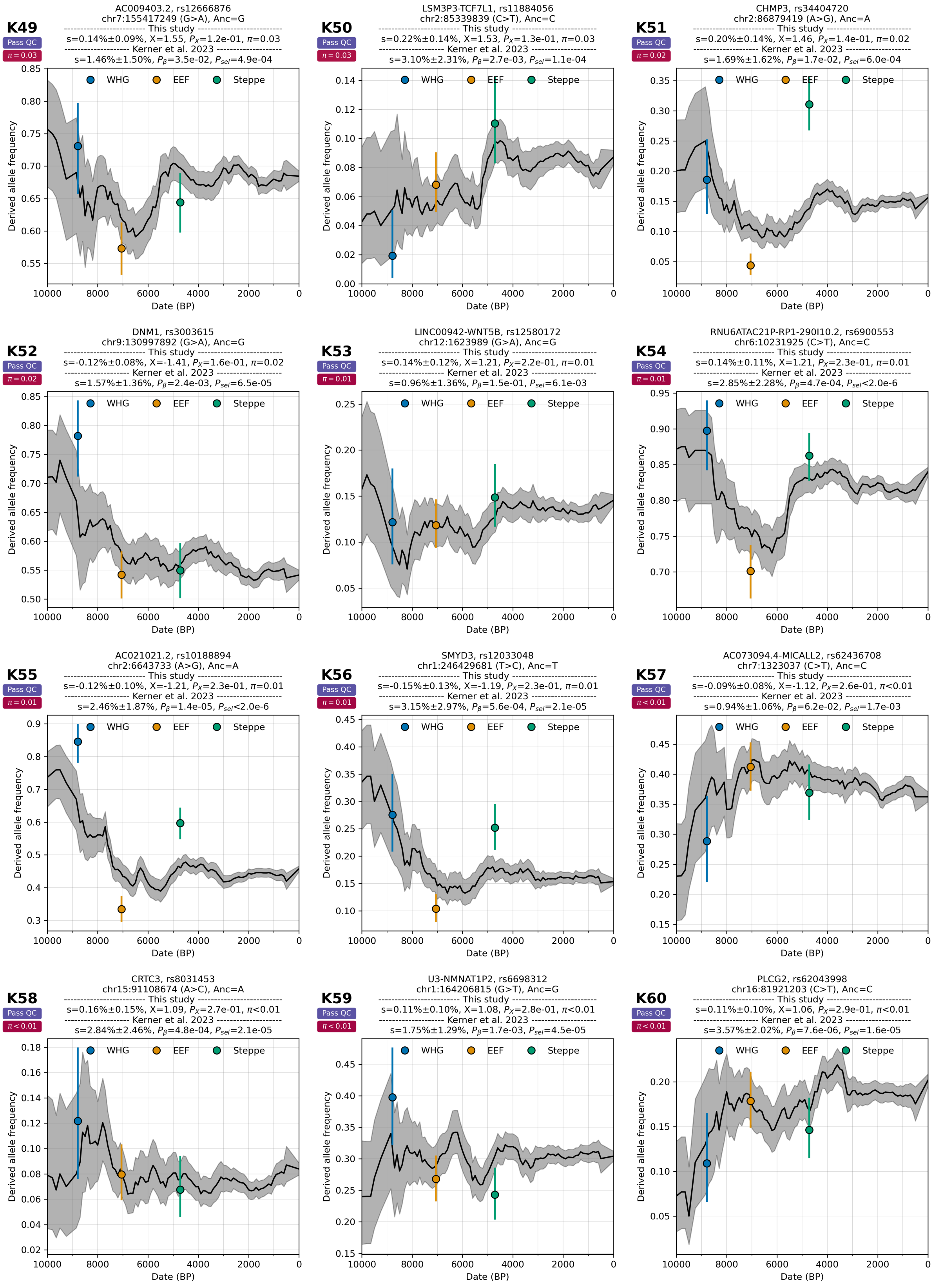

**Supplementary Figure S5.15:** Positive selection cases from Kerner et al. 2023 passing QC.

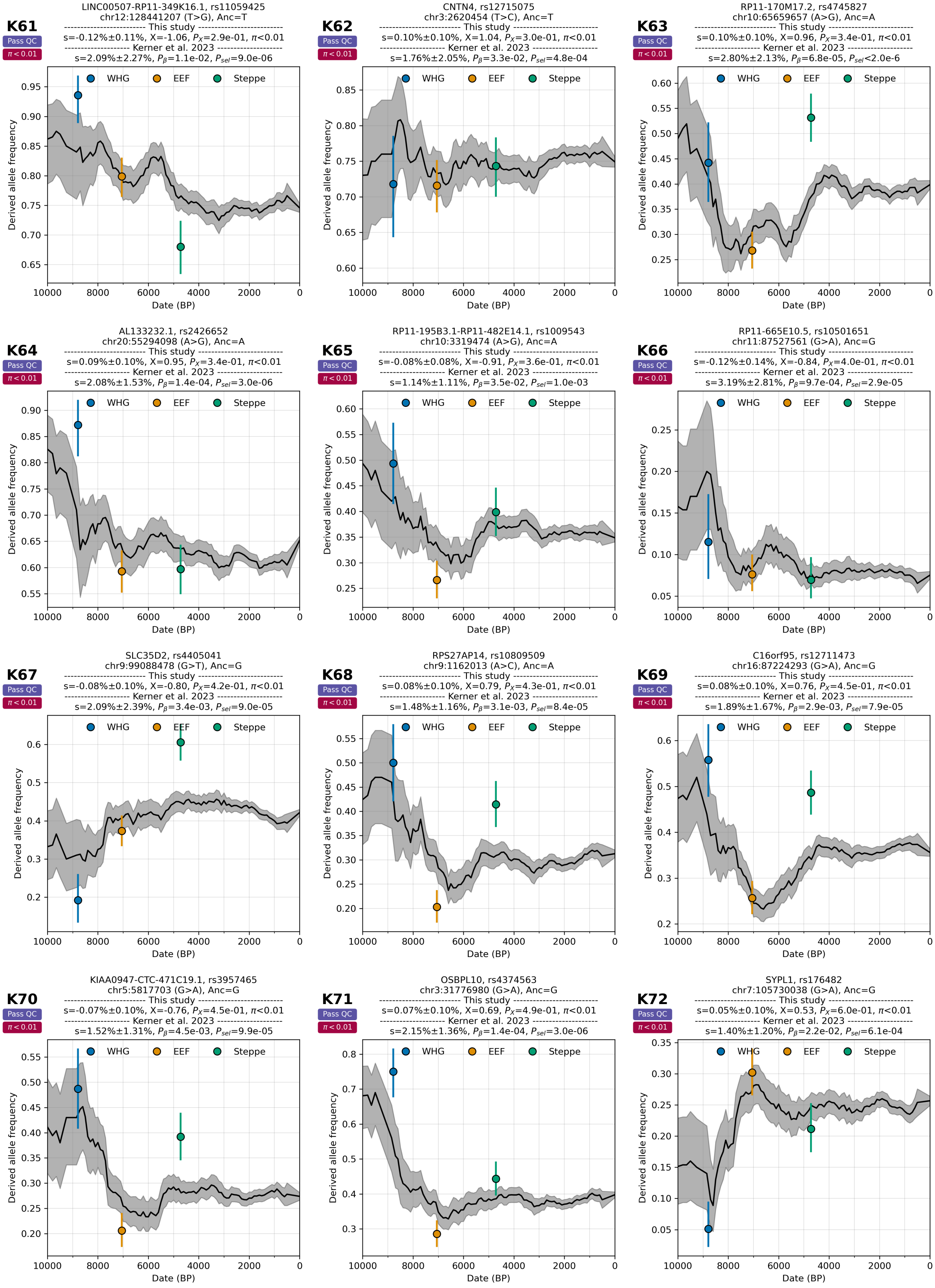

**Supplementary Figure S5.16:** Positive selection cases from Kerner et al. 2023 passing QC.

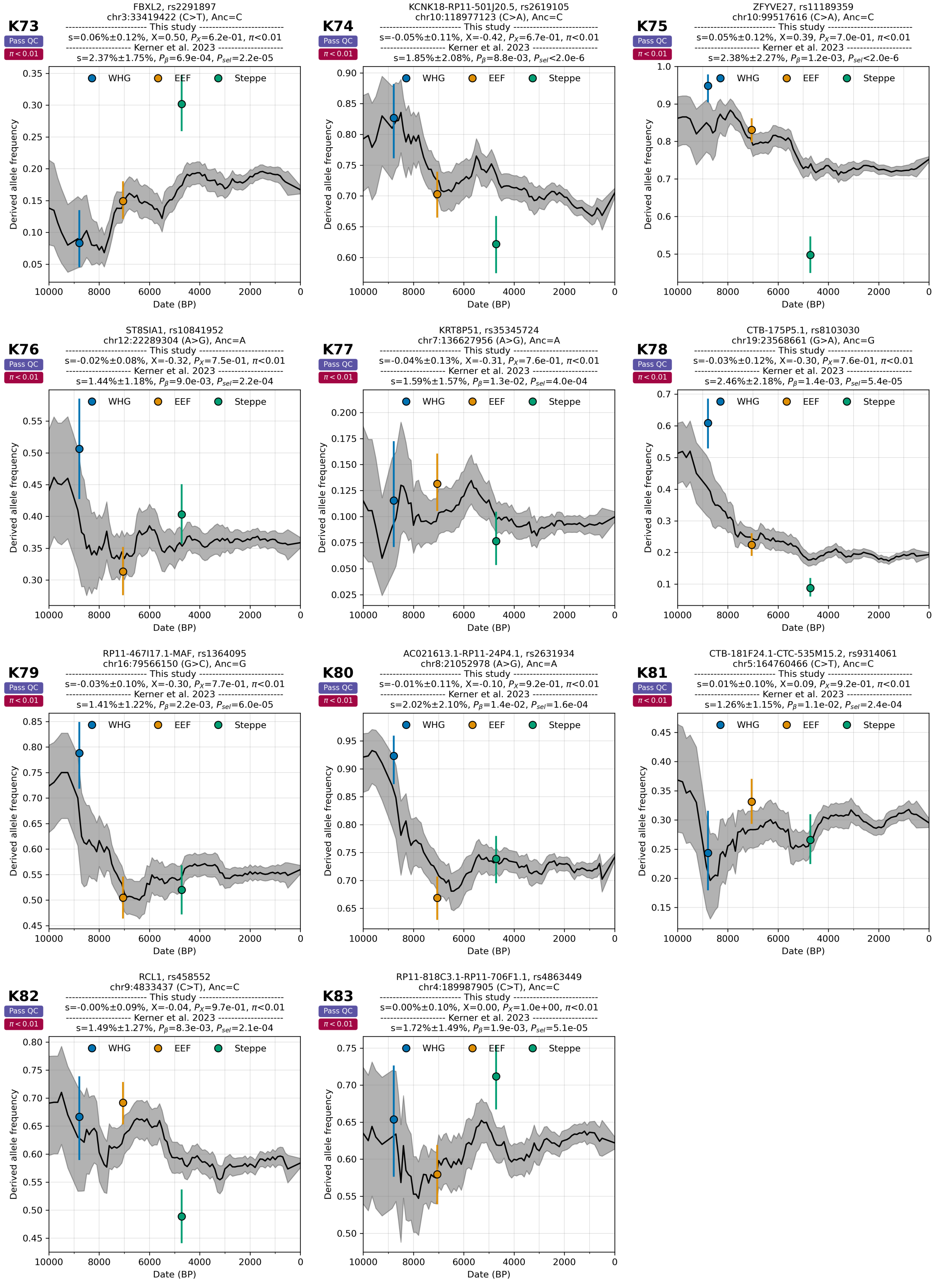

**Supplementary Figure S5.17:** Positive selection cases from Kerner et al. 2023 passing QC.

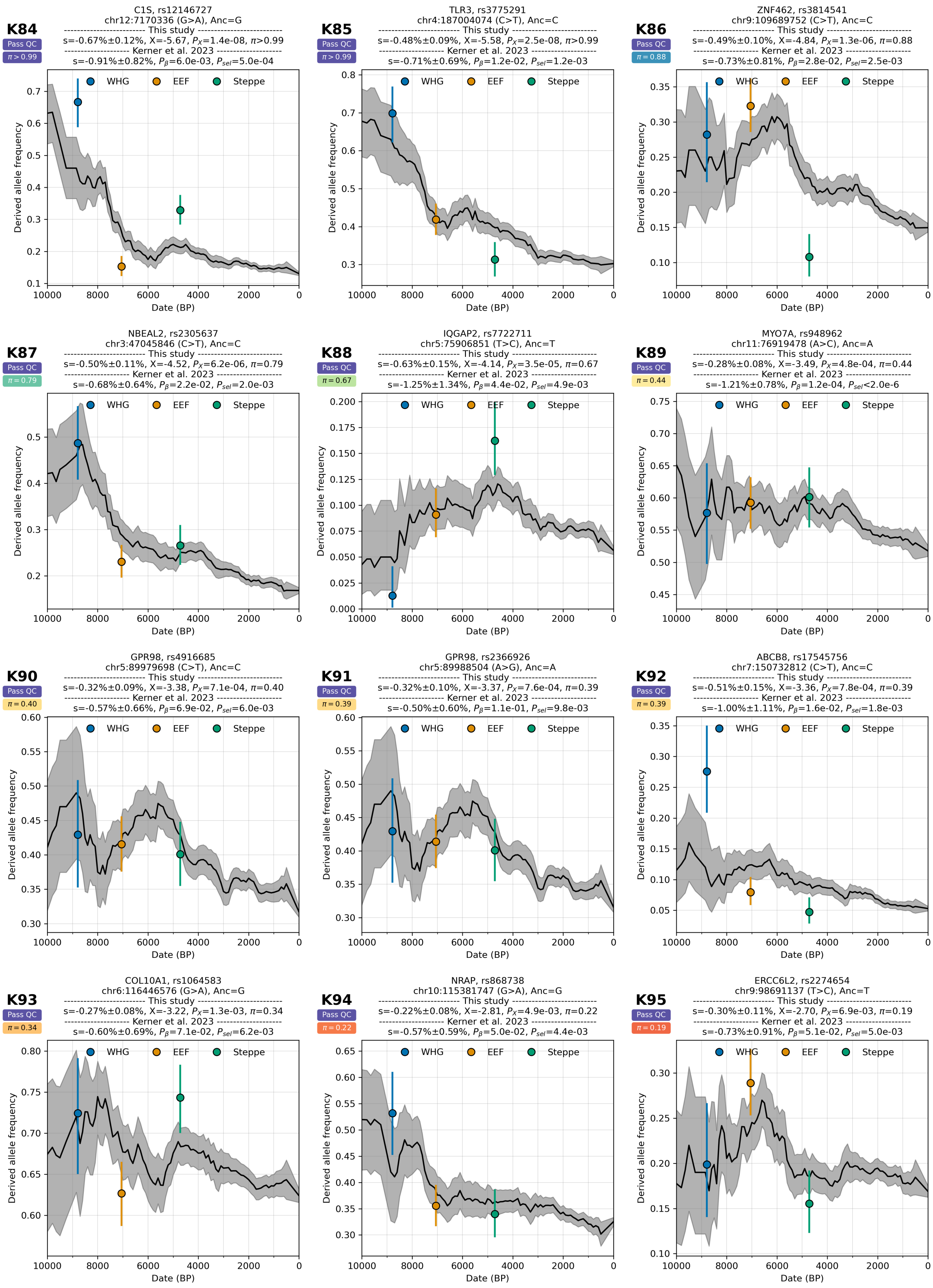

**Supplementary Figure S5.18:** Negative selection cases from Kerner et al. 2023 passing QC.

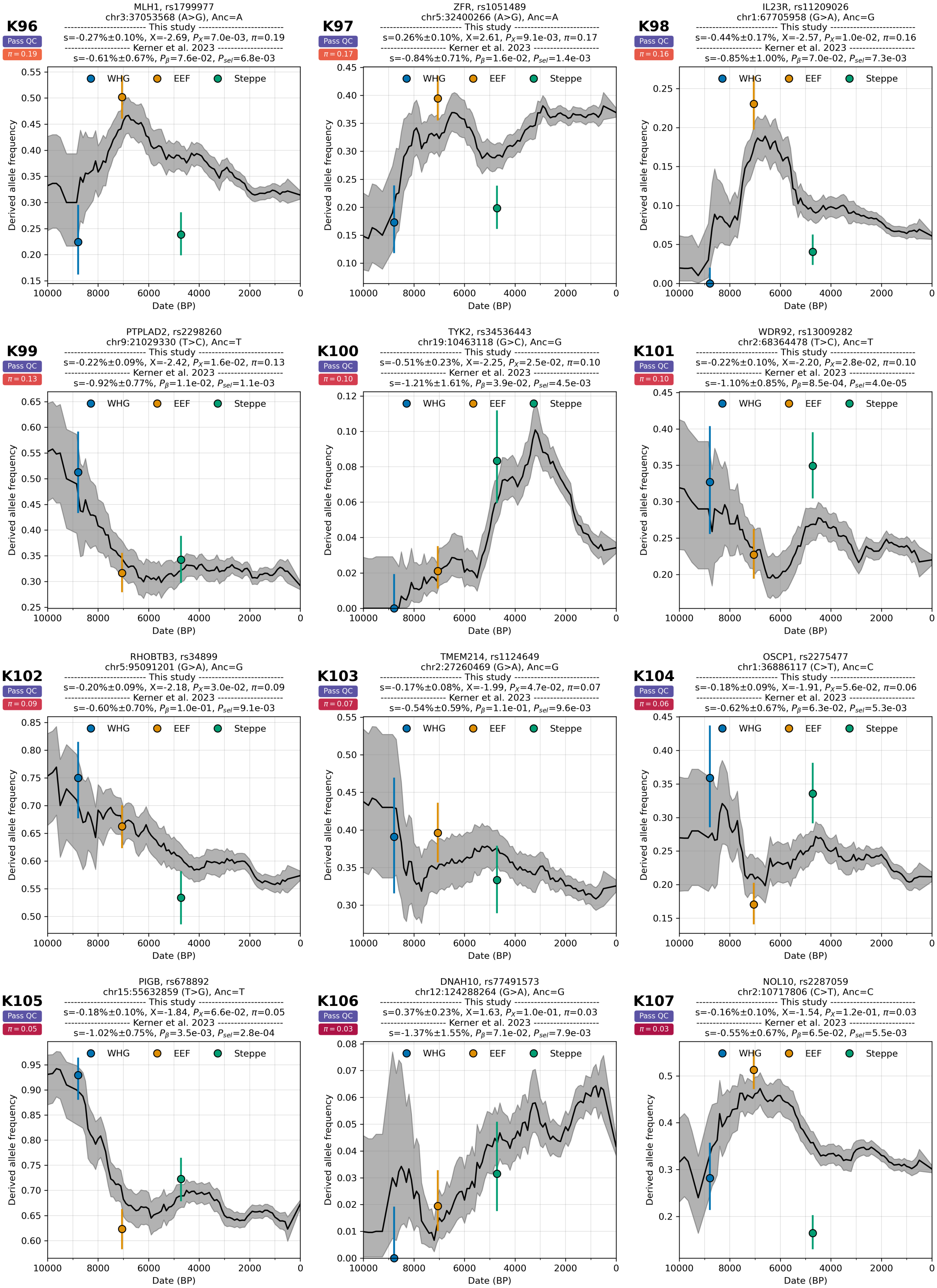

**Supplementary Figure S5.19:** Negative selection cases from Kerner et al. 2023 passing QC.

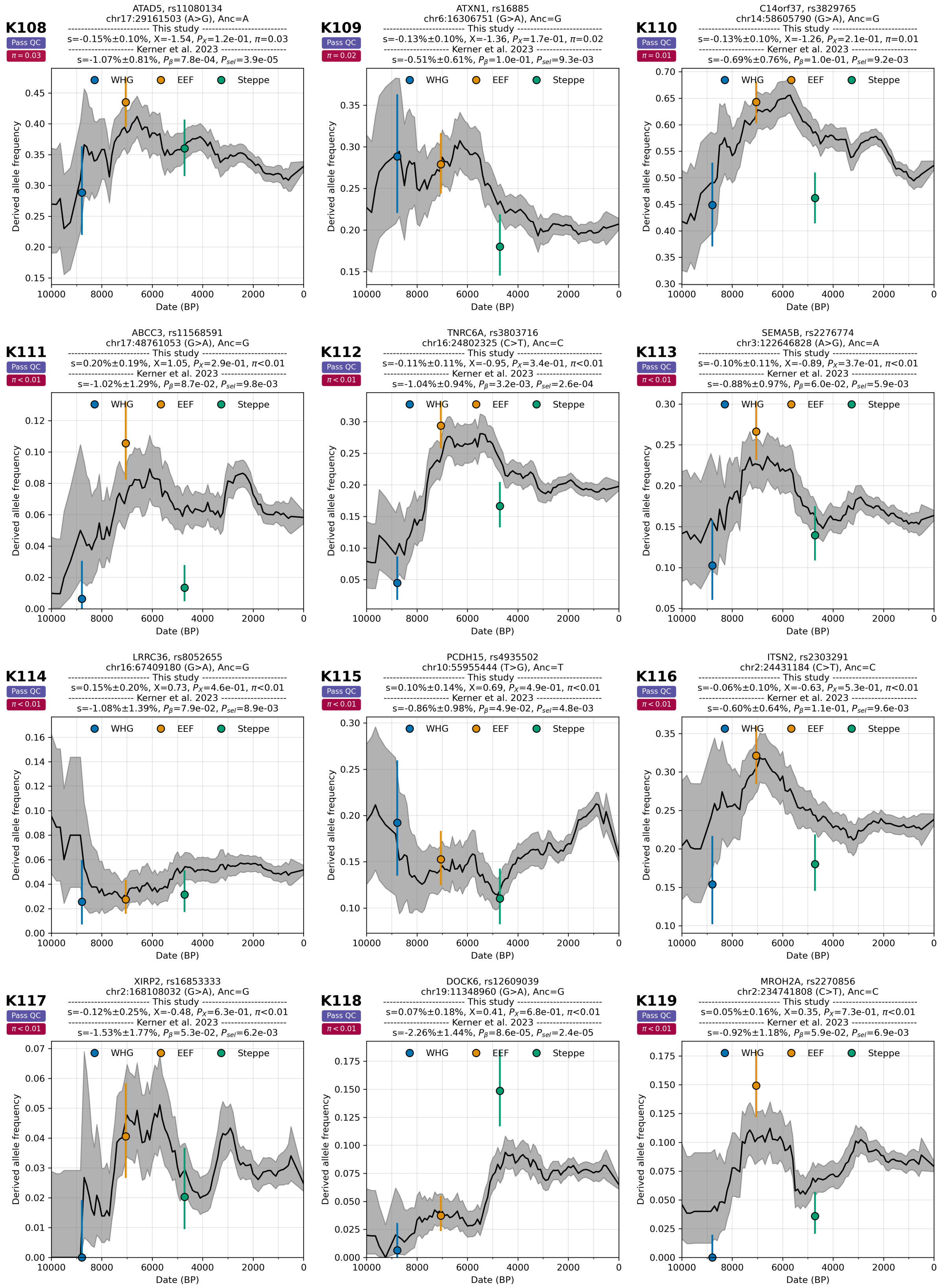

**Supplementary Figure S5.20:** Negative selection cases from Kerner et al. 2023 passing QC.

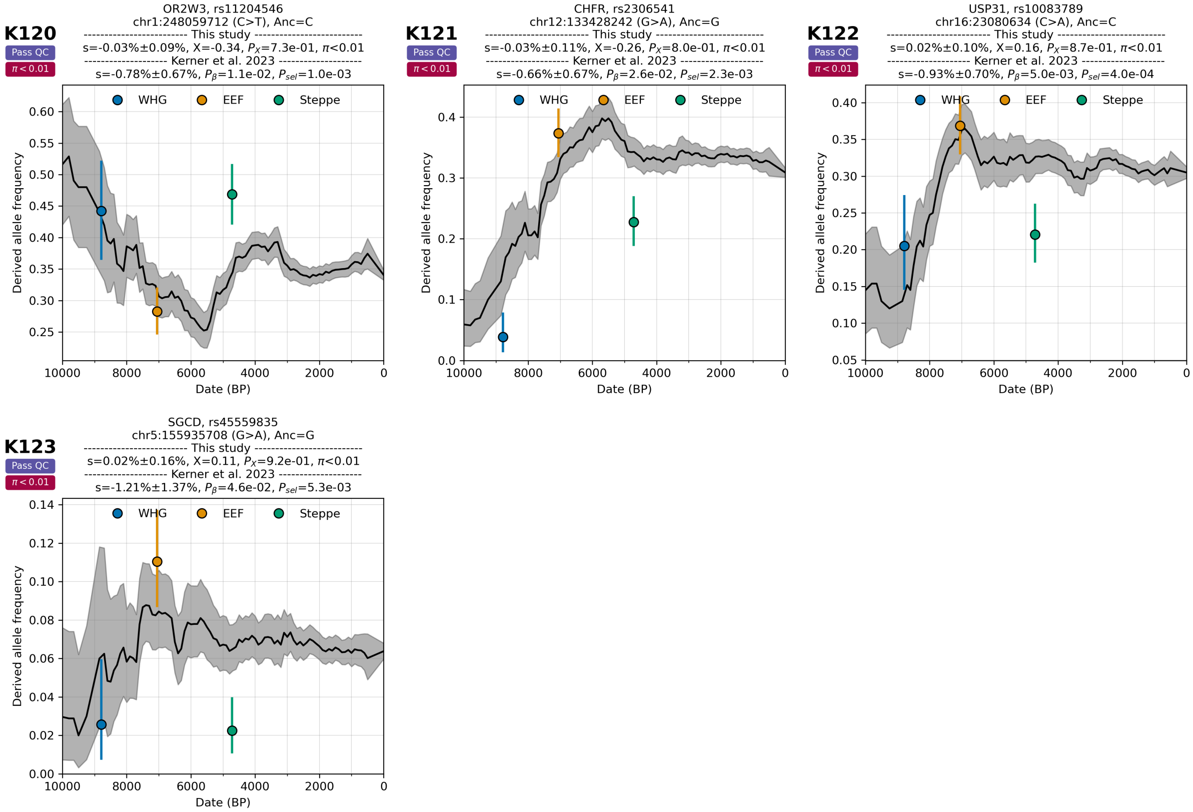

**Supplementary Figure S5.21:** Negative selection cases from Kerner et al. 2023 passing QC.

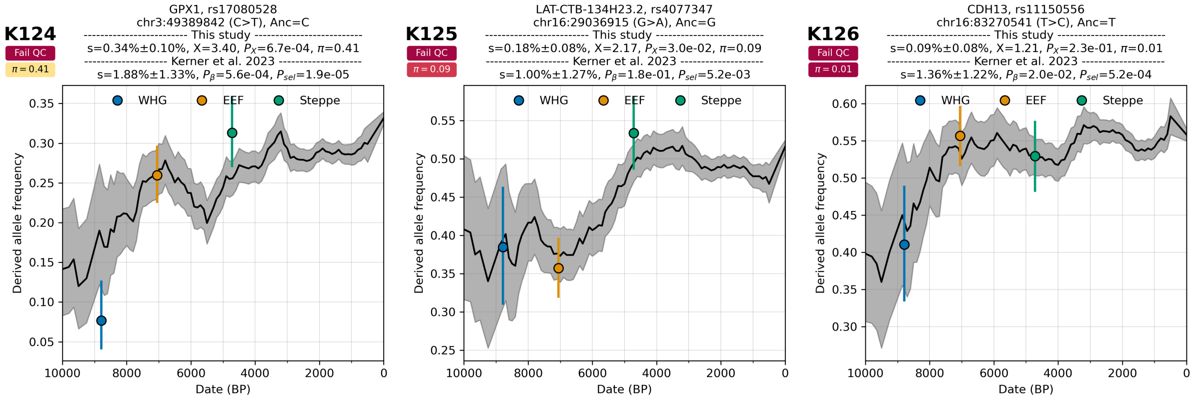

**Supplementary Figure S5.22:** Positive selection cases from Kerner et al. 2023 failing QC.

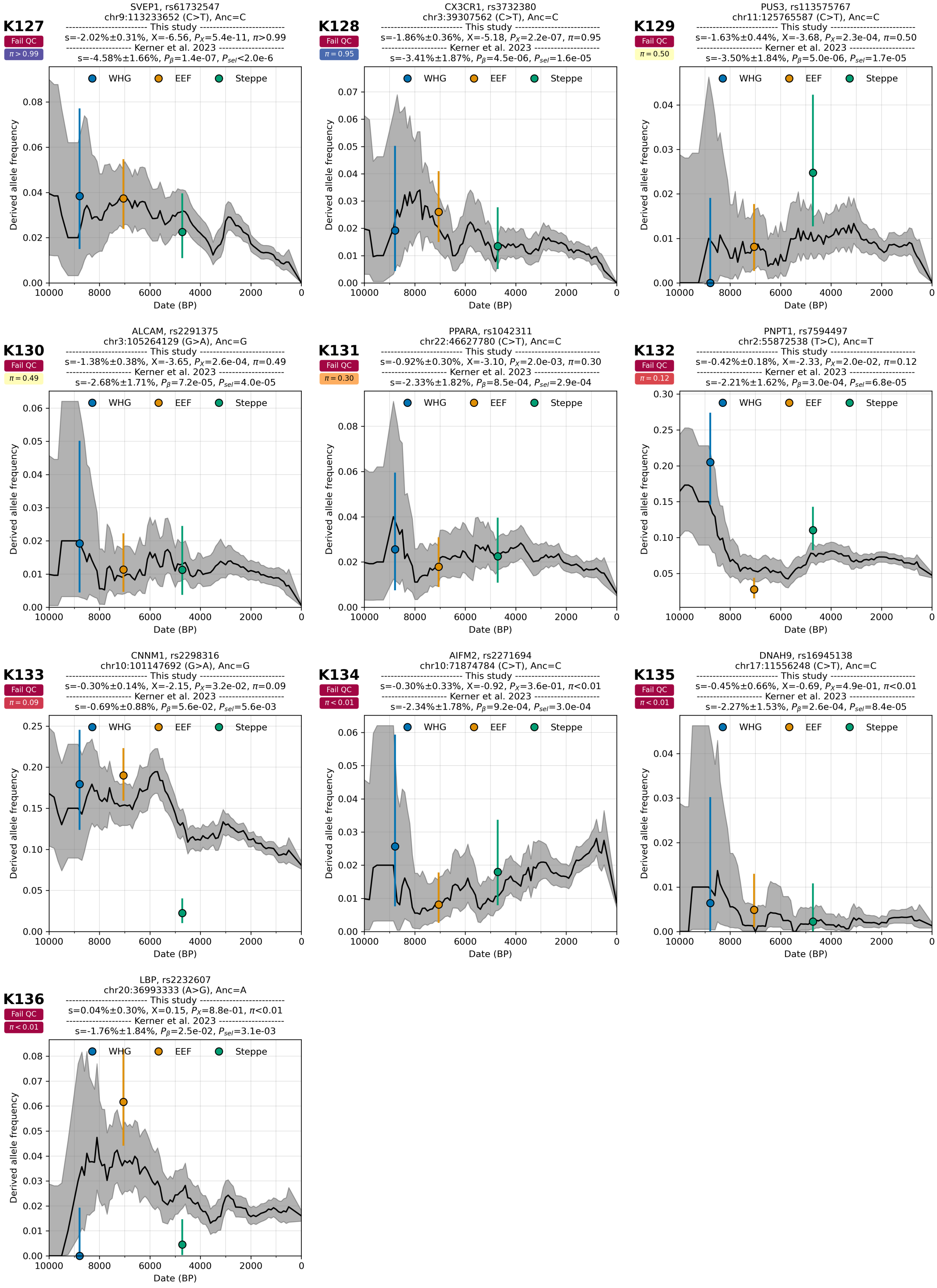

**Supplementary Figure S5.23:** Negative selection cases from Kerner et al. 2023 failing QC.

### Re-evaluation of results from Irving-Pease et al. 2024

We evaluated 21 selection signals from Figure 2a of Irving-Pease et al. 2024[11] and extracted summary statistics from Supplementary Table S2.1.4 of that study. That table includes five sets of estimated selection coefficients for different ancestry categories: pan-ancestry analysis (ALL), Western hunter-gatherers (WHG), Eastern hunter-gatherers (EHG), Caucasus hunter-gatherers (CHG), and Anatolian farmers (ANA). For each SNP, we picked $Z_{Top}$ as the Z score with the most significant value across these categories. In Supplementary Figure S5.24, we compare $Z_{Top}$ and X scores for the full time transect analysis. We evaluated five SNPs with the largest $Z_{Top}$ for each locus (Supplementary Figure S5.24). We picked these five SNPs for each locus because they are highlighted in the Extended Data Figures 1-10 and Supplementary Figures S56-S76, all from Irving-Pease et al. 2024. Re-evaluation of these SNPs is shown in our Supplementary Figures S5.25-S5.29.

In interpreting our re-evaluation of the results from Irving-Pease et al. 2024, it is important to be cognizant of the fact that for each locus, we selected the SNP with the most significant signal of selection in our study out of five candidate SNPs per locus proposed by Irving-Pease et al. While this approach lacks control for multiple testing and thus does not provide an entirely fair comparison with the other four studies re-evaluated here—it is expected to overestimate the replication rate in Irving-Pease et al. 2024 relative to those other studies—we followed this approach to maintain consistency with the approach of Irving-Pease et al. 2024.

Of the 21 candidate loci identified by Irving-Pease et al. 2024, our analysis found that 13 had at least one SNP among five candidates per locus with significant posterior probability ($\pi > 99\%$). These loci include *MCM6* (peak 3), *RNA5SP158* (peak 5), *SLC45A2* (peak 7), *IRF1* (peak 8), *SLC34A1* (peak 9), HLA (peak 10), *GATA4* (peak 11), *ABO* (peak 13), *FADS2* (peak 14), *ACAD10* (peak 15), *CYP1A1* (peak 16), *ARL17B* (peak 19), and *CENPM* (peak 21).

Additionally, *CCDC12* (peak 4, SNP 1, $\pi = 95\%$), *RAPGEFL1* (peak 18, SNP 2, $\pi = 76\%$), *WWP2* (peak 17, SNP 2, $\pi = 72\%$), and *LINC01104* (peak 2, SNP 2, $\pi = 65\%$) showed some evidence of selection on at least one of the five candidate SNPs per locus passing QC.

For the remaining four loci—*KRT18P51* (peak 6, SNP 4, $\pi = 40\%$), *RP11-415K20.1* (peak 1, SNP 1, $\pi = 28\%$), *CTD-2008O4.1* (peak 12, SNP 5, $\pi = 22\%$), and *CTC-258N23.3* (peak 20, SNP 4, $\pi = 2\%$)—there is no strong evidence of selection, as all five SNPs per locus that passed QC had posterior probabilities below 40%.

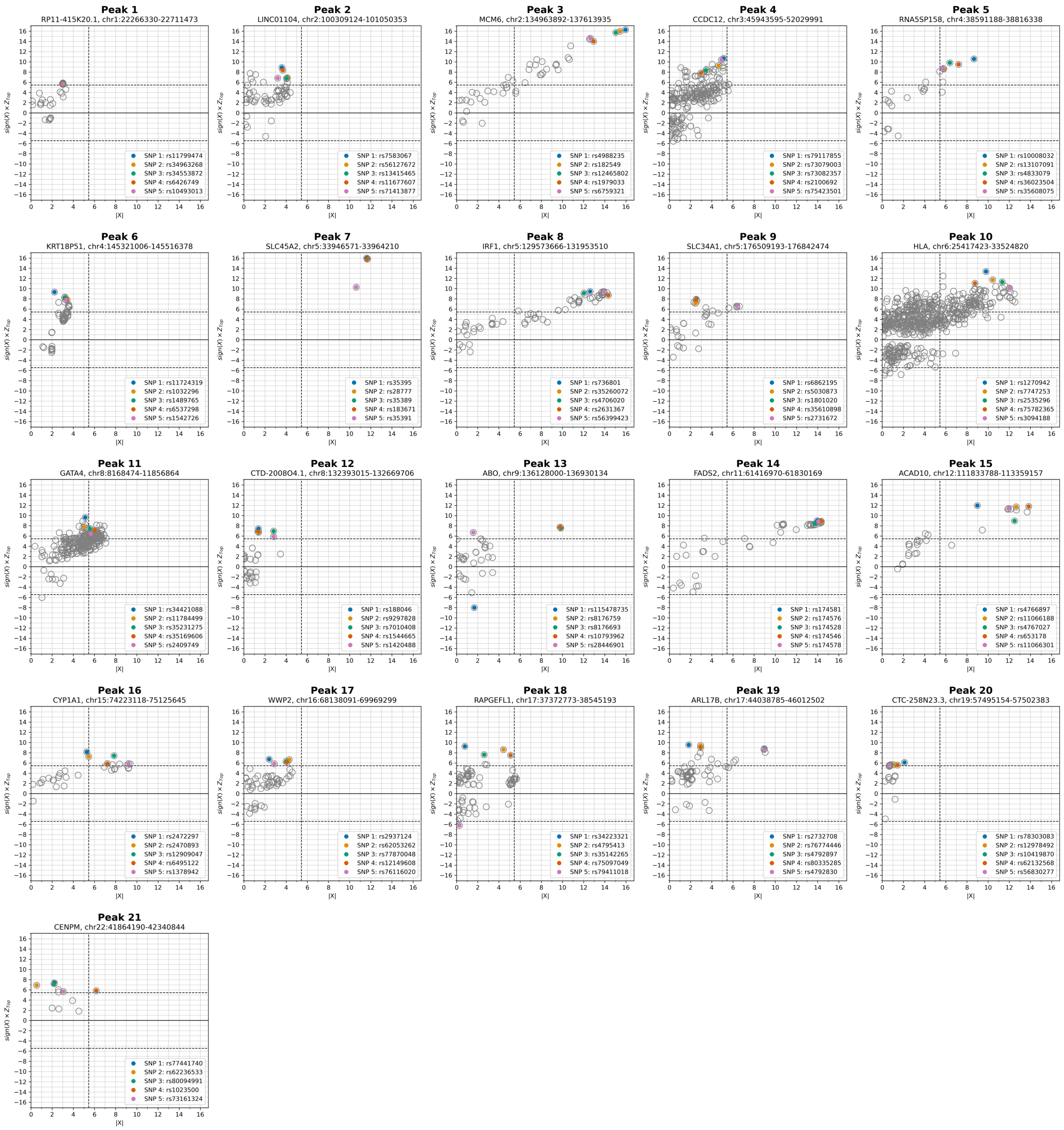

**Supplementary Figure S5.24:** Re-evaluating 21 candidate selective sweeps from Irving-Pease et al. 2024. Each marker represents a SNP, with the x-axis showing the absolute value of the X-score (|X|) from the current study. Irving-Pease et al. reported five sets of estimated selection coefficients for different ancestry categories: ALL, WHG, EHG, CHG, and ANA. For each SNP, we selected $Z_{Top}$ (y-axis), the most significant $Z$ score across these five categories. Then, we evaluated the five SNPs with the largest $Z_{Top}$ for each locus, represented by markers in different colors, as shown in the legend of each panel. We picked these five SNPs for each locus because they are highlighted in the Extended Data Figures 1-10 and Supplementary Figures S56-S76, all from Irving-Pease et al. 2024. The y-axis is polarized using the sign of the X-statistic.

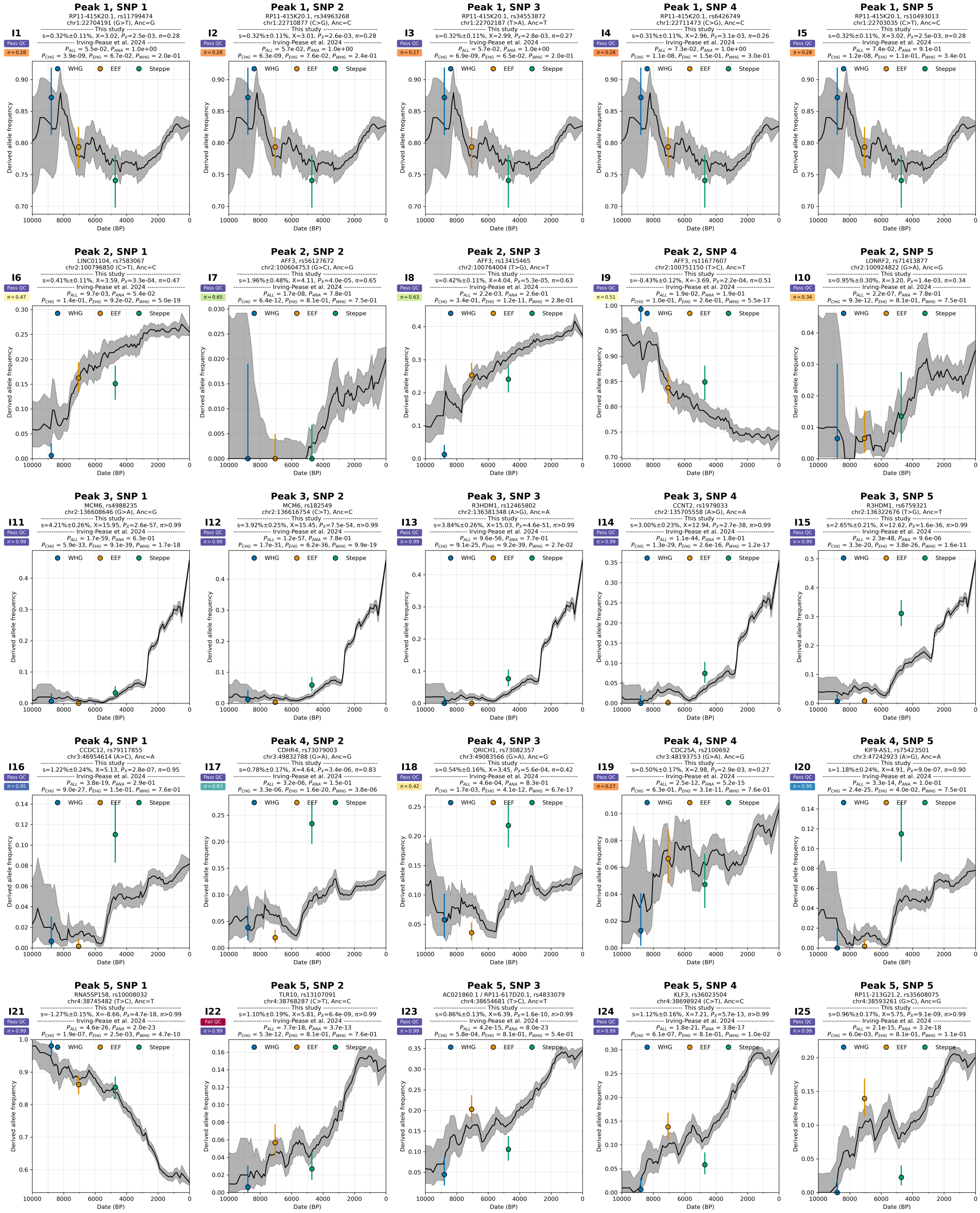

**Supplementary Figure S5.25:** Re-evaluating 21 candidate sweeps from Irving-Pease et al. 2024.

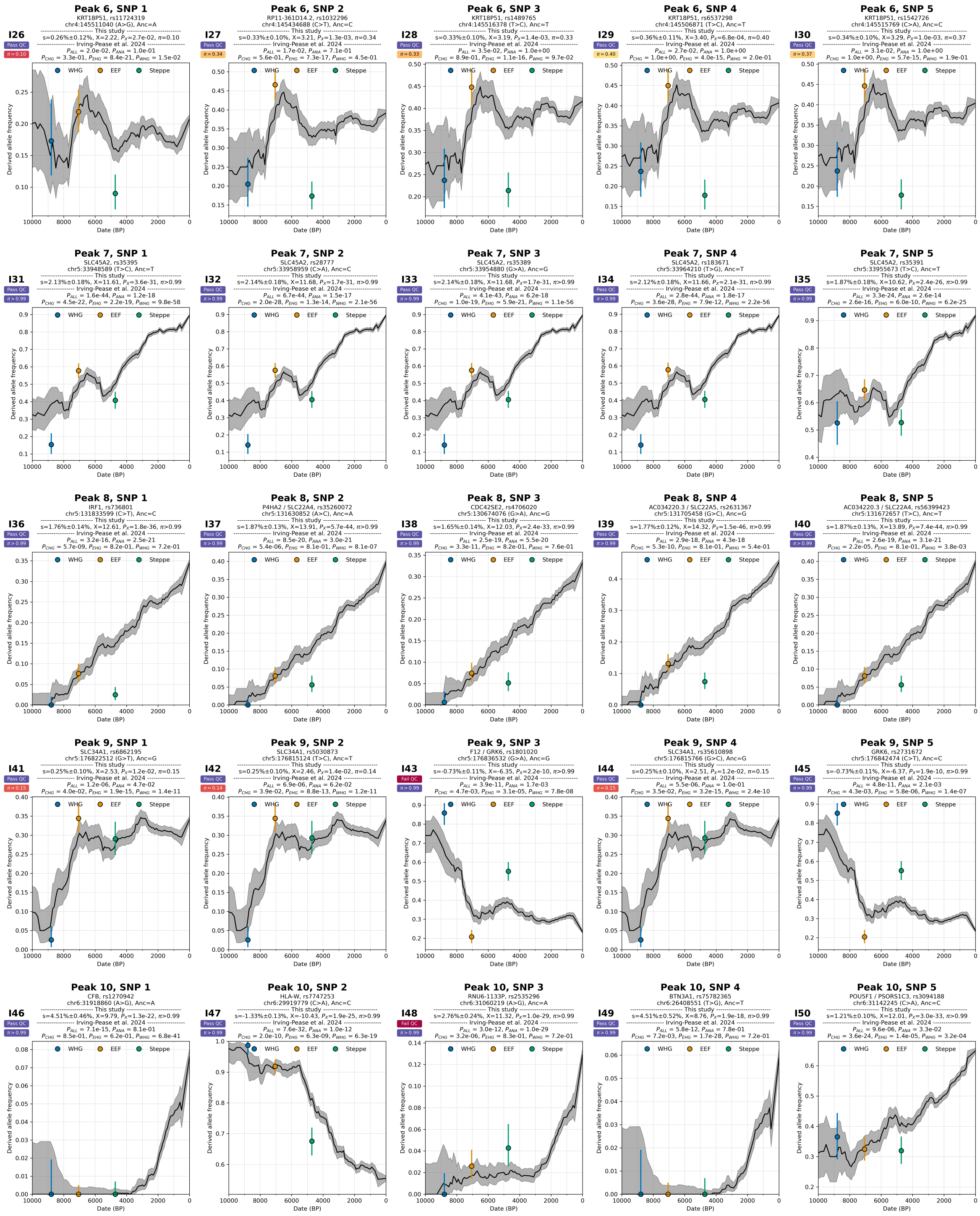

**Supplementary Figure S5.26:** Re-evaluating 21 candidate sweeps from Irving-Pease et al. 2024.

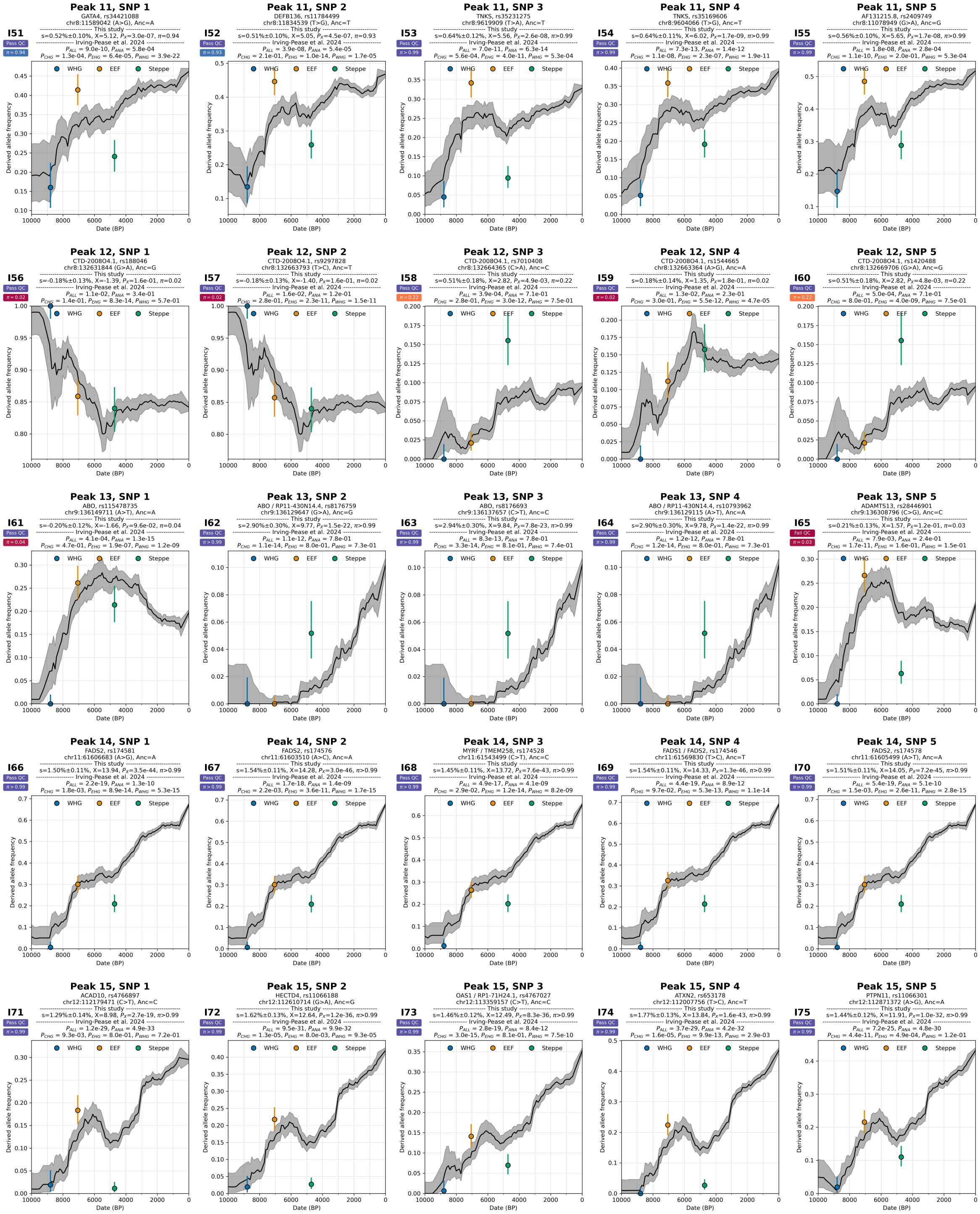

**Supplementary Figure S5.27:** Re-evaluating 21 candidate sweeps from Irving-Pease et al. 2024.

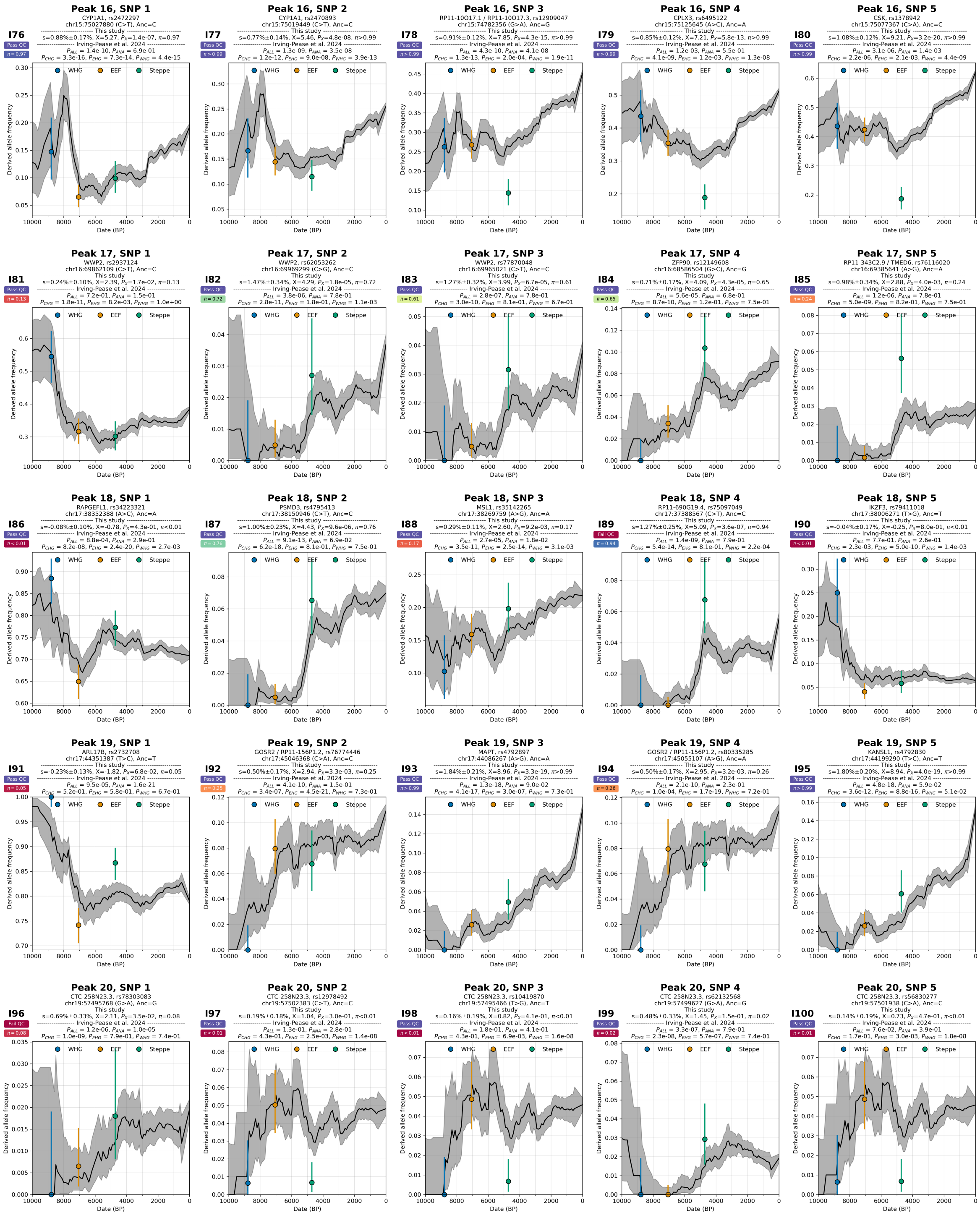

**Supplementary Figure S5.28:** Re-evaluating 21 candidate sweeps from Irving-Pease et al. 2024.

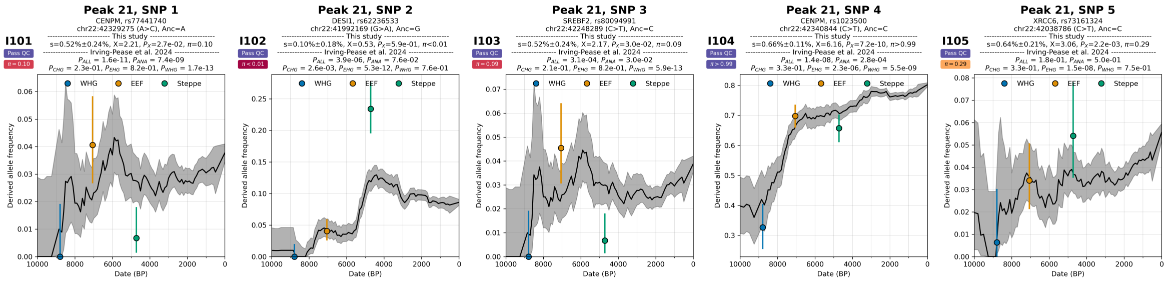

**Supplementary Figure S5.29:** Re-evaluating 21 candidate sweeps from Irving-Pease et al. 2024.

### Supplementary Information section 6

### A new picture of selection at the major risk factor for multiple sclerosis (MS)

Barrie et al. 2024[20] reported positive selection at the HLA-DRB1*15:01 allele tagged by the rs3135388 (G>A) variant, the strongest known genetic risk factor for multiple sclerosis (MS), and proposed that the elevated genetic risk for MS in Northern Europeans relative to other populations owes its origins at least in part due to the very high steppe pastoralist ancestry proportion in these populations. We confirm a strong signal of positive selection at this locus. However, we also identify features of the selection history at the locus missed by the previous study, and that together paint a qualitatively different story. The selection history was more complicated, and steppe ancestry was not in fact the main driver of the variant's frequency differences across Europe today.

First, we detect a period of strong negative selection from ~2000 years ago to the present ($s$ = -2.4%, $\pi$>99%) (Figure 3). This period of negative selection has had a primary influence on the frequency of this variant in Europeans, was missed in Barrie et al. 2024, and followed the period of positive selection from ~6000 to ~2000 years ago ($s$ = 4.0%, $\pi$>99%) that drove their finding.

Second, we show that the rise in frequency of this variant occurred initially in people without steppe ancestry living south of the Caucasus mountains, prior to the period of positive selection in Yamnaya steppe pastoralists around 5000 years ago that was the focus of the Barrie et al. 2024 study. We infer that the variant's frequency was around 10% (4.6%-18.2%; 95% confidence interval) 7000-5000 years ago south of the Caucasus mountains. Our finding of a first rise in frequency in association with Caucasus ancestry is entirely consistent with Figure 5c of Barrie et al. 2024 which infers that that the variant rose in frequency on a Caucasus ancestry background, one of the primary components of the ancestry of Yamnaya steppe pastoralists. However, our results go beyond that earlier study in showing that the rise in frequency actual likely to have occurred south of the Caucasus mountains (Supplementary Figures S6.2, S6.3), not in Eneolithic steppe hunter-gatherers in this period who also carried Caucasus ancestry.

Third, Barrie et al. 2024 observed that the frequency of the MS risk allele is highest among modern individuals in northern Europe with high steppe ancestry. They proposed that the steppe ancestry gradient, combined with environmental factors modulating genetic risk independent of genetics, address the long-standing debate regarding the north-south gradient in MS prevalence. However, our data reveal that the selection coefficient for this variant varies in both space and time, with the intensity of selection highest in northern populations compared to southern ones (Supplementary Figure S6.3). For example, for time transects older than 3500 years ago, the selection coefficient in the northern region (N; 14.5 ± 3.4% s.d.) is approximately three times higher than that for southwest region (SW; 5.1 ± 2.5% s.d.). The correlation between steppe ancestry and allele frequency in modern individuals is thus geographically confounded: the difference in selective pressure between the north and south after the spread of steppe pastoralists, not steppe ancestry, is consistent with driving the observed north-south gradient of this allele (Supplementary Figures S6.2, S6.3).

#### qpAdm modeling

We used qpAdm[21,22] (v1700) to estimate ancestry proportions for each sample.

We applied a 4-way model based on the Fernandes et al. 2020[23], using the following populations:

Right: Mbuti.DG, Ust_Ishim, ElMiron, Vestonice16, MA1, Israel_Natufian, Jordan_PPNB, Russia_Samara_EBA_Yamnaya, Morocco_LN.SG

Left: Turkey_N, EHG, WHG, Iran_GanjDareh_N

We also applied a 3-way model from Patterson et al. 2022[24], using the following populations:

Right: OldAfrica, WHGB, Russia_Afanasievo, Turkey_N

Left: WHGA, Balkan_N, OldSteppe

#### Inference of allele frequency in ancestral populations

We estimate allele frequencies in the ancestral populations by maximizing the likelihood, incorporating the estimated qpAdm ancestry proportions and imputed genotypes of all individuals.

Assume a 4-way model with unknown allele frequencies $p_A$, $p_B$, $p_C$ and $p_D$ for the source populations A, B, C, and D, respectively. For each individual $i$, the latent allele frequency $\hat{p}_i$ is calculated as a weighted average based on ancestry proportions $q_{A,i}$, $q_{B,i}$, $q_{C,i}$, and $q_{D,i}$:

$$\hat{p}_i = q_{A,i}p_A + q_{B,i}p_B + q_{C,i}p_C + q_{D,i}p_D$$

The likelihood of observing the genotype $g_i$ for each individual is then computed as:

$$P(g_i = 0 \mid \hat{p}_i) = (1 - \hat{p}_i)^2, \quad P(g_i = 1 \mid \hat{p}_i) = 2\hat{p}_i(1 - \hat{p}_i), \quad P(g_i = 2 \mid \hat{p}_i) = \hat{p}_i^2$$

The likelihood across all individuals is maximized to estimate allele frequencies in the sources.

$$L(\hat{p}_1, \hat{p}_2, \dots, \hat{p}_n) = \prod_{i=1}^{n} P(g_i \mid \hat{p}_i)$$

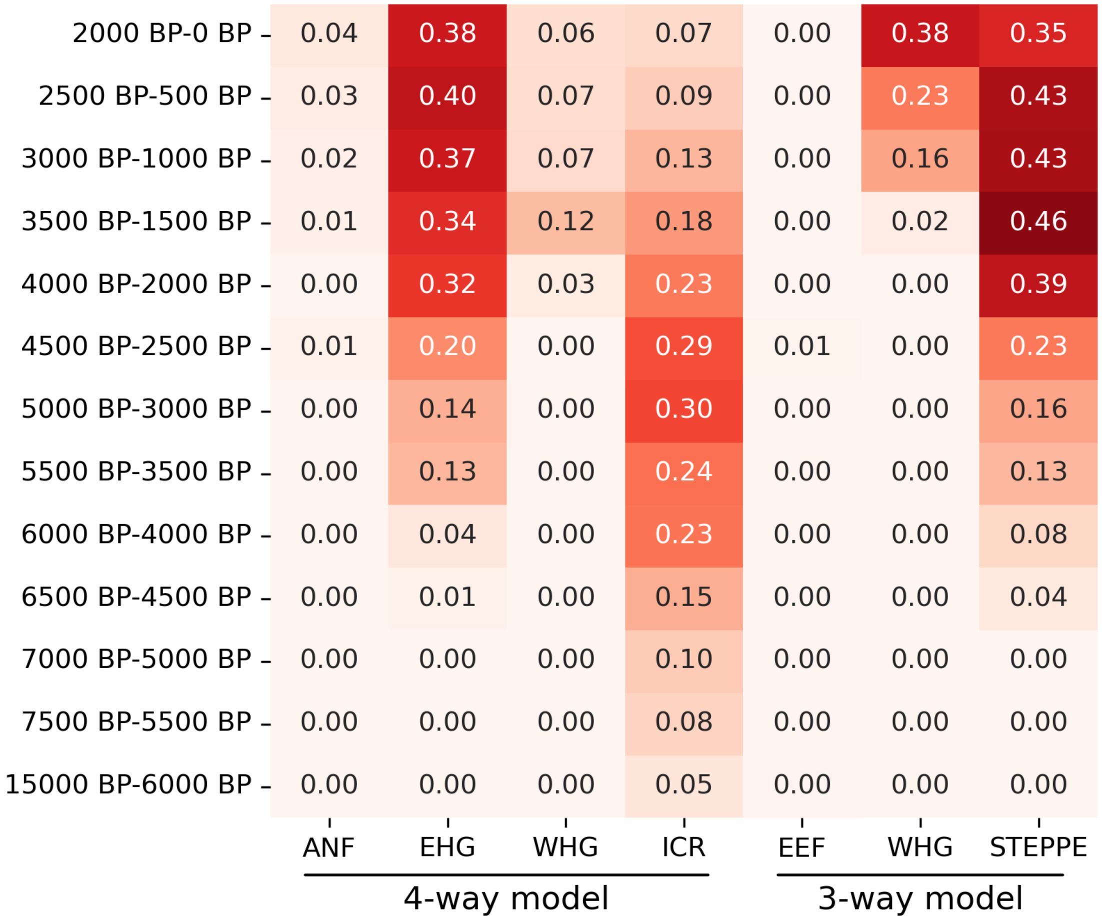

**Supplementary Figure S6.1:** Maximum likelihood estimation of allele frequencies of rs3135388 (G>A) across different time transects for ancestral populations using the 4-way and 3-way qpAdm ancestry models: ANF (Anatolian neolithic farmer), WHG (Western hunter-gatherer), ICR (Iranian/Caucasian-related), EHG (Eastern hunter-gatherer), EEF (Early European farmer), and STEPPE (Steppe pastoralists). For each qpAdm model, only individuals with a model P-value greater than 0.05 were used for the maximum likelihood estimation of allele frequencies.

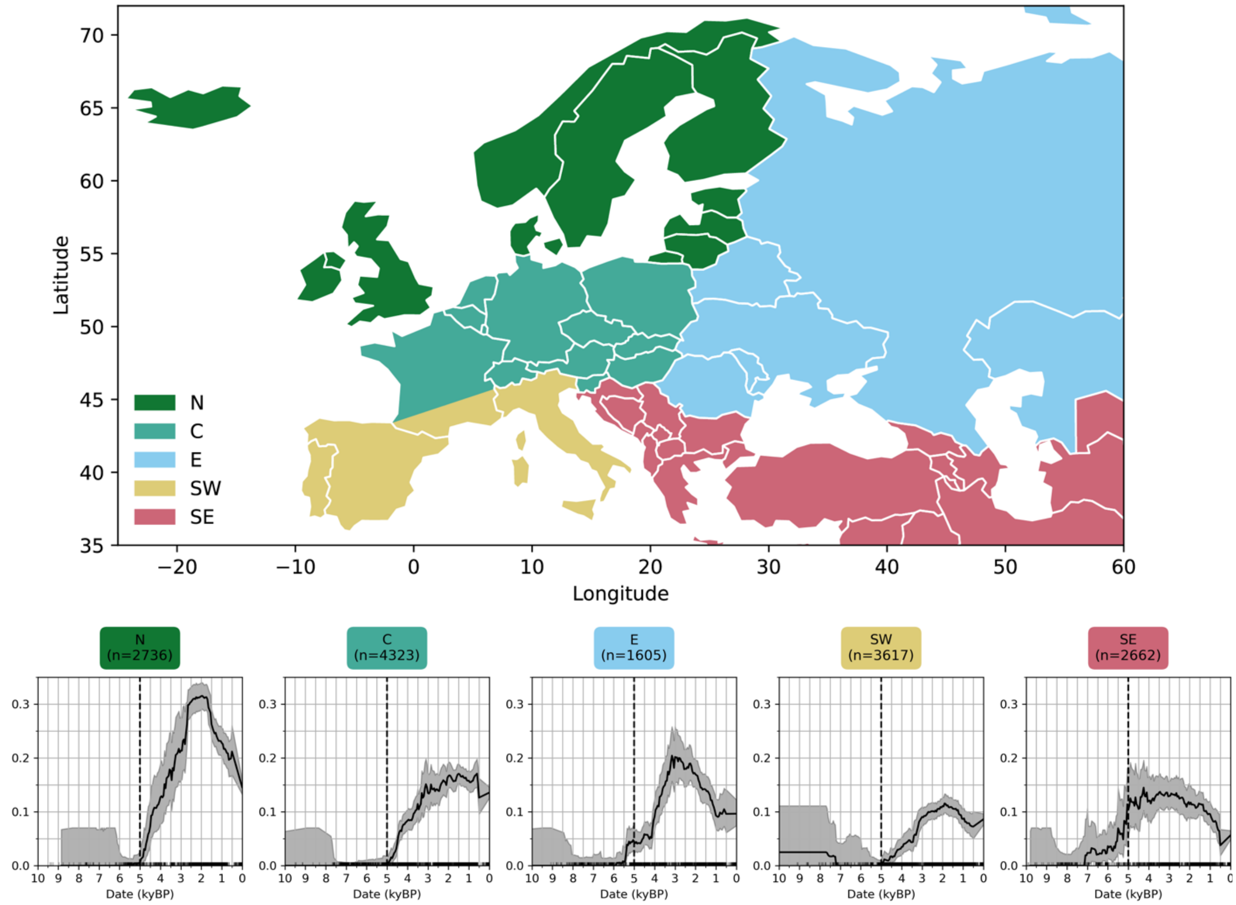

**Supplementary Figure S6.2:** Allele frequency trajectory of rs3135388 (G>A) stratified by five geographic regions: N (Northern), C (Central), E (Eastern), SW (Southwest), and SE (Southeast).

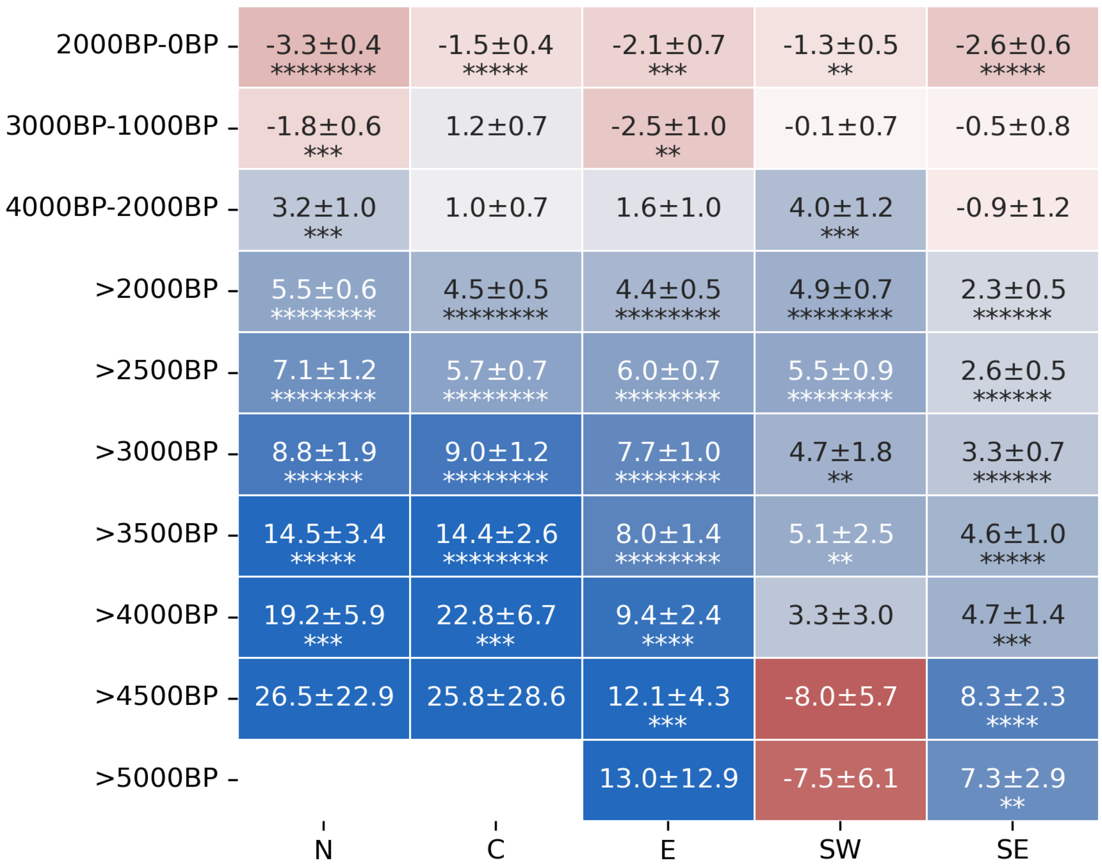

**Supplementary Figure S6.3:** Selection coefficient of rs3135388 (G>A) across different time transects, stratified by five geographic regions: N (Northern), C (Central), E (Eastern), SW (Southwest), and SE (Southeast). Selection coefficient values are presented as percentages in the format (s ± s.d.). Each star represents the level of significance. The number of stars (n stars) indicates that the P value is less than $0.5 \times 10^{-n}$, while the absence of a star means the P value is greater than 0.05.

### Supplementary Information section 7
### A fast GLMM implementation - PQLseq2

To analyze on the order of 15,000 individuals in a more time-efficient manner, we re-implemented the PQLseq[25] algorithm with several programming techniques to improve its computational speed. Specifically, we implemented the core algorithm exclusively in efficient C++ code. While the original implementation of PQLseq also used C++ for a portion of its algorithm, it alternates between R and C++ code in every iteration of the numerical optimization process, inducing repetitive type casting and object duplication. By fully implementing the core algorithm in C++, we circumvent those issues and significantly reduce the computational time. We refer to this faster re-implementation of PQLseq as PQLseq2. Furthermore, we implemented a specialized version of PQLseq2 with the heritability parameter fixed at 1, designed specifically for our analysis where the heritability parameter is expected to be close to 1. This adaptation further improves computational speed. We conducted a head-to-head comparison focusing on the computational speed and the parameter estimation between PQLseq and the faster version, PQLseq2. To do this, we randomly extracted the genotype data of 1,000 SNPs for 1,000 samples from the ancient DNA dataset, along with the sample dates and the genetic relatedness matrix. We then applied PQLseq, PQLseq2, and the specialized version of PQLseq2 to analyze the association between the allele frequency and sample dates. We analyzed one SNP at a time and compared the average computational time across the 1,000 SNPs. In the analysis, we found that PQLseq2 is approximately 15 times faster than PQLseq and the specialized version of PQLseq2 demonstrates a further 72% speed increase over PQLseq2 (Supplementary Figure S7.1A). Additionally, the estimates for all the parameters, including the fixed effect $\beta$ and its standard error, p-values for testing the presence of the fixed effect, heritability parameter $h^2$, and the total variance component $\sigma^2$, remained consistent between PQLseq and PQLseq2 (Supplementary Figure S7.1B-F). These results highlight both the efficiency and the accuracy of the PQLseq2 implementation. PQLseq2 is freely available and can be downloaded from https://github.com/zhengli09/PQLseq2.

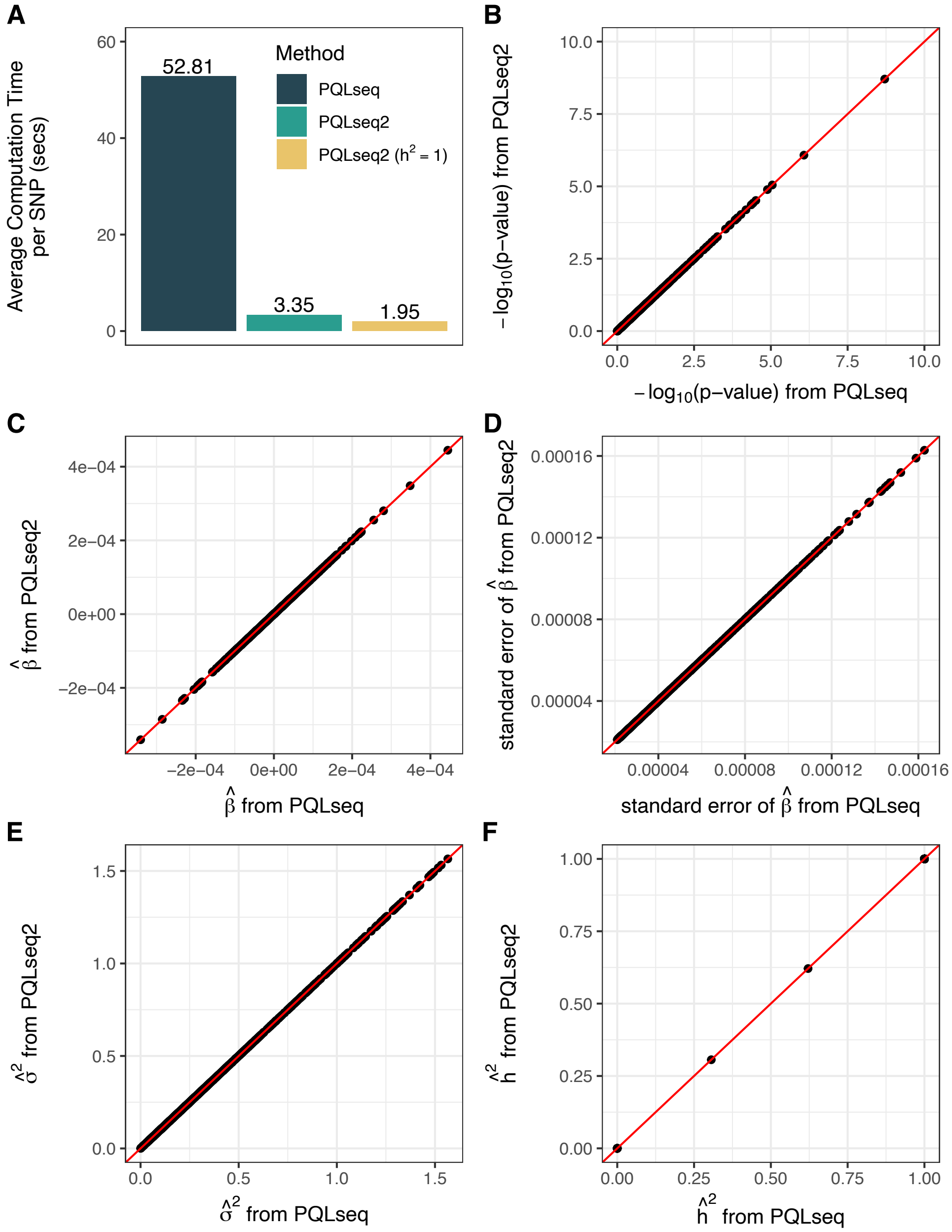

**Supplementary Figure S7.1:** Performance of a faster re-implementation of PQLseq. (**A**) Average computational time in seconds for the analysis of 1,000 randomly selected SNPs for 1,000 individuals from the ancient DNA data. Compared methods include the original implementation of PQLseq, the faster re-implementation PQLseq2, and a special version of PQLseq2 with the heritability parameter fixed at 1. (**B-F**) Scatter plots comparing the (**B**) p-values for testing the presence of the fixed effect $\beta$, (**C**) estimates of $\beta$, (**D**) standard errors of the $\beta$ estimates, (**E**) estimates of the total variance component $\sigma^2$, and (**F**) estimates of the heritability parameter $h^2$.

### References

1. Bulik-Sullivan, B. K. *et al.* LD Score regression distinguishes confounding from polygenicity in genome-wide association studies. *Nat. Genet.* **47**, 291–295 (2015).
2. Marchini, J., Howie, B., Myers, S., McVean, G. & Donnelly, P. A new multipoint method for genome-wide association studies by imputation of genotypes. *Nat. Genet.* **39**, 906–913 (2007).
3. Sousa da Mota, B. *et al.* Imputation of ancient human genomes. *Nat. Commun.* **14**, 3660 (2023).
4. Ronen, R. *et al.* Predicting Carriers of Ongoing Selective Sweeps without Knowledge of the Favored Allele. *PLoS Genet.* **11**, e1005527 (2015).
5. Akbari, A. *et al.* Identifying the Favored Mutation in a Positive Selective Sweep. *Nat. Methods* **15**, 279–282 (2018).
6. Fu, Y. X. Statistical properties of segregating sites. *Theor. Popul. Biol.* **48**, 172–197 (1995).
7. Murphy, D. A., Elyashiv, E., Amster, G. & Sella, G. Broad-scale variation in human genetic diversity levels is predicted by purifying selection on coding and non-coding elements. *eLife* **12**, e76065 (2023).
8. Mathieson, I. *et al.* Genome-wide patterns of selection in 230 ancient Eurasians. *Nature* **528**, 499–503 (2015).
9. Le, M. K. *et al.* 1,000 ancient genomes uncover 10,000 years of natural selection in Europe. *BioRxiv Prepr. Serv. Biol.* 2022.08.24.505188 (2022) doi:10.1101/2022.08.24.505188.
10. Kerner, G. *et al.* Genetic adaptation to pathogens and increased risk of inflammatory disorders in post-Neolithic Europe. *Cell Genomics* **3**, 100248 (2023).
11. Irving-Pease, E. K. *et al.* The selection landscape and genetic legacy of ancient Eurasians. *Nature* **625**, 312–320 (2024).

12. Field, Y. *et al.* Detection of human adaptation during the past 2000 years. *Science* **354**, 760–764 (2016).

13. Byrska-Bishop, M. *et al.* High-coverage whole-genome sequencing of the expanded 1000 Genomes Project cohort including 602 trios. *Cell* **185**, 3426-3440.e19 (2022).

14. Zhao, H. *et al.* CrossMap: a versatile tool for coordinate conversion between genome assemblies. *Bioinforma. Oxf. Engl.* **30**, 1006–1007 (2014).

15. Karczewski, K. J. *et al.* The mutational constraint spectrum quantified from variation in 141,456 humans. *Nature* **581**, 434–443 (2020).

16. Berg, J. J. *et al.* Reduced signal for polygenic adaptation of height in UK Biobank. *eLife* **8**, e39725 (2019).

17. Sohail, M. *et al.* Polygenic adaptation on height is overestimated due to uncorrected stratification in genome-wide association studies. *eLife* **8**, e39702 (2019).

18. Stern, A. J., Wilton, P. R. & Nielsen, R. An approximate full-likelihood method for inferring selection and allele frequency trajectories from DNA sequence data. *PLoS Genet.* **15**, e1008384 (2019).

19. Vaughn, A. H. & Nielsen, R. Fast and Accurate Estimation of Selection Coefficients and Allele Histories from Ancient and Modern DNA. *Mol. Biol. Evol.* **41**, msae156 (2024).

20. Barrie, W. *et al.* Elevated genetic risk for multiple sclerosis emerged in steppe pastoralist populations. *Nature* **625**, 321–328 (2024).

21. Haak, W. *et al.* Massive migration from the steppe was a source for Indo-European languages in Europe. *Nature* **522**, 207–211 (2015).

22. Harney, É., Patterson, N., Reich, D. & Wakeley, J. Assessing the performance of qpAdm: a statistical tool for studying population admixture. *Genetics* **217**, iyaa045 (2021).

23. Fernandes, D. M. *et al.* The Spread of Steppe and Iranian Related Ancestry in the Islands of the Western Mediterranean. *Nat. Ecol. Evol.* **4**, 334–345 (2020).

24. Patterson, N. *et al.* Large-Scale Migration into Britain During the Middle to Late Bronze Age. *Nature* **601**, 588–594 (2022).

25. Sun, S. *et al.* Heritability estimation and differential analysis of count data with generalized linear mixed models in genomic sequencing studies. *Bioinforma. Oxf. Engl.* **35**, 487–496 (2019).
